# Genome-wide determination of barriers to horizontal gene transfer

**DOI:** 10.1101/2022.06.29.498157

**Authors:** Christina L. Burch, Artur Romanchuk, Michael Kelly, Yingfang Wu, Corbin D. Jones

**Author notes:** Author for Correspondence: Christina Burch, Department of Biology, University of North Carolina, Chapel Hill, NC, USA.

## Abstract

Horizontal gene transfer (HGT) is a major contributor to bacterial genome evolution, generating phenotypic diversity, driving the expansion of protein families, and facilitating the evolution of new phenotypes, new metabolic pathways, and new species. Comparative studies of gene gain in bacteria suggest that the frequency with which individual genes successfully undergo HGT varies considerably and may be associated with the number of protein-protein interactions in which the gene participates—its connectivity. Two non-exclusive hypotheses have emerged to explain why transferability should decrease with connectivity: the Complexity Hypothesis (Jain, Rivera, & Lake, 1999) and the Balance Hypothesis (Papp, Pál, & Hurst, 2003). These hypotheses predict that the functional costs of HGT arise from a failure of divergent homologues to make normal protein-protein interactions or from gene mis-expression, respectively. Here we describe genome-wide assessments of these hypotheses in which we used 74 existing prokaryotic whole genome shotgun libraries to estimate rates of horizontal transfer of genes from taxonomically diverse prokaryotic donors into *E. coli*. We show that transferability declines as connectivity increases, but that this effect is driven primarily by the ribosomal genes. We also show that transferability declines as the divergence (% amino acid difference) between donor and recipient orthologs increases and that this effect of divergence increases with connectivity. We explain how these results, even the stronger effect of connectivity on the transferability of ribosomal compared to non-ribosomal genes, provide strong support for both the Balance and Complexity Hypotheses.

**Significance Statement:** Comparisons between prokaryotic genomes consistently show that genes with informational functions, e.g. in genome replication, transcription, and translation, have been subject to horizontal gene transfer between species more often than genes with operational functions, e.g. in metabolism and environmental sensing. In this study, we use a genome wide analysis of transferability data obtained from 74 genomes to provide the first experimental evidence that this pattern results from differences between informational and operational genes in the number of other proteins with which they interact, i.e., their connectivity, rather than from their functional differences. The importance of our exceptionally large dataset to the detection of connectivity effects on transferability explains why past experimental studies failed to replicate the consistent finding from comparative genomic studies.

## Introduction

Horizontal gene transfer (HGT) is a major contributor to bacterial genome evolution (Arnold et al. 2022; Brito 2021; Skippington & Ragan 2011), contributing 10 - 20% of the protein coding genes to most bacterial genomes (Lawrence & Ochman 1998; Nakamura et al. 2004; Soucy et al. 2015). HGT promotes diversity (Baltrus et al. 2011; Guttman 1997; Polz et al. 2013), facilitating the evolution of novel phenotypes (Moran & Jarvik 2010), metabolic pathways (Soyer & Creevey 2010), and species (Schaack et al. 2010). Numerous instances of rapid adaptation have been attributed to HGT (e.g., Lozupone et al. 2008; Dhillon et al. 2015; Arnold et al. 2020; Frazão et al. 2019; Woods et al. 2020; reviewed in Arnold et al. 2022).

Comparative studies reveal that the frequency of HGT varies among genes and among pairs of donor and recipient species in predictable ways (Nakamura et al. 2004; Pál et al. 2005; Soucy et al. 2015). For instance, transferability is observed to depend on gene function (Nakamura et al. 2004; Rivera et al. 1998), connectivity (i.e., the number of protein-protein interactions) (Cohen et al. 2011; Lercher & Pál 2008), and the divergence between donor and recipient genomes (Baltrus 2013; Soucy et al. 2015; Tuller et al. 2011).

Two non-exclusive hypotheses have been proposed to explain why the fitness cost of gene transfer increases with connectivity: the Balance Hypothesis (Papp et al. 2003) and the Complexity Hypothesis (Jain et al. 1999). These hypotheses predict that the fitness costs of HGT arise from gene mis-regulation (Balance Hypothesis) or from the failure of transferred orthologs to engage in normal protein-protein interactions (Complexity Hypothesis). The central predictions of these hypotheses are illustrated in fig. 1. Whereas costs associated with gene misregulation are expected regardless of the divergence between the resident and transferred orthologs (figs. 1A and 1C), costs associated with protein-protein interaction failure are expected to increase in frequency with divergence (figs. 1B and 1D). In both hypotheses, the average magnitude of the fitness cost is expected to increase with connectivity.

**Figure 1.**
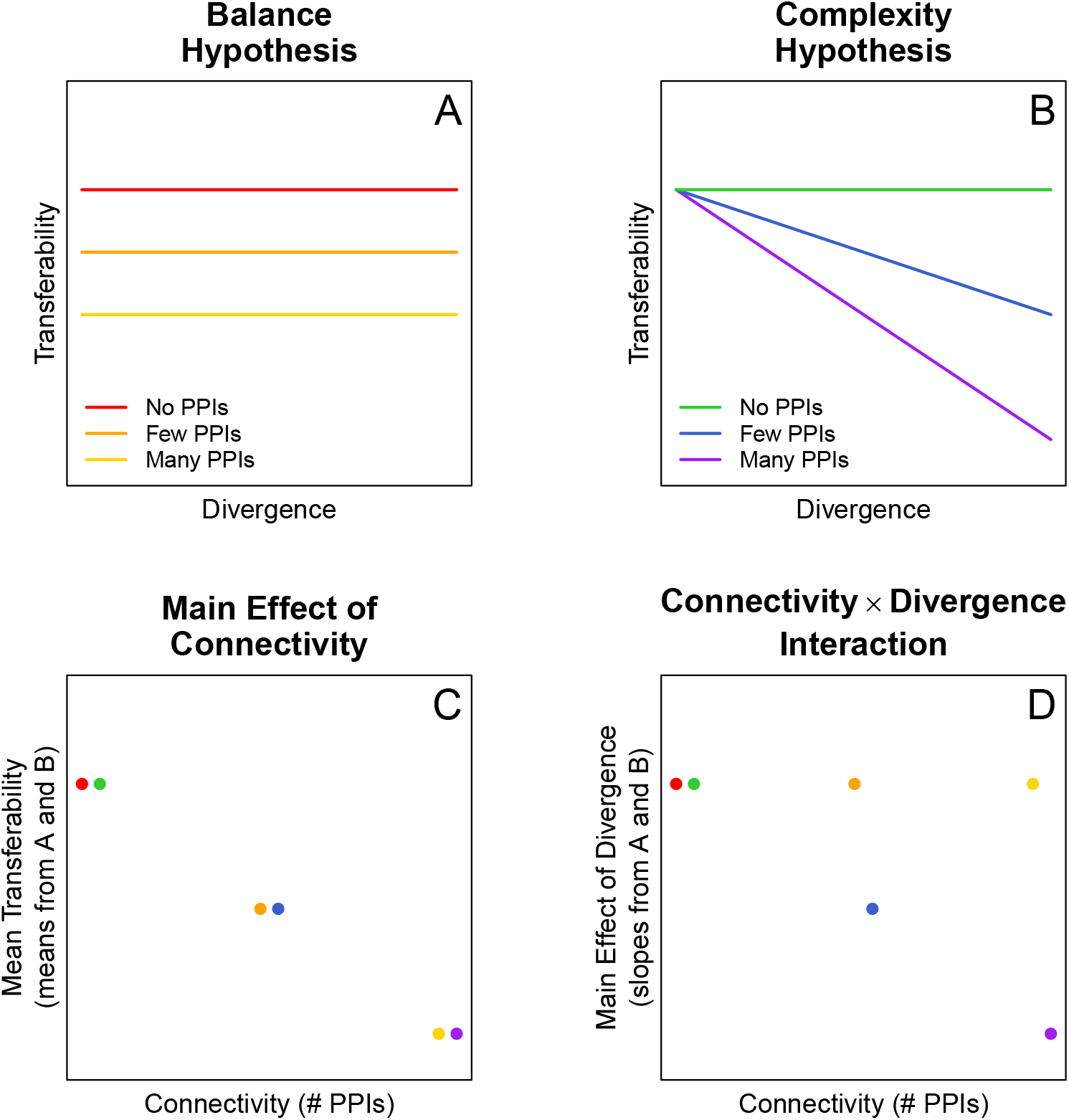
Central predictions of the Complexity and Balance Hypotheses. The Balance Hypothesis (A) posits that transferred genes with resident orthologs may cause an imbalance in the expression of genes that participate in multi-protein complexes and must be produced in stoichiometric amounts. The cost of expression imbalance is predicted to increase with the number of protein-protein interactions (PPIs), but not with divergence. As a result, genes that engage in PPIs (orange and yellow lines) are less transferable than genes that lack PPIs (red line). The Complexity Hypothesis (B) posits that transferred genes may interfere in the normal protein-protein interactions (PPIs) of the resident ortholog, and that the probability of interference increases with the divergence between the resident and transferred genes. As a result, transferability decreases with divergence, but only for genes that engage in PPIs (blue and purple lines). Two testable predictions result from these hypotheses. The central prediction of the Balance Hypothesis (C) is that the transferability of a particular gene, averaged over all donor orthologs, should decrease as the connectivity of the gene increases. In statistical lingo, the Balance Hypothesis predicts a *main effect of connectivity*. Points in (C) are mean values of lines of the same color in (A) and (B), illustrating that this prediction does not distinguish between the Balance and Complexity Hypotheses. The central prediction of the Complexity Hypothesis (D) is that the effect of divergence on transferability (i.e., the *main effect of divergence*) should become more negative as connectivity increases. In statistical lingo, the Complexity Hypothesis predicts a *connectivity × divergence interaction*. Points in (D) are slopes of lines of the same color in (A) and (B), illustrating that only the Complexity Hypothesis predicts a connectivity × divergence interaction, shown here as a decreasing main effect of divergence from green to blue to purple. The Balance Hypothesis makes no such prediction: the main effect of divergence does not differ between yellow, orange, and red.

Because comparative data often rely on the existence of sequence divergence to detect HGT, comparative data can document only cases where gene transfer had the potential to affect both gene regulation and protein-protein interactions. Thus, experimental data are needed to distinguish between these two effects. To date, experimental data have confirmed that the fitness costs of HGT vary among genes (Acar Kirit et al. 2020; Knöppel et al. 2014; Sorek et al. 2007) and depend on the divergence between donor and recipient genomes (Sorek et al. 2007). However, previous experimental tests have failed to detect an effect of connectivity on transferability (Acar Kirit et al. 2020).

Here we use a genome-wide quantitative approach to provide a direct test of the Complexity and Balance Hypotheses. Following (Sorek et al. 2007), we analyze the data generated during whole genome shotgun sequencing of 75 bacterial and 4 archaeal genomes. In the shotgun approach, random genome fragments were sequenced only after they were successfully cloned into a plasmid and transformed (i.e. transferred) into *E. coli*, thus genes that imposed fitness costs may have been underrepresented in the shotgun libraries. Sorek et al. (2007) investigated qualitative differences in gene representation (presence or absence) in the shotguns to explain the inability of some specific genes from some specific species to transfer into *E. coli*. We build on their analysis by investigating quantitative differences in gene representation in the shotgun sequencing data, to test general hypotheses for differences in transferability among genes and genomes (as in Knöppel et al. 2014; Acar Kirit et al. 2020). We test the Complexity and Balance Hypotheses by investigating the effects of connectivity and divergence on the number of times individual genes, in their entirety, were successfully cloned, transformed, and sequenced in the whole genome shotguns. The strong statistical power of our genome-wide quantitative approach enabled detection of both a main effect of connectivity and of an interaction effect between connectivity and divergence on the transferability (representation in the shotgun data) of individual genes, confirming the central predictions of the Balance and Complexity Hypotheses, respectively (fig. 1).

## Results

We acquired the shotgun library sequences of 74 prokaryotic species (70 bacteria and 4 archaea, described in Supplementary Table 1) from the NCBI Trace Archive, following (Sorek et al. 2007). We then calculated the coverage of each coding sequence in each shotgun library as the number of plasmid inserts in the library that contained the gene in its entirety (as described in *Methods: Gene coverage in the shotguns*). A visual examination of the coverage variation among genes within individual shotgun libraries (fig. 2 and Supplementary fig. S1) revealed two general patterns: 1) long genes are less well covered than short genes in all shotguns, and 2) genes near the origin of replication are better covered than genes near the terminus of replication, but to different degrees in different shotguns. These patterns suggested strong effects of the particular pre-transformation methods used to generate the shotgun libraries; thus, we first determined the extent to which variance in the gene coverage within libraries could be explained by the particular methods used to generate the libraries. Below, we first describe how the shotgun library methods were likely to bias gene coverage and how we controlled for these pre-transformation methodological effects on variance in coverage. We then describe our investigation of the ability of connectivity, divergence from the *E. coli* ortholog, and their statistical interaction to explain the remaining variance in coverage, that we infer to have resulted from differences in transformation efficiency, i.e., transferability.

**Figure 2.**
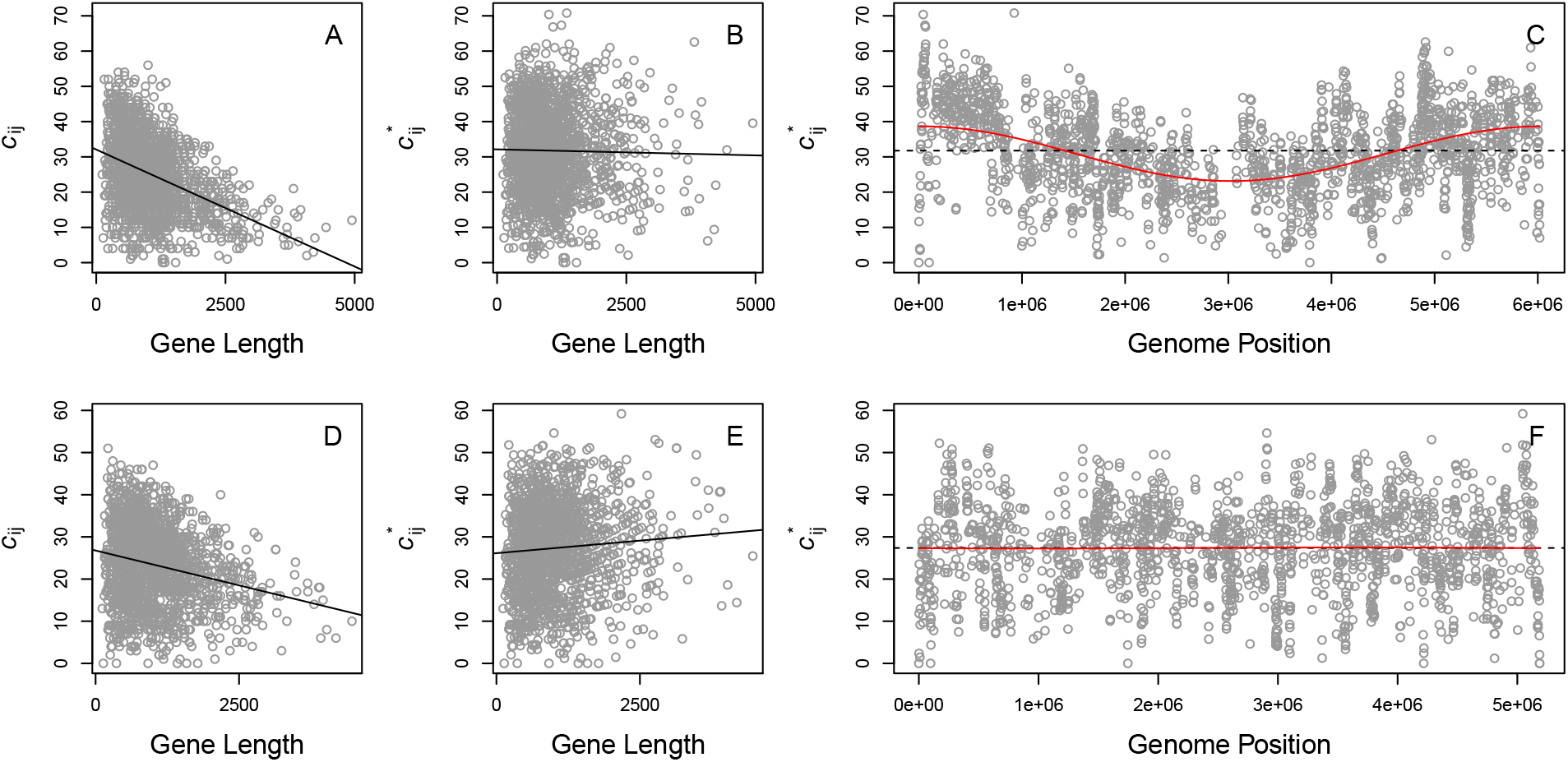
Methodological effects on coverage (*c_ij_*) in whole genome shotguns. As shown here for *Pseudomonas syringae* (A) and *Pseudoalteromonas atlantica* (D), long genes are substantially less well covered than short genes in all shotguns. By adjusting the raw coverage values of each gene solely by the Likelihood of complete coverage given the gene’s length (to obtain 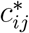; see Methods), the bias against long genes is dramatically reduced, as shown here for *P. syringae* (B) and *P. atlantica* (E). These adjusted coverage values, 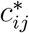, show positional biases in some genomes, e.g., *P. syringae* (C), but not others, e.g. *P. atlantica* (F). In all panels, points represent the coverage or adjusted coverage of individual genes (i.e., coding sequences) in the P. *syringae* (A, B, C) or *P. atlantica* (D, E, F) shotgun. Solid black lines (A, B, D, E) show the best fit linear relationship between coverage or adjusted coverage and gene length. Dashed black lines (C, F) are the mean adjusted coverage across all genes in the individual shotguns and red lines (C, F) are the sine curves that yield the best fit to the adjusted coverage data. Supplementary fig. S1 shows analogous data and model fits for the complete set of genome shotguns.

### Effect of gene length on coverage

In figs. 2A, 2D, and Supplementary fig. S1, we show the relationship between coverage and gene length in each shotgun library (note that 2 species possess 2 chromosomes and we analyzed data for each chromosome separately). It is apparent from these figures that long genes are less well covered than short genes in all shotgun libraries. The likely reason is straightforward. Long genes are less likely to be entirely contained within short cloned fragments.

To correct for this bias against long genes, we calculated the expected bias, *b_ij_*, against each gene *i* from genome *j*, given the length of the gene, Lij, and the distribution of cloned fragment lengths from genome *j* (Supplementary fig. S1; details in *Methods: Effect of gene length on coverage*). We then divided the observed raw coverage value, *c_ij_*, by the expected bias, *b_ij_*, to obtain an unbiased, length-corrected estimate *c_ij_*\* for that gene (figs. 2B, 2E, and Supplementary fig. 1). Before correcting for length bias, the effect of gene length on coverage was negative for 74 out of 74 bacterial chromosomes (Supplementary fig. S1). After correcting for length bias, the effect of gene length on coverage was negative in 21 and positive in 53 shotguns (the unbiased expectation is 37 negative and 37 positive), and has a much smaller statistical effect on coverage. Whereas gene length explains up to 43% of variance in raw coverage within individual shotgun libraries (median R^2^ = 0.092; range = 0.0023 to 0.43), gene length explains only up to 1.8% of variance in length-adjusted coverage (median R^2^ = 0.0020; range = 1.0 × 10^-5^ to 0.036).

### Effect of gene position on coverage

In figs. 2C, 2F, and Supplementary fig. S1, we show the adjusted coverage values, *c_ij_*\*, plotted against position in the genome for individual shotguns. The most obvious pattern is that genes near the origin of replication had higher coverage residuals than genes near the terminus of replication in some shotguns. Again, the likely reason for this pattern is straightforward. Because bacterial genome replication begins at the origin, genes near the origin are replicated before genes near the terminus. Thus, in actively dividing cells, genes near the origin are present in more copies within the cell than genes near the terminus. We infer that the difference in this pattern between shotguns resulted from a methodological difference in the bacterial growth phase, exponential or stationary, at the time genomes were harvested for use in the shotgun.

We estimated the positional bias in each genome j by fitting a sine curve to the *c_ij_*\* values as a function of the start position of each gene *i* (red lines in figs. 2C, 2F, and Supplementary fig. S1). The sine curves explained between 0.007% (fig. 2F; *Pseudomonas syringae pv. syringae*) and 45% (Supplementary fig. S1; *Arthrobacter sp. fb24*) of the variance in *c_ij_*\* (interquartile range = 1.32% to 7.57% variance explained). To correct for this positional bias, we divided the *c_ij_*\* values by the coverage expectation given the position of gene *i* in genome *j*, E(*c_ij_*\* | position_*ij*_), obtained from the best fit sine curve. For all downstream analyses, we consider the effects of various independent variables on the resulting dependent variable *c_ij_* = *c_ij_*\*/E(*c_ij_*\* | position_*ij*_), which we refer to as relative coverage.

### Biological effects on coverage

We next examined the effect of amino acid divergence between donor and recipient copies of orthologous transferred genes on relative coverage in the shotgun libraries. To increase statistical power, we limited this analysis to protein coding genes in the *E. coli* K12 genome for which we identified likely orthologs in a large number of bacterial genomes and for which the likely orthologs spanned a wide range of divergence (% amino acid difference) from the *E. coli* gene. Specifically, we required that each set of orthologous genes include at least 16 orthologs and exhibit a standard deviation greater than 12% amino acid difference. These criteria were based on a power analysis of the entire dataset (described in *Methods: Power Analysis*), but the results presented below were not qualitatively sensitive to the choice of criteria (Supplementary figs. S2 and S3). For the 1108 sets of orthologous genes that met these criteria, we calculated mean relative coverage and the slope of the relationship between relative coverage and divergence (% amino acid difference) from the *E. coli* gene (figs. 3A, 3E and Supplementary fig. S4; see *Methods: Effect of divergence on coverage*).

**Figure 3.**
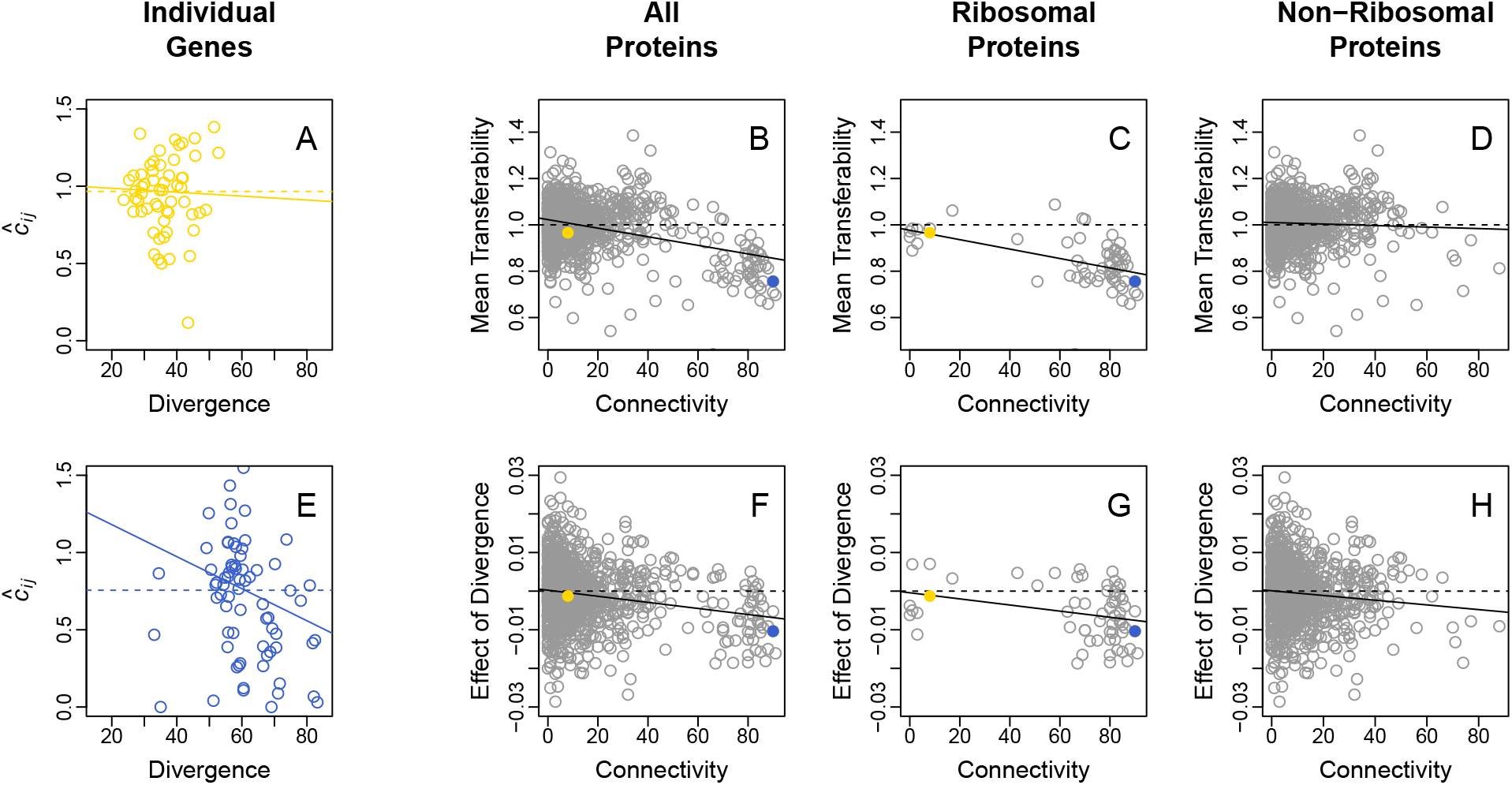
Biological effects on transferability in whole genome shotguns. We first estimated the mean value and the effect of divergence on transferability among the set of orthologs of each protein coding gene in the *E. coli* K12 genome, illustrated here for orthologs of the *E. coli* genes rbfA (A), and rplB (E). For a particular set of orthologs, mean transferability was estimated as mean relative coverage (mean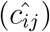, dashed lines in A and E), and the effect of divergence was estimated as the slope of the best fit linear relationship between relative coverage 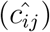 and divergence (% amino acid difference) from the *E. coli* gene (solid lines in A and E). Next we examined the effect of connectivity on the resulting estimates of mean transferability and of the effect of divergence, shown here for all of the genes in our collection (B and F), for ribosomal genes only (C and G), and for all genes except ribosomal genes (D and H). Points in A and E show the relative coverage and divergence from the *E. coli* gene of likely orthologs of the *E. coli* genes rbfA (A) and rplB (E). Each point is estimated from a different genome shotgun library. Points in B-D show mean transferability (dashed lines in A, E, and Supplementary fig. S4) and points in F-H show the effect of divergence (slope of the solid lines in A, E, and Supplementary fig. S4) for different orthologous gene sets as a function of the connectivity (number of protein-protein interactions) of the *E. coli* ortholog. Colored points in B, C, F, and G correspond to the orthologous gene sets shown in the same color in A and B. Solid black lines show the best fit linear relationship between mean transferability (B-D) or the effect of divergence (F-H) and connectivity.

For a particular set of orthologous genes, mean relative coverage (dashed lines in figs. 3A, 3E, and Supplementary fig. S2) provides a measure of the average transferability of divergent copies of that ortholog. Among the complete set of protein coding genes in our dataset, we found a significant negative relationship between this measure of transferability and connectivity (fig. 3B; estimate = −0.0019, *F*_1,1106_ = 127.2, *p* = 2 × 10^-16^). This main effect of connectivity appears to be largely driven by the ribosomal proteins. Narrowing our focus to the 61 genes that include the word “ribosomal” in the product field of the *E. coli* K12 genbank record, the negative relationship between transferability and connectivity is just as strong (fig. 3C; estimate = - 0.0020, *F*_1,59_ = 19.07, *p* = 5 10^-5^). By contrast, among the 1047 non-ribosomal genes, the relationship remains negative, but is weaker and not statistically significant (fig. 3D; estimate = - 0.0003, *F*_1,1045_ = 1.56, *p* = 0.2116).

For a particular set of orthologous genes, the slope of the relationship between relative coverage and divergence from the *E. coli* gene (solid lines in figs. 3A, 3E, and Supplementary fig. S2) estimates the main effect of divergence on transferability of the orthologs. Among the complete set of 1108 protein coding genes in our dataset, we found a significant negative relationship between this main effect of divergence and connectivity (Figure 3F, estimate = −7.87 × 10^-5^, *F*_1,1106_ = 41.96, *p* = 1.4 × 10^-10^). That is, we found a significant divergence × connectivity interaction. In contrast to the main effect of connectivity, the divergence × connectivity interaction appears to be a general characteristic of protein coding genes, that is not limited to ribosomal proteins. The divergence × connectivity interaction is negative and of similar magnitude among the 61 ribosomal proteins (Figure 3G; estimate = −7.92 × 10^-5^, *F*_1,59_ = 5.771, *p* = 0.0195) and the 1047 non-ribosomal proteins (Figure 3H; estimate = −6.1 10^-5^, *F*_1,1045_ = 9.329, *p* = 0.0023). The results described in this and the previous paragraph do not appear to be constrained to the 1108 genes we analyzed here. All of these results were qualitatively unchanged in an analysis of the 2420 orthologous gene sets that met the less stringent criteria of including a minimum of 5 orthologs and a standard deviation greater than 5% amino acid difference (Supplementary fig. S5).

## Discussion

In this paper, we built on the recognition by Sorek et al. (2007) that the shotgun libraries used to generate the earliest whole prokaryotic genome sequences could be used as experimental tests of the Complexity Hypothesis. Sorek et al. (2007) observed across 85 finished microbial genomes that a subset of genes appeared “unclonable”, as indicated by an absence or reduced number of sequencing reads spanning a gene. They then followed a bottom-up approach, investigating particular genes from particular genomes that failed to transfer and inferring the molecular mechanism that explained each failure. Here, we focused on a top-down approach, using the shotgun libraries to estimate relative rates of transfer for large numbers of genes from large numbers of genomes. We then tested the ability of two general hypotheses – the Balance and Complexity hypotheses – to explain differences in the estimated transfer rates between genes and genomes. Our quantitative analysis confirmed that 1) transferability decreased with the connectivity of the transferred gene, and 2) divergence between transferred and native orthologs reduced transferability more for genes with higher connectivities than for genes with lower connectivities. The first observation is predicted by both the Balance and Complexity hypotheses; the second observation is predicted *only* by the Complexity hypothesis (fig. 1). Thus, although our analysis is consistent with the Balance Hypothesis that the cost of protein overexpression increases with connectivity, it provides stronger support for the Complexity Hypothesis. Specifically, our analysis suggests that the success or failure of a transferred gene to engage in normal protein-protein interactions is an important determinant of HGT, and that the probability of HGT failure increases via a negative (i.e., synergistic) interaction between increasing connectivity and increasing divergence.

Our results highlight the difficulty inherent in testing the Balance and Complexity Hypotheses experimentally. A previous experimental test (Acar Kirit et al. 2020) did not detect a statistically significant main effect of connectivity on transferability (as defined in fig. 1C). Acar Kirit et al. (2020) achieved more precise measures of transferability, albeit from a substantially smaller number of genes (44 compared to 1108) and from a smaller number of donor genomes (1 compared to ≥16 for each gene). Thus, the difference in outcome is likely to have resulted from their limited statistical power to detect a weak or noisy main effect of connectivity on transferability. Although their precise fitness assays enabled the detection of fitness costs imposed by increasing the number of disordered regions and the length of transferred proteins, their use of only a single donor genome prevented an examination of the statistical interaction between connectivity and divergence in their effects on transferability (as in fig. 1D). Because the detection of interaction effects requires more statistical power than the detection of main effects, our ability both to differentiate between the two hypotheses and to support the Complexity Hypothesis was likely only possible because our dataset was compiled from 74 whole genome shotgun libraries.

The observation that ribosomal proteins drive the signal associated with the Balance hypothesis (figs. 3B – 3D) may provide additional support for this idea. As the most highly expressed proteins in the cell, the transfer and resulting overexpression of a highly connected ribosomal protein might be expected to result in a greater stoichiometric imbalance among its interaction partners than the transfer and resulting overexpression of a non-ribosomal protein that is expressed at a lower level. Thus, inherently high levels of ribosomal gene expression may explain why we had greater statistical power and estimated a larger main effect of connectivity on transferability among ribosomal than non-ribosomal genes.

By contrast, our strongest support of the Complexity Hypothesis comes from the *consistency* of the interaction between divergence and connectivity among both ribosomal and non-ribosomal genes (fig. 3), like transferases and kinases (Supplementary fig. S3). The Complexity Hypothesis was proposed to explain the observation from comparative data that HGT has happened less often among informational genes, like the subunits of ribosomes and polymerase complexes, than among operational genes, like enzymes (Rivera et al. 1998; Jain et al. 1999). The Complexity Hypothesis posits that the observed difference in the rate of HGT resulted, not from the difference in function between informational and operational genes, but rather from the large difference in the connectivities of these different types of genes. Its central prediction is that increases in connectivity and the divergence between donor and recipient orthologs will interact synergistically to reduce transferability. In short, highly divergent genes with many protein-protein interactions will exhibit the lowest rates of HGT, regardless of function. Thus, our finding that connectivity interacts with divergence to reduce transferability, not only among the ribosomal (i.e., informational) genes but also among non-ribosomal (i.e. non-informational) genes supports both the central prediction of the Complexity Hypothesis and its underlying logic, that informational genes exhibit lower rates of HGT specifically because they are highly connected, not because they perform an informational function.

Our results confirm the central predictions of both of the two non-exclusive hypotheses used to explain why transferability should decrease with connectivity. We provide the strongest support for the role of protein-protein interaction failure among divergent orthologs of highly-connected genes (the Complexity Hypothesis), but we also provide support for a role of gene misexpression (the Balance Hypothesis). Together, our results support the emerging “rule” that deleterious interactions among protein partners in bacteria may govern the frequency with which individual genes successfully undergo HGT. Given that HGT is a major contributor to bacterial genome evolution, our work suggests that the Balance and Complexity Hypotheses may shape phenotypic diversity, drive the expansion of protein families, and affect the evolution of new phenotypes, new metabolic pathways, and new species in bacteria.

## Materials and Methods

### Data Sources

The complete set of whole genome shotguns and assembled genomes used in this work are described in Supplementary Table S1.

We obtained whole genome shotgun reads from NCBI’s trace archive (ftp://ftp.ncbi.nih.gov/pub/TraceDB). We included all of the shotguns examined in (Sorek et al. 2007) except Candidatus Koribacter versatilis Ellin345, whose shotgun reads were no longer available in the trace archive.

We obtained complete bacterial genome sequences from NCBI’s microbial genome database (https://www.ncbi.nlm.nih.gov/genome/microbes/). From these sequence files, we determined the protein sequence, start position, end position, and length for each coding sequence in each genome.

We obtained protein interaction data from the STRING database version 11.5 (http://string-db.org). We downloaded two files containing detailed confidence scores associated with evidence of protein-protein interactions in *Escherichia coli* strain K12 substrain MG1655, one describing evidence of any type of interaction and the other describing evidence only of physical interactions. Confidence scores range from 0 (low confidence) to 999 (high confidence). For the analysis described here, we considered pairs of proteins to be interacting if their confidence score was greater than 900.

### Gene Coverage in the Shotguns

The coverage, *c_ij_*, of gene *i* in the whole genome shotgun of species *j* was calculated as the number of plasmid inserts in shotgun *j* that contain gene *i* in its entirety. Coverage values were determined by mapping the paired reads from each whole genome shotgun to the corresponding assembled genome using the BWA-SW algorithm from the Burrows-Wheeler Aligner (Li & Durbin 2010). A small minority of reads mapped to more than one location. We identified and eliminated most of the incorrectly mapped reads by requiring a phred-scaled map quality score greater than 150 and a distance between paired reads of fewer than 100,000 bases. For the few multiply mapped reads that remained, one of the mapping locations was chosen at random. Read pairs for which one read mapped upstream of gene *i* and one read mapped downstream of gene *i* were counted toward the coverage of gene *i*. Read pairs that entirely spanned more than one gene were counted toward all of the spanned genes. Although we mapped read pairs to all of the replicons that comprised each genome, only genes contained on large replicons (i.e., chromosomes) were included in downstream analyses. We ignored genes on plasmids.

### Effect of gene length on coverage

To control for the bias against long genes we calculated for each genome *j* a Likelihood of observing a gene of a particular length, *l*, given the actual distribution of cloned fragment lengths, *f_j_*, that comprised whole genome shotgun *j*.

For an individual cloned fragment of length *x*, the probability that it contained the entirety of a gene of length *l* is proportional to the difference in the lengths between the cloned fragment and the gene, if the cloned fragment is at least as long as the gene, or 0 otherwise:

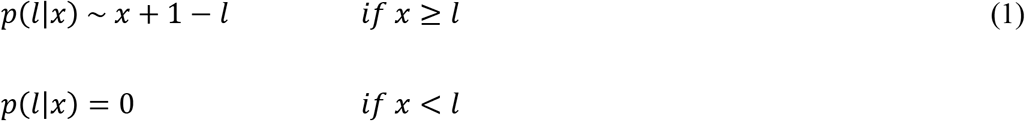

For each genome *j*, the Likelihood of observing each gene, *g_ij_*, of length *l_ij_*, given the distribution of cloned fragment lengths, *f_j_*, is then calculated by summing these individual probabilities over all of the observed cloned fragments lengths *x_j_* in the shotgun of genome *j*:

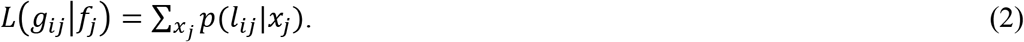

For each genome *j*, the bias against long genes, *b_j_*(*l*), was then calculated as the Likelihood of observing a gene of length *l* relative to that of the most likely (i.e. the shortest) gene in the same genome:

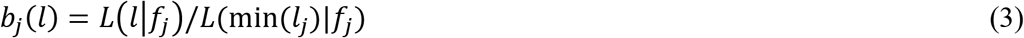

For each genome, coverage values for each gene were adjusted by the length biases calculated for that genome, so that the data used for all downstream analysis were of the form:

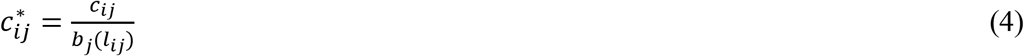

### Effect of gene position on coverage

To control for the long-range positional biases we fit the adjusted coverage data from individual genomes to a sine curve using the *lm* function in R (Figure S3). For gene *i* in genome *j*, we calculated relative coverage, *c_ij_*, by dividing its length-adjusted coverage value, *c_ij_*\*, by the fitted value at that position in the genome, E(*c_ij_*\* | position_*ij*_):

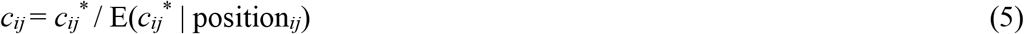

### Effect of divergence on coverage

For each protein-coding gene in the *E. coli* K12 genome, we identified the coding sequences in each shotgun library that were reciprocal best hits to the *E. coli* K12 gene using BLASTp (Altschul et al. 1990). Coding sequences that were not reciprocal best hits could not be identified as the most likely ortholog of a particular coding sequence in *E. coli* K12 and were, therefore, excluded from downstream analyses. The protein sequences of reciprocal best hits were aligned to the *E. coli* K12 gene using the software ProbCons (Do et al. 2005) and the alignments were used to calculate protein divergence as % amino acid difference from the *E. coli* K12 ortholog. For each of the resulting sets of orthologous genes, we used the *lm* function from the *stats* package in R (version 4.1.2) to determine the best-fit linear relationship between the relative coverage of each ortholog in the set and its divergence from the *E. coli* ortholog.

### Statistical Tests of the Balance and Complexity Hypotheses

We used the *lm* function from the *stats* package in R (version 4.1.2) to investigate the effects of connectivity and divergence on the transferability into *E. coli* of orthologs of the protein coding genes in the *E. coli* genome (as illustrated in fig. 1). For each *E. coli* gene, we estimated mean transferability (the x-axis in fig. 1C) as the mean relative coverage across all orthologs of the gene. We measured the effect of divergence on transferability (i.e. the main effect of divergence; x-axis in fig. 1D) as the slope of the linear relationship between relative coverage and divergence for all orthologs of the gene (as described above in *Methods: Effect of divergence on coverage*). In essence, this statistical approach considers connectivity and divergence as fixed effects and the identity of the *E. coli* ortholog as a random effect. The experimental units are the sets of gene orthologs and the analysis considers the individual orthologs in each set to be repeated measures of a particular orthologous gene along a divergence gradient.

### Power Analysis

A visual examination of the data in Supplementary fig. S4 revealed that the highest positive and lowest negative estimates of the effects of divergence on relative coverage were most common among genes for which the set of identified orthologs spanned only a narrow range of divergence values or for which we identified only very few orthologs. In addition, the smaller sets of orthologous genes often included at least one gene with an exceptionally high divergence (> 80% amino acid difference) from the *E. coli* ortholog, suggesting that these sets included donor genes that were not true orthologs. To examine the sensitivity of our statistical tests to the noisy data that resulted from these issues we repeated the analysis described above, requiring that each set of orthologous genes include at least X orthologs and exhibit a standard deviation greater than Y% amino acid difference, where X and Y were both varied between 2 and 22. For each pair of X and Y, we estimated the number of genes that met both requirements, the main effect of connectivity on mean transferability, and the *F* statistic and *p* value associated with the statistical test of the effect (Supplementary fig. S2). For each pair of X and Y, we also estimated the connectivity × divergence interaction effect (i.e., the effect of connectivity on the main effect of divergence; see fig. 1D), as well as the *F* statistic and *p* value associated with the statistical test (Supplementary fig. S3).

## Data and Code Availability

The data are publicly available and described in Supplementary Table S1.

The analysis pipeline consists of programs written in Python 3.6 and R version 4.1.2 (R Development Core Team 2013), and UNIX shell scripts that call those programs, BEDTools (Quinlan & Hall 2010), and SAMtools (Li et al. 2009). All of our code and scripts are publically available at https://bitbucket.org/cburch/burch-et-al-2022.

