## Supplementary Material for "Genome-wide determination of barriers to horizontal gene transfer"

Supplementary Table S1

| Accession # | Replicon Name | Replicon Size | # CDS |
|---|---|---|---|
| CP000481.1 | Acidothermus cellulolyticus 11B | 2,443,540 | 2,157 |
| CP000453.1 | Alkalilimnicola ehrlichii MLHE-1 | 3,275,944 | 2,865 |
| CP000251.1 | Anaeromyxobacter dehalogenans 2CP-C | 5,013,479 | 4,346 |
| CP000455.1 | Arthrobacter sp. FB24 plasmid 1 | 159,538 | 158 |
| CP000456.1 | Arthrobacter sp. FB24 plasmid 2 | 115,507 | 108 |
| CP000457.1 | Arthrobacter sp. FB24 plasmid 3 | 96,488 | 94 |
| CP000454.1 | Arthrobacter sp. FB24 | 4,698,945 | 4,146 |
| NC_003997.3 | Bacillus anthracis str. Ames | 5,227,293 | 5,337 |
| AE017195.1 | Bacillus cereus ATCC 10987 plasmid pBc10987 | 208,369 | 241 |
| AE017194.1 | Bacillus cereus ATCC 10987 | 5,224,283 | 5,603 |
| CP000238.1 | Baumannia cicadellinicola str. Hc (Homalodisca coagulata) | 686,194 | 595 |
| NC_004310.3 | Brucella suis 1330 chromosome I | 2,107,794 | 1,979 |
| AE014292.2 | Brucella suis 1330 chromosome II | 1,207,381 | 1,150 |
| CP000440.1 | Burkholderia ambifaria AMMD chromosome 1 | 3,556,545 | 3,206 |
| CP000441.1 | Burkholderia ambifaria AMMD chromosome 2 | 2,646,969 | 2,346 |
| CP000442.1 | Burkholderia ambifaria AMMD chromosome 3 | 1,281,472 | 1,013 |
| CP000443.1 | Burkholderia ambifaria AMMD plasmid 1 | 43,581 | 45 |
| NC_008061.1 | Burkholderia cenocepacia AU 1054 chromosome 2 | 2,788,459 | 2,497 |
| NC_007510.1 | Burkholderia lata chromosome 1 | 3,694,126 | 3,349 |
| NC_007511.1 | Burkholderia lata chromosome 2 | 3,587,082 | 3,203 |
| NC_007509.1 | Burkholderia lata chromosome 3 | 1,395,069 | 1,207 |
| NZ_CP009727.1 | Burkholderia mallei strain Turkey2 chromosome 1 | 3,471,484 | 3,088 |
| NZ_CP009728.1 | Burkholderia mallei strain Turkey2 chromosome 2 | 2,120,565 | 1,754 |
| CP000086.1 | Burkholderia thailandensis E264 chromosome I | 3,809,201 | 3,276 |
| CP000085.1 | Burkholderia thailandensis E264 chromosome II | 2,914,771 | 2,358 |
| NC_003912.7 | Campylobacter jejuni RM1221 | 1,777,831 | 1,818 |
| CP000141.1 | Carboxydothermus hydrogenoformans Z-2901 | 2,401,520 | 2,620 |
| NC_008242.1 | Chelativorans sp. BNC1 plasmid 1 | 343,931 | 330 |
| NC_008243.1 | Chelativorans sp. BNC1 plasmid 2 | 131,247 | 122 |
| NC_008244.1 | Chelativorans sp. BNC1 plasmid 3 | 47,561 | 49 |
| NC_008254.1 | Chelativorans sp. BNC1 | 4,412,446 | 4,150 |
| NC_004720.1 | Chlamydia caviae GPIC plasmid pCpGP1 | 7,966 | 9 |
| NC_003361.3 | Chlamydia caviae GPIC | 1,173,390 | 986 |
| NC_002932.3 | Chlorobaculum tepidum TLS | 2,154,946 | 2,029 |
| NC_003910.7 | Colwellia psychrerythraea 34H | 5,373,180 | 4,436 |
| NC_004704.2 | Coxiella burnetii RSA 493 plasmid pQpH1 | 37,319 | 35 |
| NC_002971.4 | Coxiella burnetii RSA 493 | 1,995,488 | 1,798 |
| NC_007974.2 | Cupriavidus metallidurans CH34 megaplasmid | 2,580,084 | 2,305 |
| NC_007972.2 | Cupriavidus metallidurans CH34 plasmid pMOL28 | 171,459 | 167 |
| NC_007971.2 | Cupriavidus metallidurans CH34 plasmid pMOL30 | 233,720 | 246 |
| NC_007973.1 | Cupriavidus metallidurans CH34 | 3,928,089 | 3,596 |
| NC_007347.1 | Cupriavidus pinatubonensis JMP134 chromosome 1 | 3,806,533 | 3,490 |
| NC_007348.1 | Cupriavidus pinatubonensis JMP134 chromosome 2 | 2,726,152 | 2,467 |
| NC_007336.1 | Cupriavidus pinatubonensis JMP134 megaplasmid | 634,917 | 570 |
| NC_007337.1 | Cupriavidus pinatubonensis JMP134 plasmid 1 | 87,688 | 89 |
| NC_007298.1 | Dechloromonas aromatica RCB | 4,501,104 | 4,237 |
| CP000027.1 | Dehalococcoides mccartyi 195 chromosome | 1,469,720 | 1,580 |
| CP000358.2 | Deinococcus geothermalis DSM 11300 plasmid pDGEO01 | 574,127 | 519 |
| CP000856.1 | Deinococcus geothermalis DSM 11300 plasmid pDGEO02 | 205,686 | 205 |

| | | | |
|---|---|---|---|
| CP000359.1 | Deinococcus geothermalis DSM 11300 | 2,467,205 | 2,330 |
| NC_007354.1 | Ehrlichia canis str. Jake | 1,315,030 | 947 |
| CP000236.1 | Ehrlichia chaffeensis str. Arkansas | 1,176,248 | 1,105 |
| NC_000913.3 | Escherichia coli str. K-12 substr. MG1655 | 4,641,652 | 4,302 |
| NC_007777.1 | Frankia casuarinae | 5,433,628 | 4,659 |
| CP000149.1 | Geobacter metallireducens GS-15 plasmid | 13,762 | 17 |
| CP000148.1 | Geobacter metallireducens GS-15 | 3,997,420 | 3,551 |
| NC_002939.5 | Geobacter sulfurreducens PCA | 3,814,128 | 3,424 |
| NC_007520.2 | Hydrogenovibrio crunogenus XCL-2 | 2,427,734 | 2,257 |
| CP000265.1 | Jannaschia sp. CCS1 plasmid1 | 86,072 | 71 |
| CP000264.1 | Jannaschia sp. CCS1 | 4,317,977 | 4,212 |
| NC_003210.1 | Listeria monocytogenes EGD-e chromosome | 2,944,528 | 2,867 |
| NC_008347.1 | Maricaulis maris MCS10 | 3,368,780 | 3,120 |
| NC_007955.1 | Methanococcoides burtonii DSM 6242 | 2,575,032 | 2,539 |
| NC_007349.1 | Methanosarcina barkeri str. Fusaro plasmid 1 | 36,358 | 25 |
| NC_007355.1 | Methanosarcina barkeri str. Fusaro | 4,837,408 | 3,995 |
| NC_007796.1 | Methanospirillum hungatei JF-1 | 3,544,738 | 3,365 |
| CP000556.1 | Methylibium petroleiphilum PM1 plasmid RPME01 | 599,444 | 630 |
| CP000555.1 | Methylibium petroleiphilum PM1 | 4,044,195 | 3,819 |
| NC_007947.1 | Methylobacillus flagellatus KT | 2,971,517 | 2,798 |
| NC_002977.6 | Methylococcus capsulatus str. Bath | 3,304,561 | 3,028 |
| NC_007644.1 | Moorella thermoacetica ATCC 39073 | 2,628,784 | 2,631 |
| NC_008147.1 | Mycobacterium sp. MCS Plasmid1 | 215,075 | 236 |
| NC_008146.1 | Mycobacterium sp. MCS | 5,705,448 | 5,468 |
| NC_007798.1 | Neorickettsia sennetsu str. Miyayama | 859,006 | 747 |
| CP000320.1 | Nitrobacter hamburgensis X14 plasmid 1 | 294,829 | 239 |
| CP000321.1 | Nitrobacter hamburgensis X14 plasmid 2 | 188,318 | 172 |
| CP000322.1 | Nitrobacter hamburgensis X14 plasmid 3 | 121,408 | 111 |
| CP000319.1 | Nitrobacter hamburgensis X14 | 4,406,967 | 3,804 |
| CP000115.1 | Nitrobacter winogradskyi Nb-255 | 3,402,093 | 3,122 |
| CP000126.1 | Nitrosococcus oceani ATCC 19707 plasmid A | 40,420 | 43 |
| CP000127.1 | Nitrosococcus oceani ATCC 19707 | 3,481,691 | 2,976 |
| CP000676.1 | Novosphingobium aromaticivorans DSM 12444 plasmid pNL1 | 184,462 | 182 |
| CP000677.1 | Novosphingobium aromaticivorans DSM 12444 plasmid pNL2 | 487,268 | 431 |
| CP000248.1 | Novosphingobium aromaticivorans DSM 12444 | 3,561,584 | 3,324 |
| CP000483.1 | Pelobacter propionicus DSM 2379 plasmid pPRO1 | 202,397 | 195 |
| CP000484.1 | Pelobacter propionicus DSM 2379 plasmid pPRO2 | 30,722 | 33 |
| CP000482.1 | Pelobacter propionicus DSM 2379 | 4,008,000 | 3,576 |
| CP000317.1 | Polaromonas sp. JS666 plasmid 1 | 360,405 | 326 |
| CP000318.1 | Polaromonas sp. JS666 plasmid 2 | 338,007 | 310 |
| CP000316.1 | Polaromonas sp. JS666 | 5,200,264 | 4,817 |
| NC_007577.1 | Prochlorococcus marinus str. MIT 9312 | 1,709,204 | 1,858 |
| NC_007335.2 | Prochlorococcus marinus str. NATL2A | 1,842,899 | 2,040 |
| NC_008228.1 | Pseudoalteromonas atlantica T6c | 5,187,005 | 4,356 |
| NZ_CP068034.2 | Pseudomonas syringae strain BIM B-268 chromosome | 6,018,586 | 5,081 |
| CP000324.1 | Psychrobacter cryohalolentis K5 plasmid 1 | 41,221 | 44 |
| CP000323.1 | Psychrobacter cryohalolentis K5 | 3,059,876 | 2,467 |
| CP000268.1 | Rhodoferax ferrireducens T118 plasmid1 | 257,447 | 248 |
| CP000267.1 | Rhodoferax ferrireducens T118 | 4,712,337 | 4,170 |
| NC_008435.1 | Rhodopseudomonas palustris BisA53 | 5,505,494 | 4,948 |
| NC_007925.1 | Rhodopseudomonas palustris BisB18 | 5,513,844 | 4,984 |
| CP000283.1 | Rhodopseudomonas palustris BisB5 | 4,892,717 | 4,397 |

| | | | |
|---|---|---|---|
| CP000250.1 | Rhodopseudomonas palustris HaA2 | 5,331,656 | 4,683 |
| CP000386.1 | Rubrobacter xylanophilus DSM 9941 | 3,225,748 | 3,140 |
| CP000377.1 | Ruegeria sp. TM1040 chromosome | 3,200,938 | 3,030 |
| CP000302.1 | Shewanella denitrificans OS217 | 4,545,906 | 3,754 |
| CP000447.1 | Shewanella frigidimarina NCIMB 400 | 4,845,257 | 4,029 |
| CP000469.1 | Shewanella sp. ANA-3 chromosome 1 | 4,972,204 | 4,111 |
| CP000470.1 | Shewanella sp. ANA-3 plasmid 1 | 278,942 | 249 |
| CP000376.1 | Silicibacter sp. TM1040 mega plasmid | 821,788 | 728 |
| CP000375.1 | Silicibacter sp. TM1040 plasmid | 130,973 | 106 |
| CP000357.1 | Sphingopyxis alaskensis RB2256 F plasmid | 28,543 | 30 |
| CP000356.1 | Sphingopyxis alaskensis RB2256 | 3,345,170 | 3,165 |
| NZ_CP035288.1 | Staphylococcus epidermidis strain ATCC 14990 chromosome | 2,466,502 | 2,241 |
| NZ_CP035289.1 | Staphylococcus epidermidis strain ATCC 14990 plasmid unnamed1 | 20,117 | 23 |
| NZ_CP035290.1 | Staphylococcus epidermidis strain ATCC 14990 plasmid unnamed2 | 4,439 | 3 |
| NZ_CP012480.1 | Streptococcus agalactiae strain NGBS128 chromosome | 2,074,179 | 1,952 |
| NZ_CP012480.1 | Streptococcus agalactiae strain NGBS128 chromosome | 2,074,179 | 1,952 |
| NZ_CP012742.1 | Streptococcus agalactiae strain NGBS128 plasmid pNGBS128 | 4,944 | 7 |
| NZ_CP012742.1 | Streptococcus agalactiae strain NGBS128 plasmid pNGBS128 | 4,944 | 7 |
| CP000153.1 | Sulfurimonas denitrificans DSM 1251 | 2,201,561 | 2,096 |
| NZ_CP033061.1 | Synechococcus elongatus UTEX 3055 chromosome | 2,767,524 | 2,751 |
| NZ_CP033062.1 | Synechococcus elongatus UTEX 3055 plasmid unnamed1 | 89,249 | 97 |
| NZ_CP033063.1 | Synechococcus elongatus UTEX 3055 plasmid unnamed2 | 24,450 | 24 |
| NC_007516.1 | Synechococcus sp. CC9605 | 2,510,659 | 2,751 |
| NC_007333.1 | Thermobifida fusca YX | 3,642,249 | 3,088 |
| NC_002967.9 | Treponema denticola ATCC 35405 | 2,843,201 | 2,567 |
| NC_008312.1 | Trichodesmium erythraeum IMS101 | 7,750,108 | 5,263 |
| NC_007410.1 | Trichormus variabilis ATCC 29413 plasmid A | 366,354 | 360 |
| NC_007411.1 | Trichormus variabilis ATCC 29413 plasmid B | 35,762 | 33 |
| NC_007412.1 | Trichormus variabilis ATCC 29413 plasmid C | 300,758 | 252 |
| NC_014000.1 | Trichormus variabilis ATCC 29413 | 37,151 | 58 |
| NC_007413.1 | Trichormus variabilis ATCC 29413 | 6,365,727 | 5,185 |

**Figure S1. Methodological effects on coverage in whole genome shotguns.** The raw and adjusted data from each bacterial chromosome is described by a set of 4 panels, that show (A) the length distribution of successfully transformed genome fragments and the relationships between (B) raw coverage, $c_{ij}$, and gene length, (C) length-corrected coverage, $c^*_{ij}$, and gene length, and (D) length-corrected coverage, $c^*_{ij}$, and position in the genome. Long genes are substantially less well covered than short genes in all libraries (i.e., the best fit linear relationship between coverage and gene length (black line) has a negative slope for 74 out of 74 chromosomes; B panels). By adjusting the raw coverage values of each gene solely by the Likelihood of complete coverage given the gene's length (to obtain $c^*_{ij}$; see Methods), the bias against long genes is dramatically reduced (the relationship between $c^*_{ij}$ and gene length is negative for 21 out of 74 chromosomes; C panels). The adjusted coverage values, $c^*_{ij}$, show positional biases in some genomes but not others (D panels). In all panels, points represent the coverage or adjusted coverage of individual genes (i.e., coding sequences) in the shotgun library indicated above each row of panels. Dashed black lines are the mean adjusted coverage across all genes in the individual shotguns and red lines are the sine curves that yield the best fit to the adjusted coverage data (D panels).

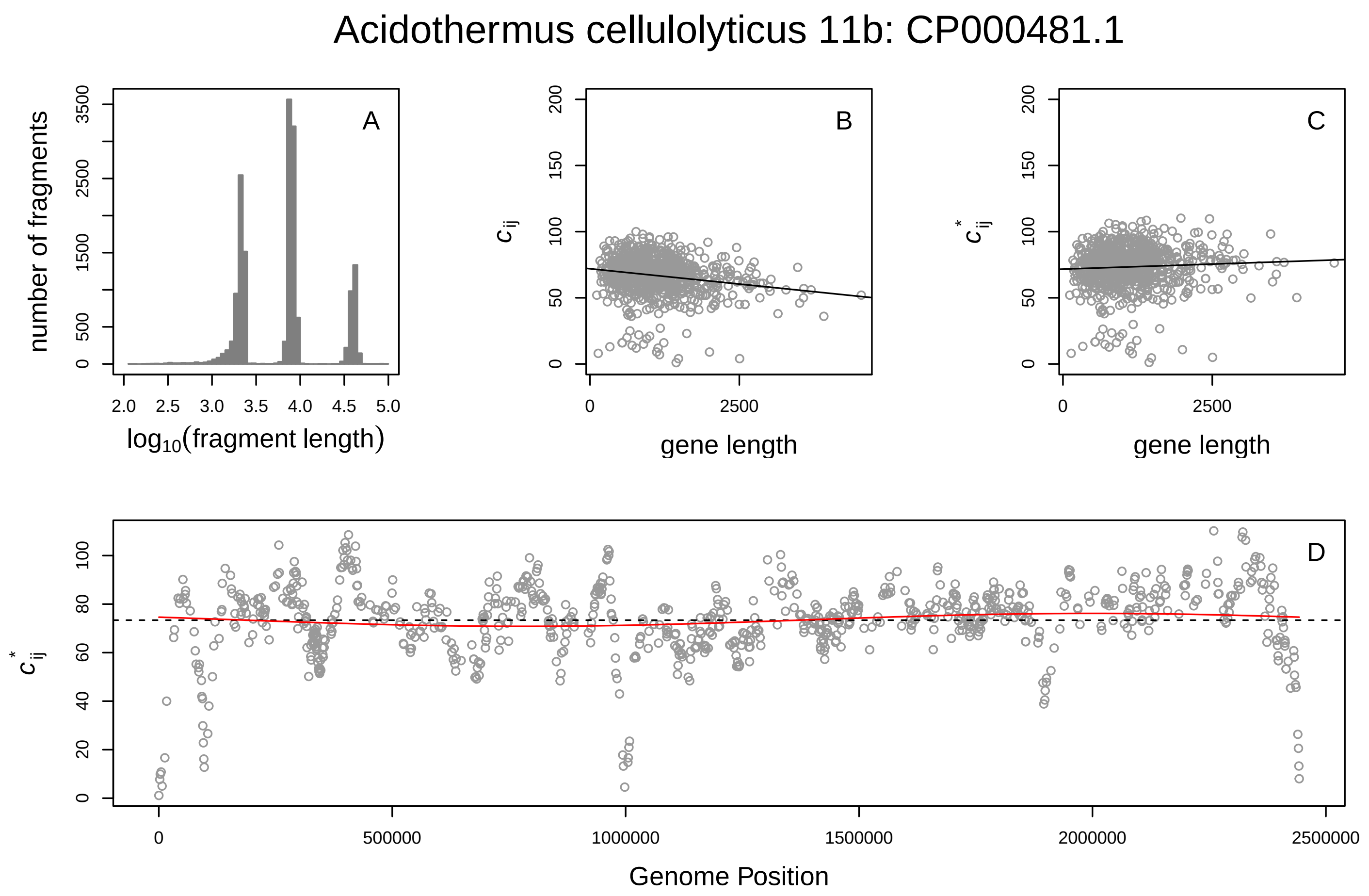
Acidothermus cellulolyticus 11b: CP000481.1
A
number of fragments
log10(fragment length)
B
c_ij
gene length
C
c*_ij
gene length
D
c*_ij
Genome Position

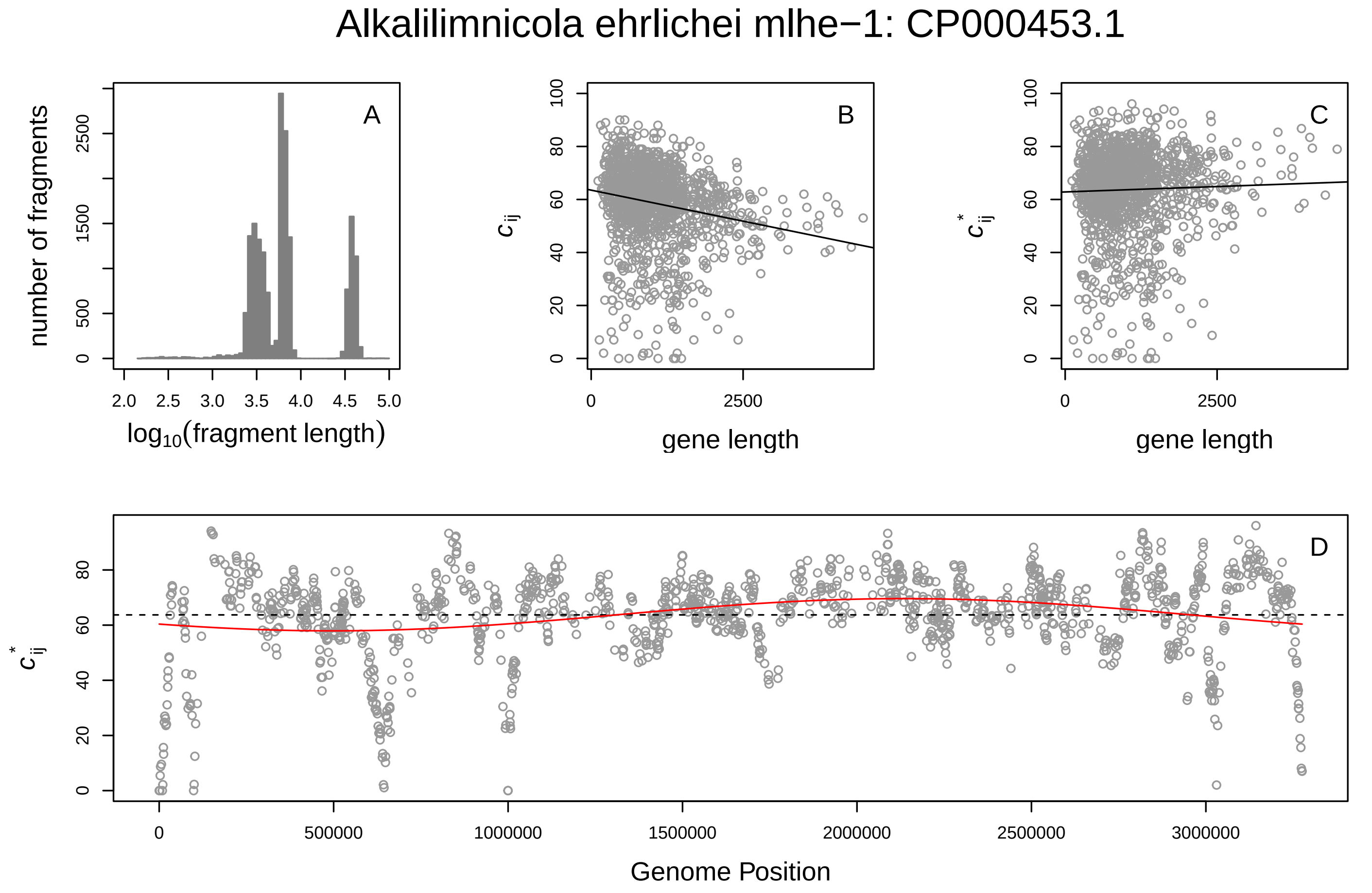
Alkalilimnicola ehrlichei mlhe−1: CP000453.1
A
number of fragments
log10(fragment length)
B
c_ij
gene length
C
c*_ij
gene length
D
c*_ij
Genome Position

#### Anabaena variabilis atcc 29413: NC_007413.1

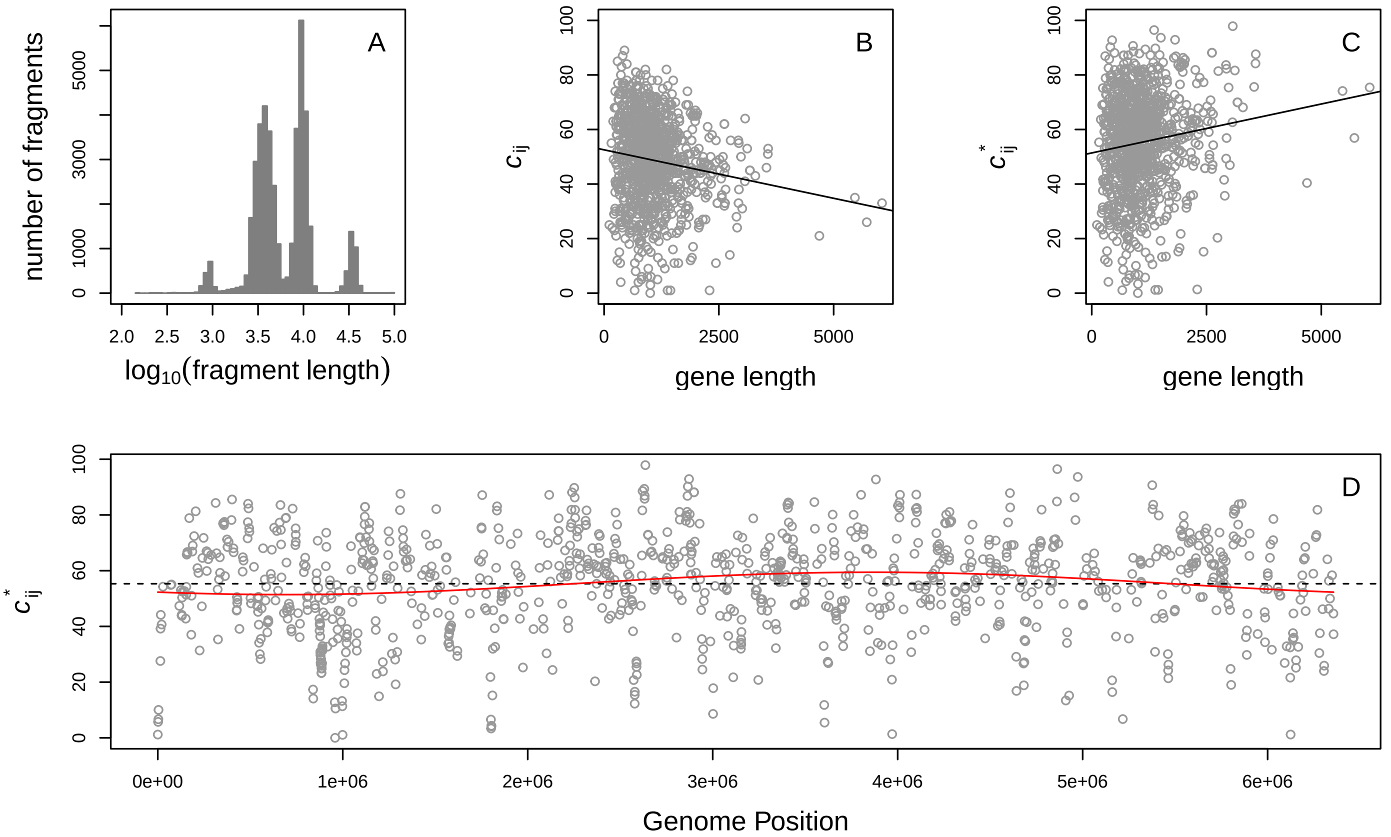


#### Anaeromyxobacter dehalogenans 2cp−c: CP000251.1

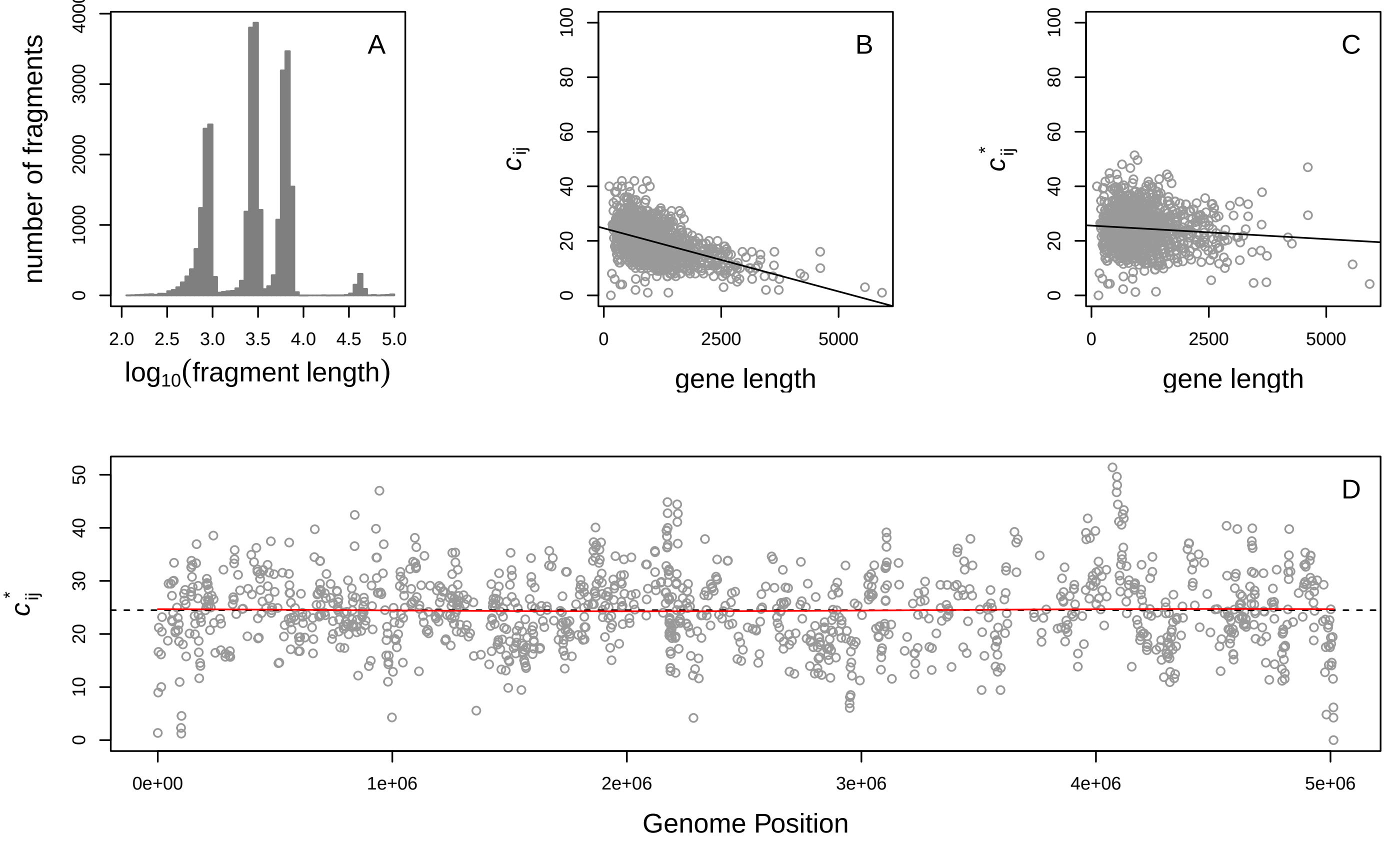

Arthrobacter sp fb24: CP000454.1

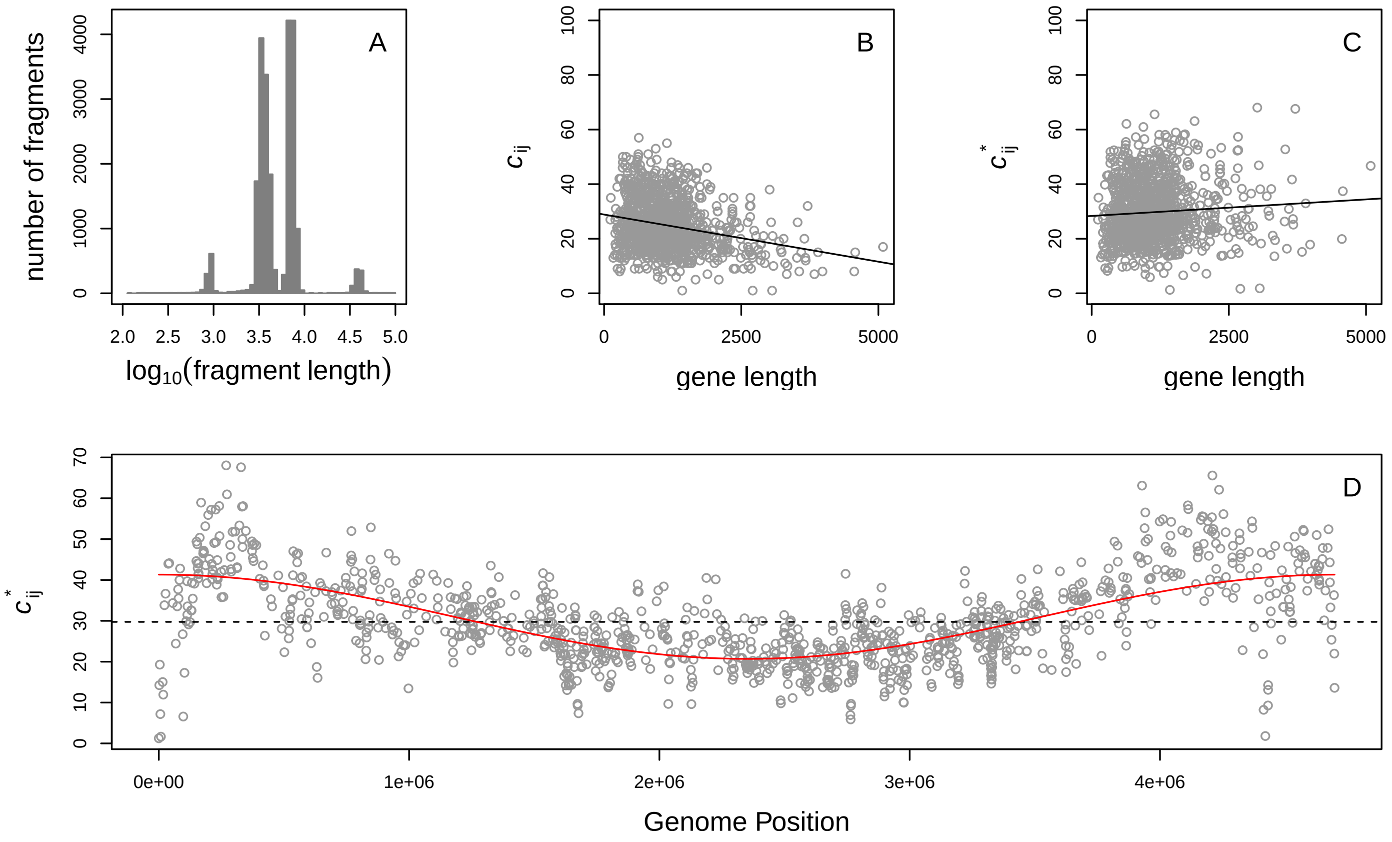


Bacillus anthracis Ames: NC_003997.3

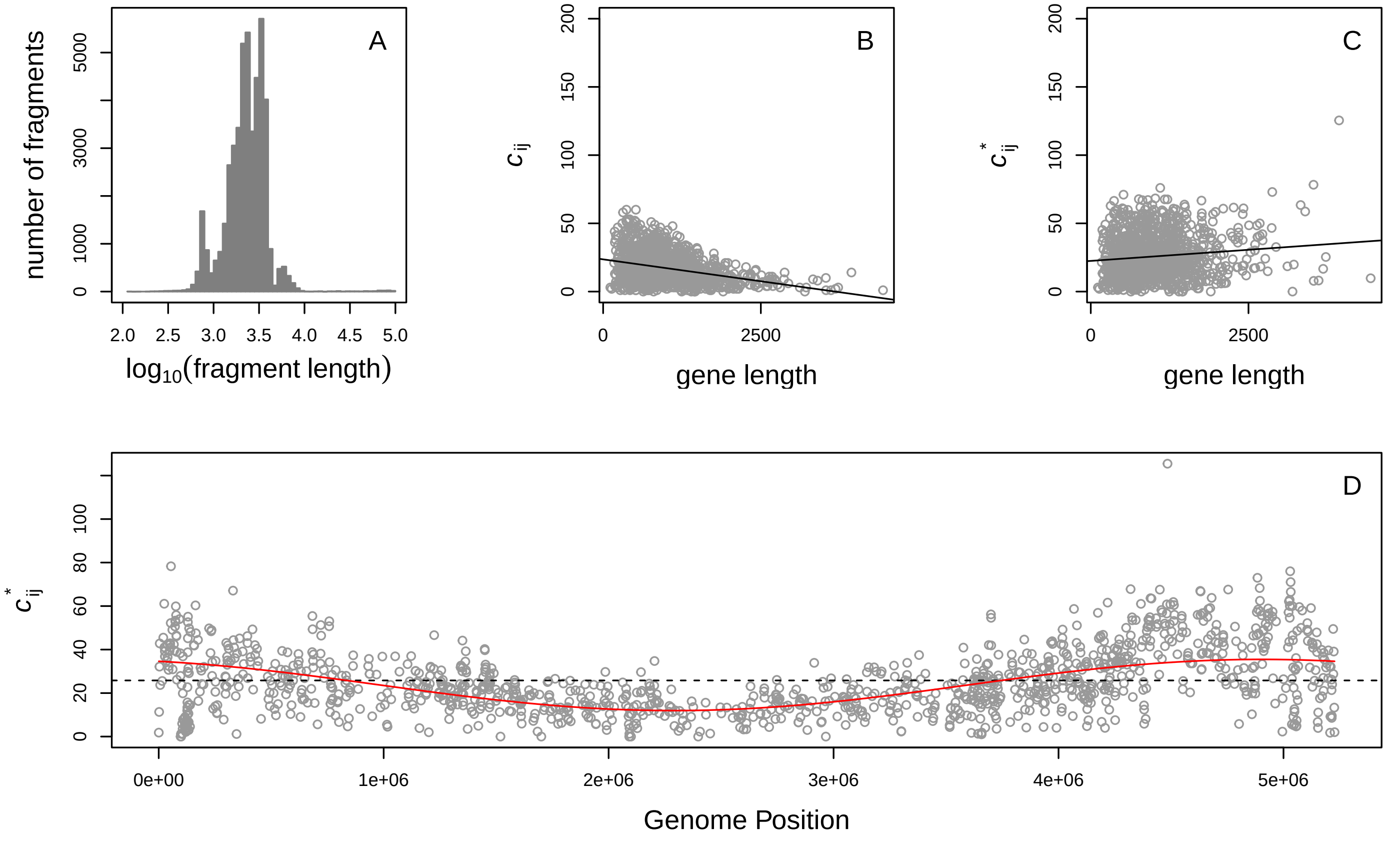

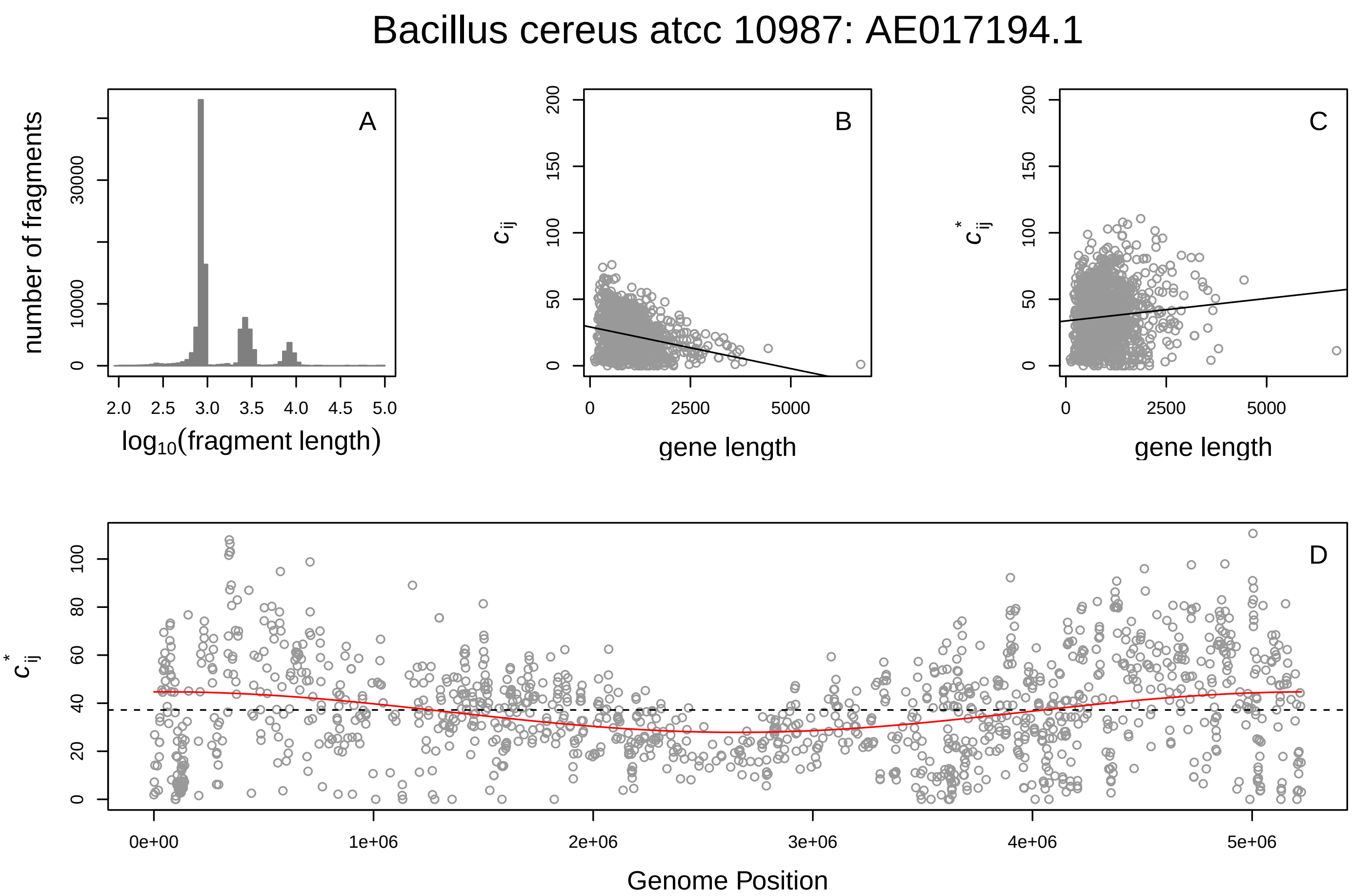
Bacillus cereus atcc 10987: AE017194.1
A
number of fragments
log10(fragment length)
B
$c_{ij}$
gene length
C
$c^*_{ij}$
gene length
D
$c^*_{ij}$
Genome Position

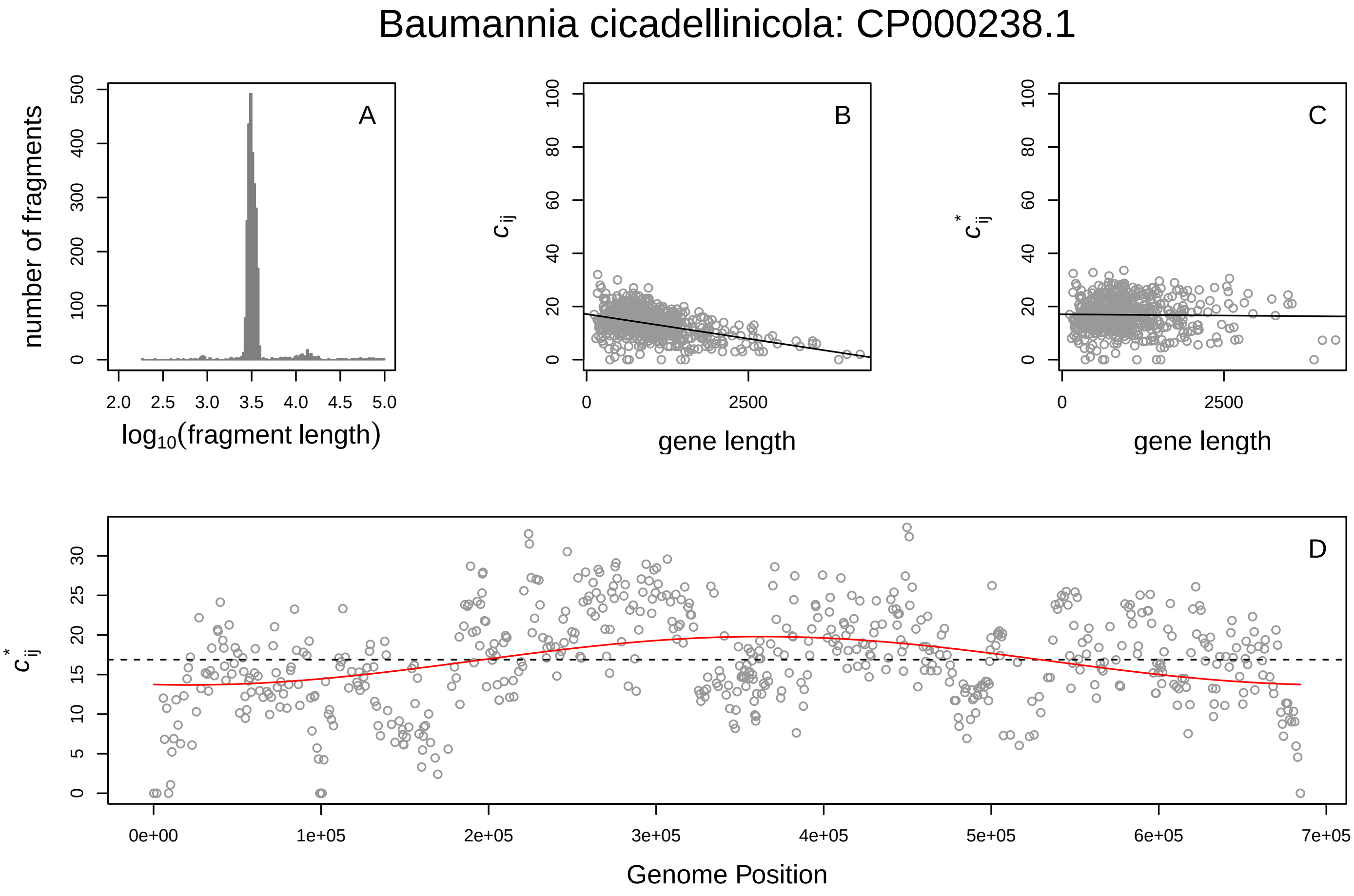
Baumannia cicadellinicola: CP000238.1
A
number of fragments
log10(fragment length)
B
$c_{ij}$
gene length
C
$c^*_{ij}$
gene length
D
$c^*_{ij}$
Genome Position

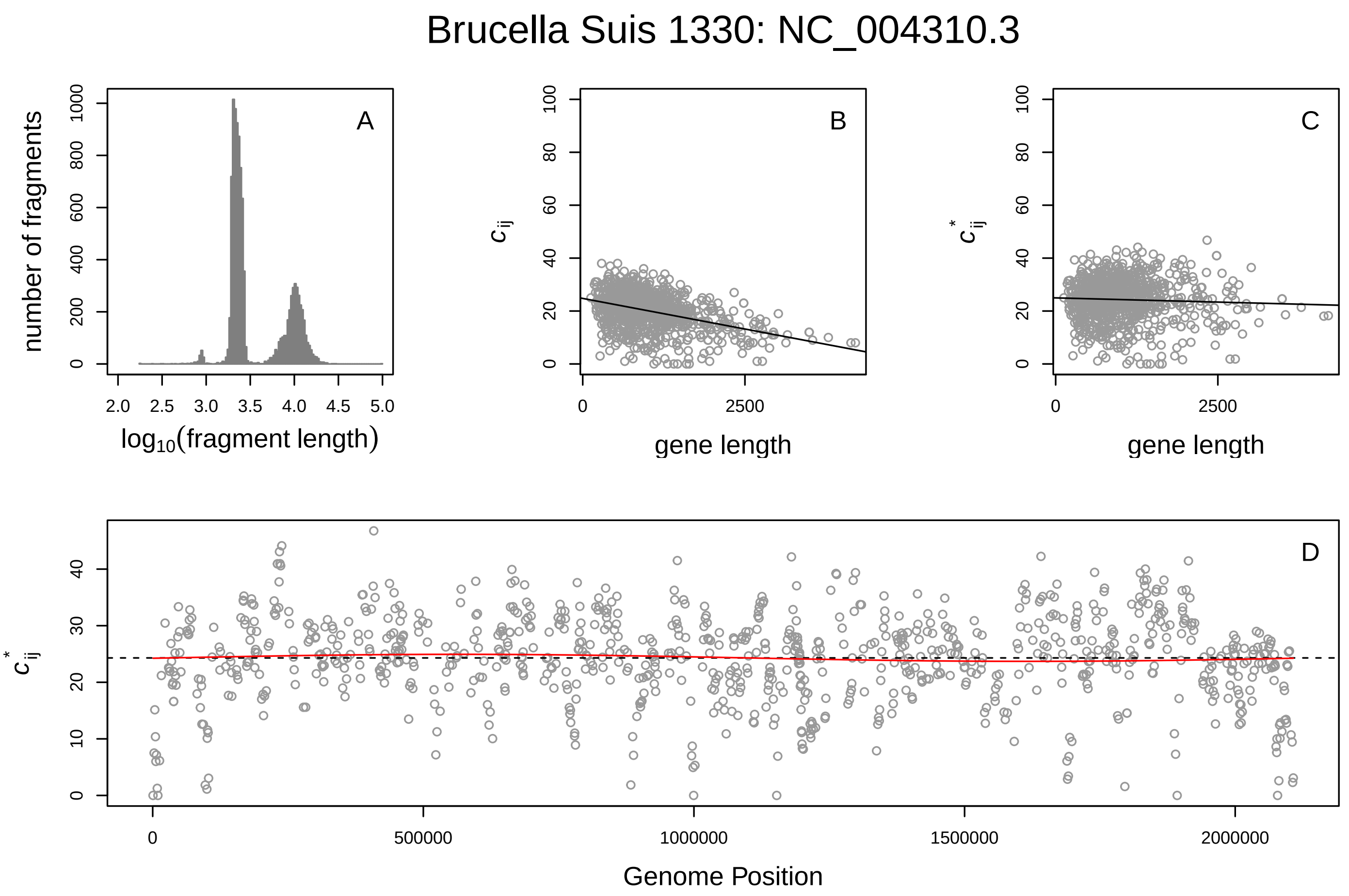
Brucella Suis 1330: NC_004310.3
A
B
C
D
number of fragments
log10(fragment length)
gene length
Genome Position
c_ij
c*_ij

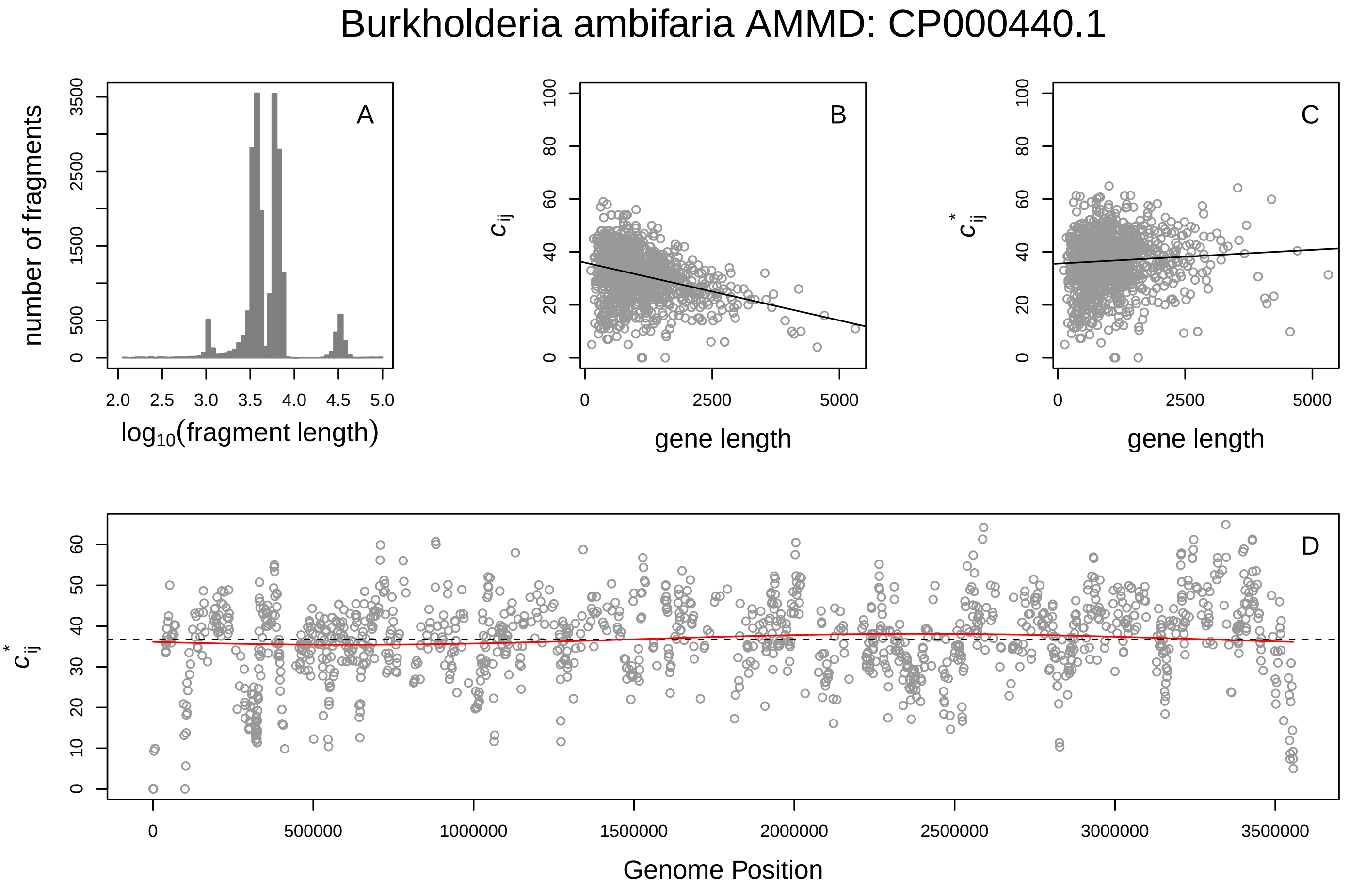
Burkholderia ambifaria AMMD: CP000440.1
A
B
C
D
number of fragments
log10(fragment length)
gene length
Genome Position
c_ij
c*_ij

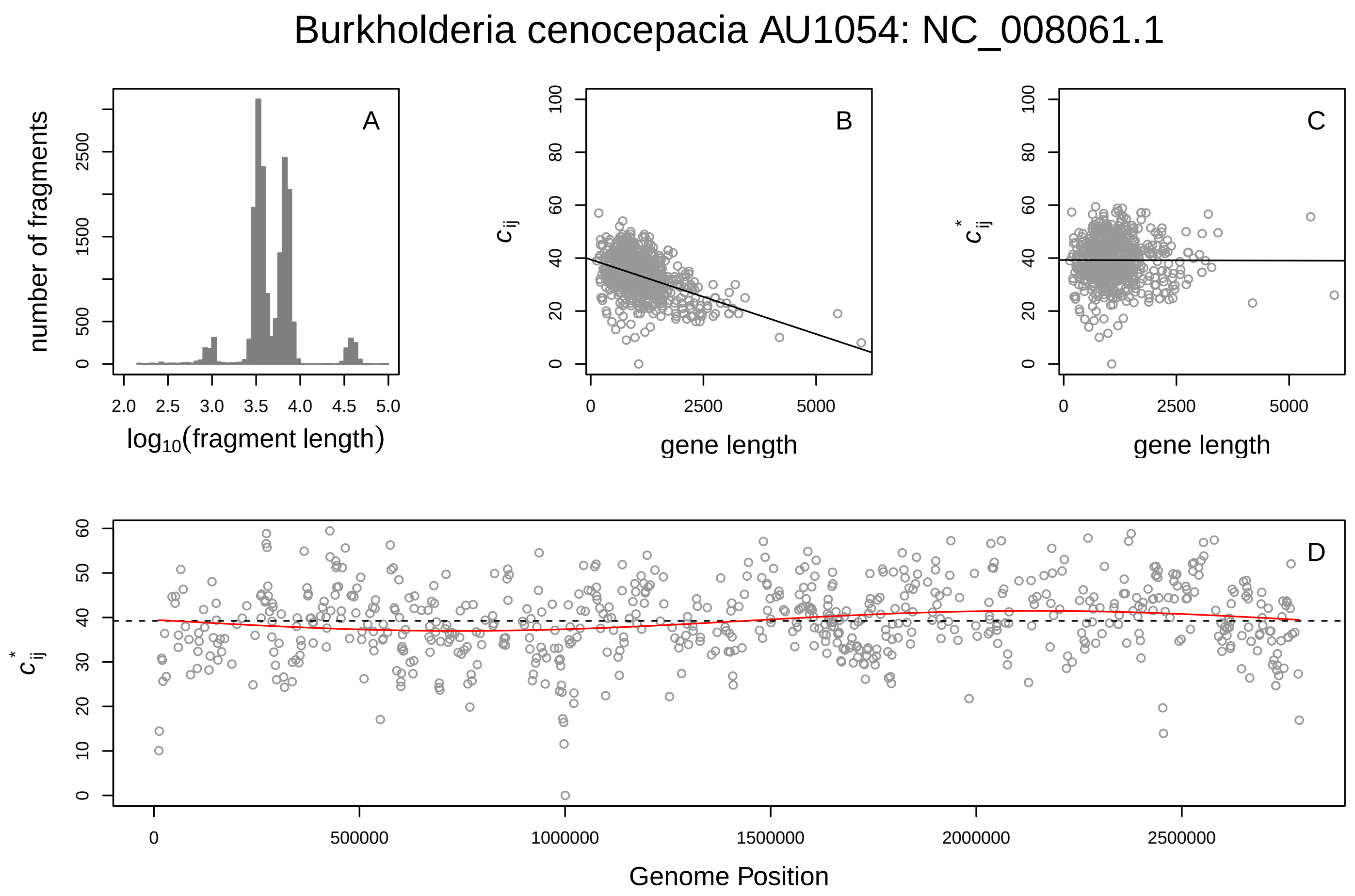

Burkholderia cenocepacia AU1054: NC_008061.1
A
number of fragments
log10(fragment length)
B
$c_{ij}$
gene length
C
$c^*_{ij}$
gene length
D
$c^*_{ij}$
Genome Position


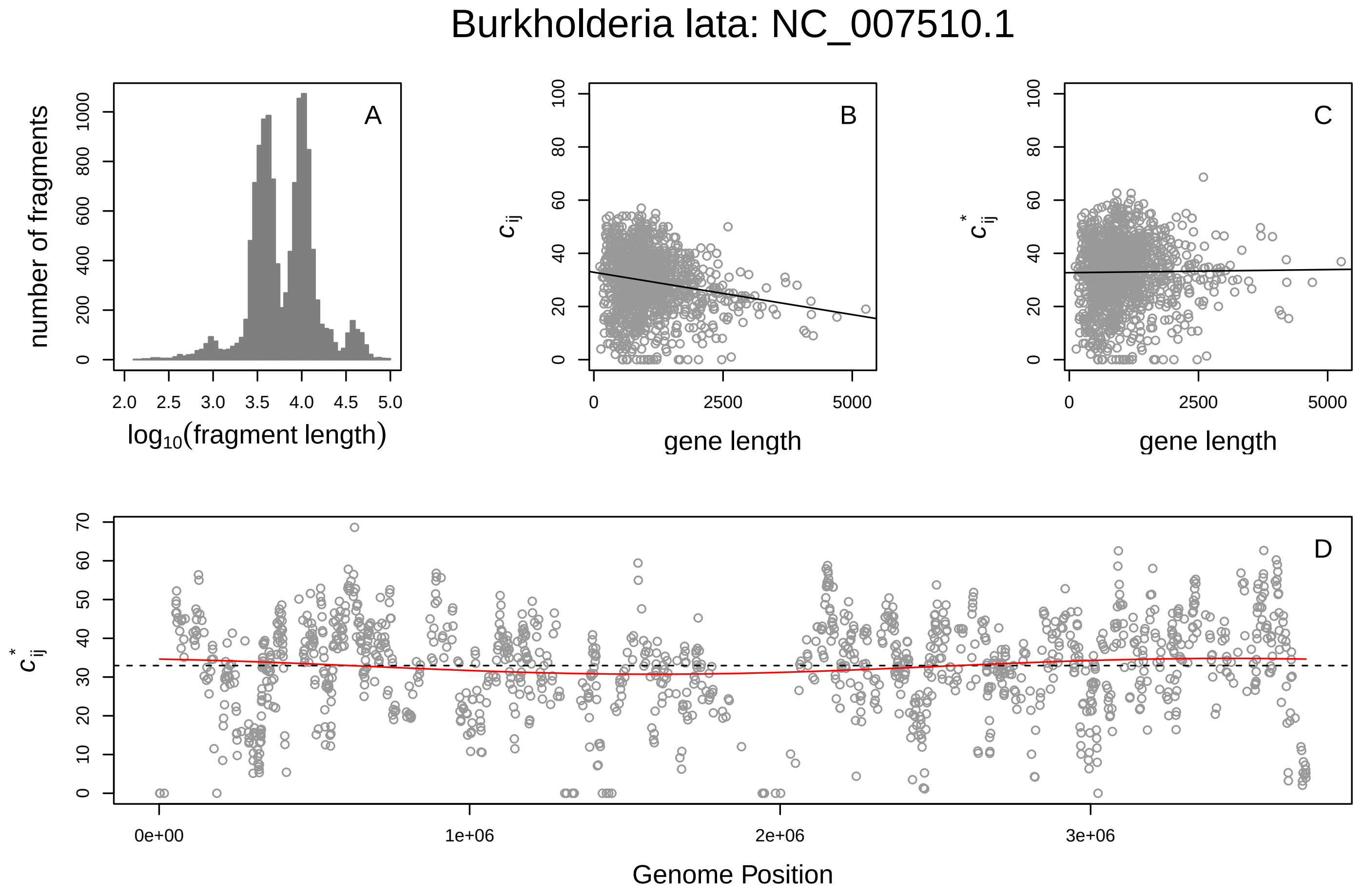

Burkholderia lata: NC_007510.1
A
number of fragments
log10(fragment length)
B
$c_{ij}$
gene length
C
$c^*_{ij}$
gene length
D
$c^*_{ij}$
Genome Position

Burkholderia lata: NC_007511.1

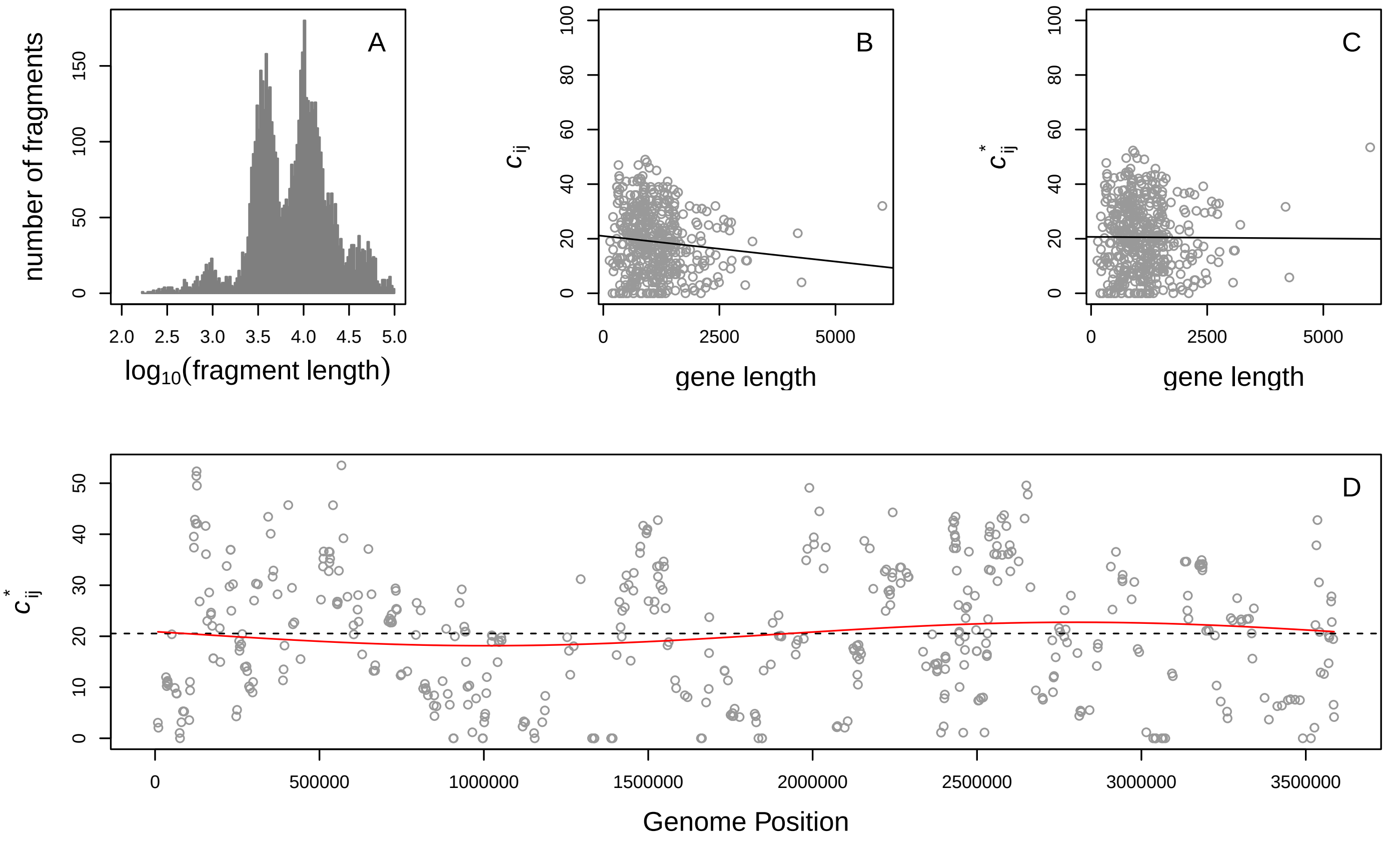


Burkholderia mallei atcc 23344: NZ_CP009727.1

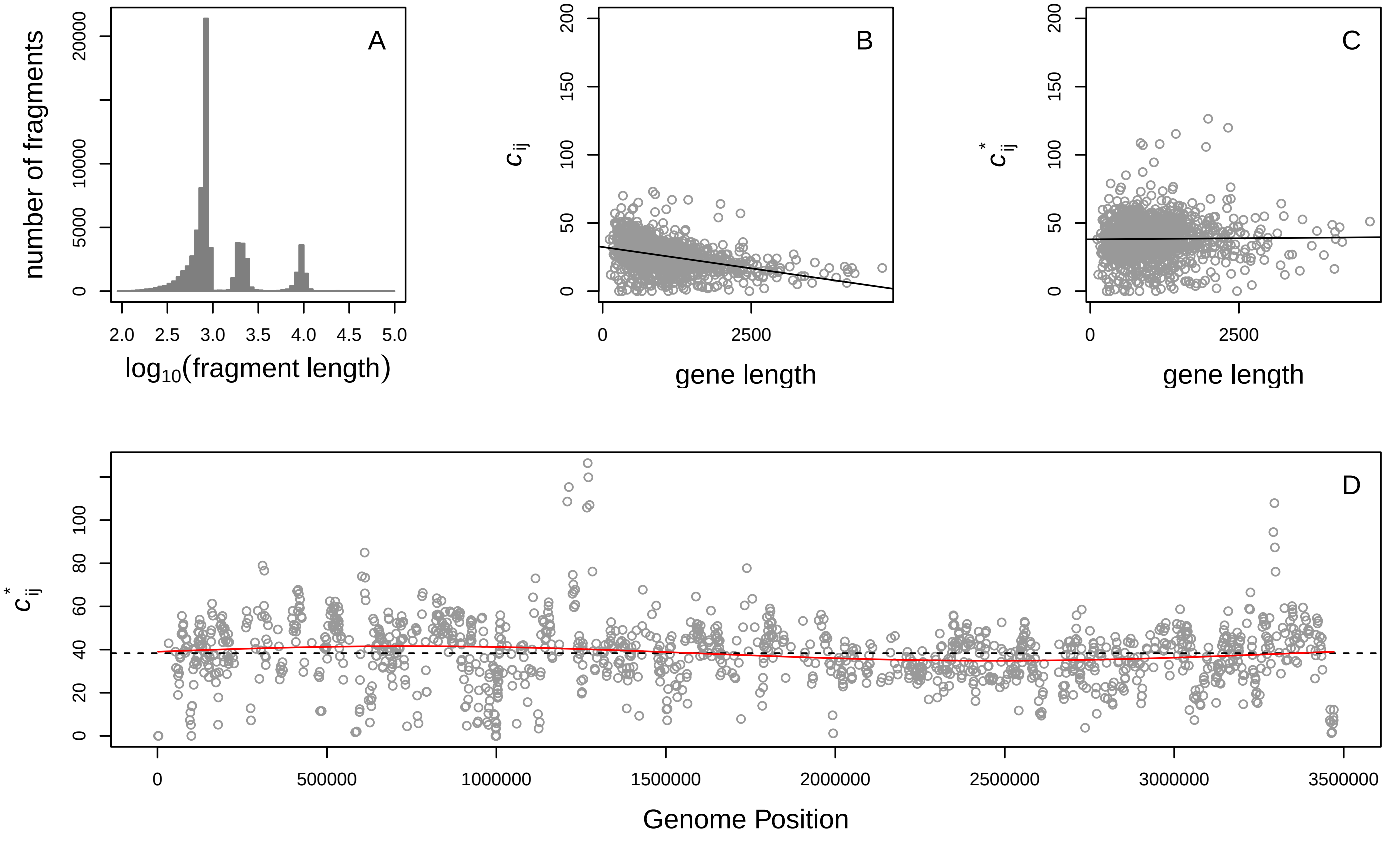

Burkholderia thailandensis e264: CP000086.1

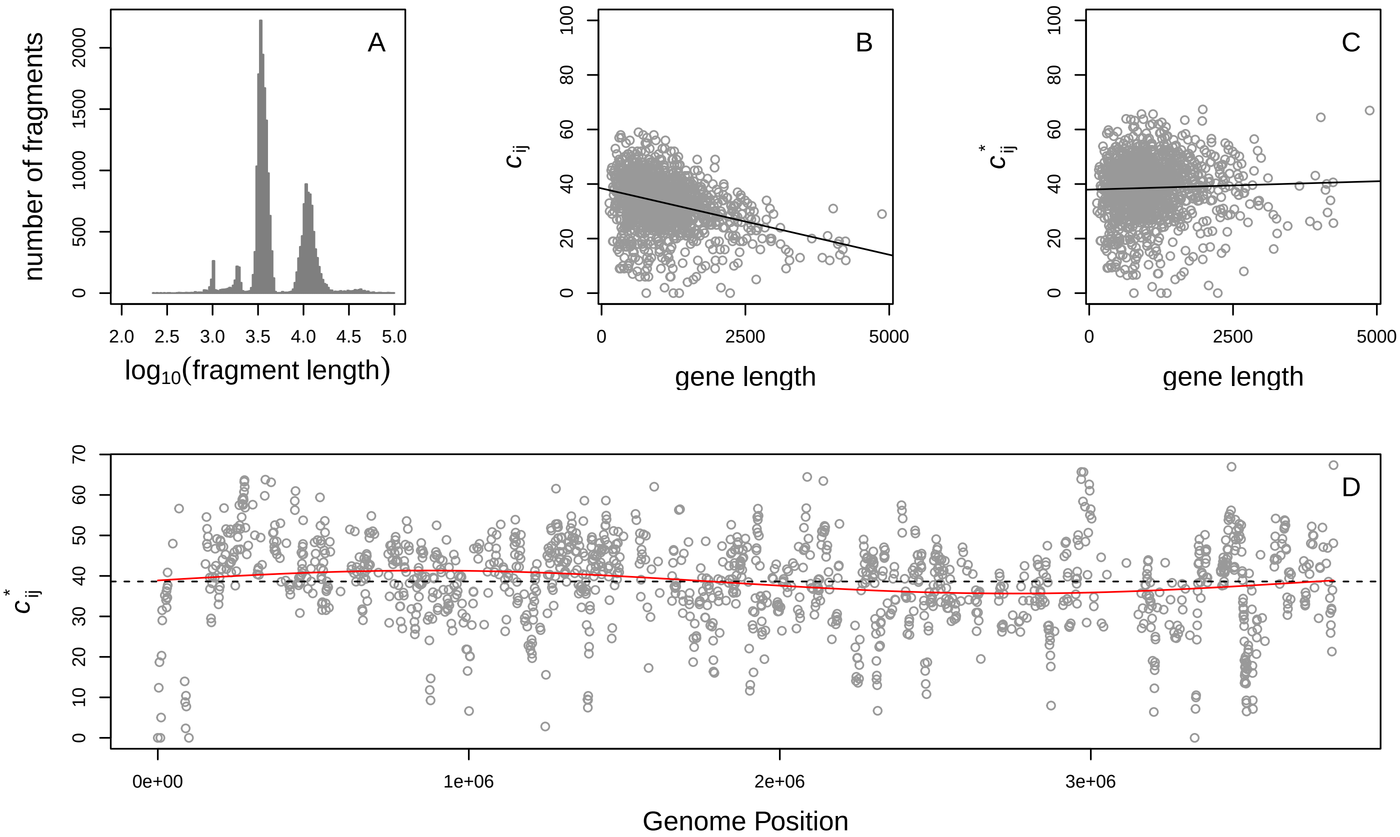


Campylobacter jejuni RM1221: NC_003912.7

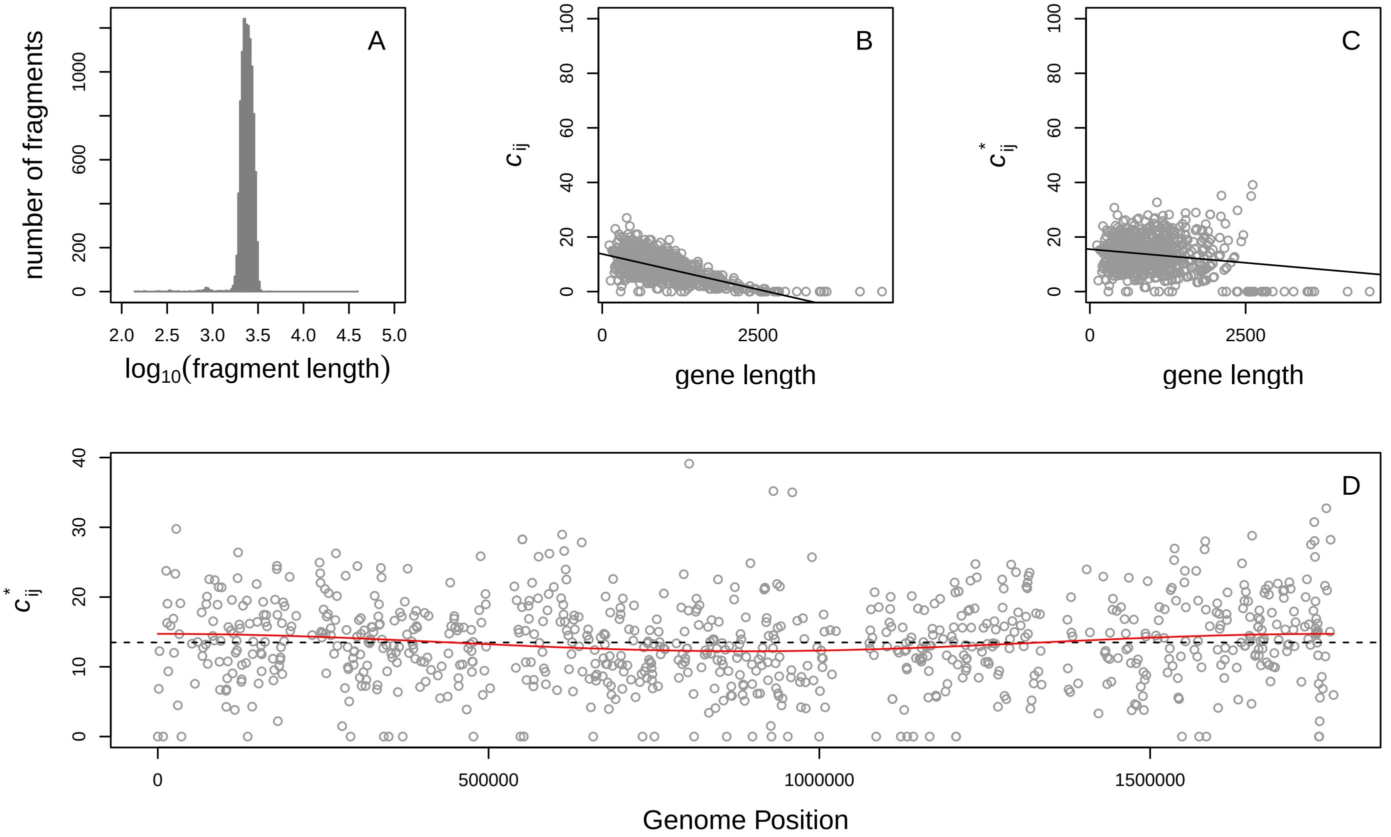

#### Carboxydothermus hydrogenoformans z−2901: CP000141.1

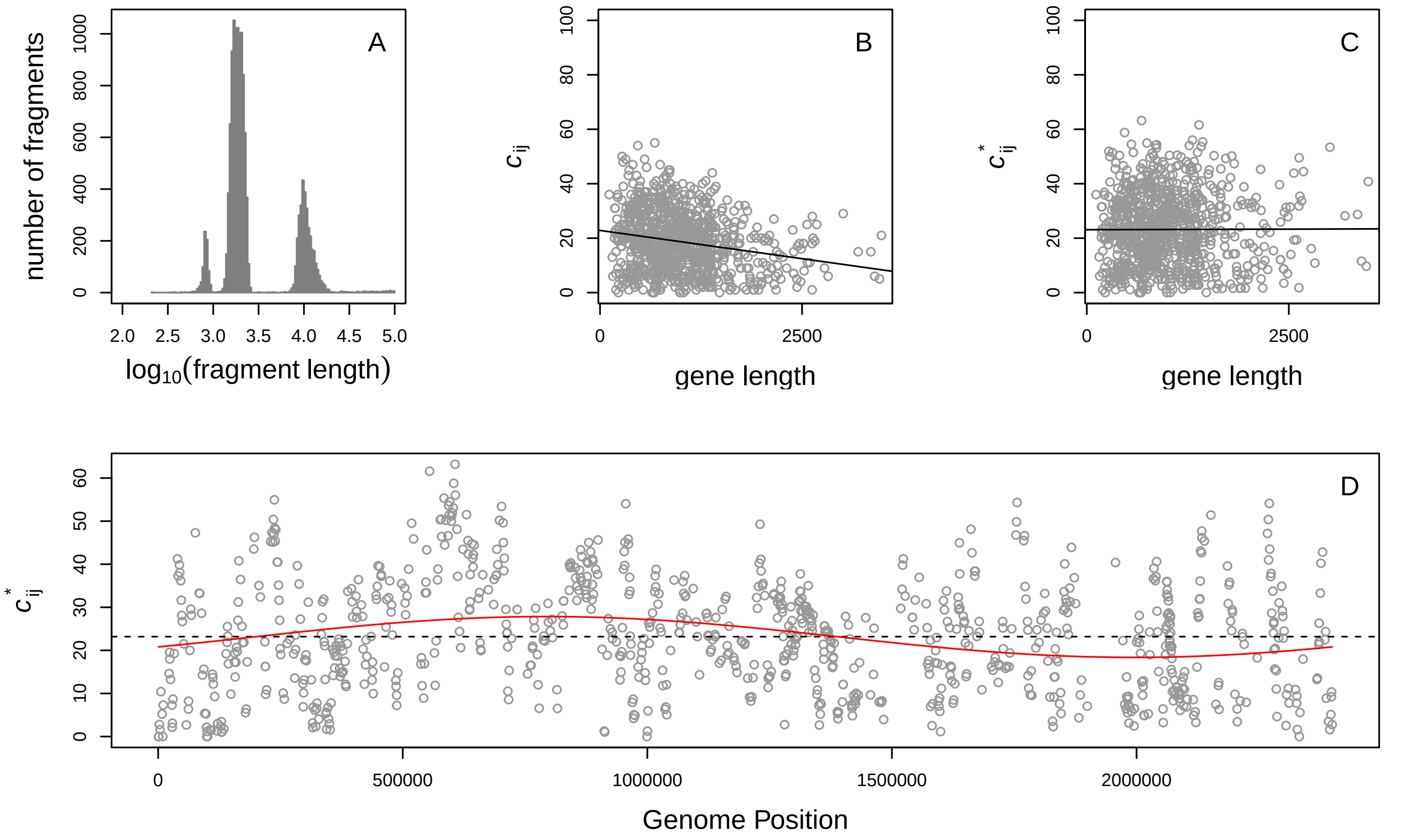


#### Chlamydophila caviae gpic: NC_003361.3

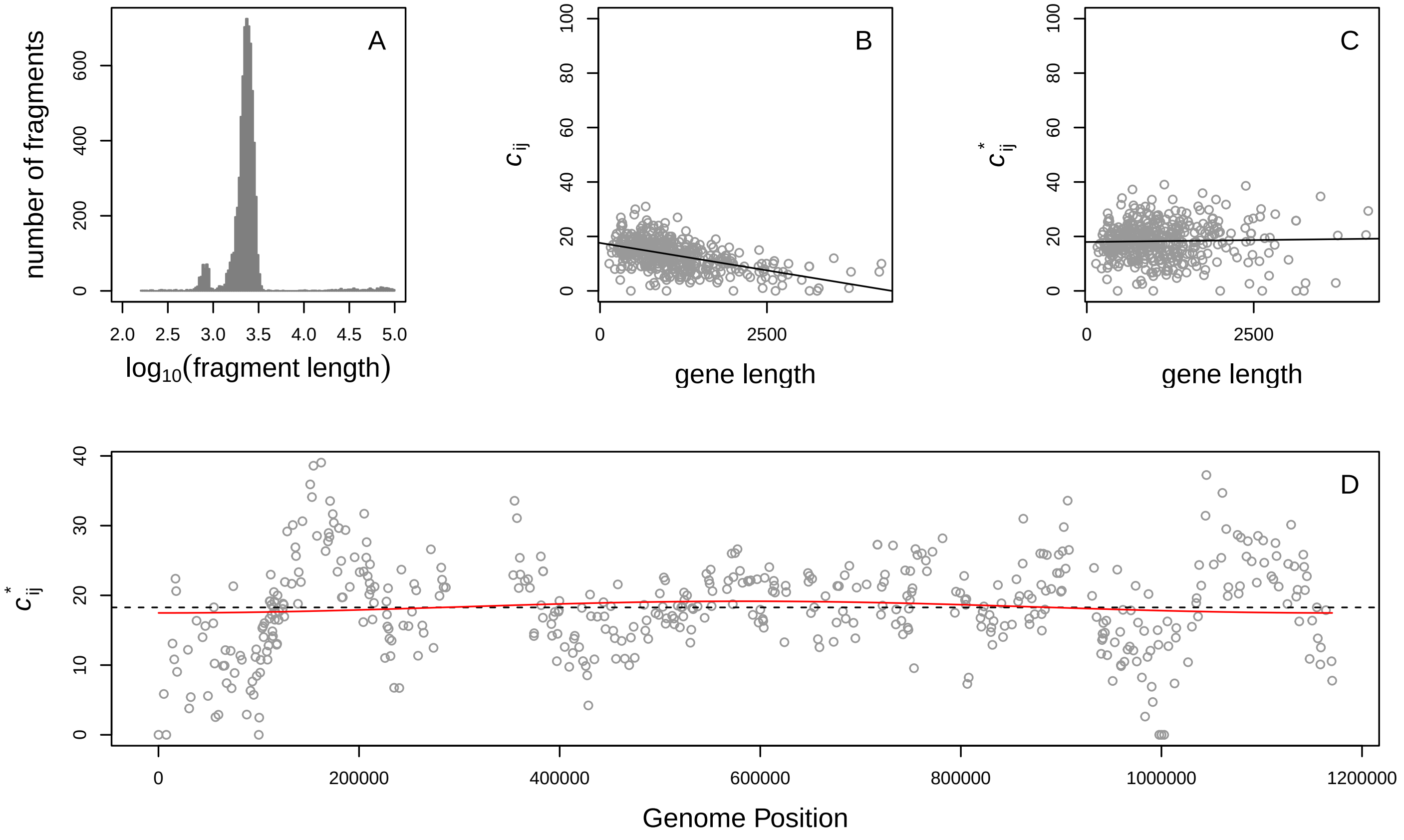

### Chlorobium tepidum tls: NC_002932.3

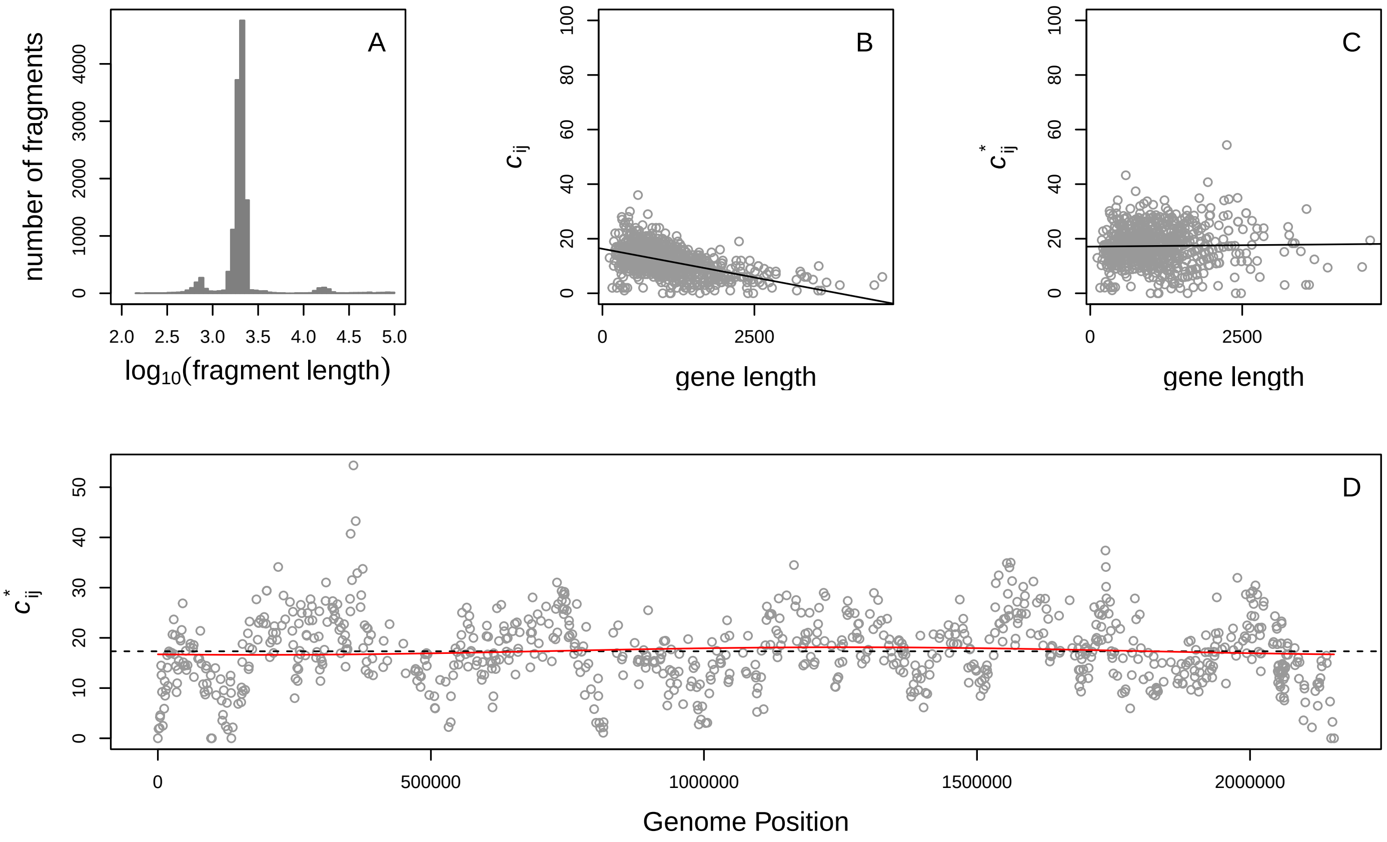


### Colwellia psychrerythraea 34h: NC_003910.7

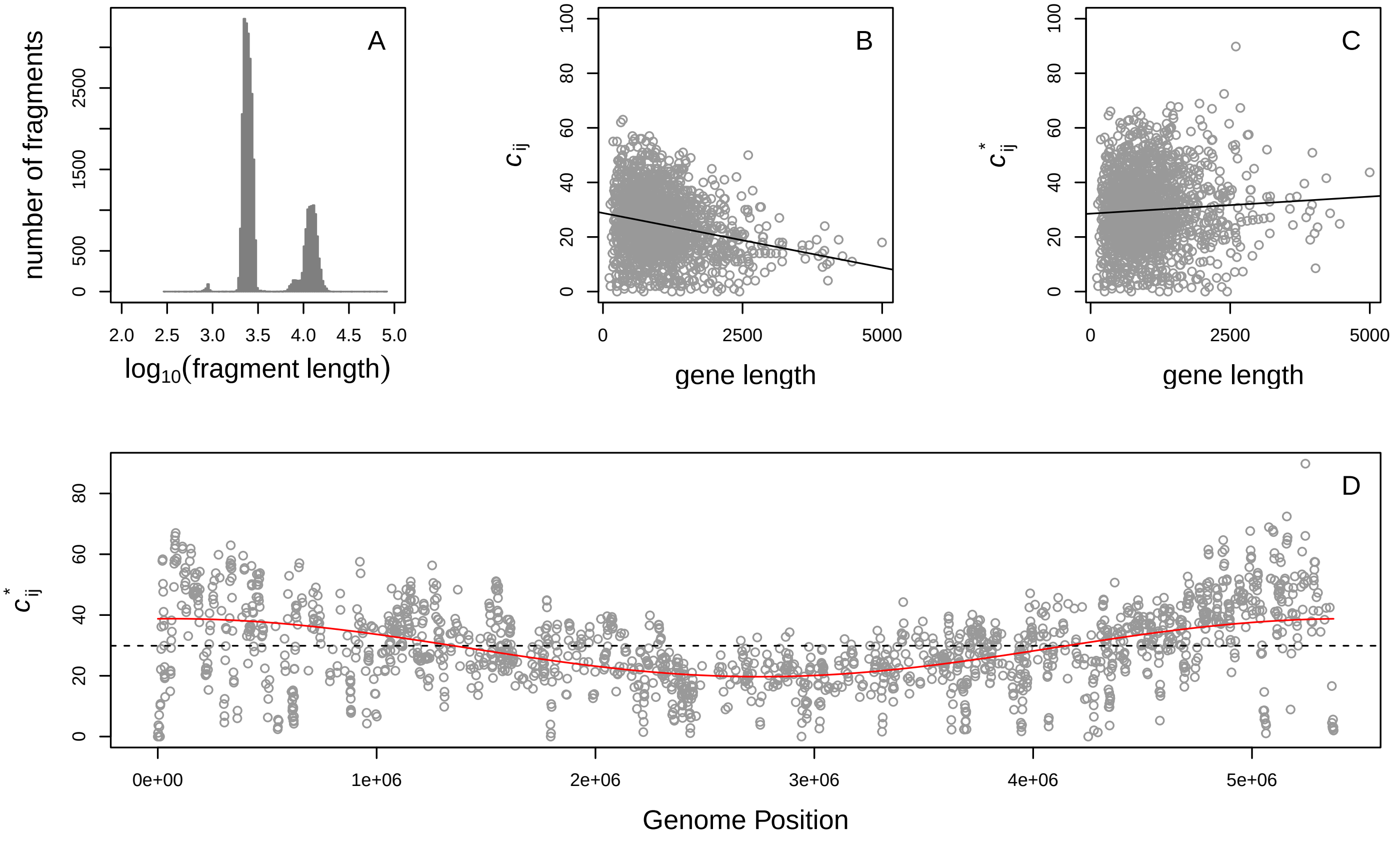

#### Coxiella burnetii rsa 493: NC_002971.4

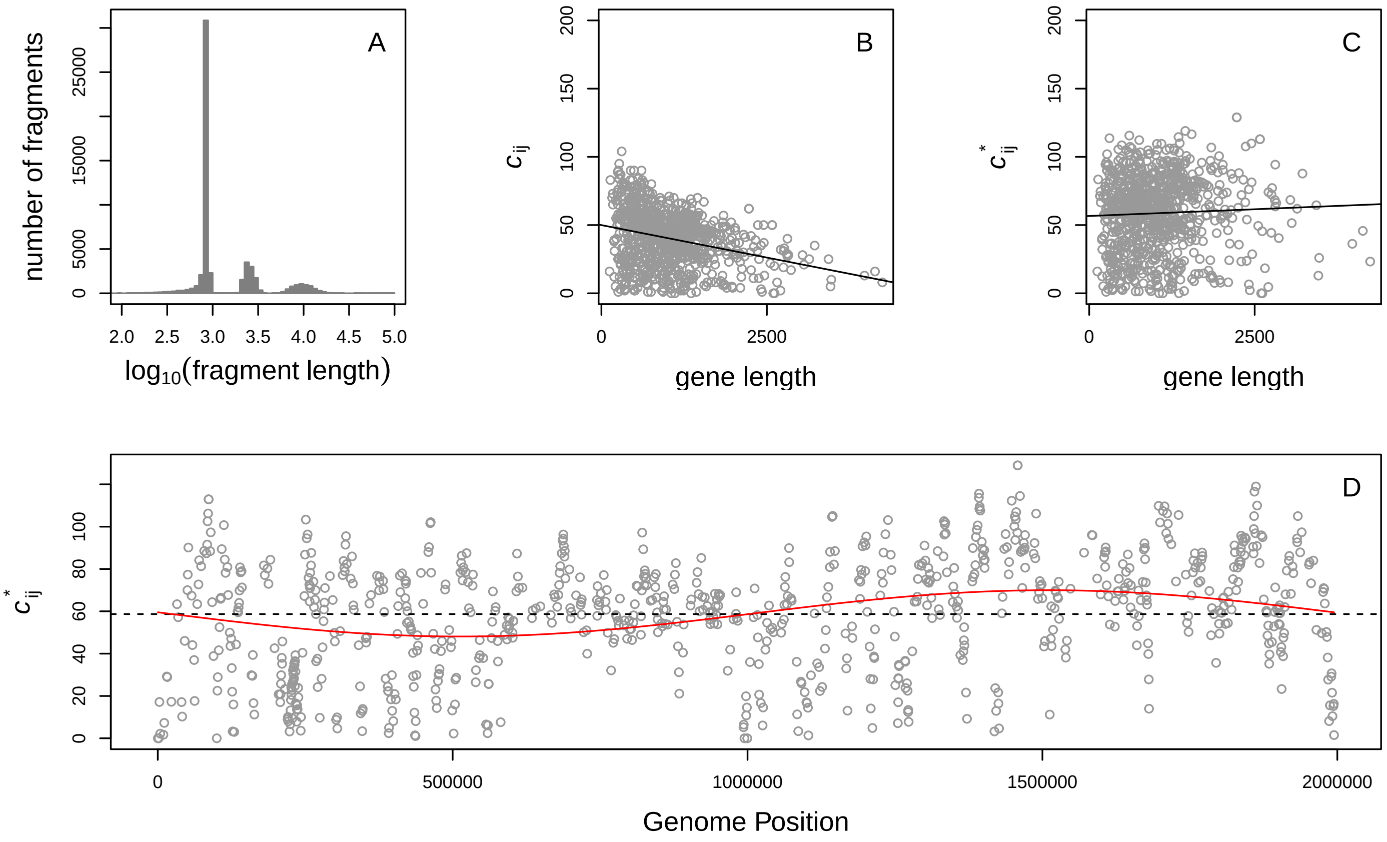


#### Dechloromonas aromatica rcb: NC_007298.1

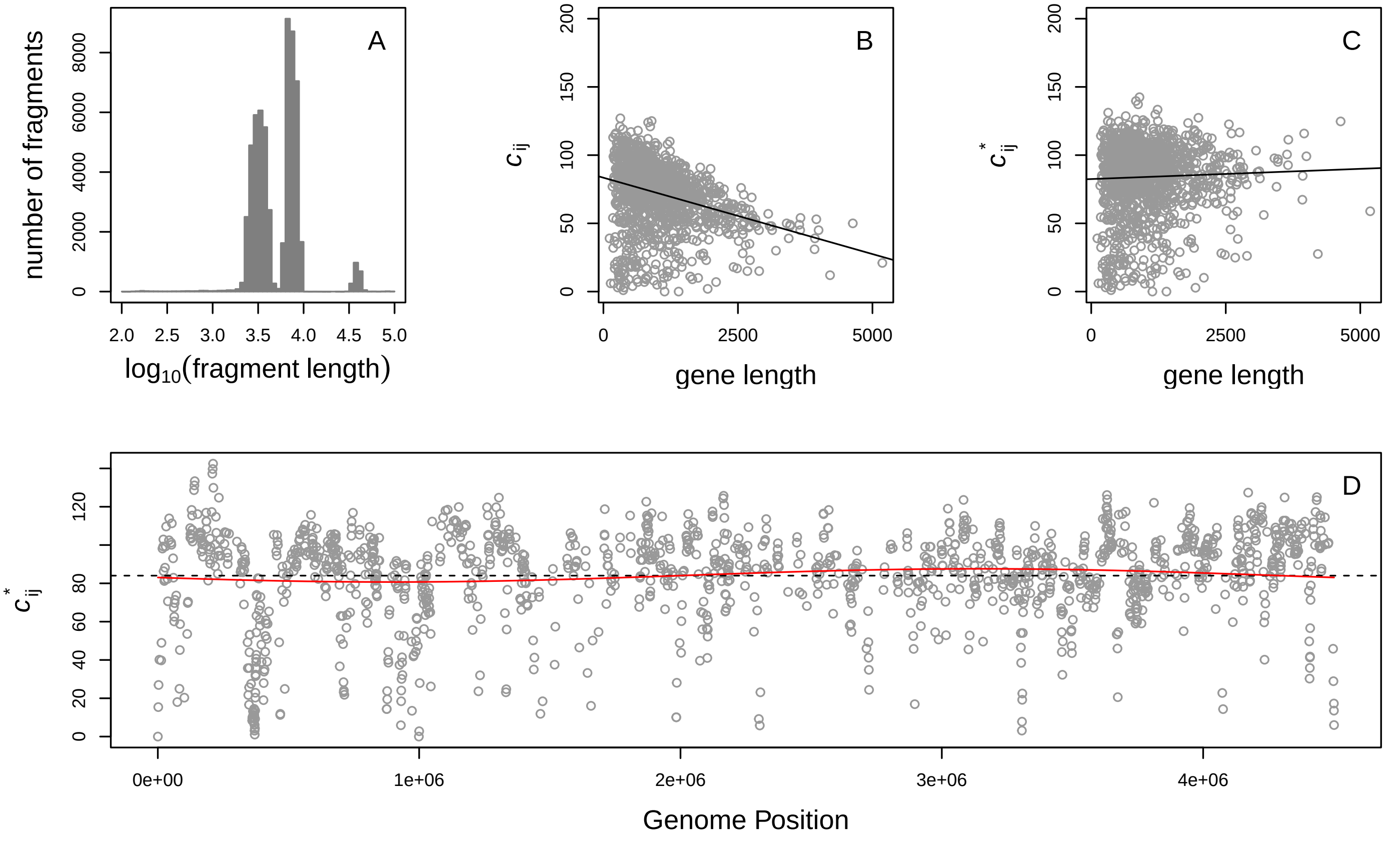

#### Dehalococcoides ethenogenes 195: CP000027.1

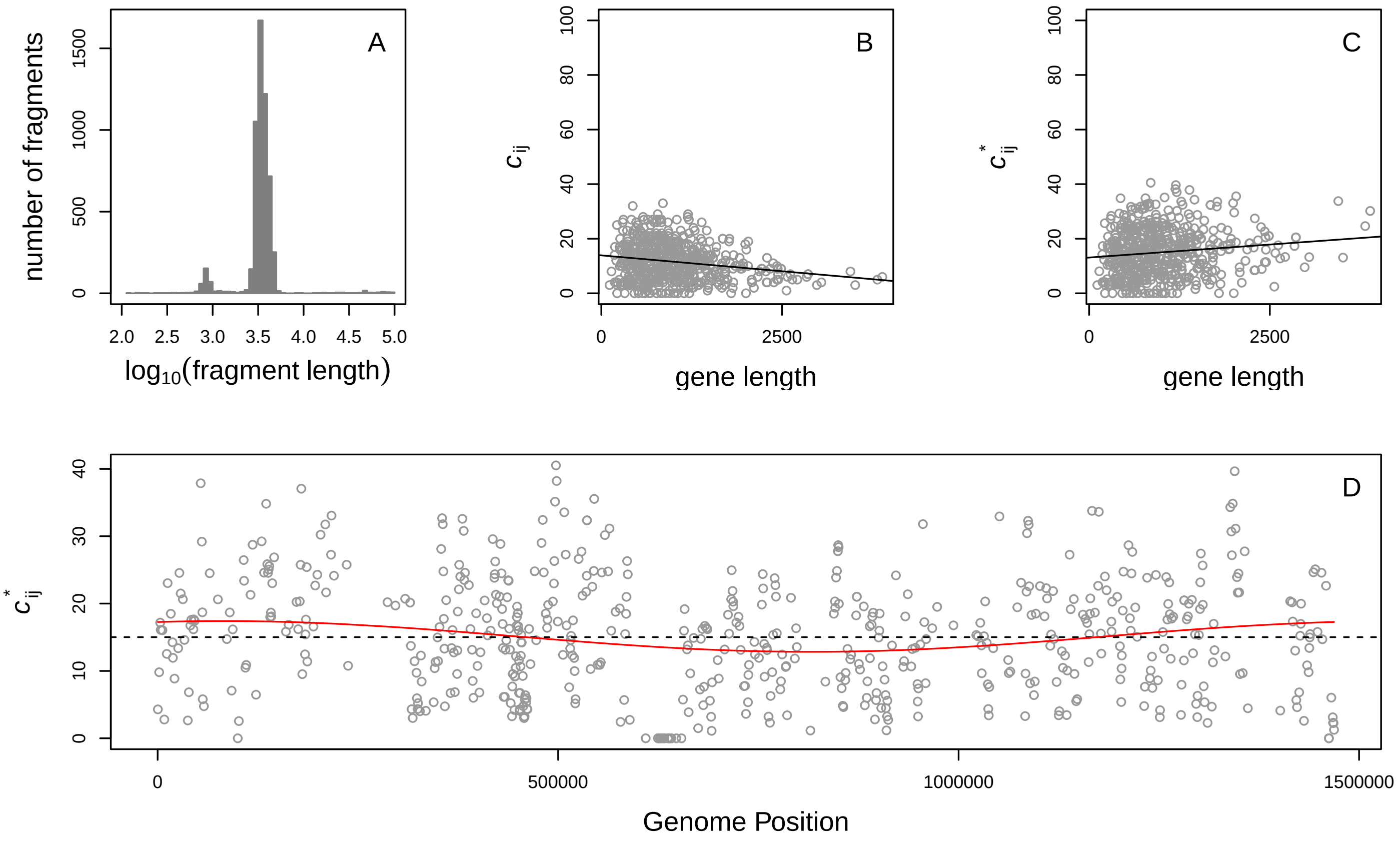


#### Deinococcus geothermalis dms 11300: CP000359.1

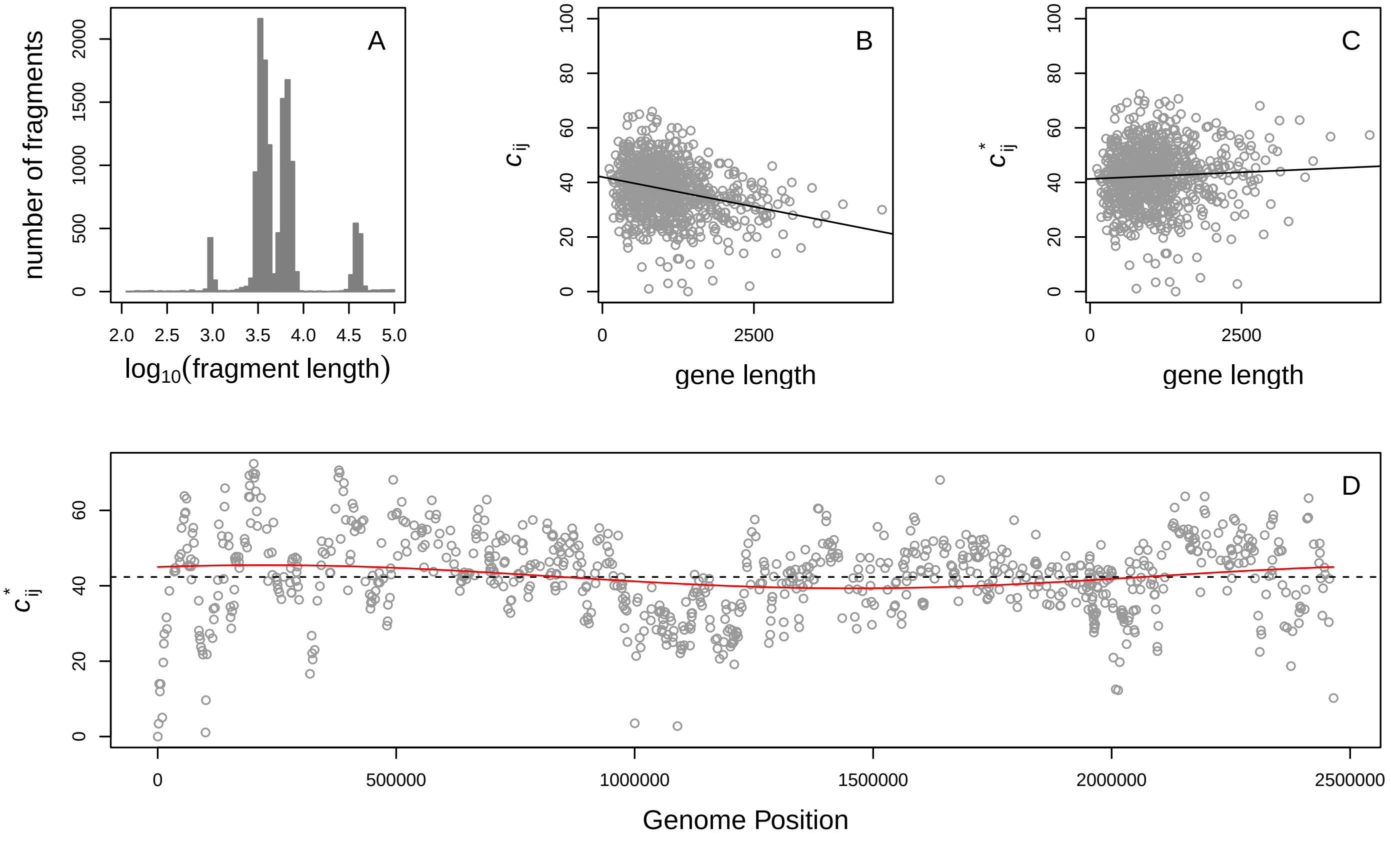

Ehrlichia canis str jake: NC_007354.1

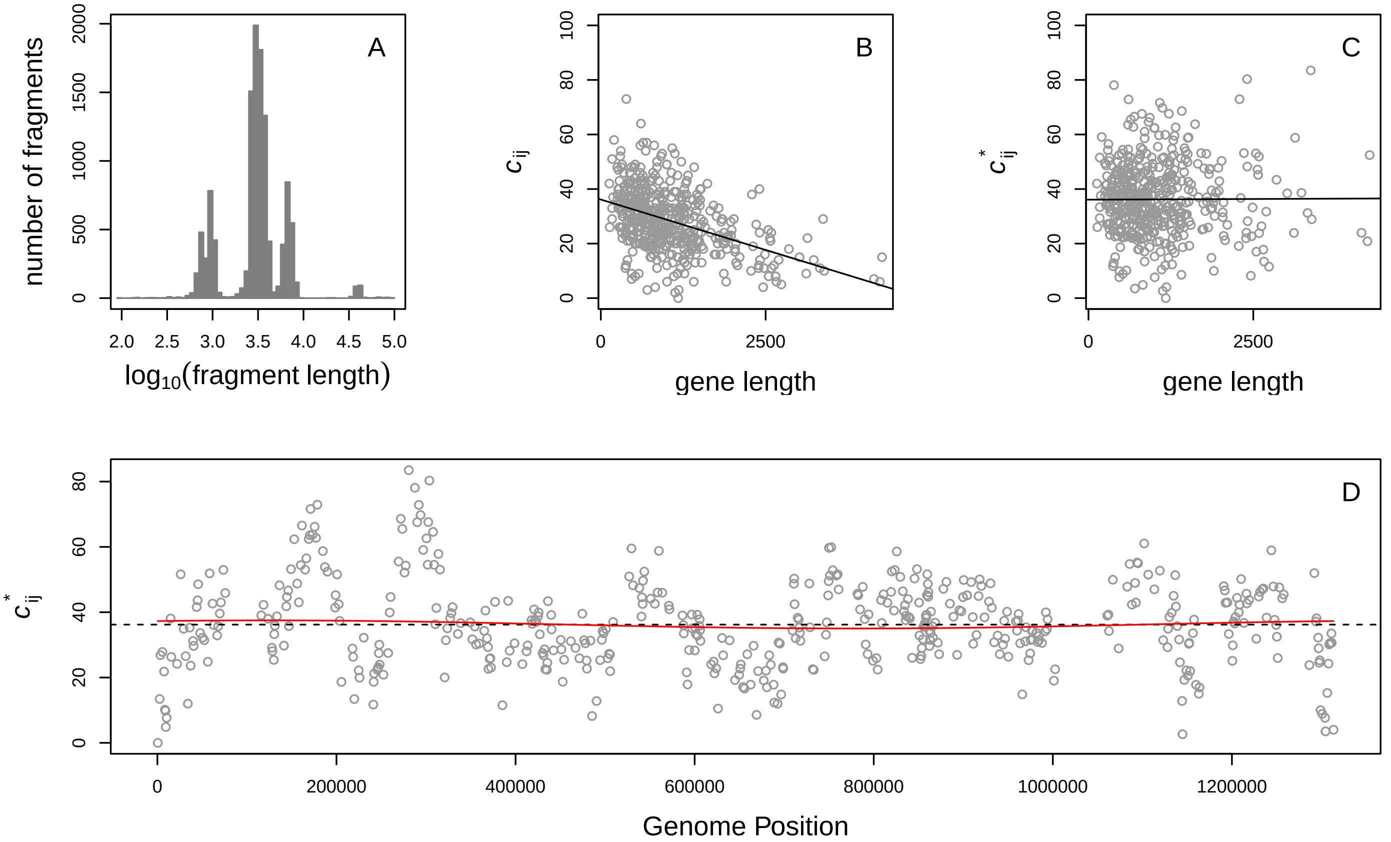


Ehrlichia chaffeensis str Arkansas: CP000236.1

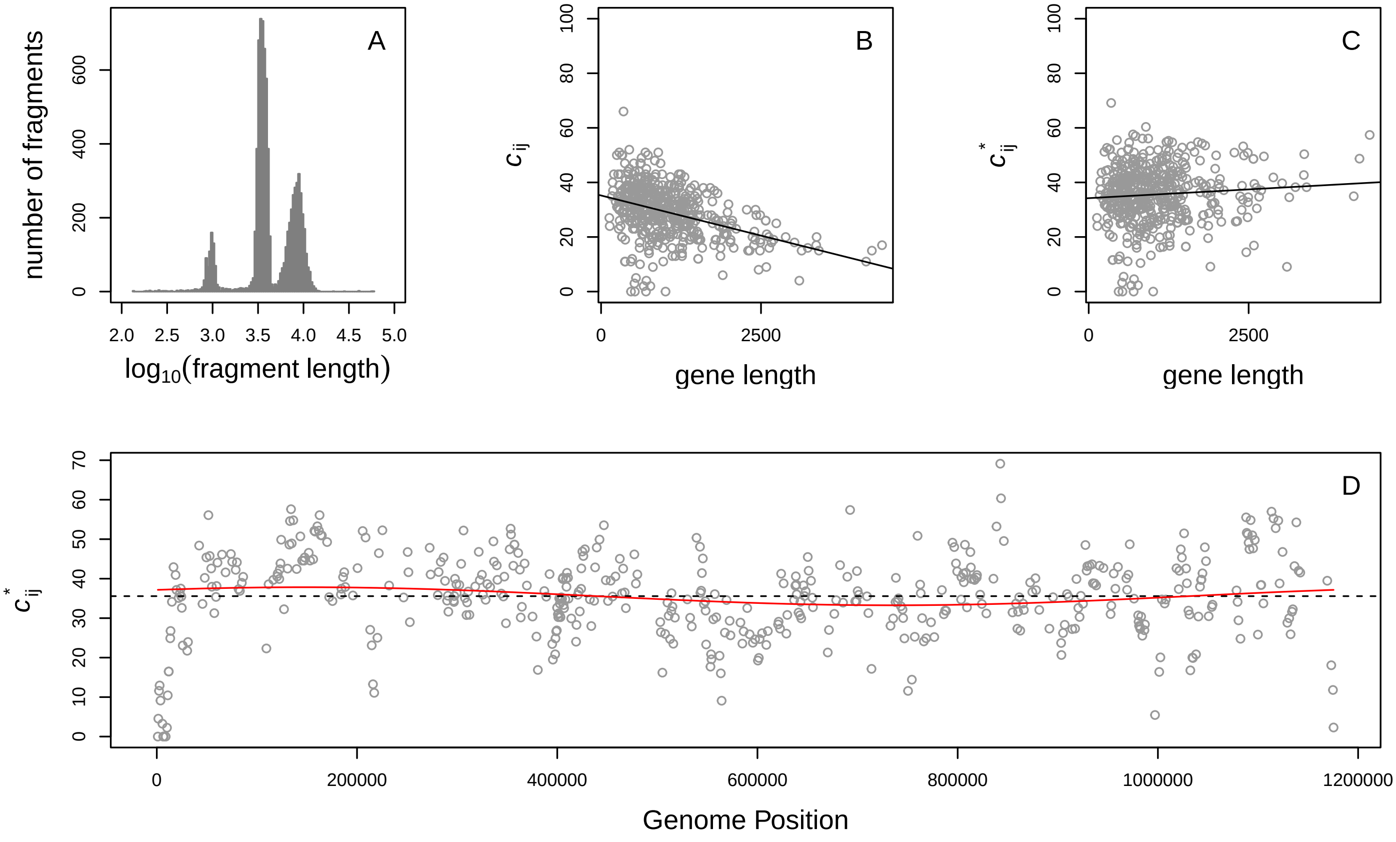

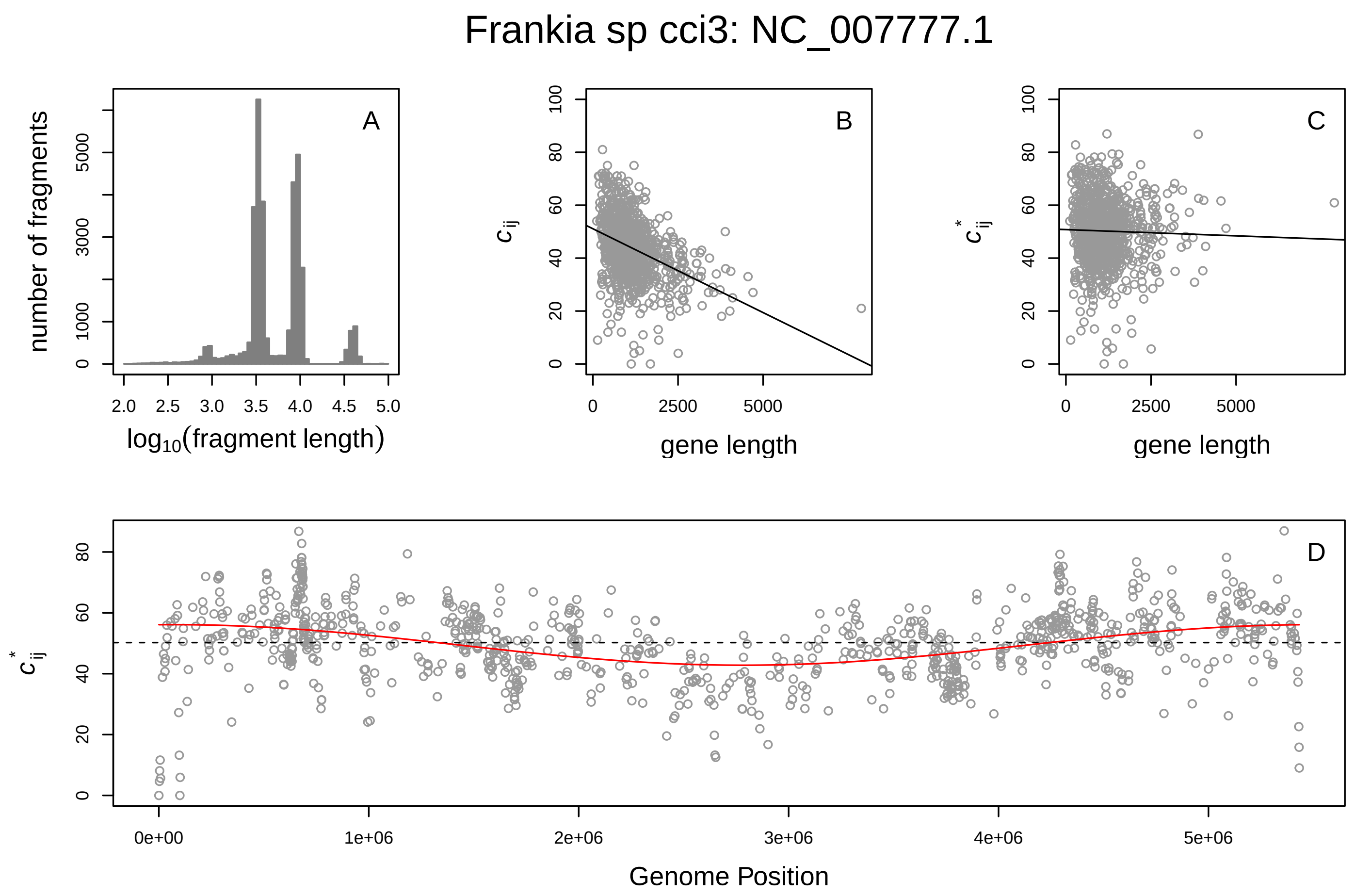
Frankia sp cci3: NC_007777.1
A
B
C
D
number of fragments
log10(fragment length)
gene length
Genome Position

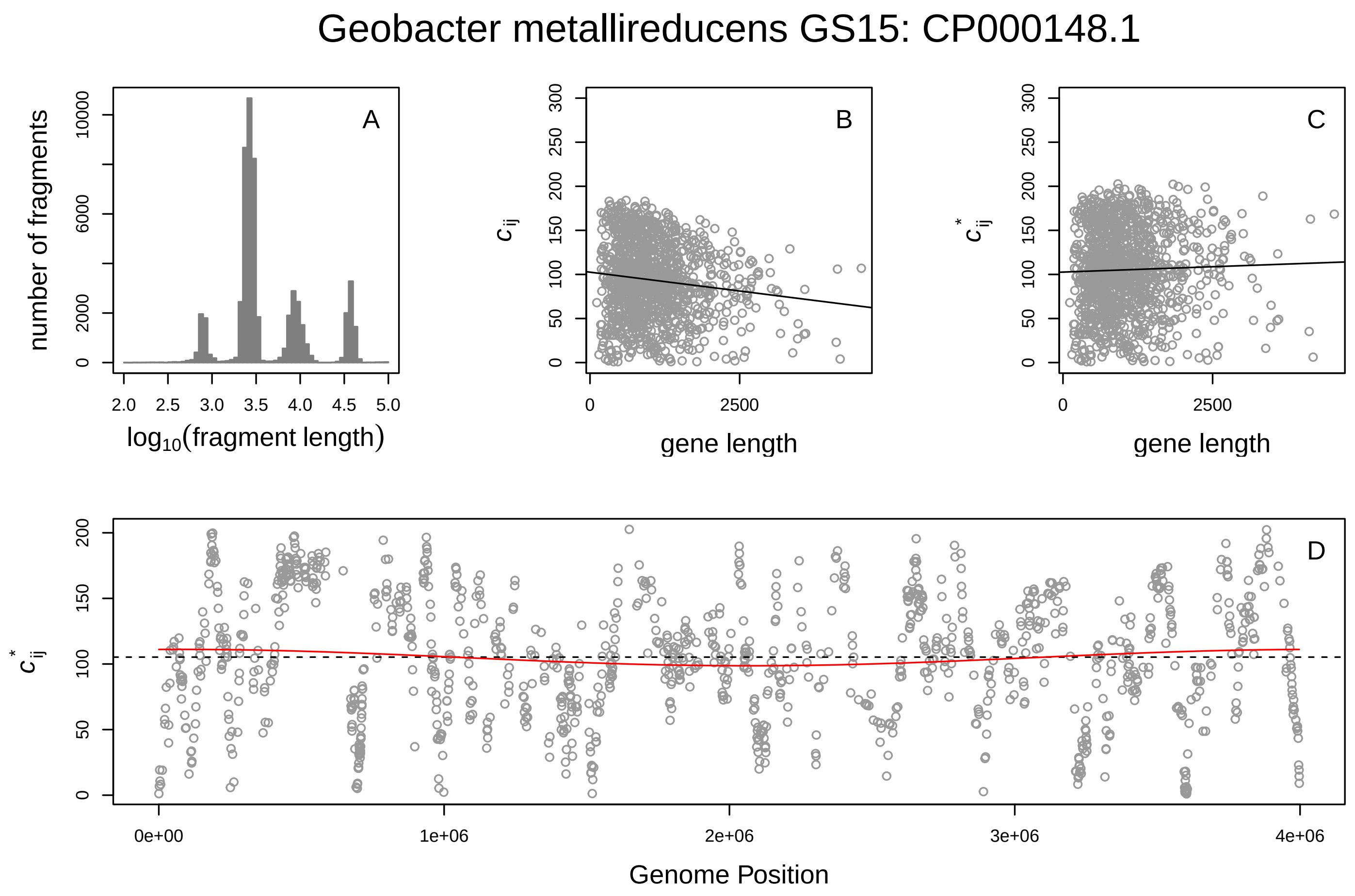
Geobacter metallireducens GS15: CP000148.1
A
B
C
D
number of fragments
log10(fragment length)
gene length
Genome Position

Geobacter sulfurreducens pca: NC_002939.5

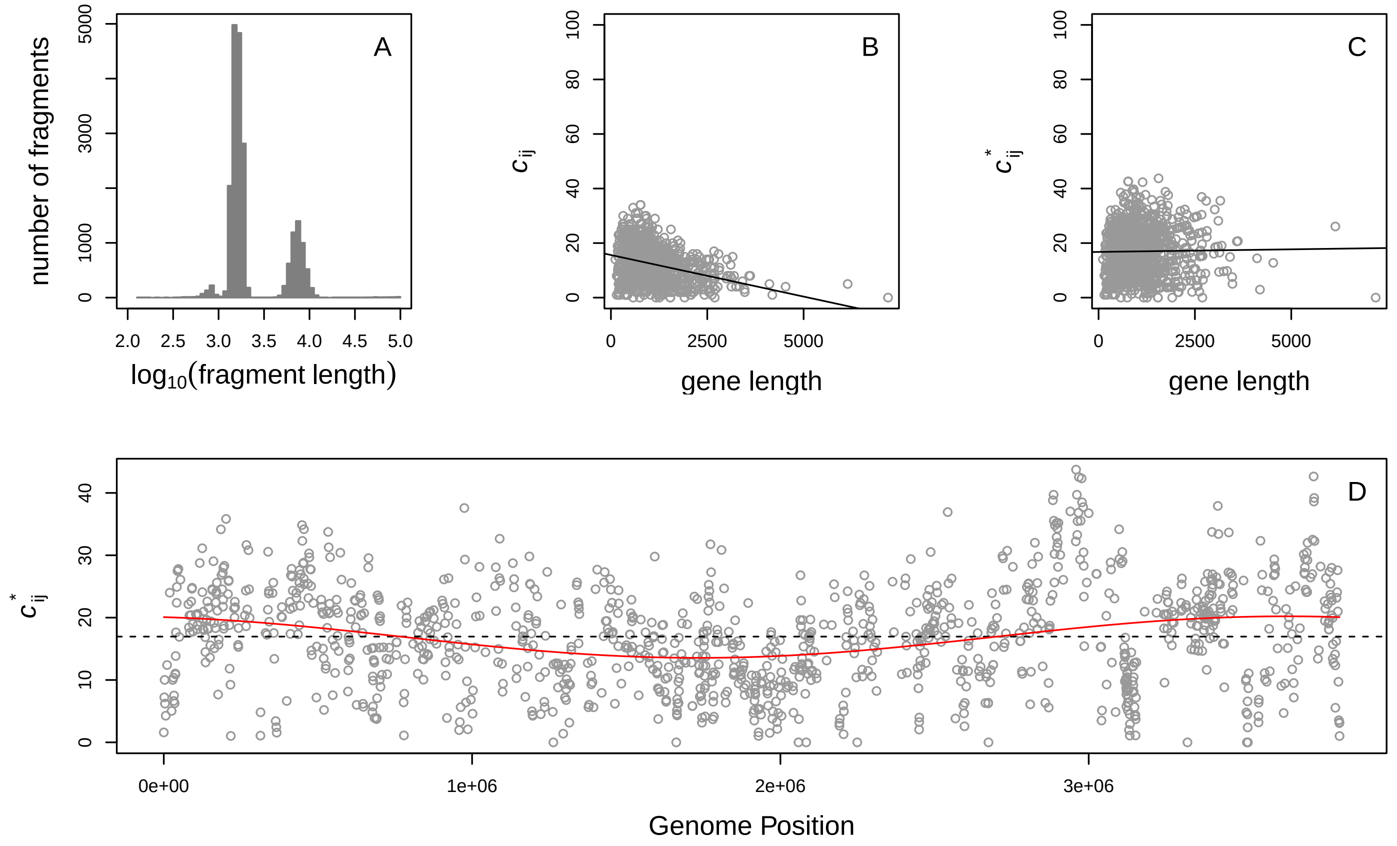


Jannaschia sp. CCS1: CP000264.1

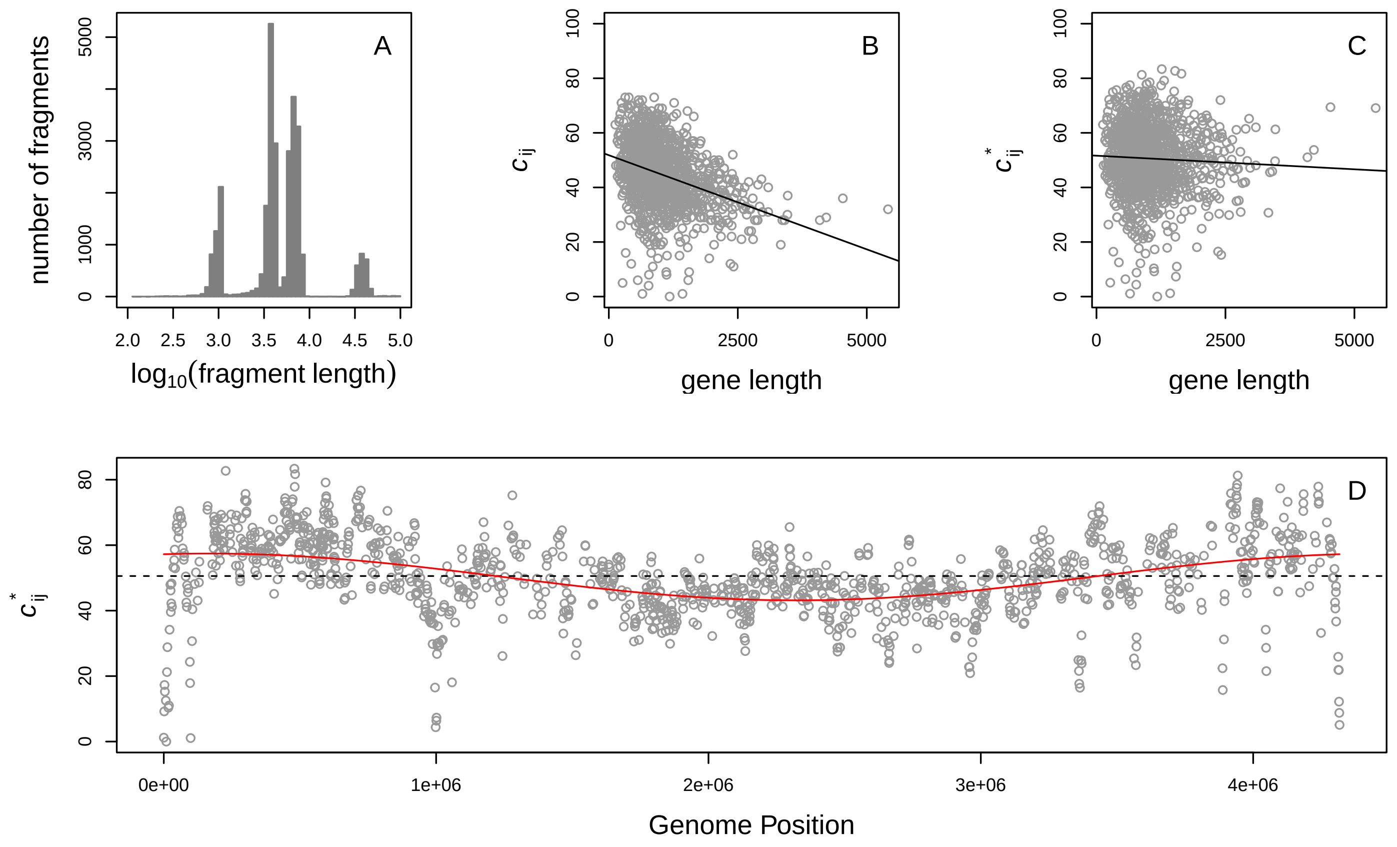

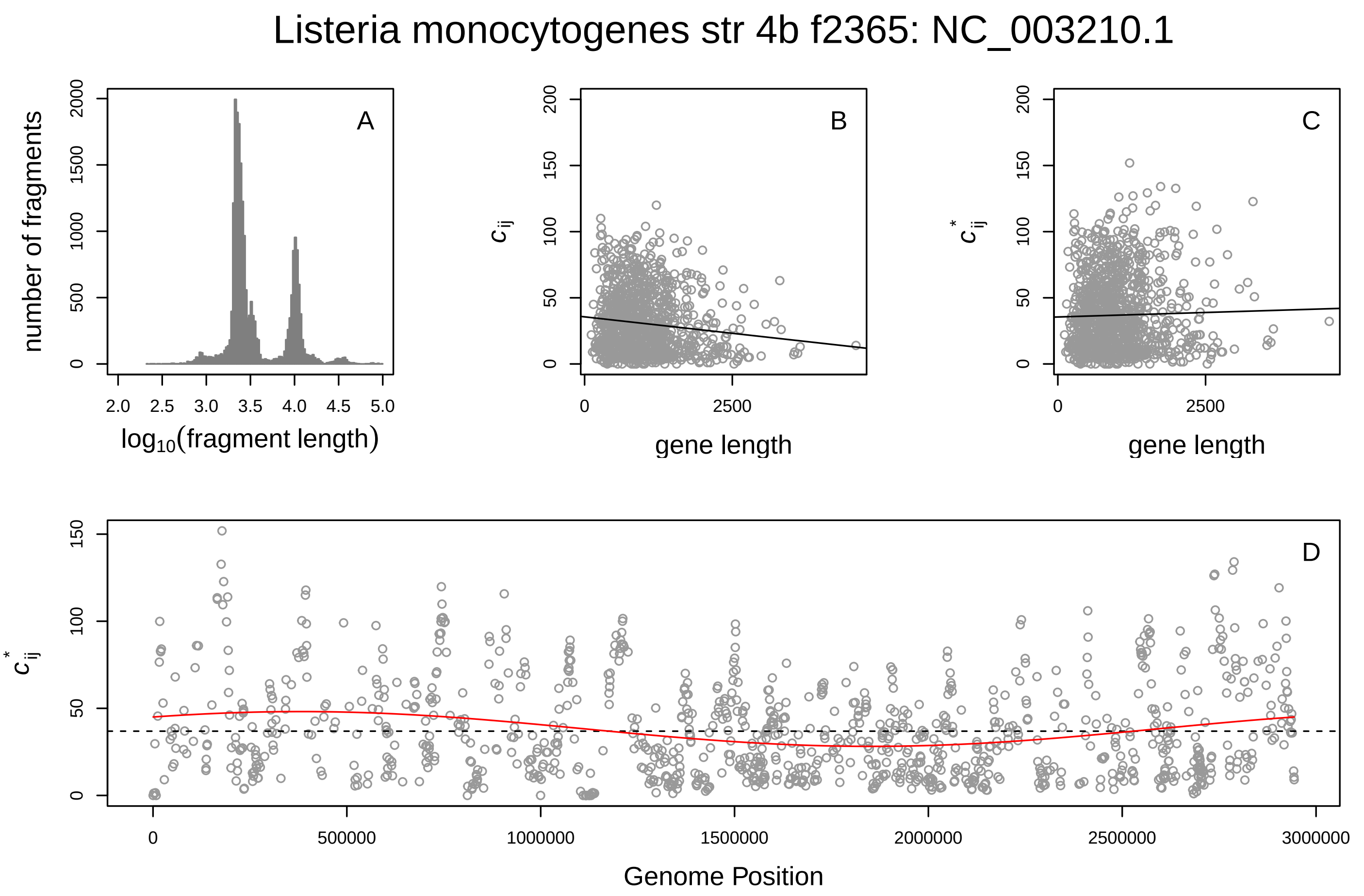
Listeria monocytogenes str 4b f2365: NC_003210.1
A
number of fragments
log10(fragment length)
B
$c_{ij}$
gene length
C
$c^*_{ij}$
gene length
D
$c^*_{ij}$
Genome Position

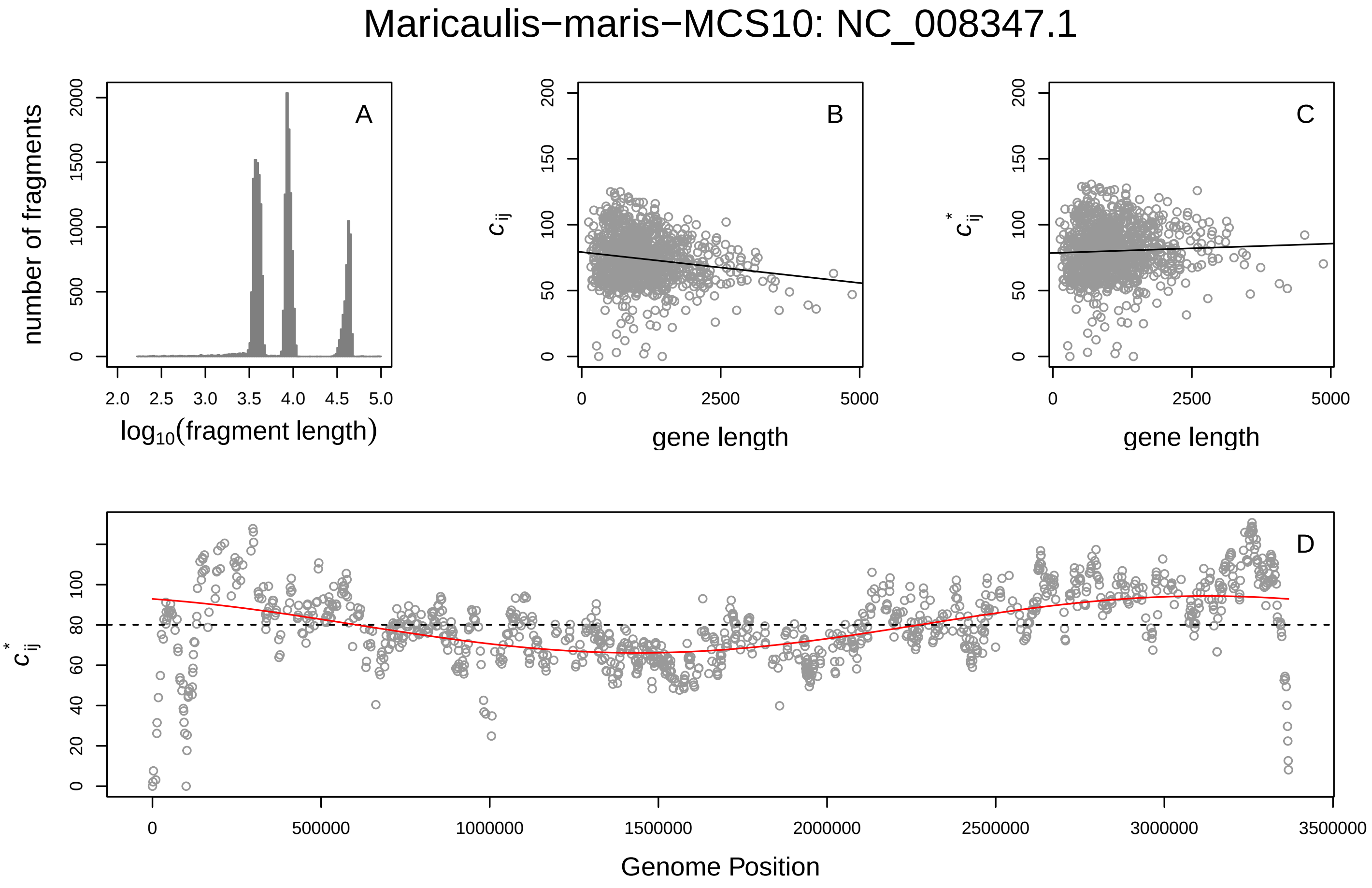
Maricaulis−maris−MCS10: NC_008347.1
A
number of fragments
log10(fragment length)
B
$c_{ij}$
gene length
C
$c^*_{ij}$
gene length
D
$c^*_{ij}$
Genome Position

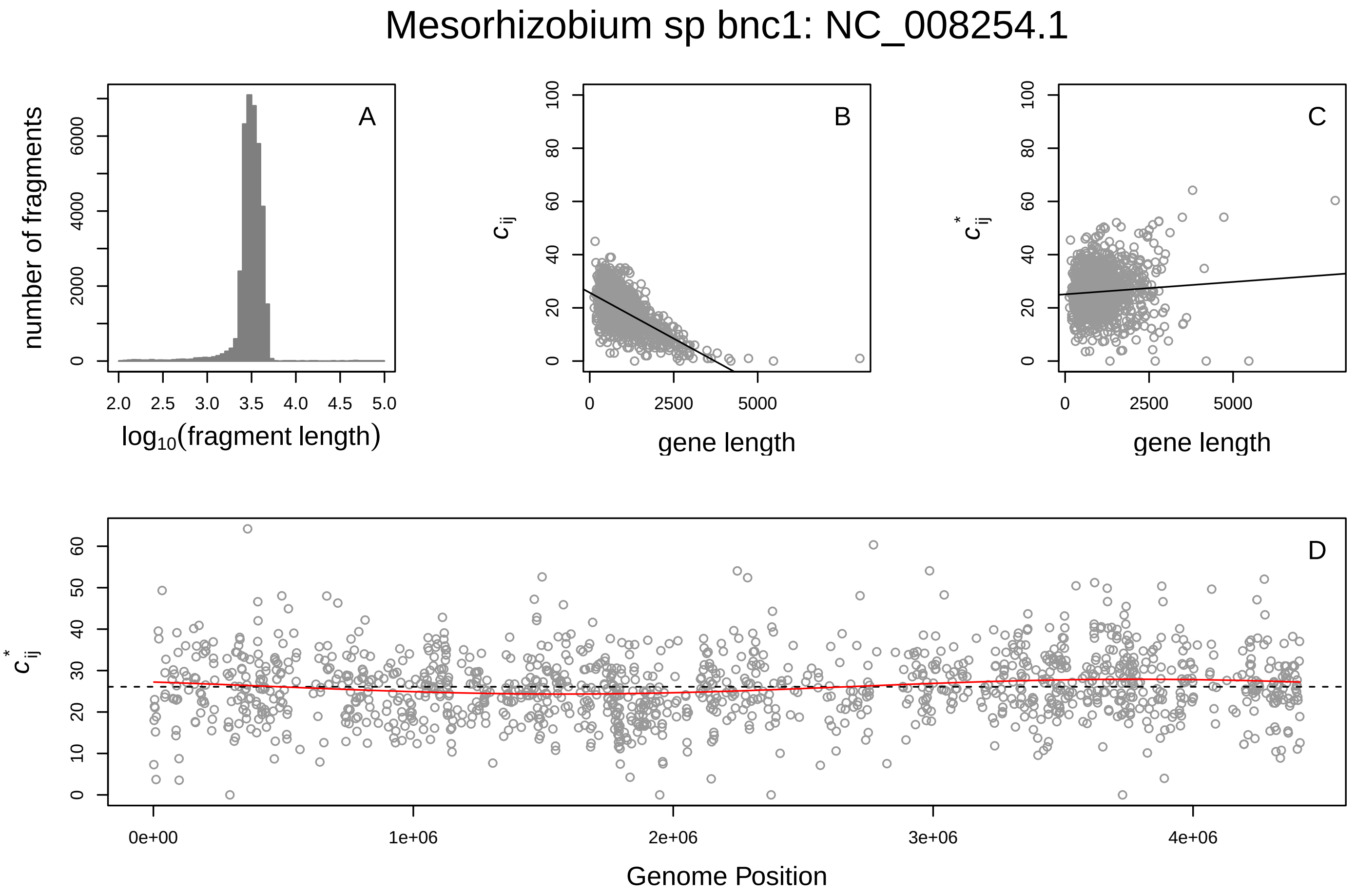
Mesorhizobium sp bnc1: NC_008254.1
A
B
C
D
number of fragments
log10(fragment length)
c_ij
c*_ij
gene length
Genome Position

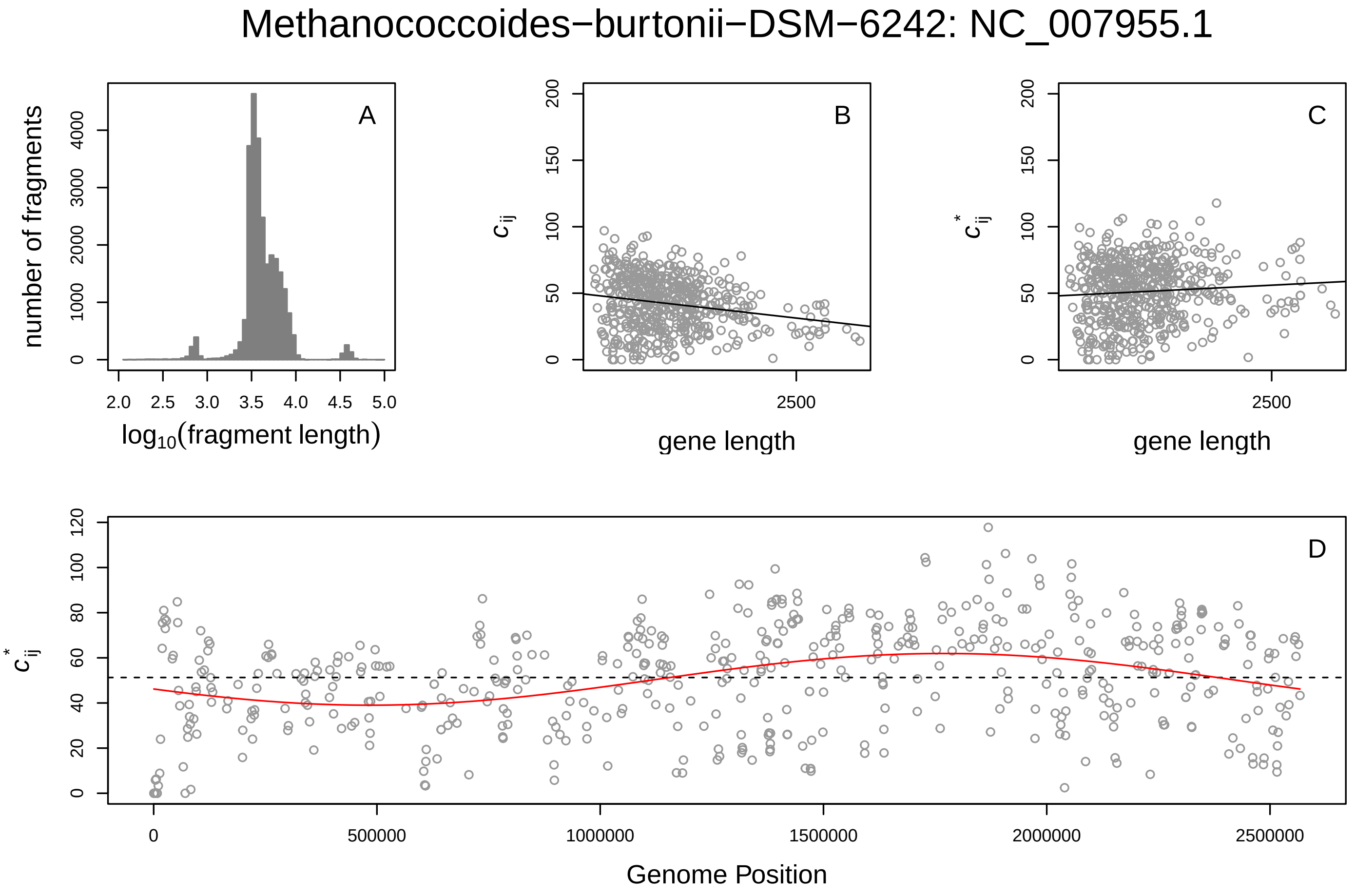
Methanococcoides−burtonii−DSM−6242: NC_007955.1
A
B
C
D
number of fragments
log10(fragment length)
c_ij
c*_ij
gene length
Genome Position

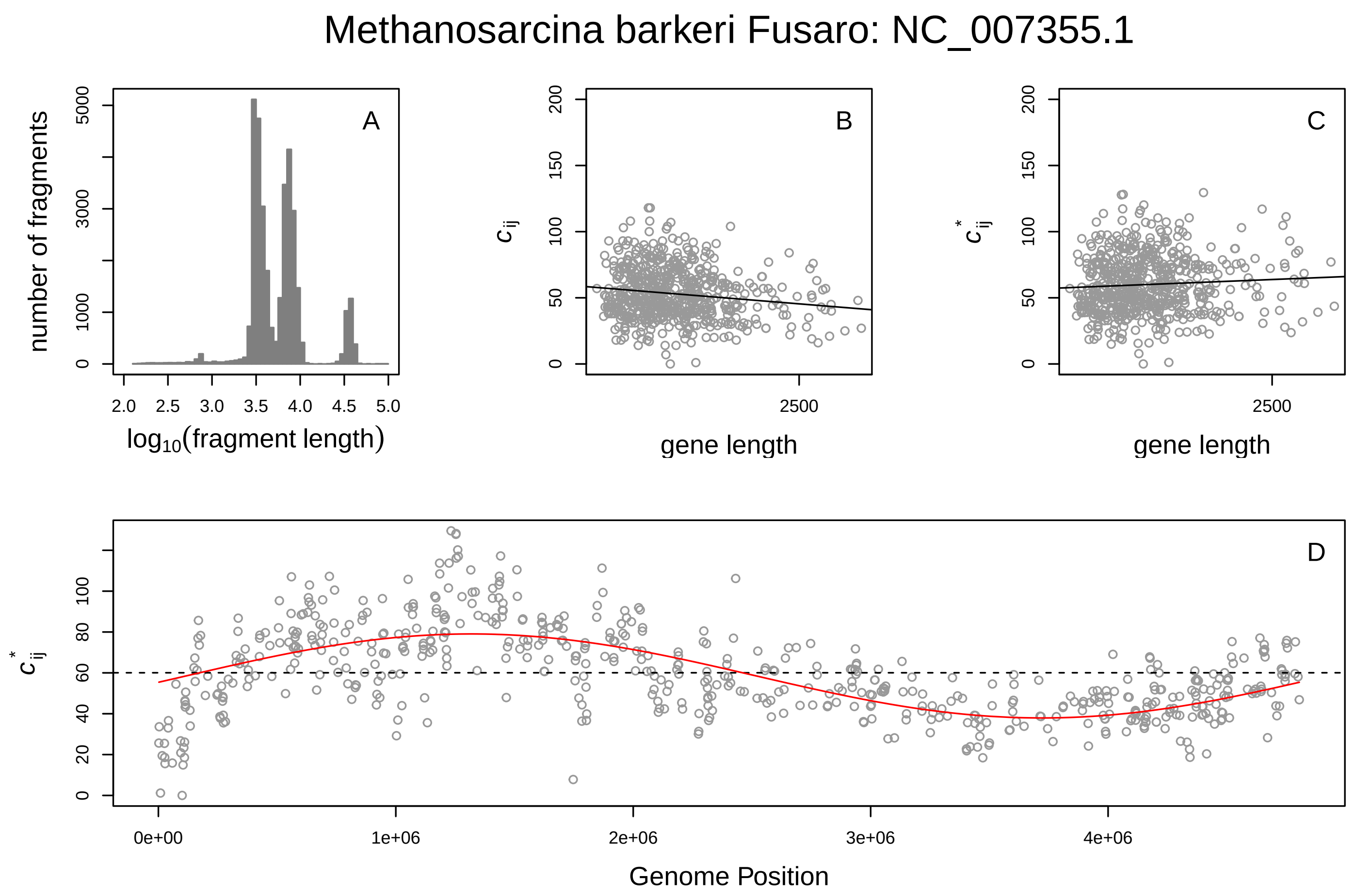
Methanosarcina barkeri Fusaro: NC_007355.1
A
number of fragments
log10(fragment length)
B
cij
gene length
C
c*ij
gene length
D
c*ij
Genome Position
0e+00
1e+06
2e+06
3e+06
4e+06

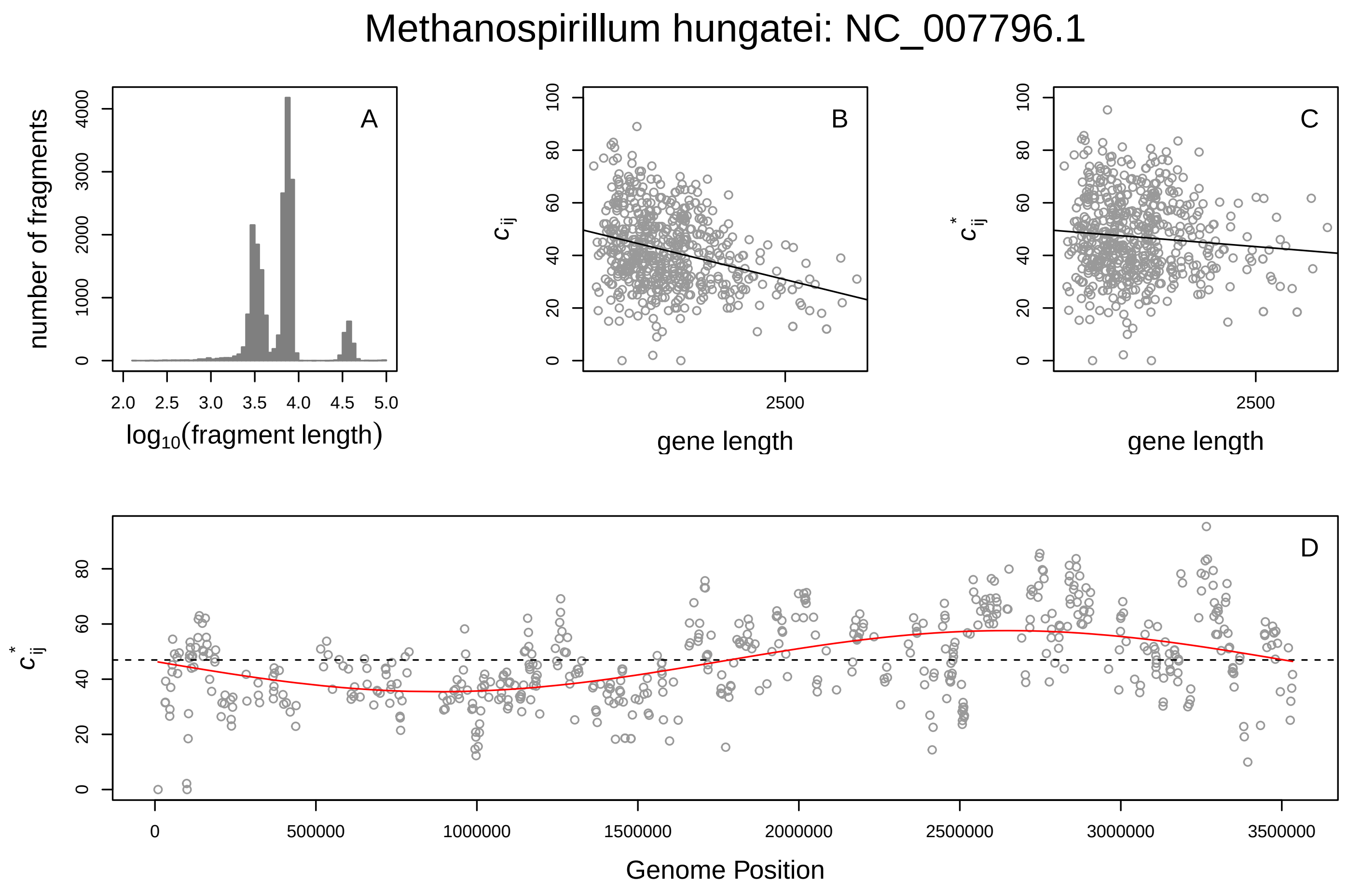
Methanospirillum hungatei: NC_007796.1
A
number of fragments
log10(fragment length)
B
cij
gene length
C
c*ij
gene length
D
c*ij
Genome Position
0
500000
1000000
1500000
2000000
2500000
3000000
3500000

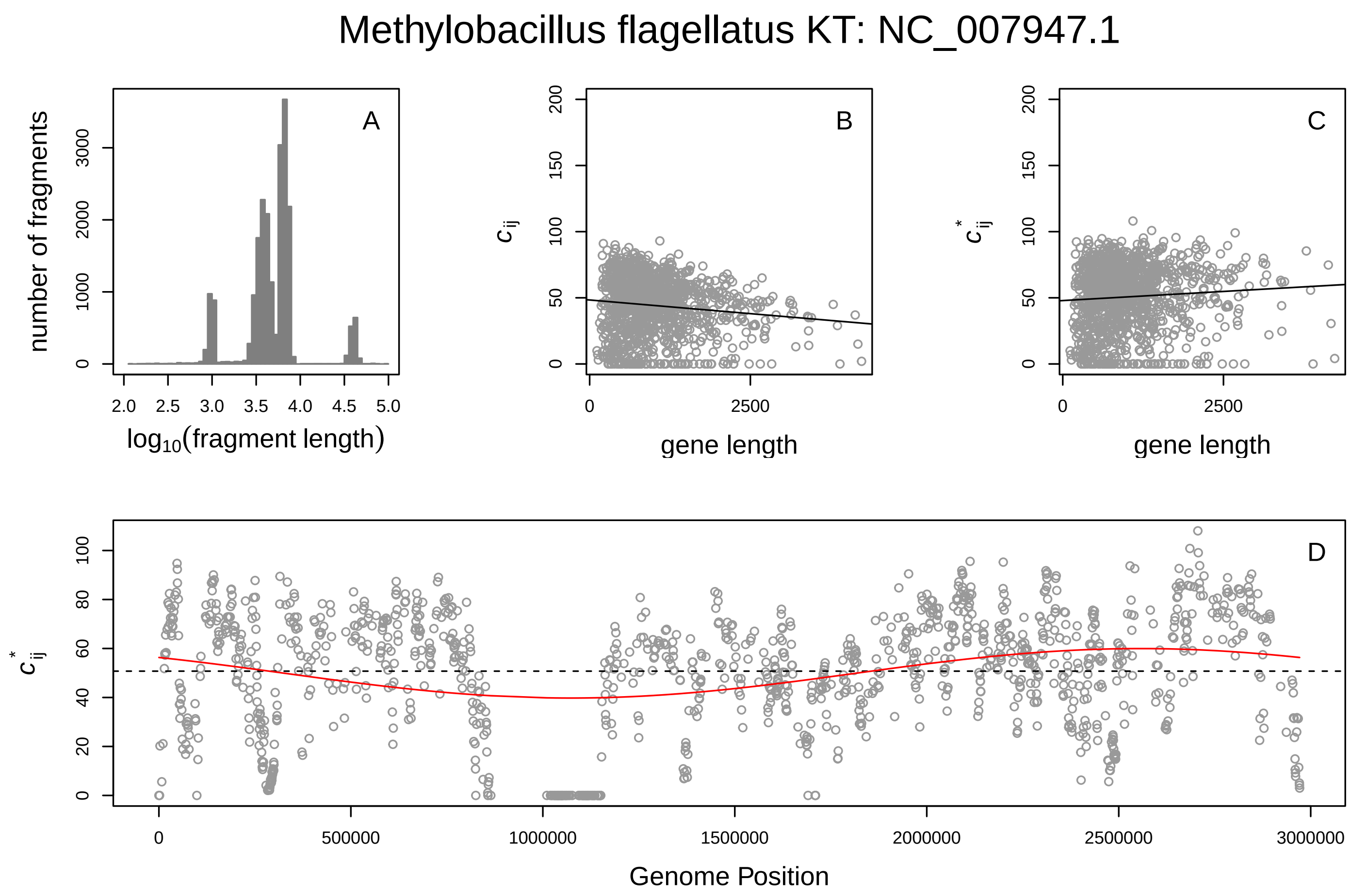
Methylobacillus flagellatus KT: NC_007947.1
A
B
C
D
number of fragments
log10(fragment length)
c_ij
c*_ij
gene length
Genome Position

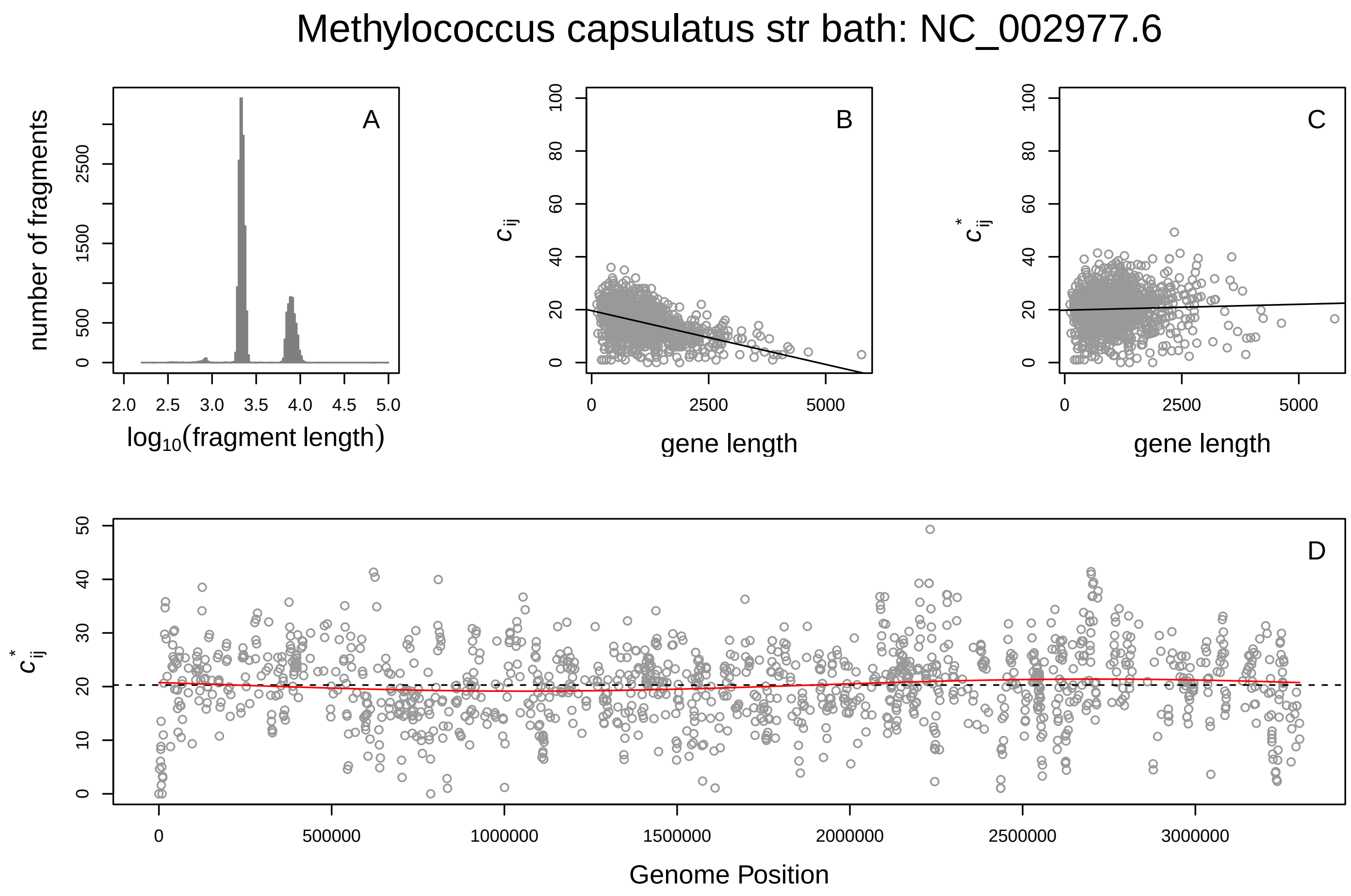
Methylococcus capsulatus str bath: NC_002977.6
A
B
C
D
number of fragments
log10(fragment length)
c_ij
c*_ij
gene length
Genome Position

#### Moorella thermoacetica ATCC 39073: NC_007644.1

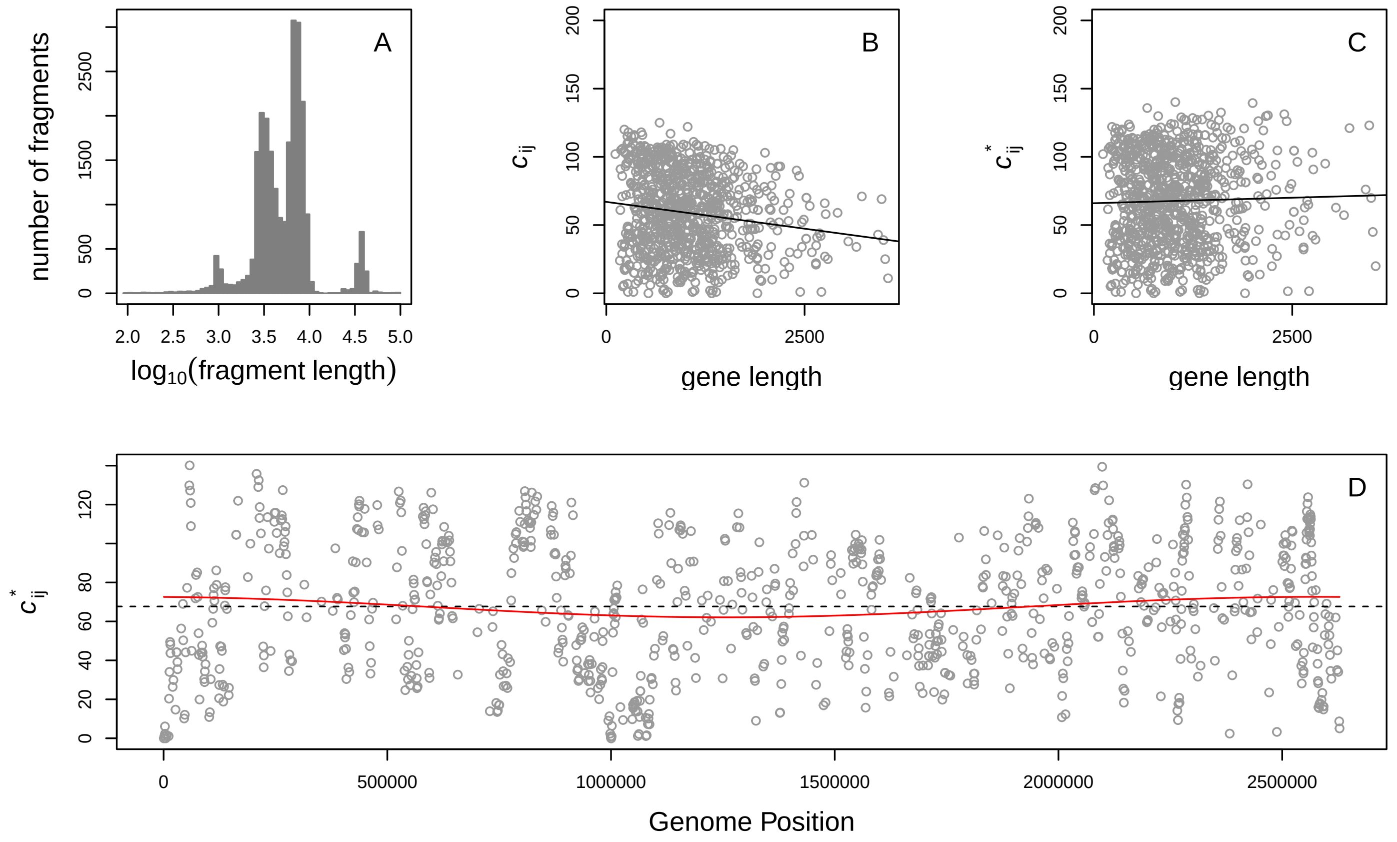


#### Mycobacterium sp MCS: NC_008146.1

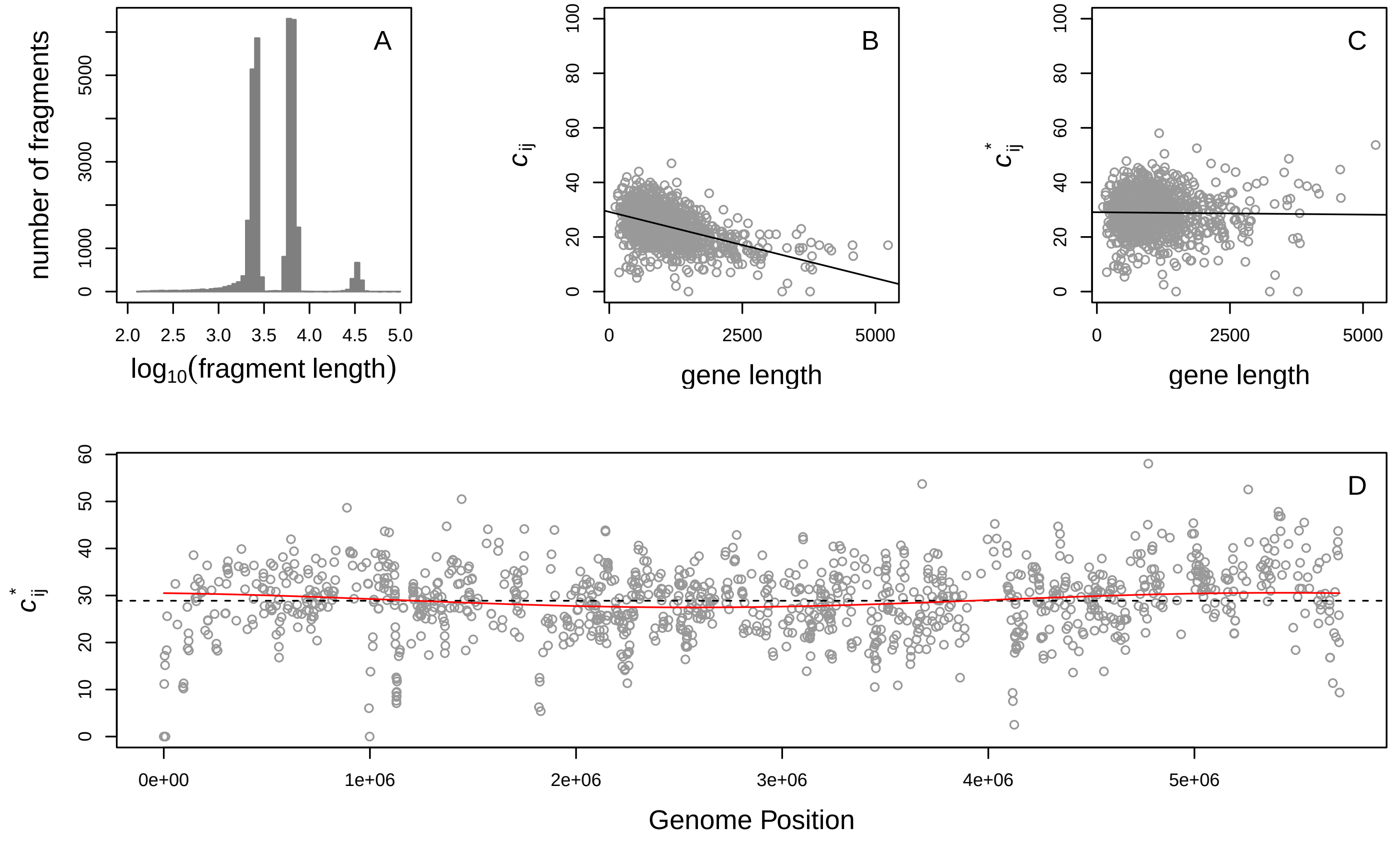

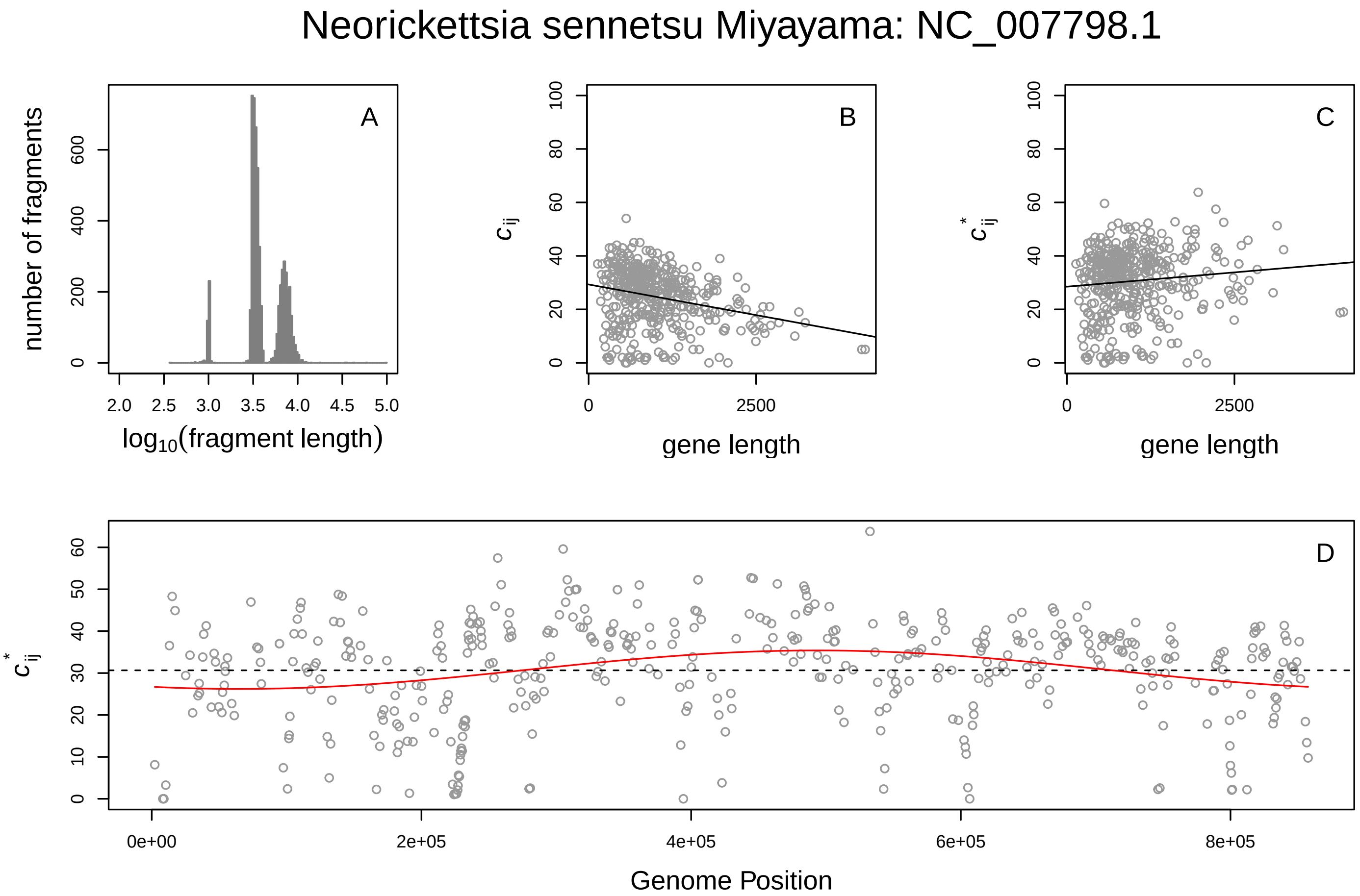
Neorickettsia sennetsu Miyayama: NC_007798.1
A
number of fragments
log10(fragment length)
B
c_ij
gene length
C
c*_ij
gene length
D
c*_ij
Genome Position

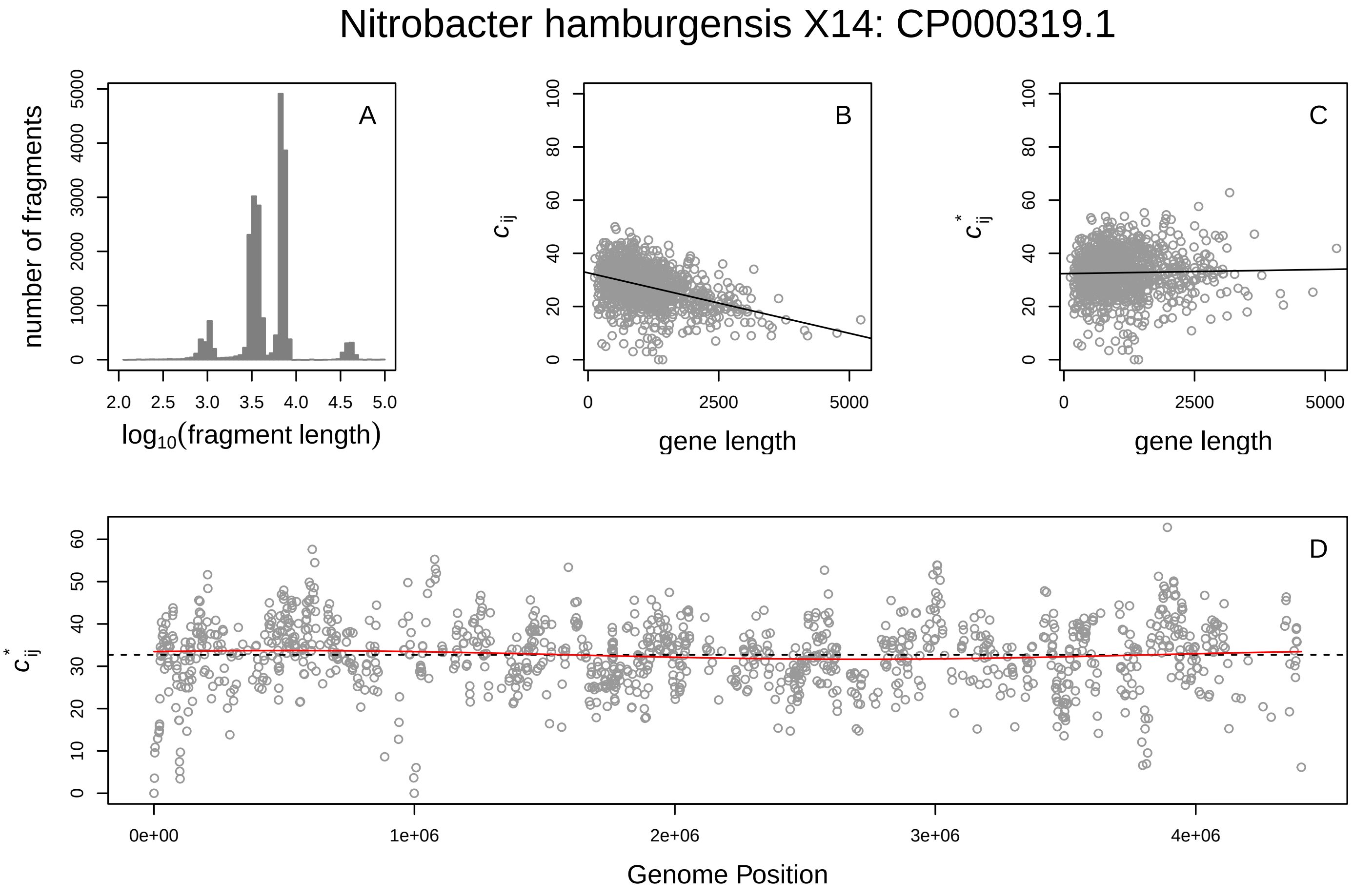
Nitrobacter hamburgensis X14: CP000319.1
A
number of fragments
log10(fragment length)
B
c_ij
gene length
C
c*_ij
gene length
D
c*_ij
Genome Position

Nitrobacter winogradskyi Nb 255: CP000115.1

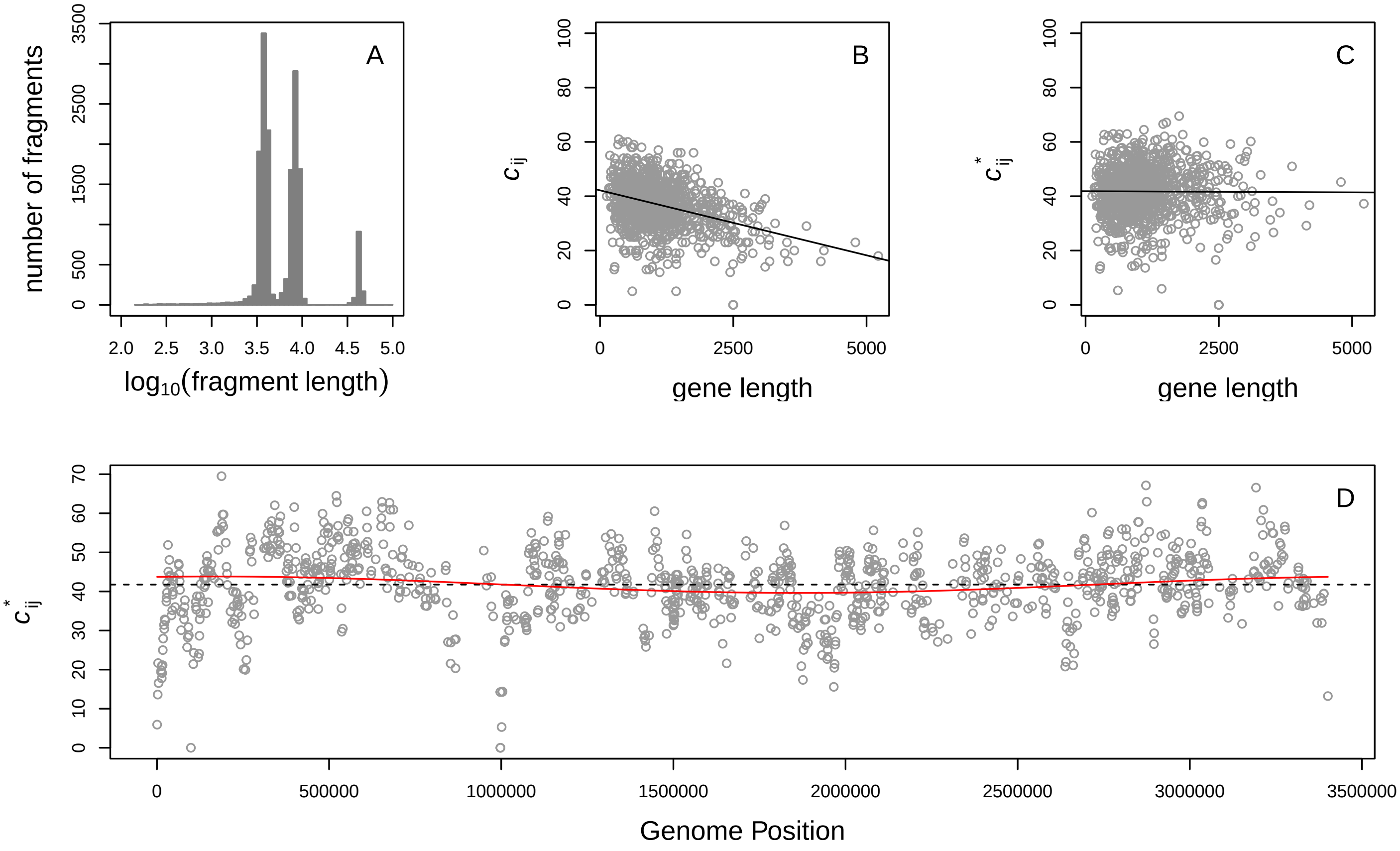


Nitrosococcus oceani ATCC 19707: CP000127.1

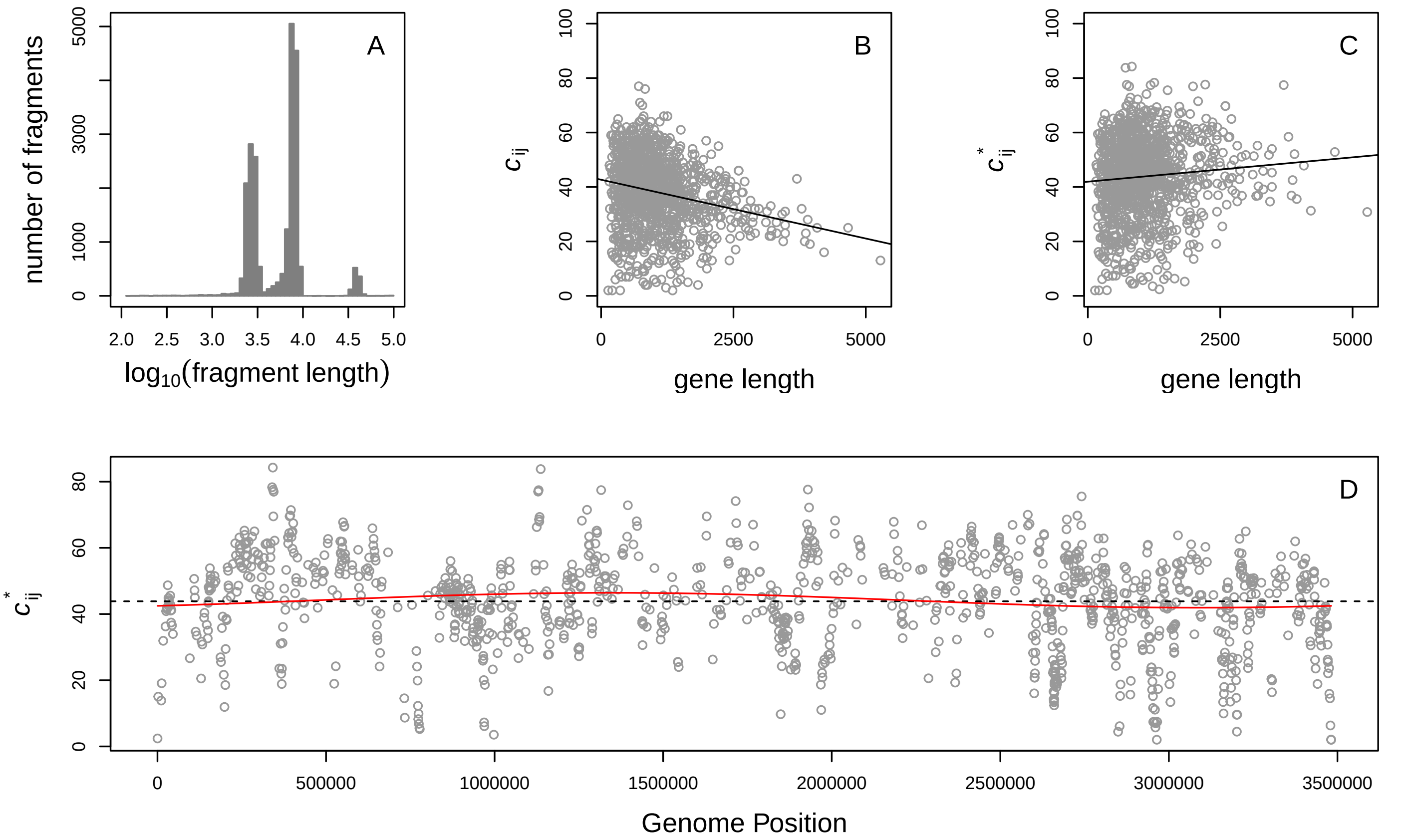

#### Novosphingobium aromaticivorans DSM 12444: CP000248.1

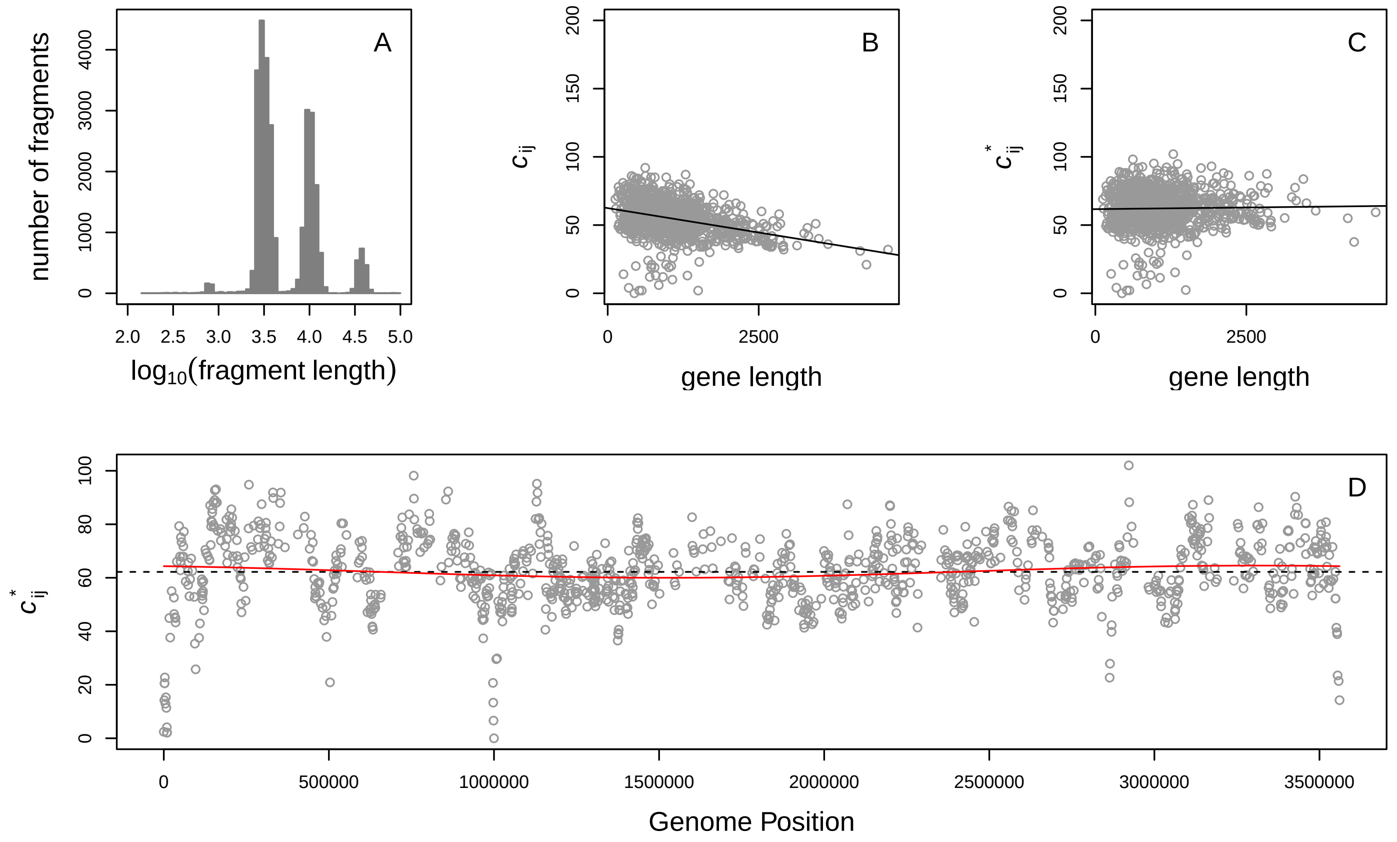


#### Pelobacter propinonicus DSM 2379: CP000482.1

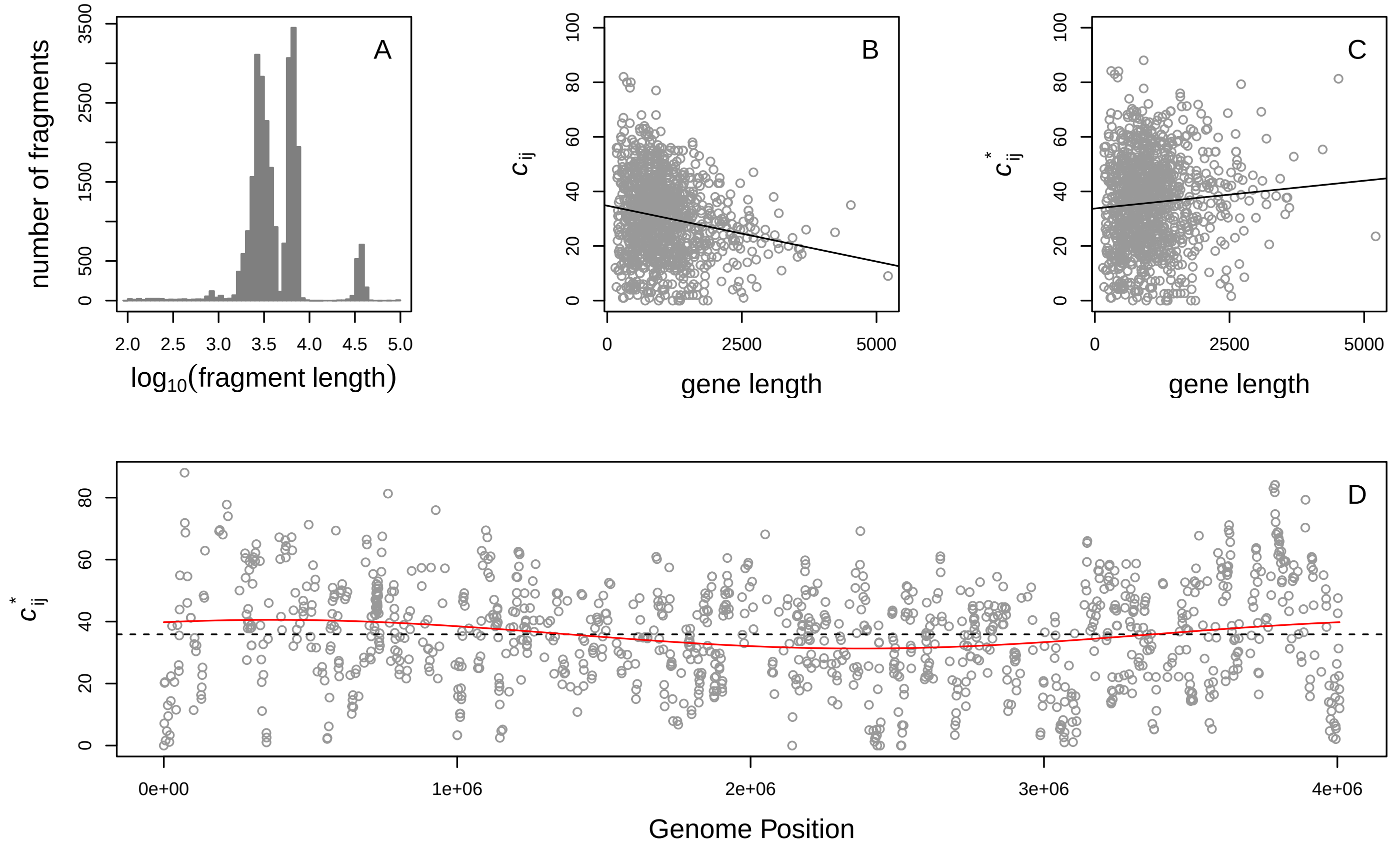

Polaromonas sp JS666: CP000316.1

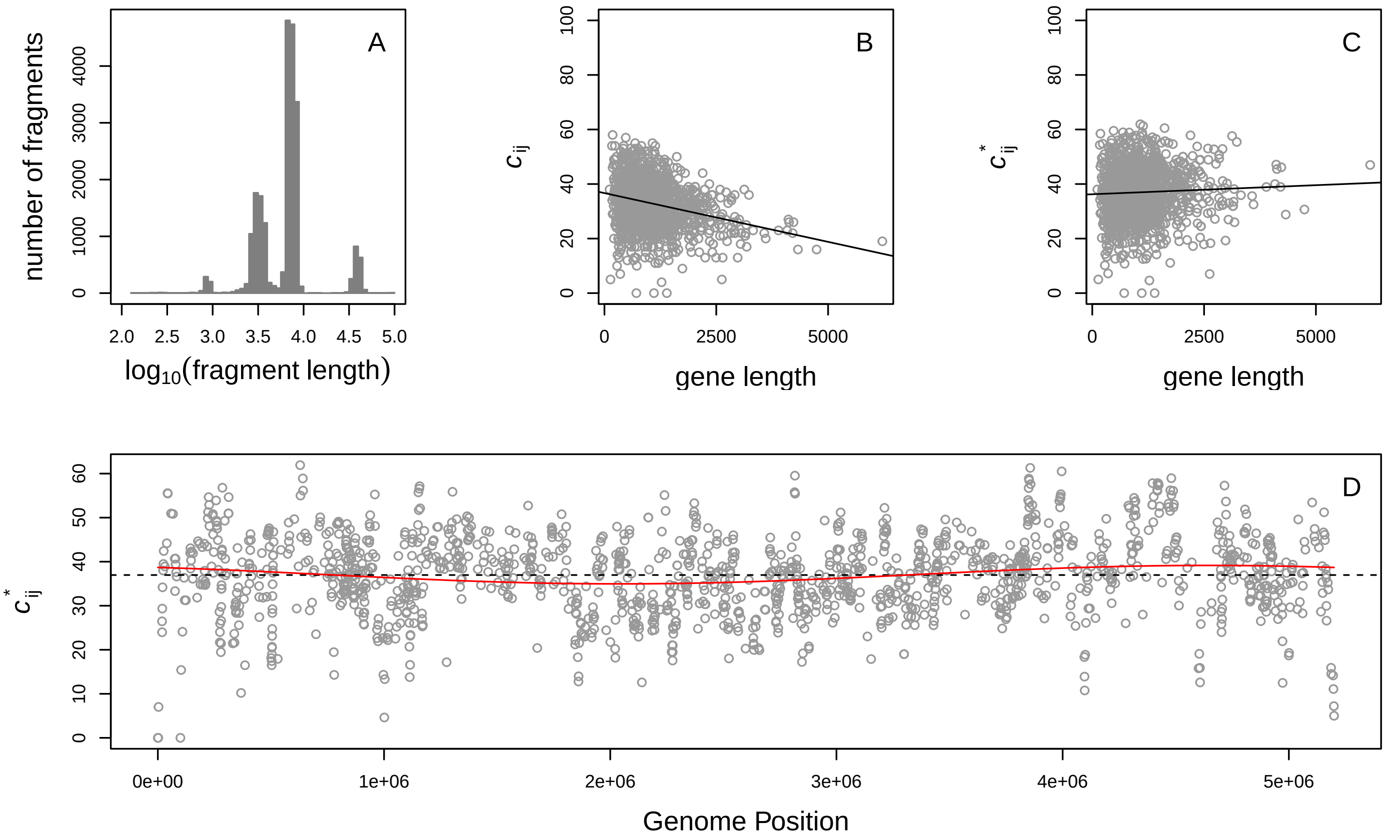


Prochlorococcus marinus NATL2A: NC_007335.2

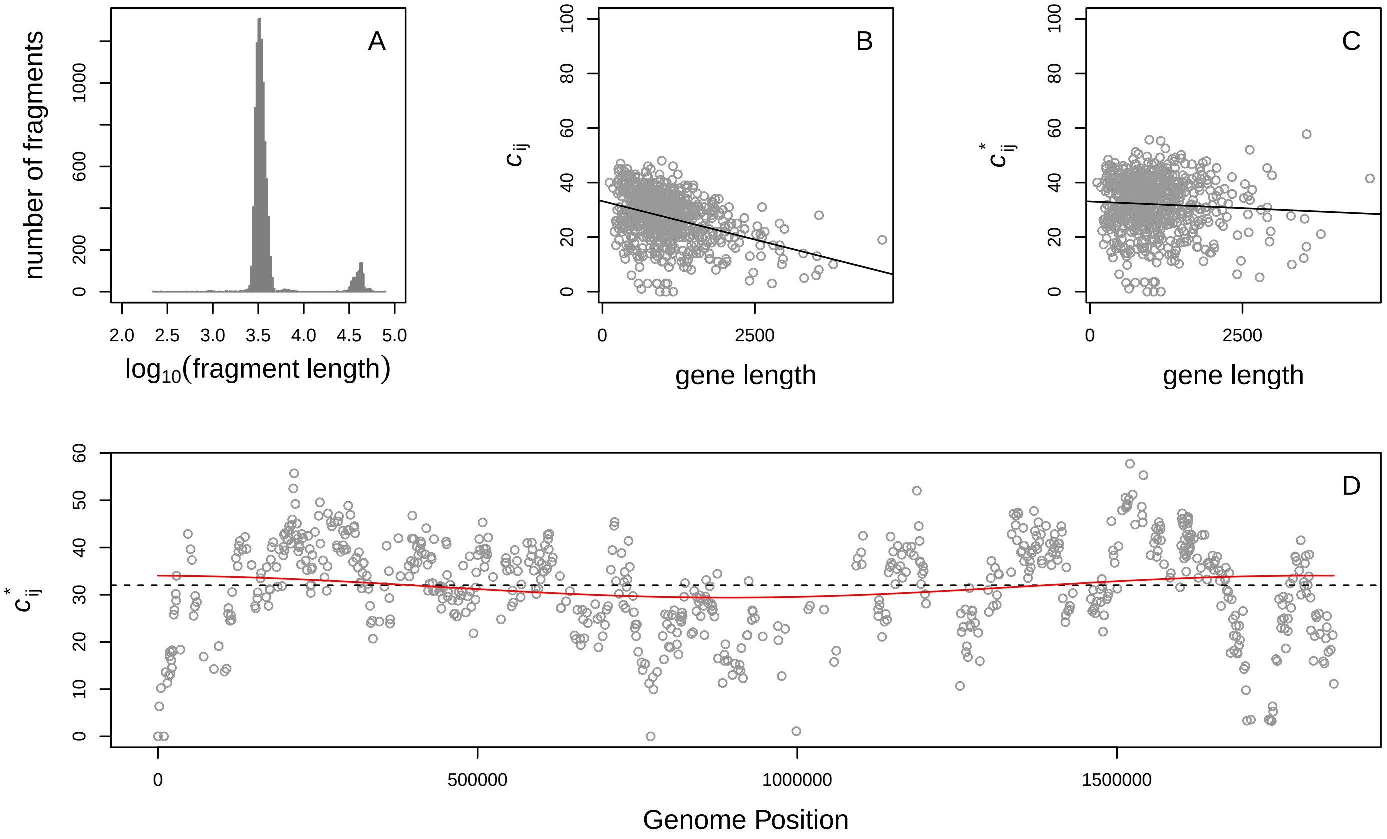

Pseudoalteromonas atlantica T6c: NC_008228.1

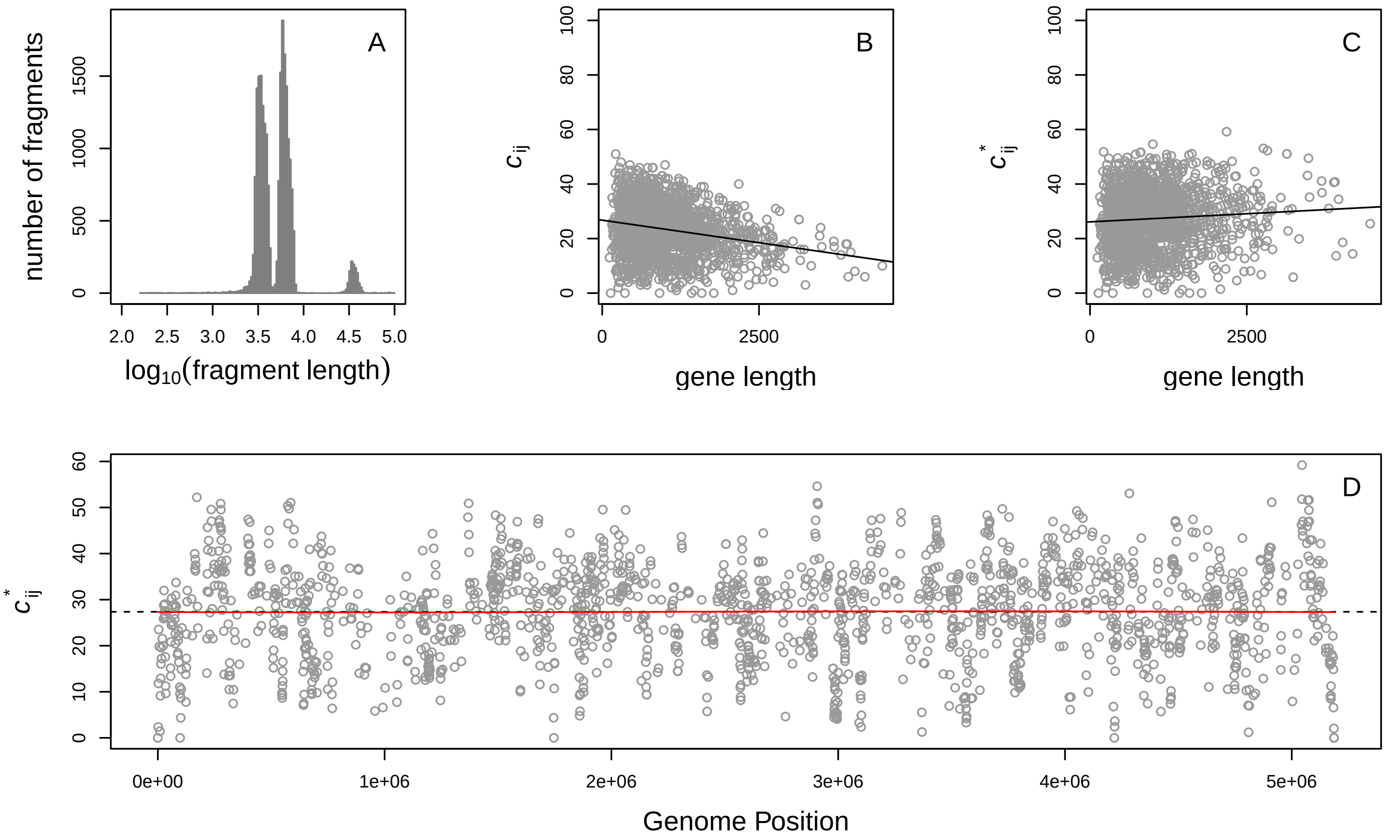


Pseudomonas syringae pv syringae b728a: NZ_CP068034.2

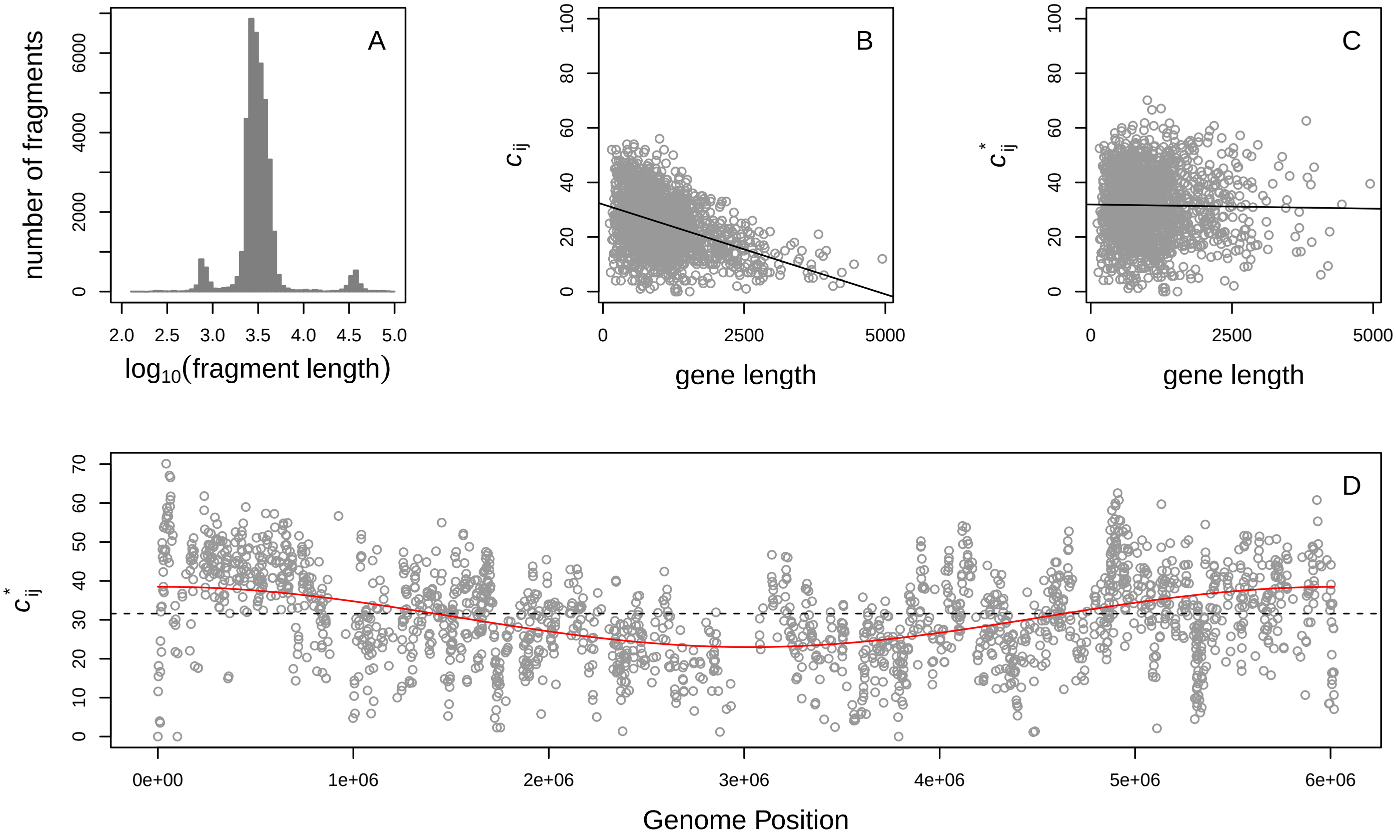

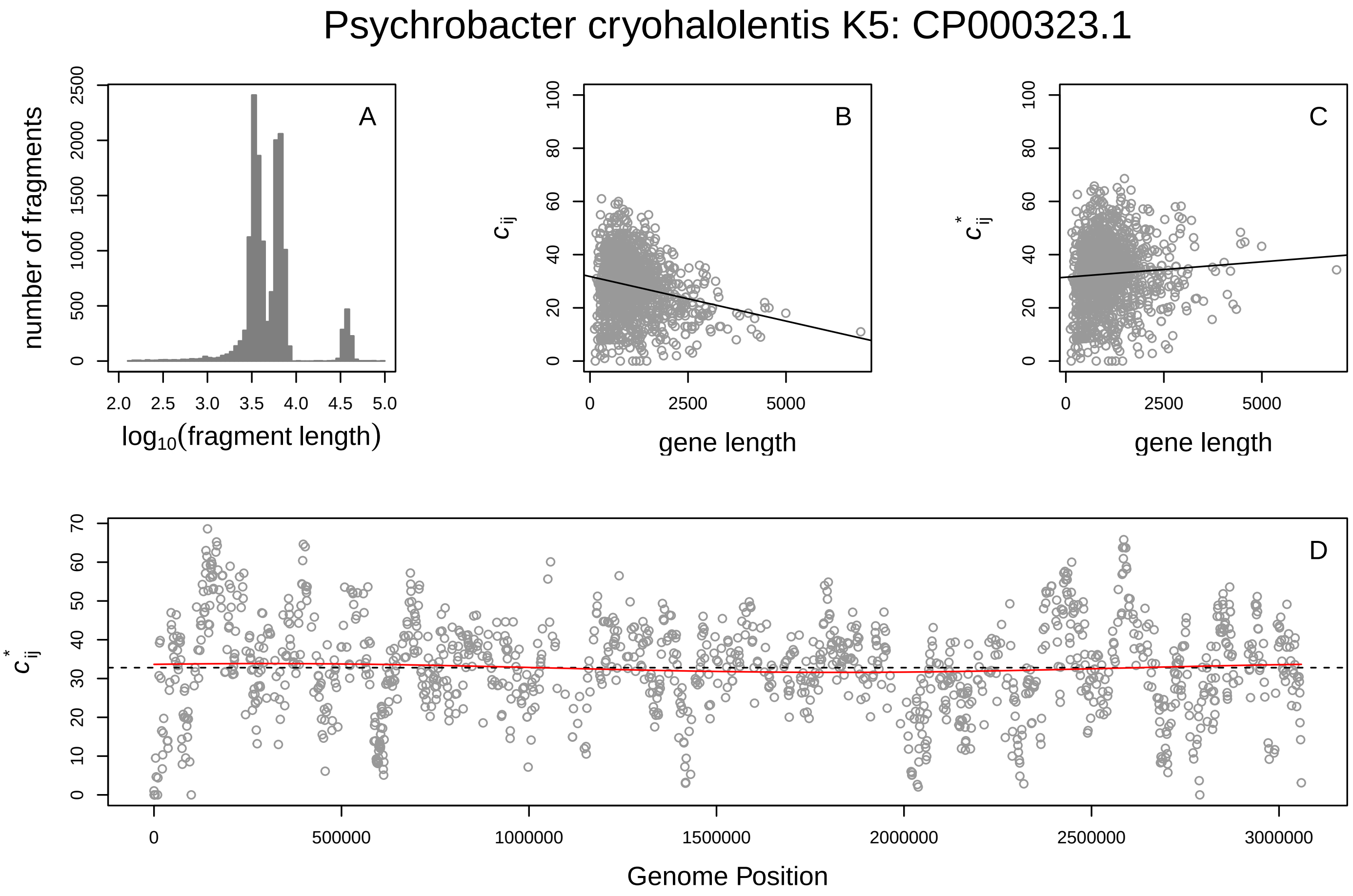
Psychrobacter cryohalolentis K5: CP000323.1
A
number of fragments
log10(fragment length)
B
$c_{ij}$
gene length
C
$c^*_{ij}$
gene length
D
$c^*_{ij}$
Genome Position

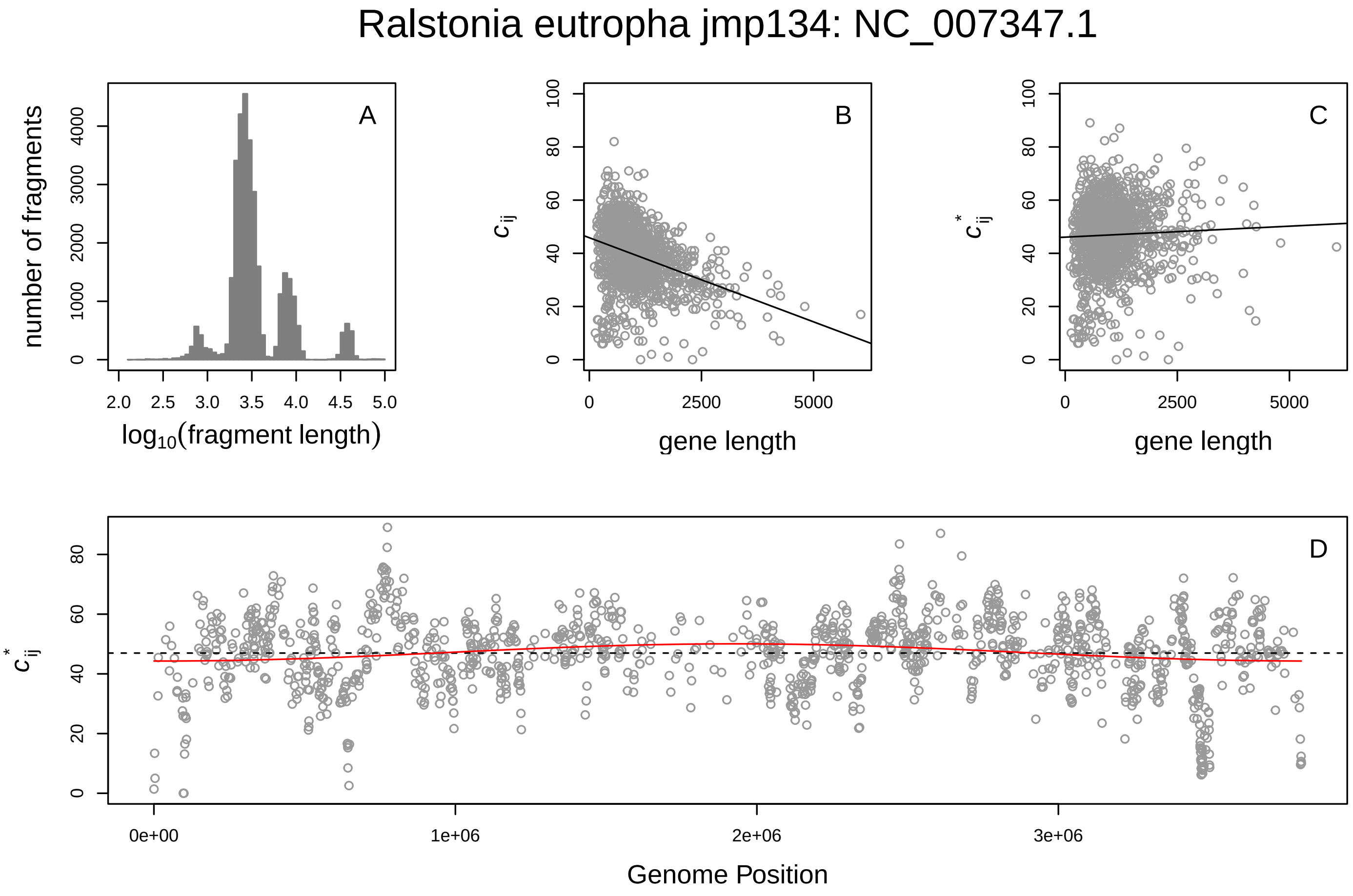
Ralstonia eutropha jmp134: NC_007347.1
A
number of fragments
log10(fragment length)
B
$c_{ij}$
gene length
C
$c^*_{ij}$
gene length
D
$c^*_{ij}$
Genome Position

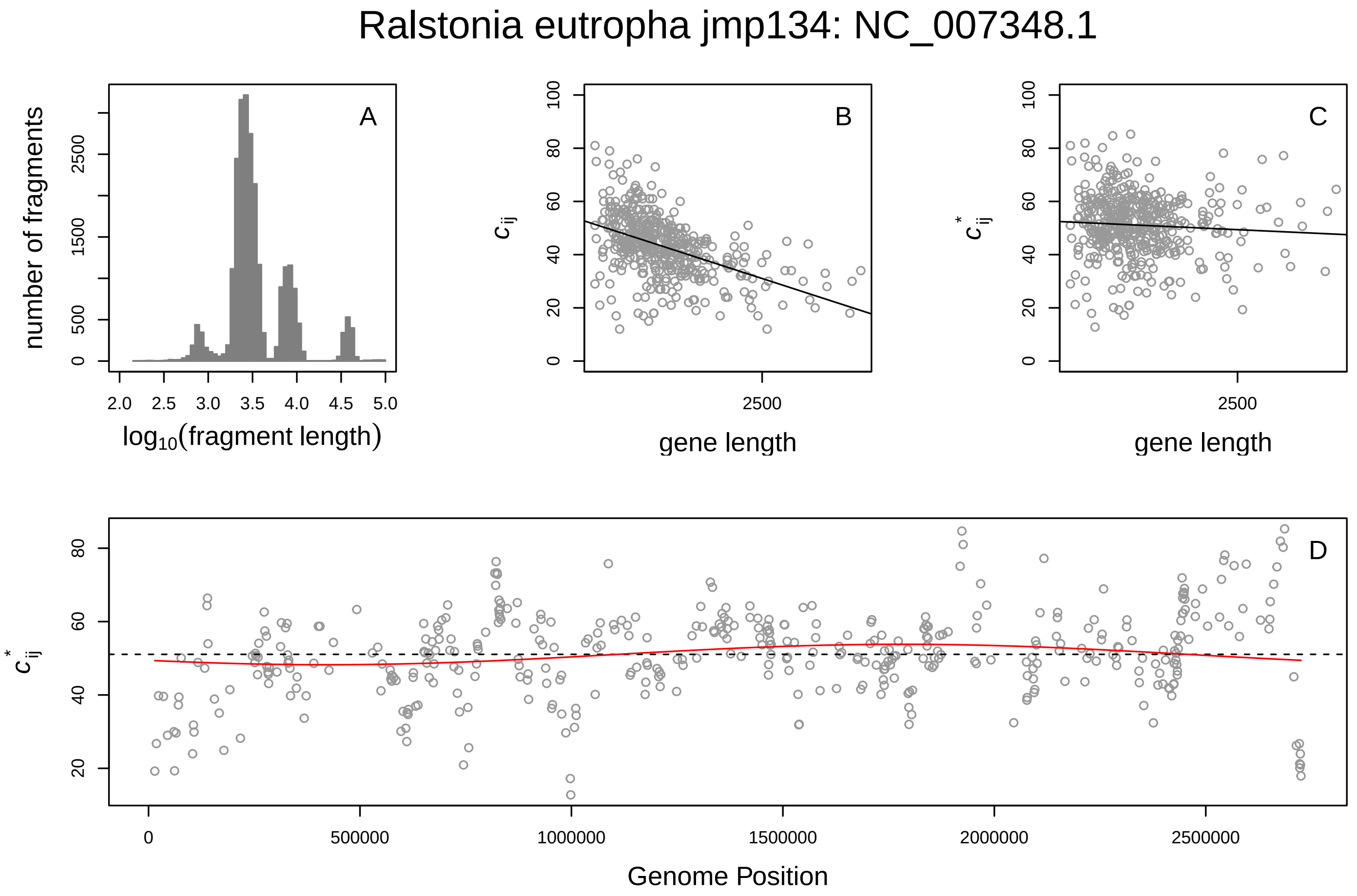
Ralstonia eutropha jmp134: NC_007348.1
A
number of fragments
log10(fragment length)
B
$c_{ij}$
gene length
C
$c^*_{ij}$
gene length
D
$c^*_{ij}$
Genome Position

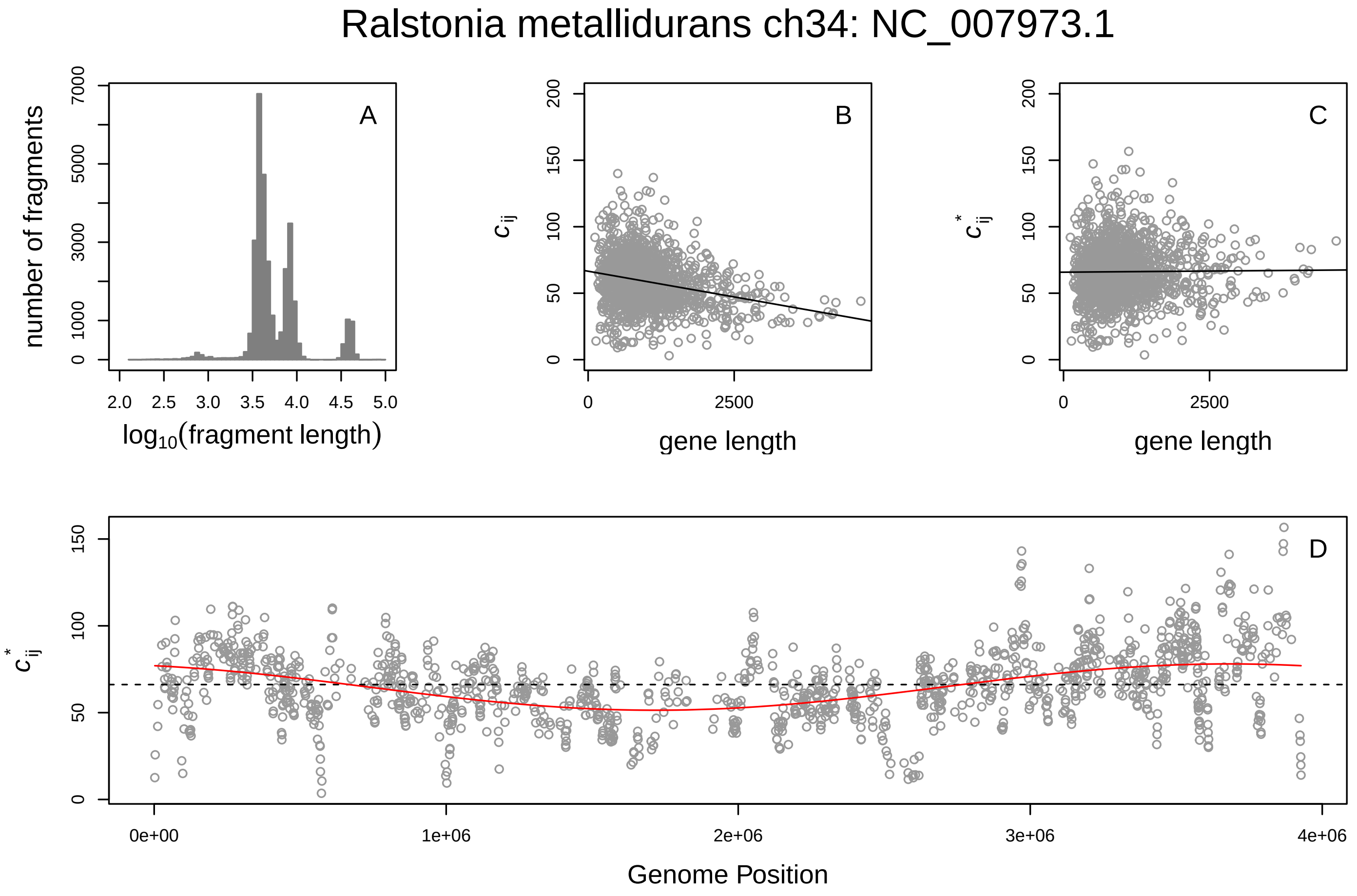
Ralstonia metallidurans ch34: NC_007973.1
A
number of fragments
log10(fragment length)
B
$c_{ij}$
gene length
C
$c^*_{ij}$
gene length
D
$c^*_{ij}$
Genome Position

#### Rhodoferax ferrireducens DSM 15236: CP000267.1

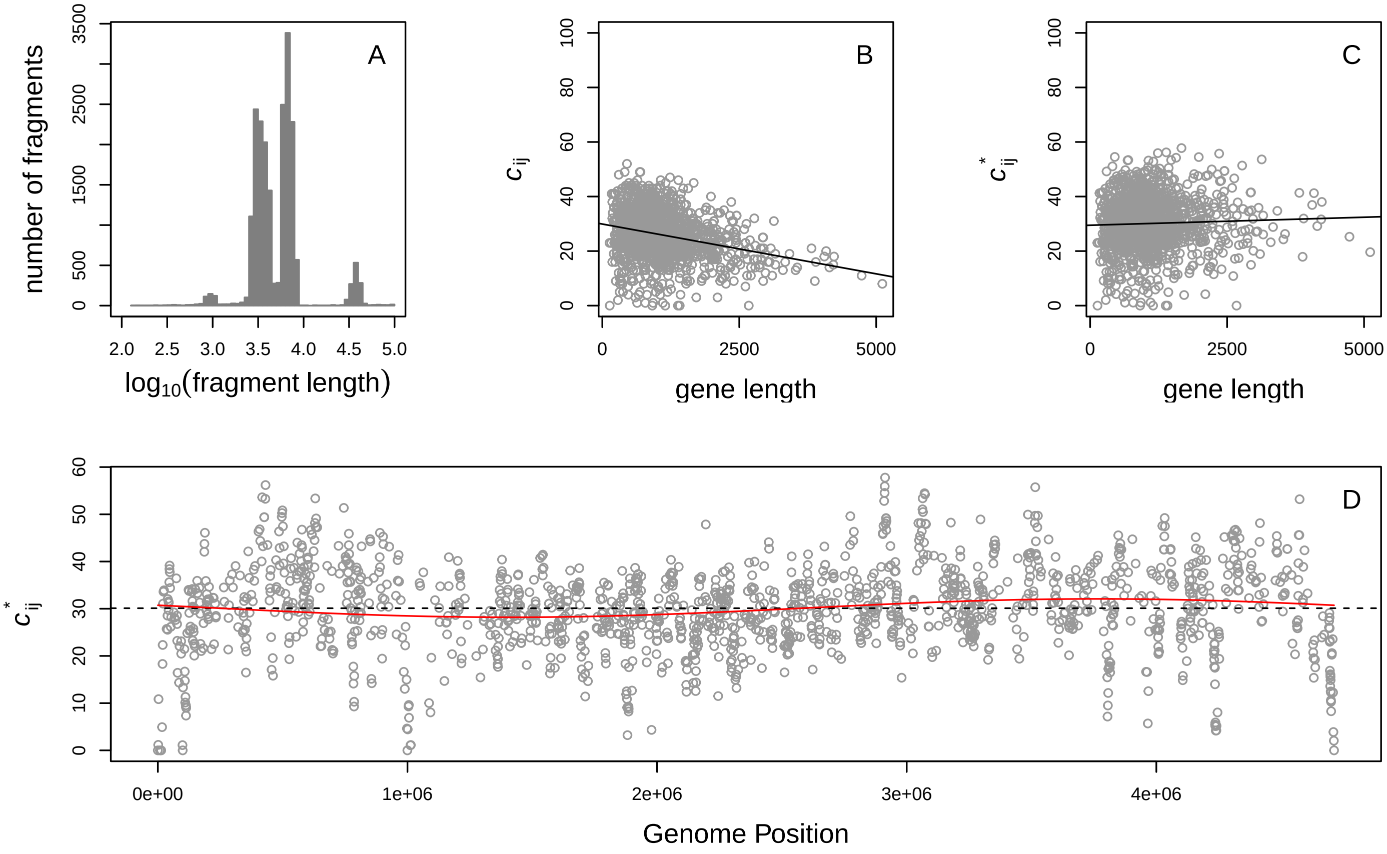


#### Rhodopseudomonas Palustris Bisb18: NC_007925.1

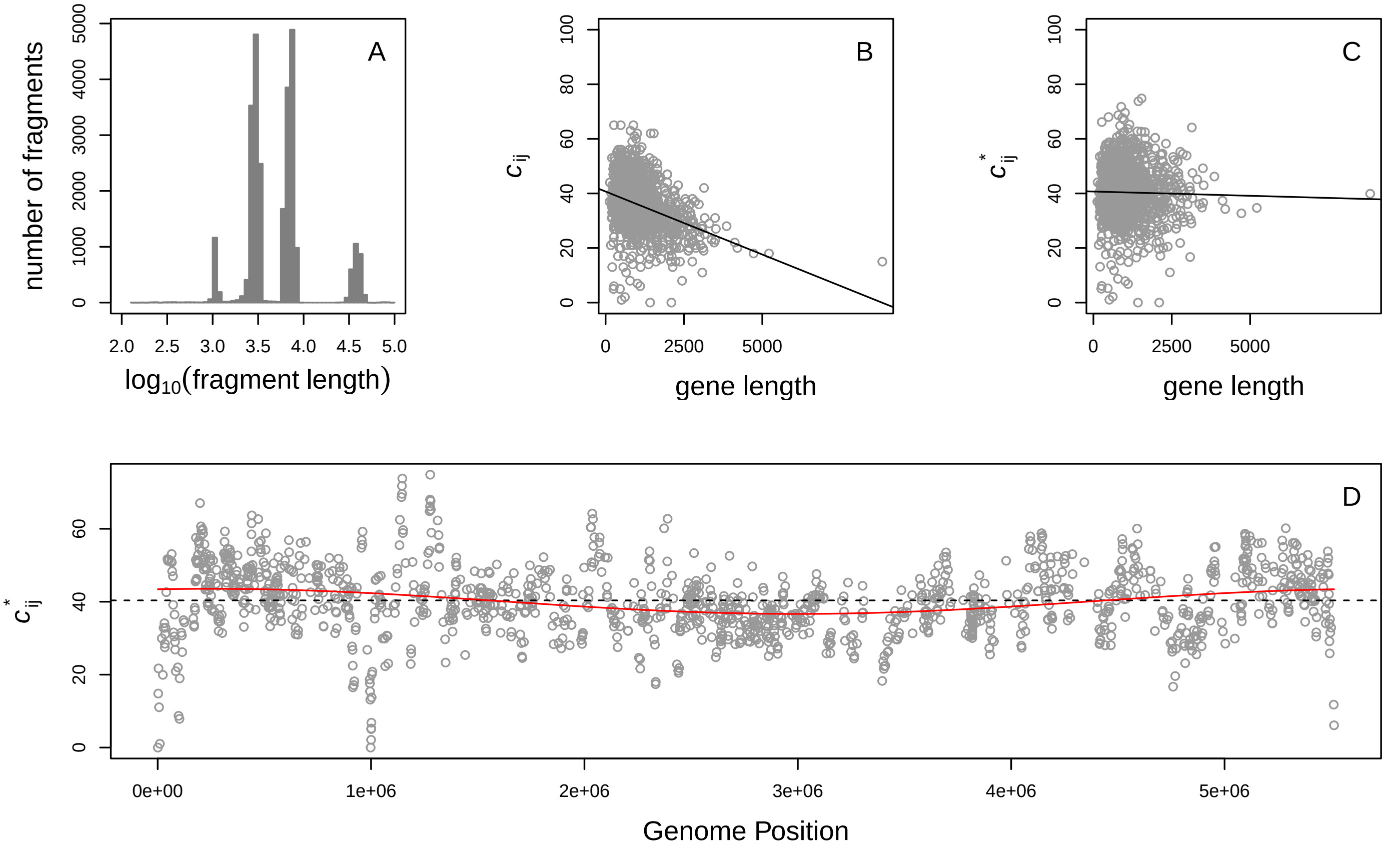

#### Rhodopseudomonas Palustris Bisb5: CP000283.1

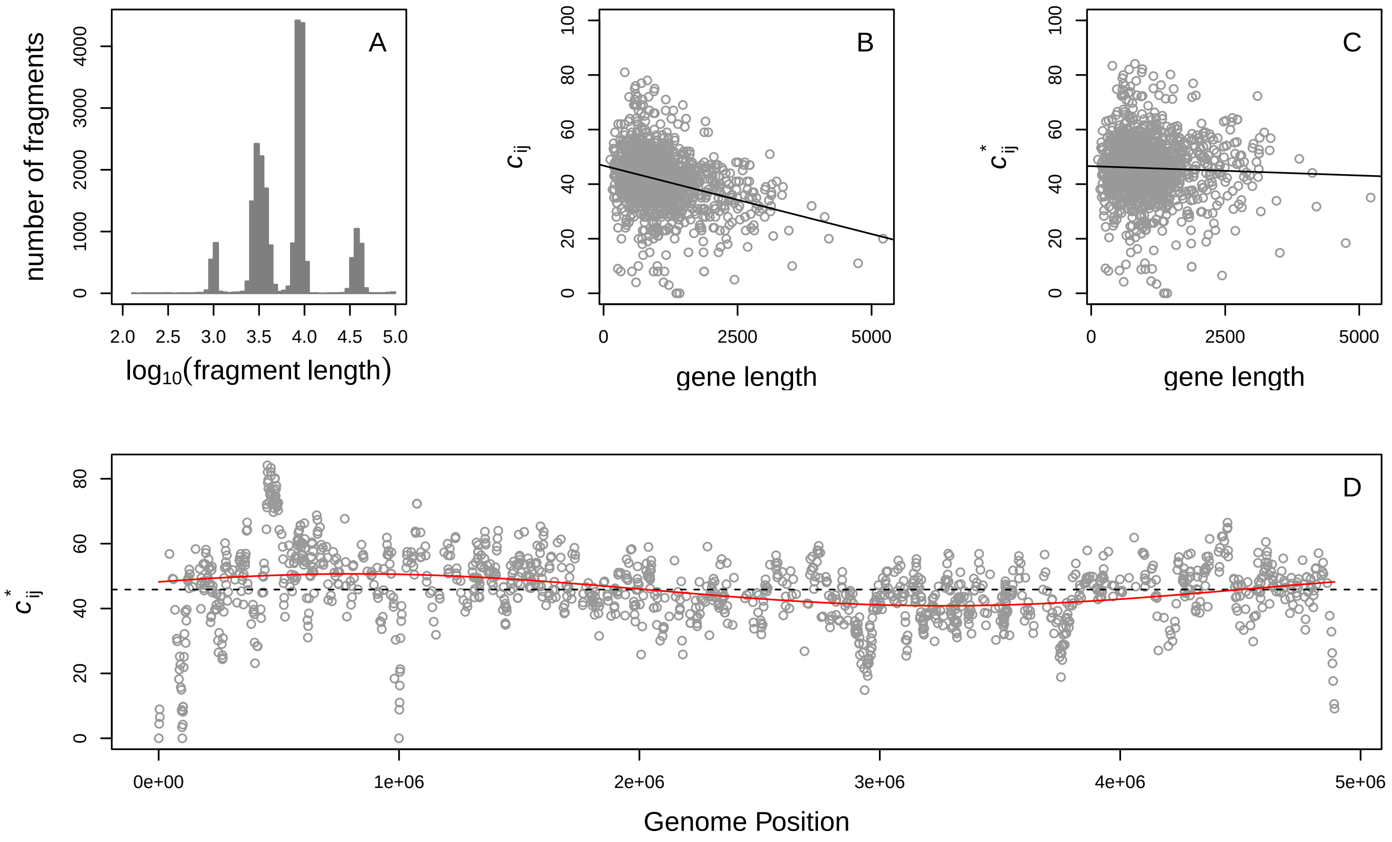


#### Rhodopseudomonas palustris tie 1: NC_008435.1

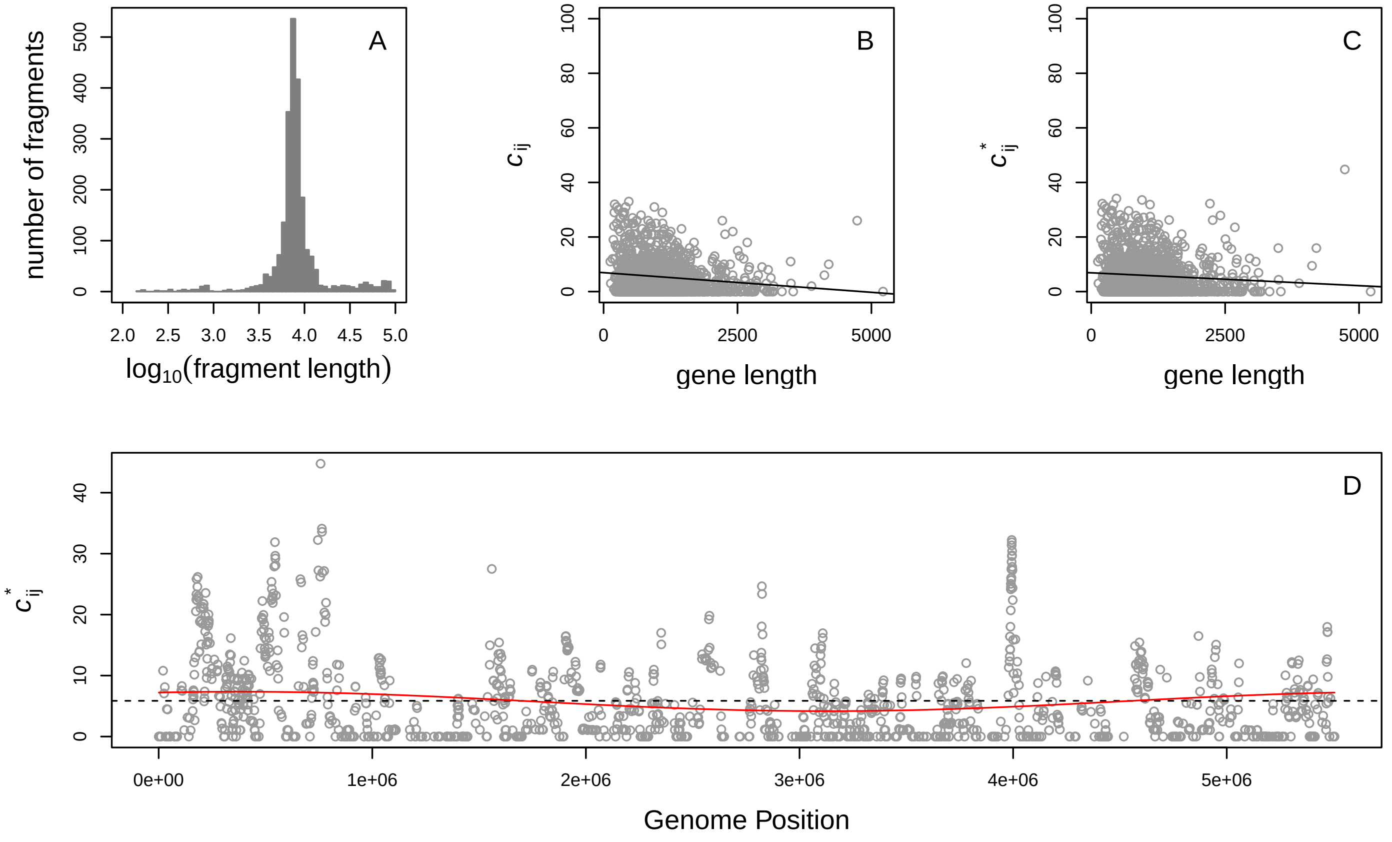

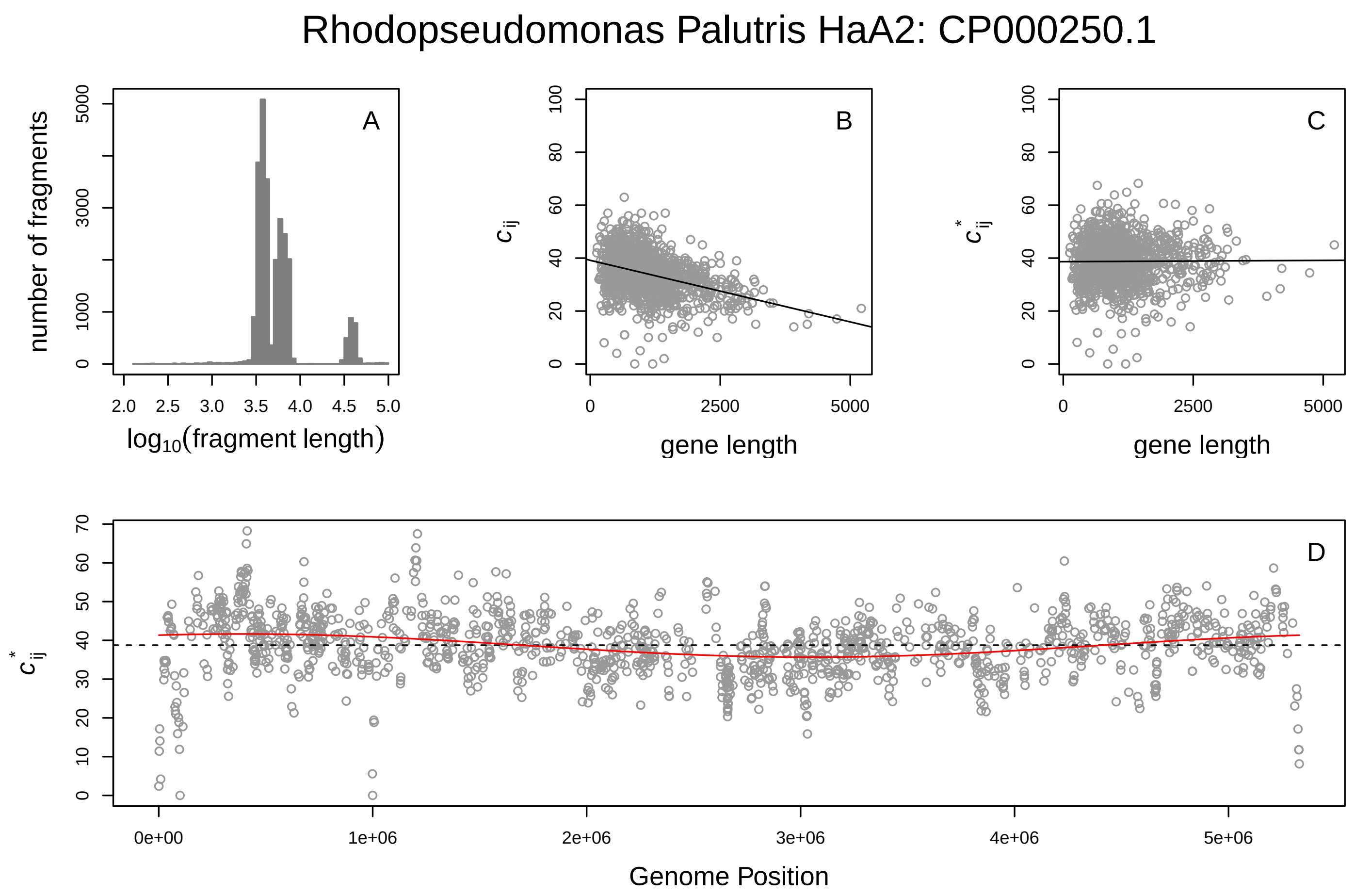
Rhodopseudomonas Palutris HaA2: CP000250.1
A
number of fragments
log10(fragment length)
B
c_ij
gene length
C
c*_ij
gene length
D
c*_ij
Genome Position

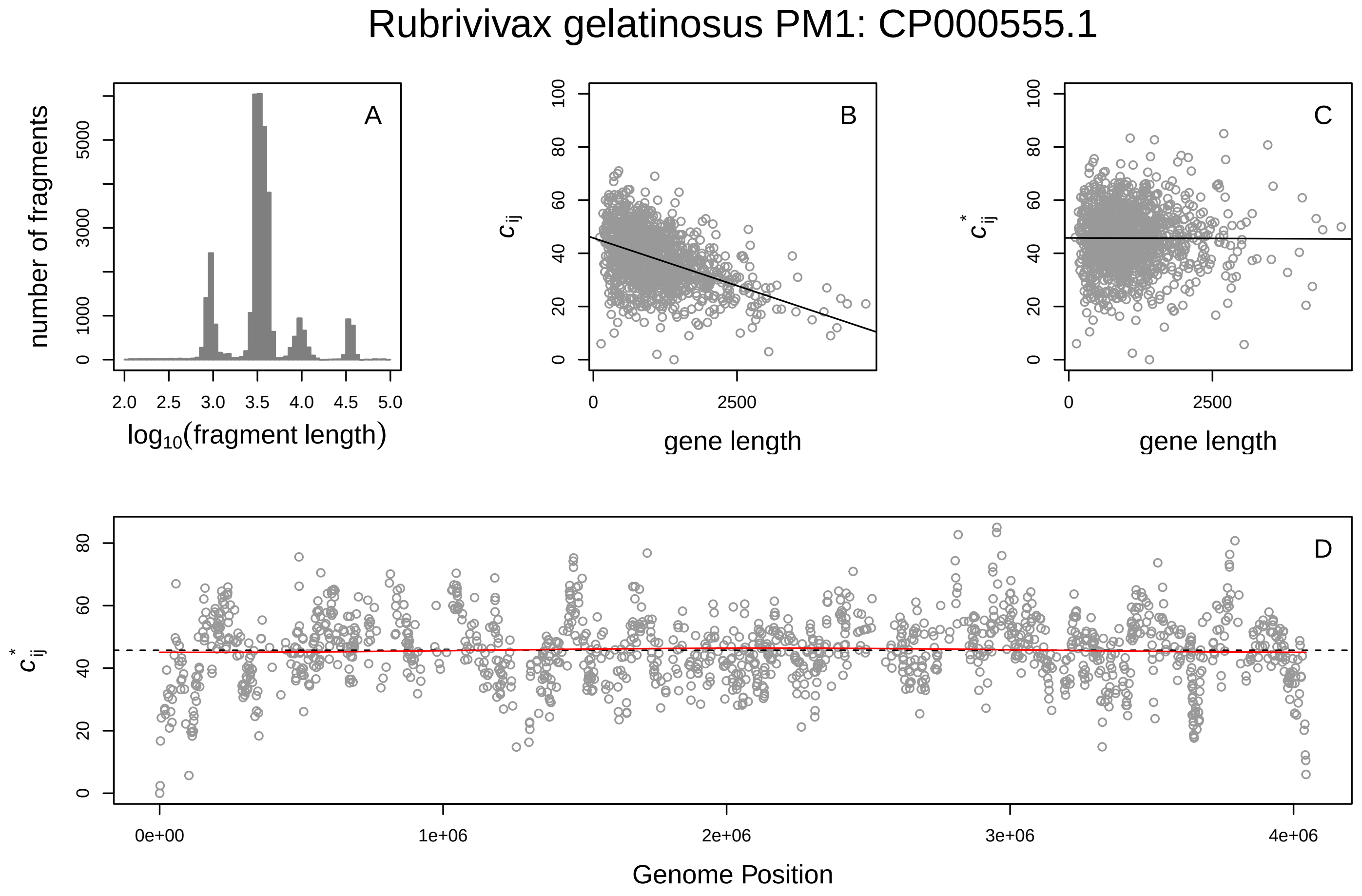
Rubrivivax gelatinosus PM1: CP000555.1
A
number of fragments
log10(fragment length)
B
c_ij
gene length
C
c*_ij
gene length
D
c*_ij
Genome Position

### Rubrobacter xylanophilus DSM 9941: CP000386.1

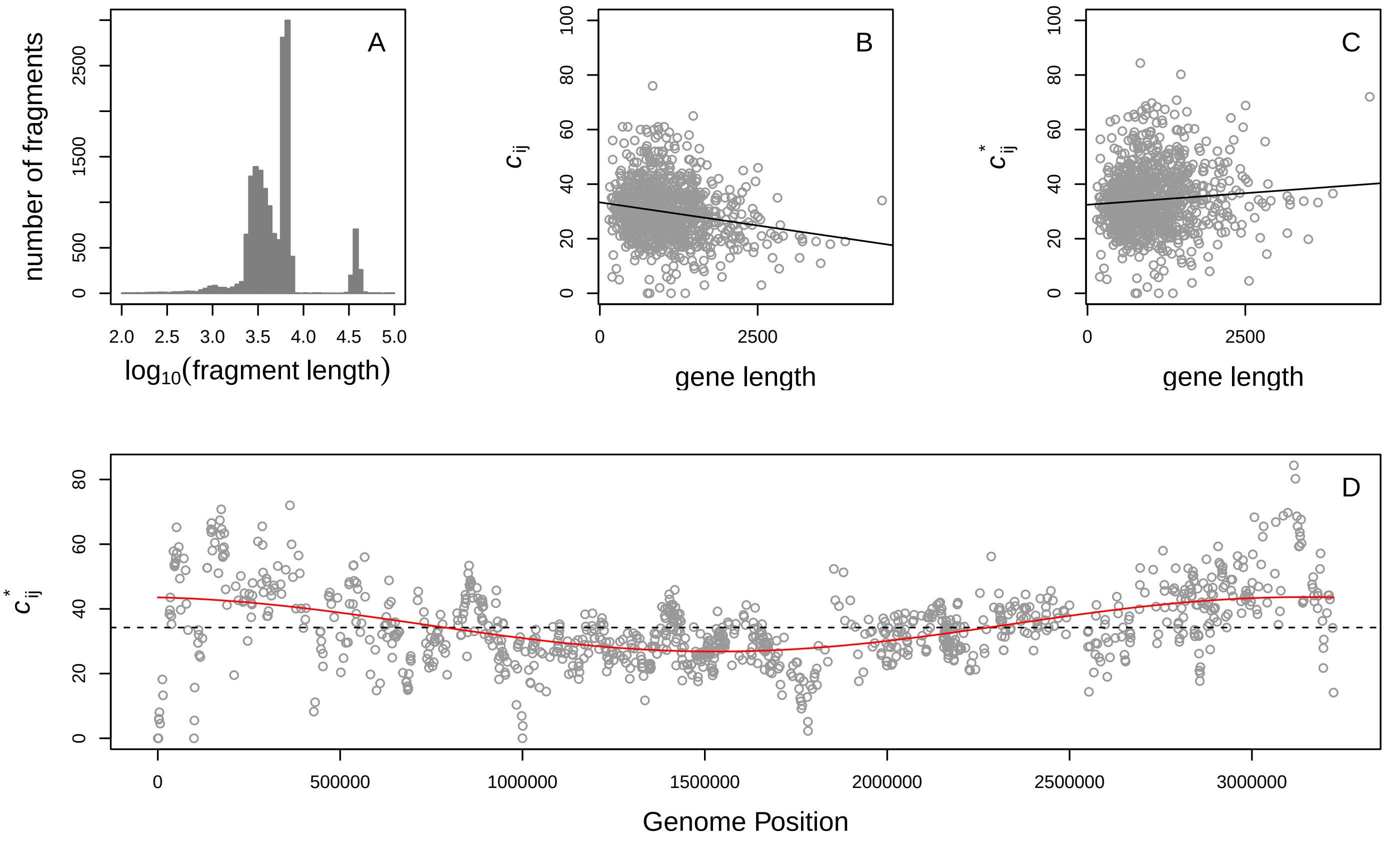


### Shewanella frigidimarina NCNM400: CP000447.1

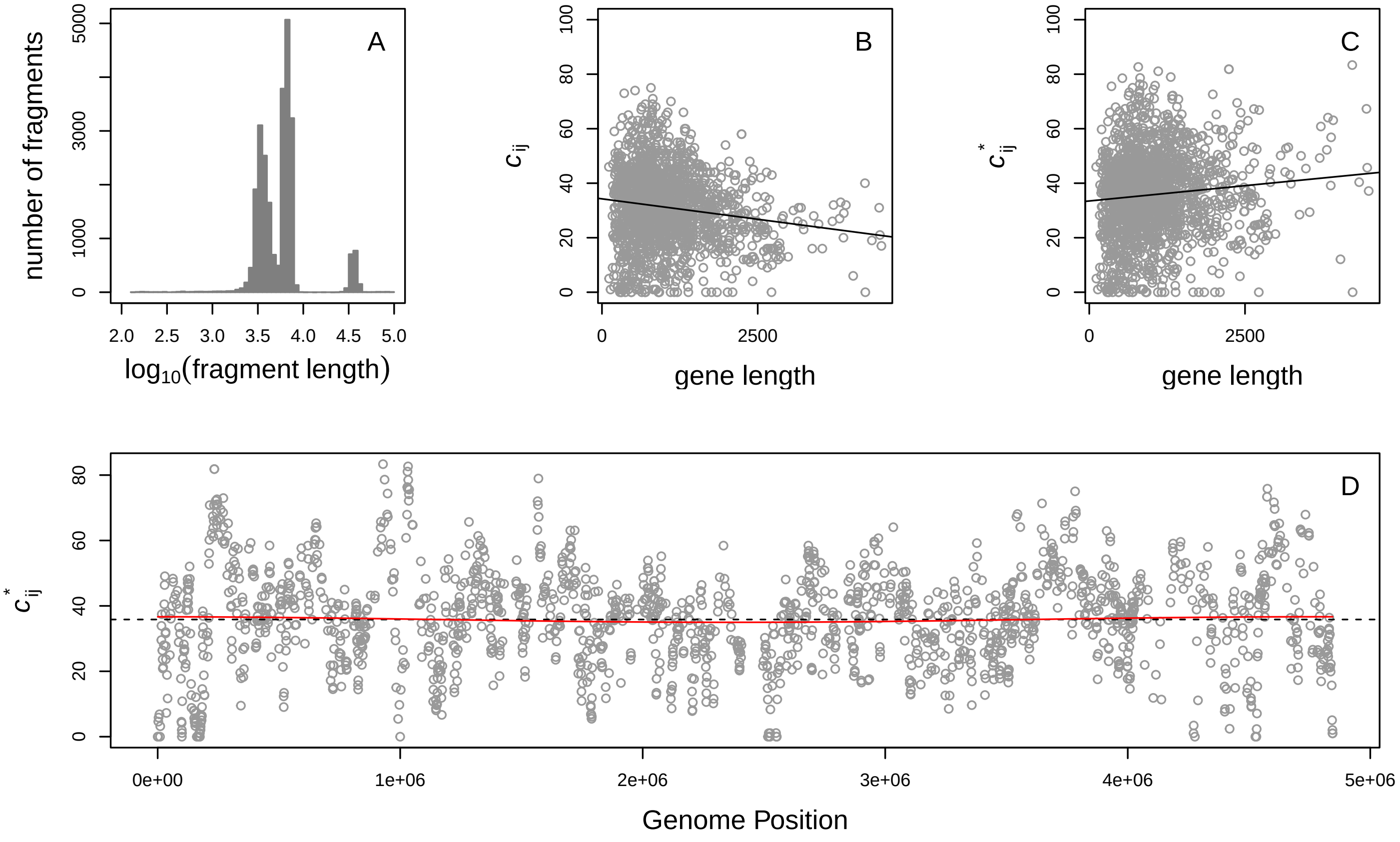

Shewanella sp ANA−3: CP000469.1

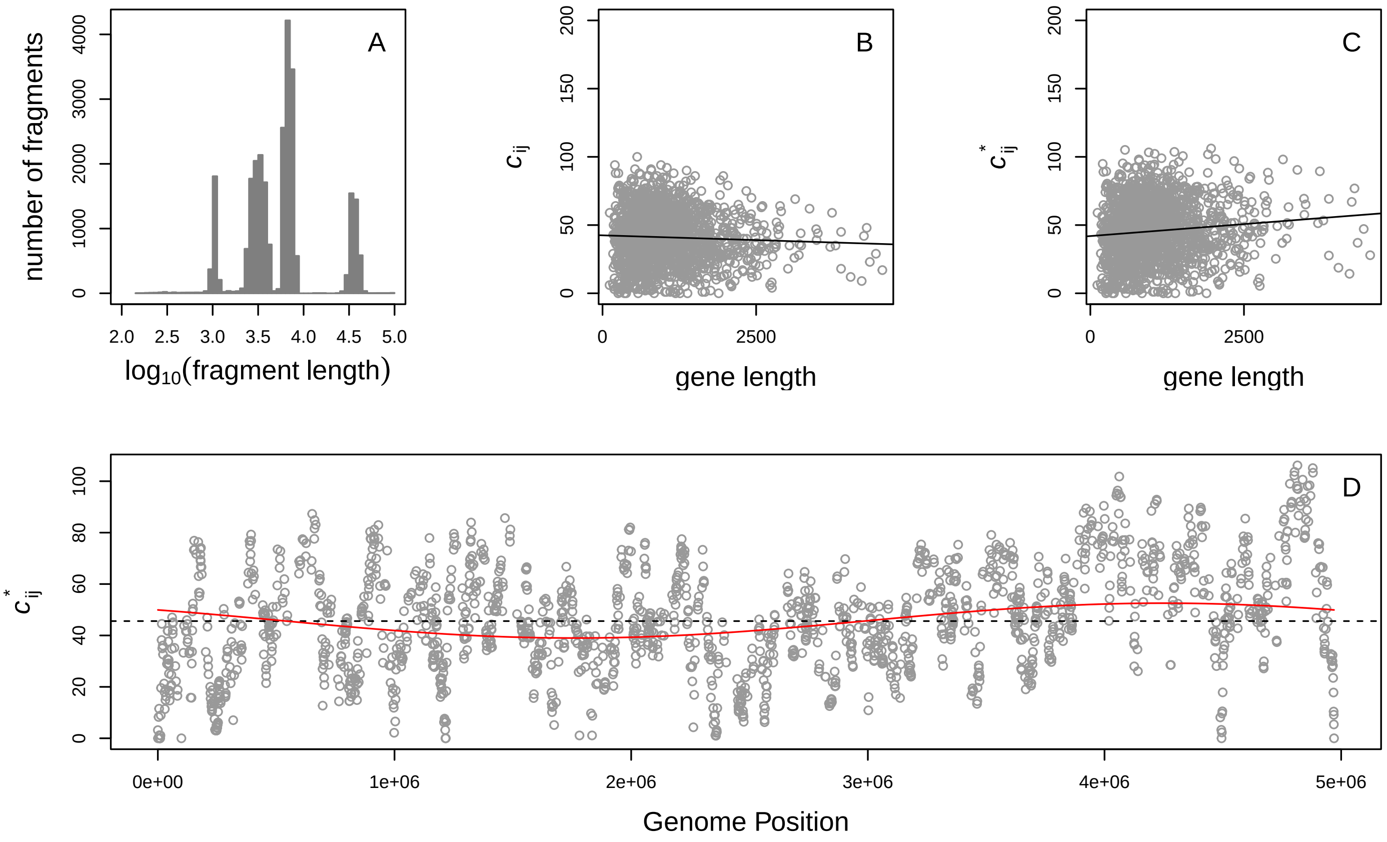


Silicibacter sp TM1040: CP000377.1

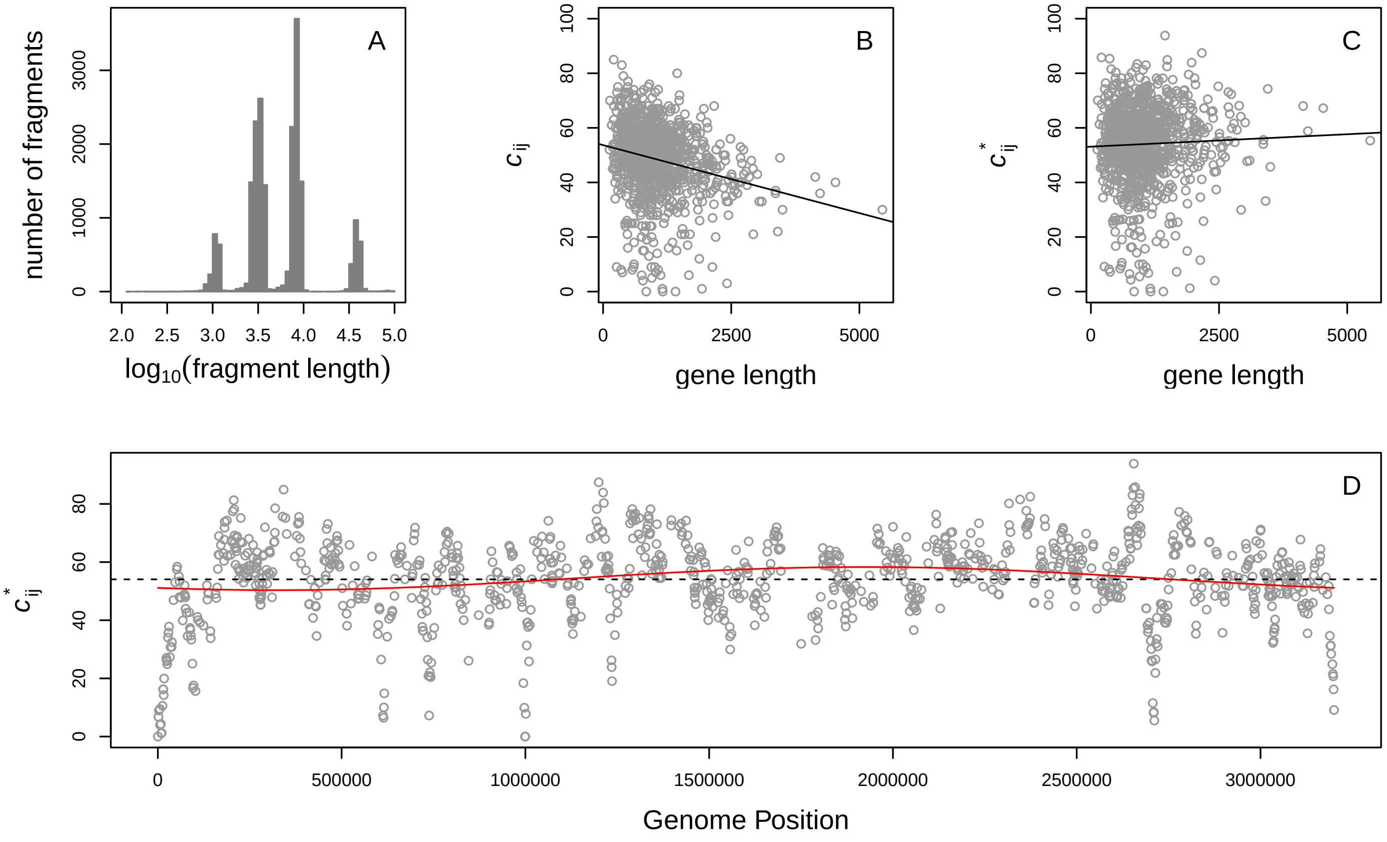

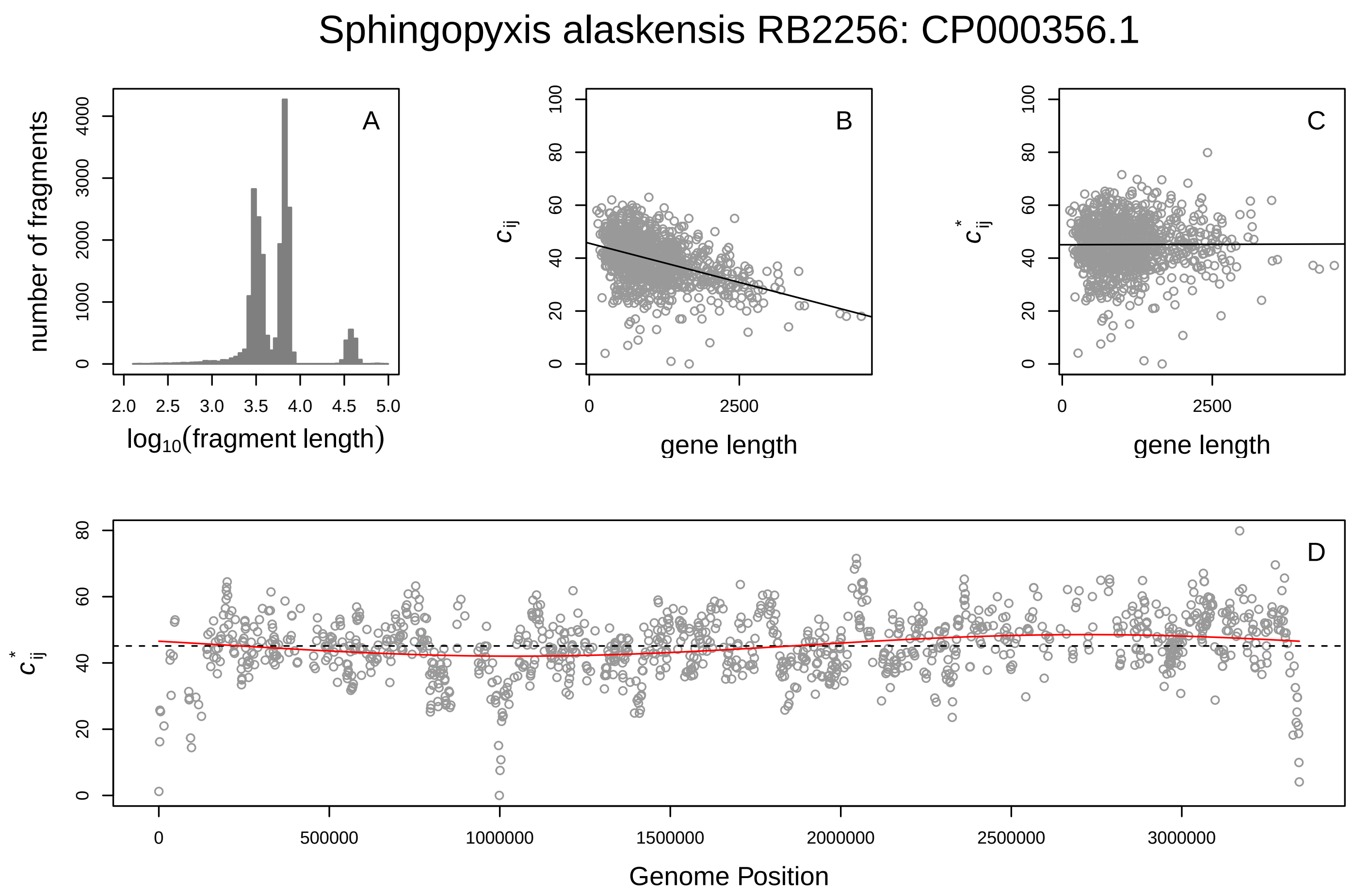
Sphingopyxis alaskensis RB2256: CP000356.1
A
number of fragments
log10(fragment length)
B
c_ij
gene length
C
c*_ij
gene length
D
c*_ij
Genome Position

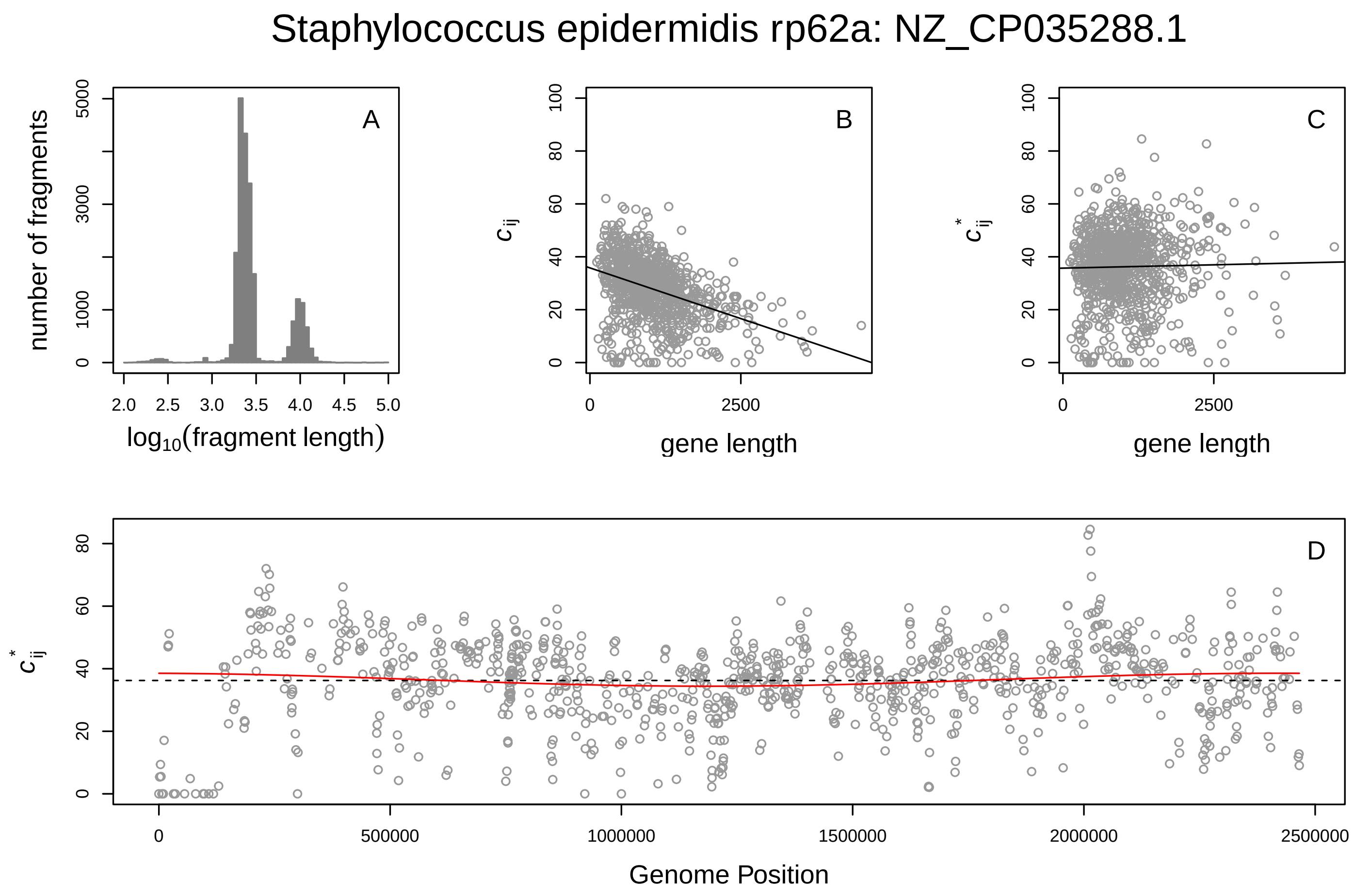
Staphylococcus epidermidis rp62a: NZ_CP035288.1
A
number of fragments
log10(fragment length)
B
c_ij
gene length
C
c*_ij
gene length
D
c*_ij
Genome Position

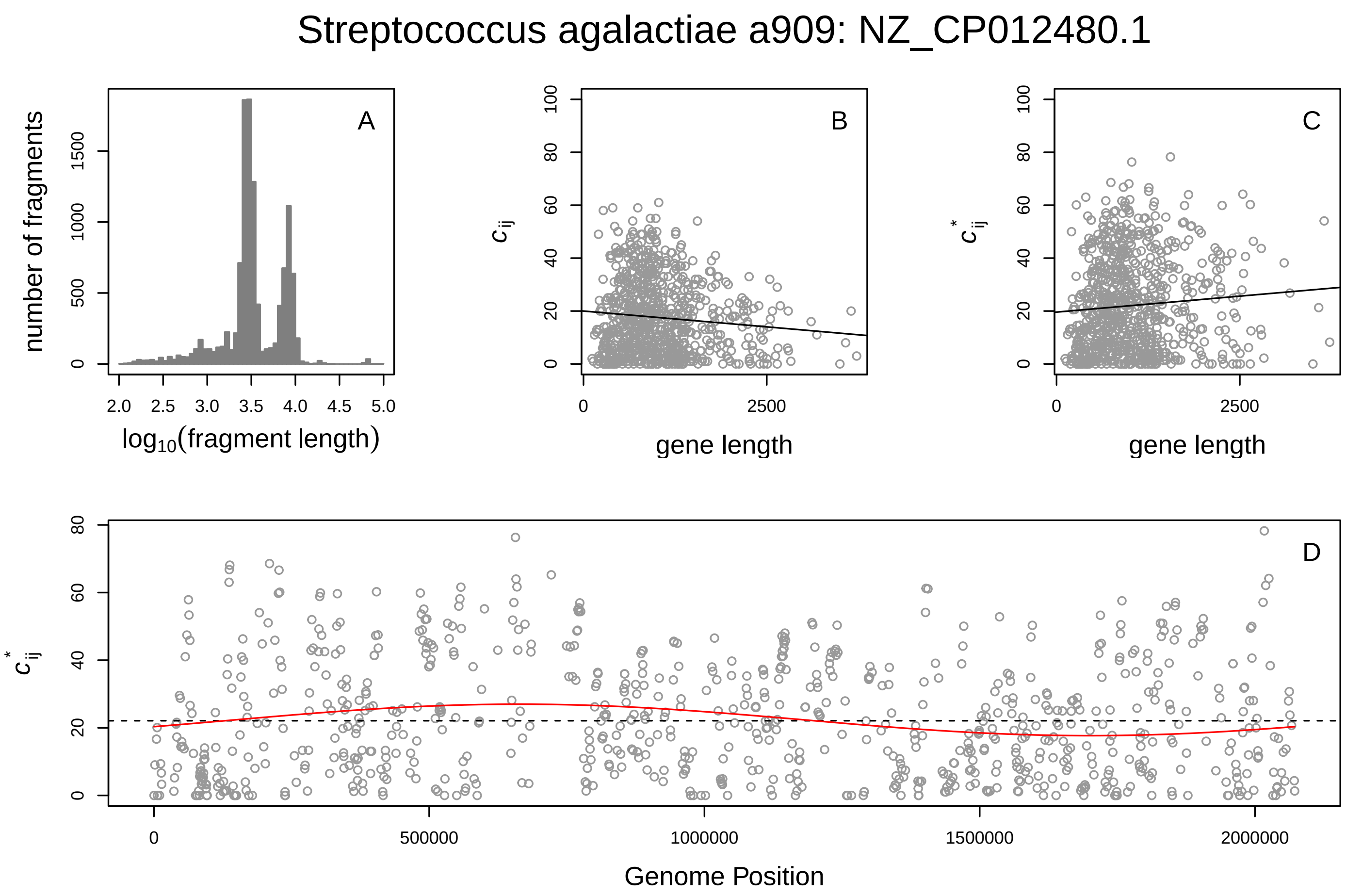

Streptococcus agalactiae a909: NZ_CP012480.1
A
B
C
D
number of fragments
log10(fragment length)
$c_{ij}$
$c^*_{ij}$
gene length
Genome Position


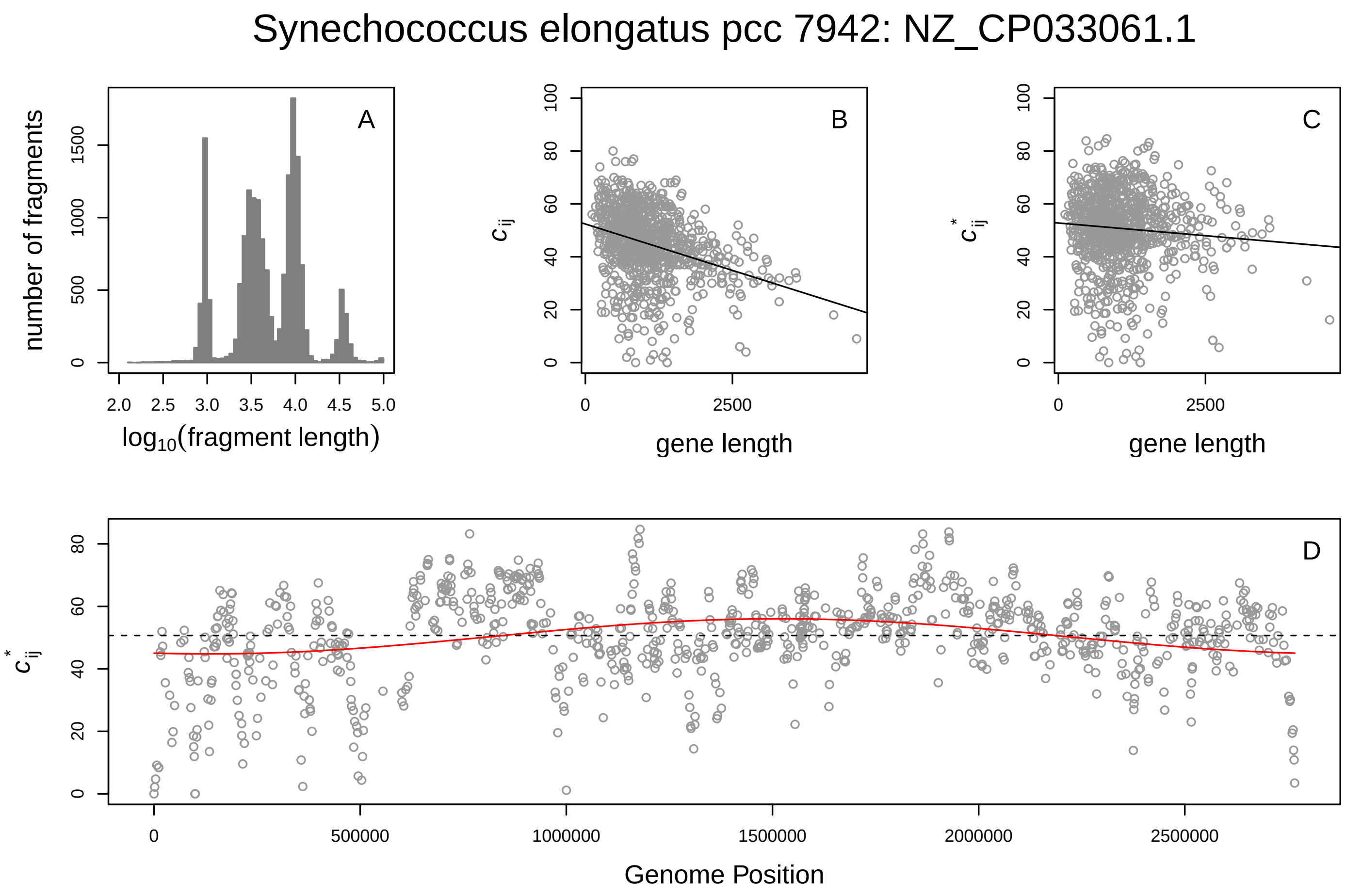

Synechococcus elongatus pcc 7942: NZ_CP033061.1
A
B
C
D
number of fragments
log10(fragment length)
$c_{ij}$
$c^*_{ij}$
gene length
Genome Position

#### Synechococcus sp cc9605: NC_007516.1

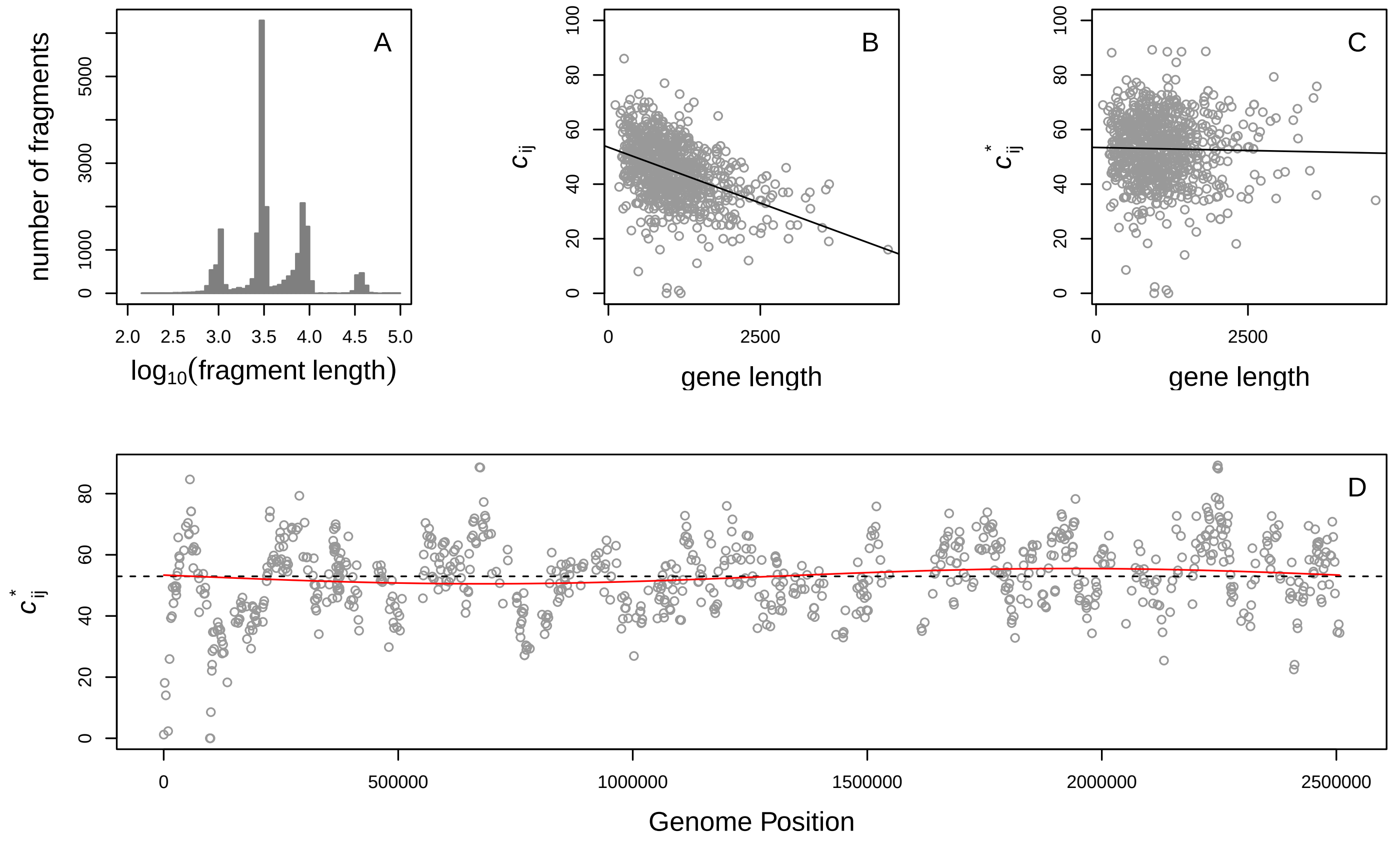


#### Thermobifida fusca YX: NC_007333.1

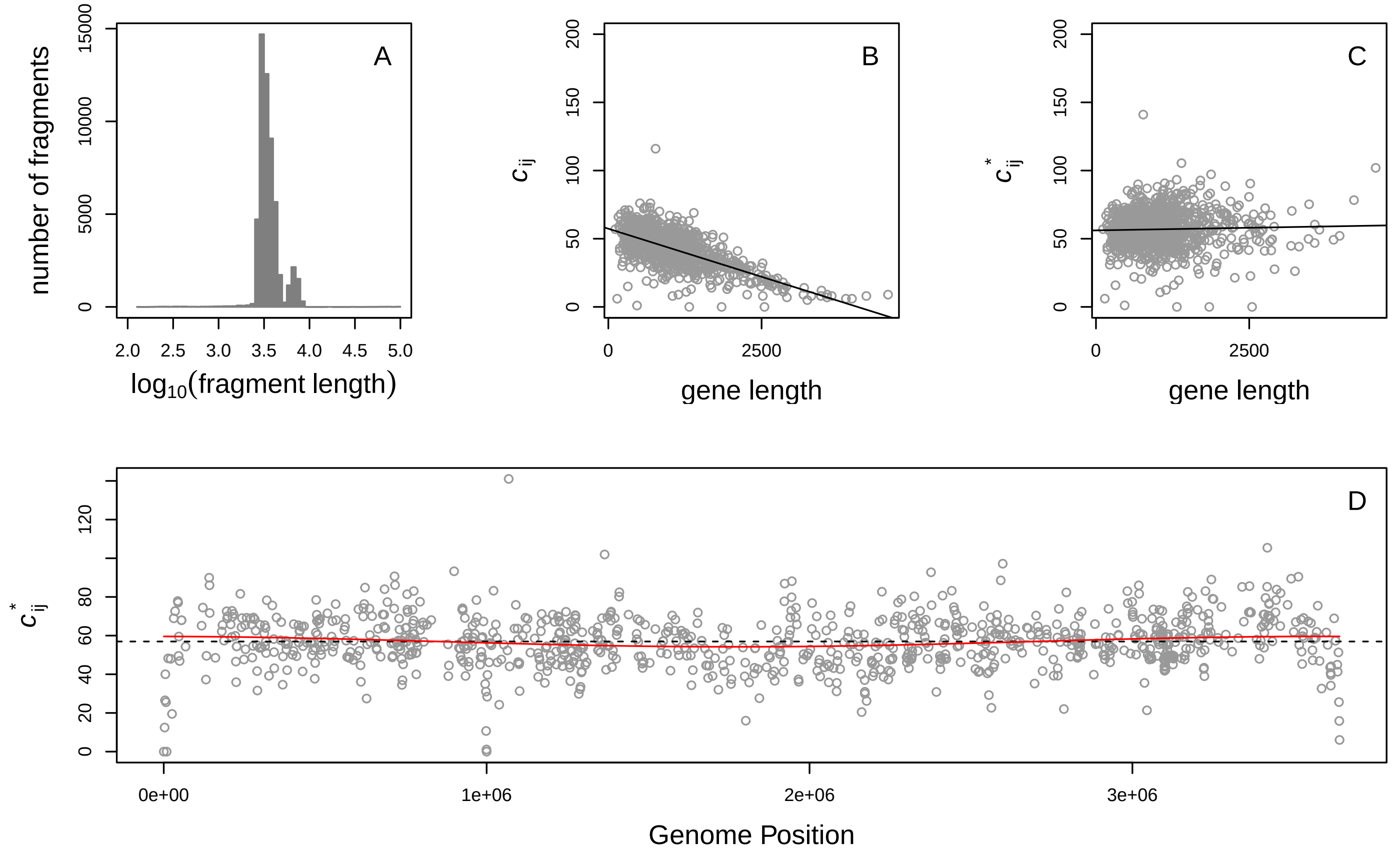

Thiomicrospira crunogena xcl−2: NC_007520.2

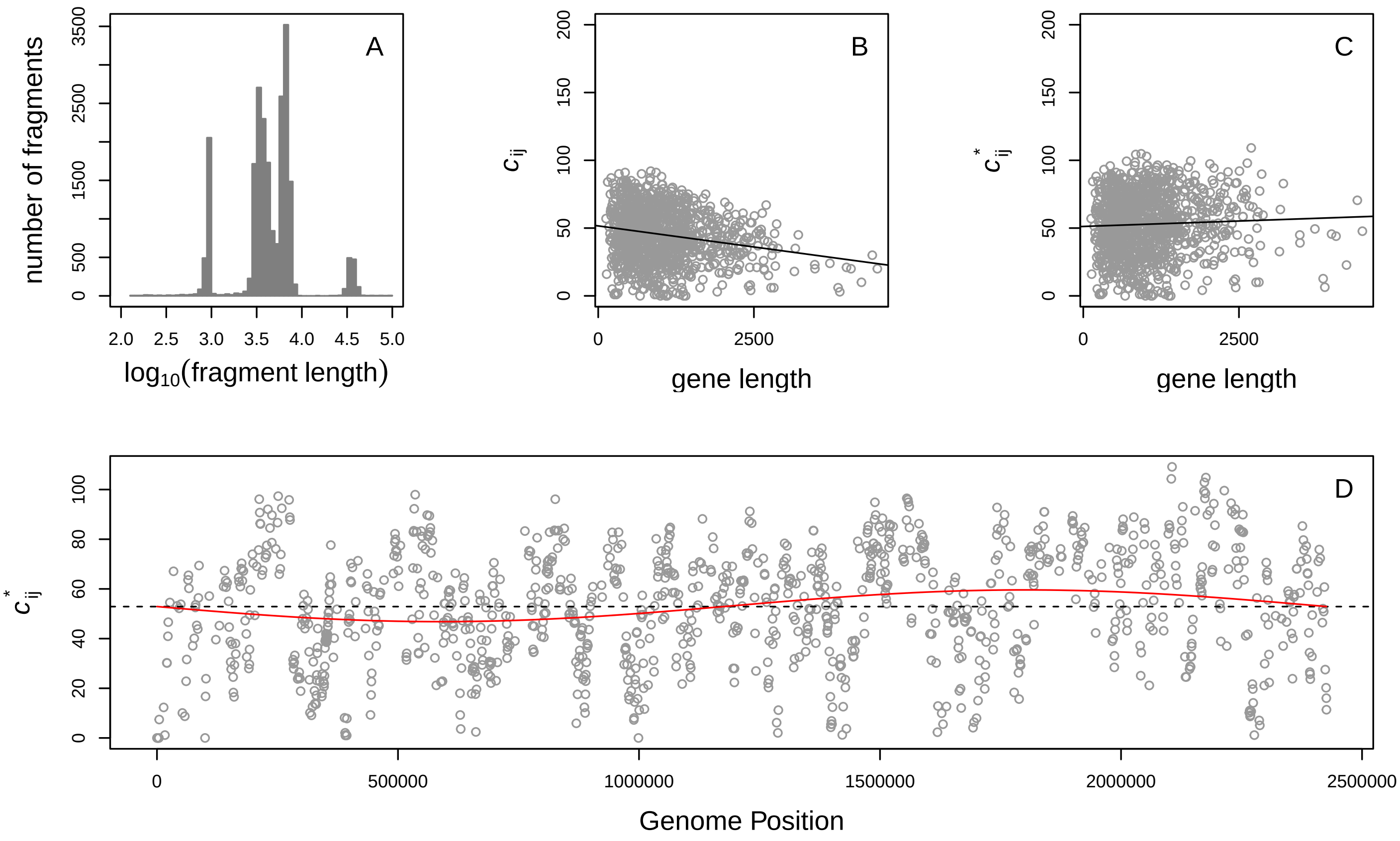


Thiomicrospira denitrificans ATCC 33889: CP000153.1

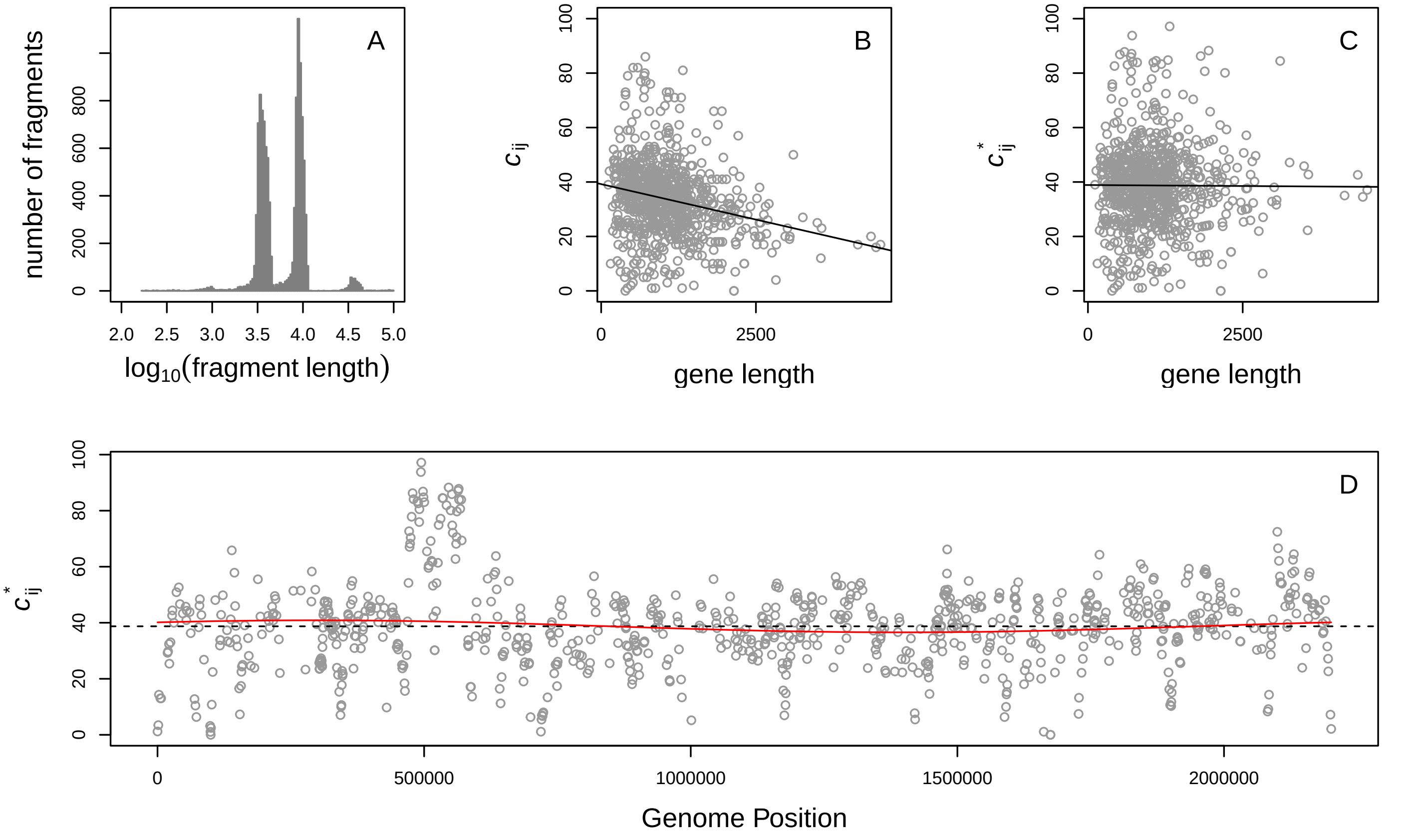

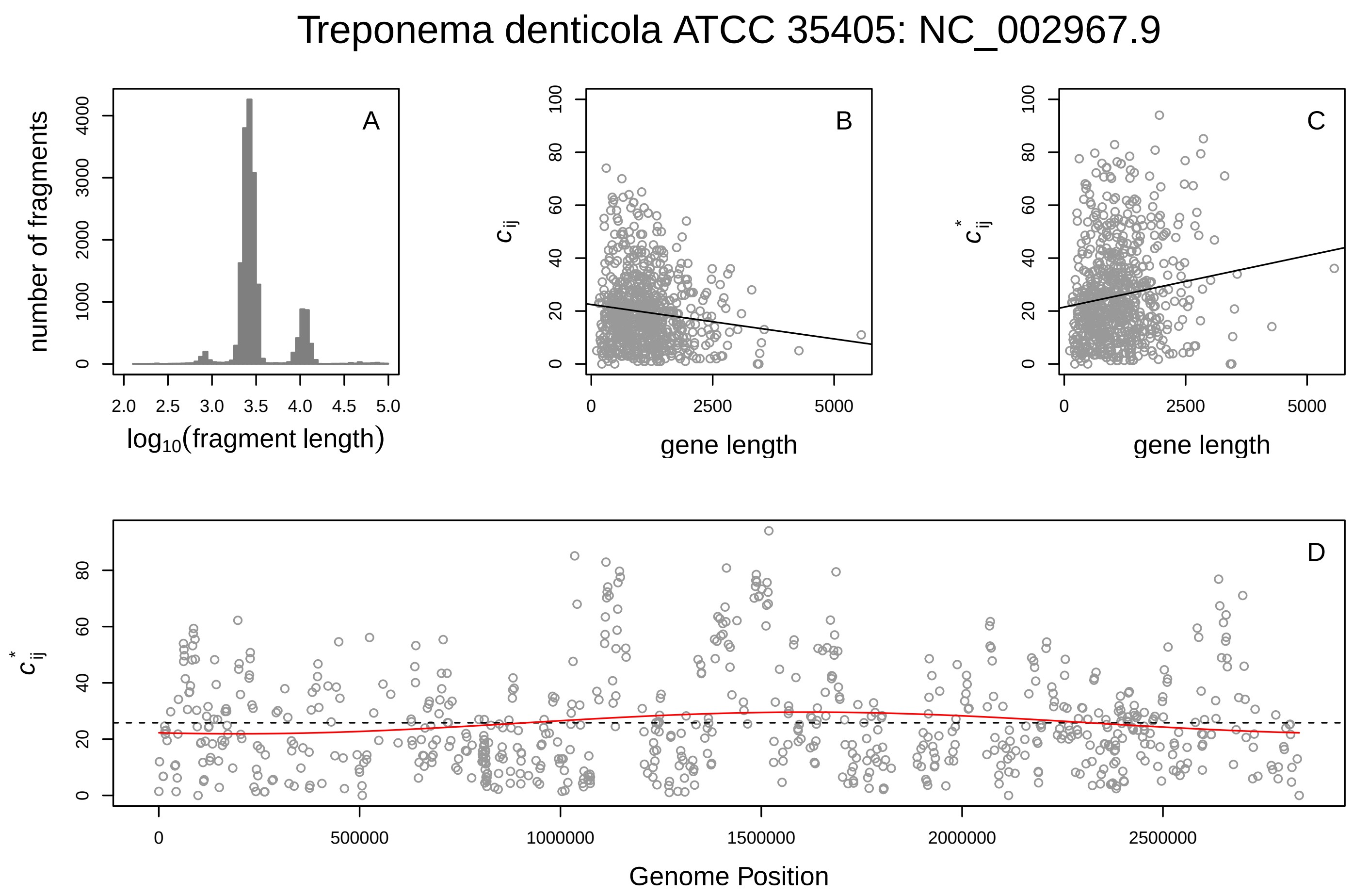
Treponema denticola ATCC 35405: NC_002967.9
A
number of fragments
log10(fragment length)
B
c_ij
gene length
C
c*_ij
gene length
D
c*_ij
Genome Position

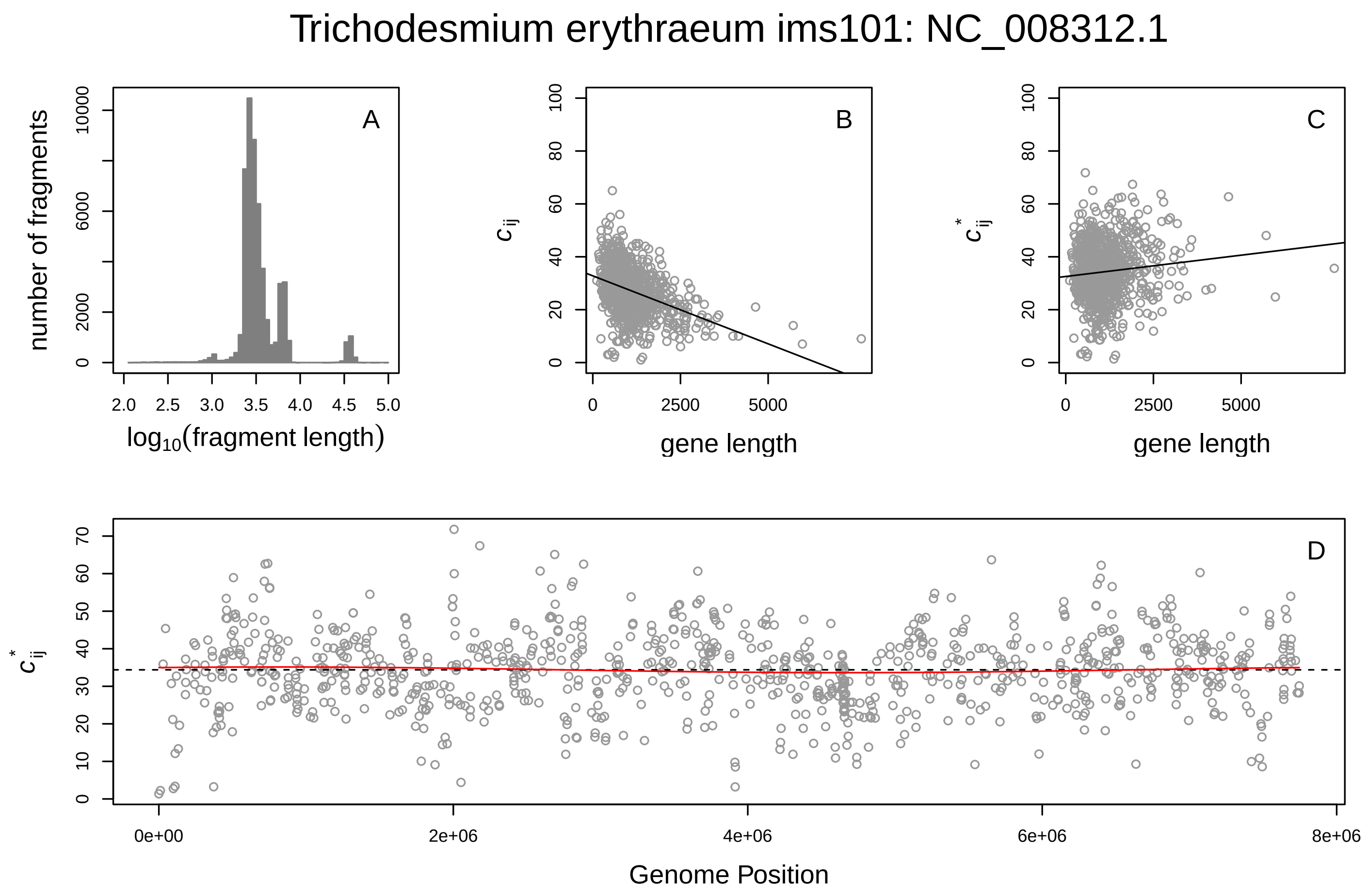
Trichodesmium erythraeum ims101: NC_008312.1
A
number of fragments
log10(fragment length)
B
c_ij
gene length
C
c*_ij
gene length
D
c*_ij
Genome Position

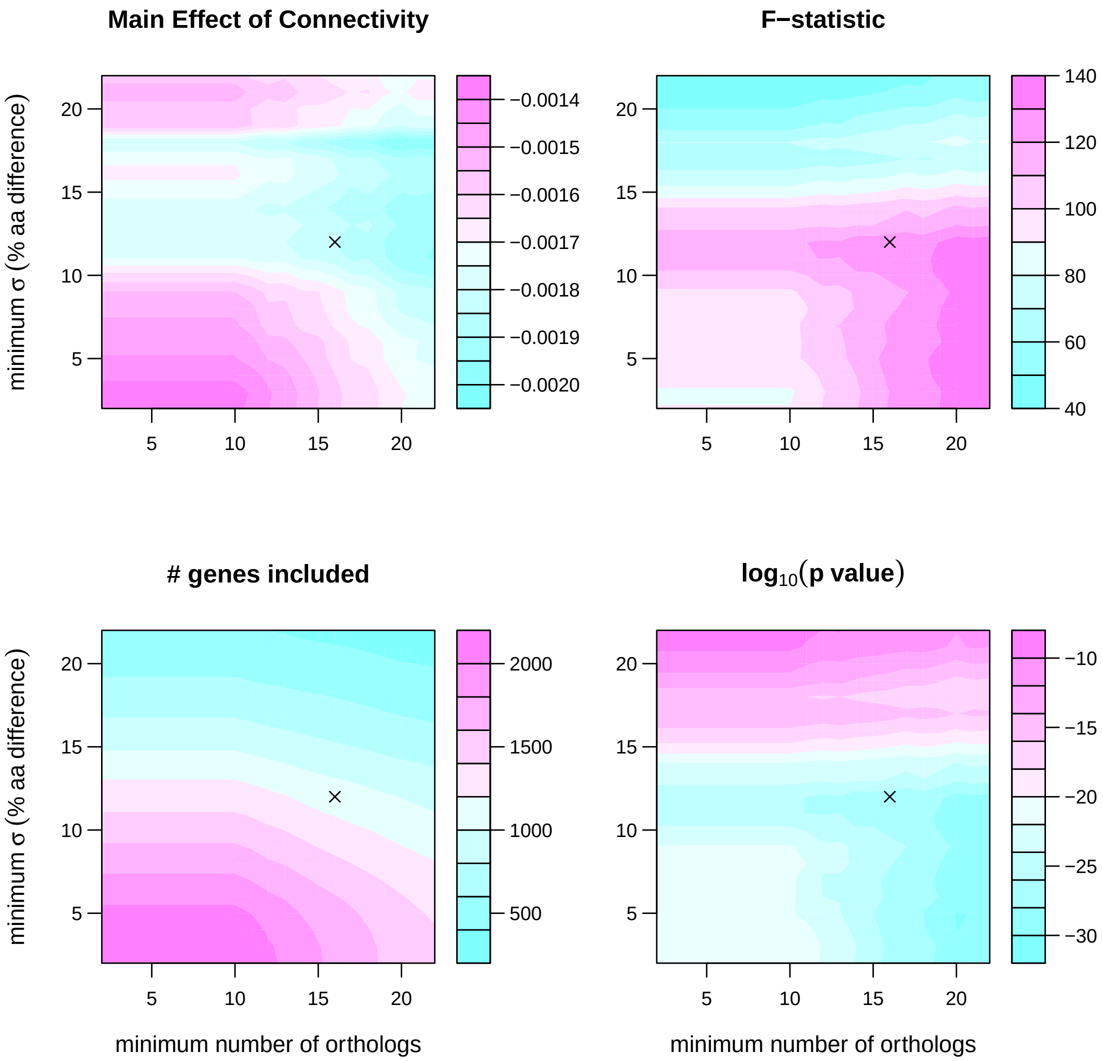


**Figure S2. Statistical sensitivity of the estimated main effect of connectivity on transferability.** We repeated the analysis of the effect of connectivity on transferability described in the main text, after restricting the dataset included in the analysis to different degrees. The repeated analyses required that each set of orthologs (of a particular gene in the *E. coli* K12 genome) include at least X minimum number of orthologs and exhibit a standard deviation greater than Y% amino acid difference. X and Y were both varied between 2 and 22. For each pair of X and Y, we estimated the number of *E. coli* genes that met both requirements (A), the estimated main effect of connectivity on transferability (B), and the F statistic (C) and p value (D) associated with the statistical test of the effect. The $\times$ in each plot shows the analysis described in the main text, that required $\geq 16$ orthologs and $\geq 12\%$ amino acid divergence.

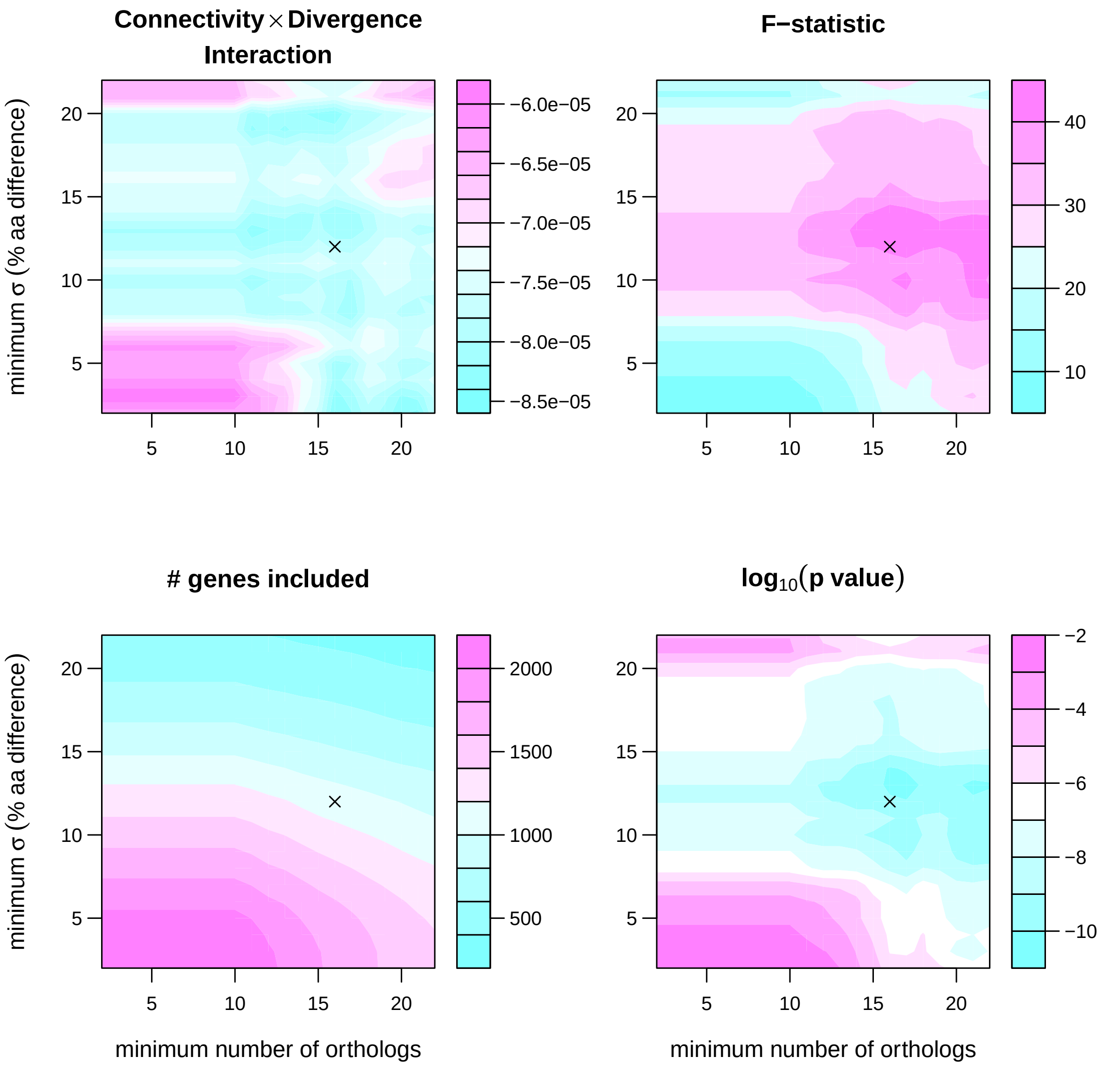


**Figure S3. Statistical sensitivity of the estimated effect on transferability of the connectivity × divergence interaction.** We repeated the analysis of the effect of connectivity on the slope of the relationship between divergence and transferability (see fig. 1), after restricting the dataset included in the analysis to different degrees. The repeated analyses required that each set of orthologs (of a particular gene in the *E. coli* K12 genome) include at least X minimum number of orthologs and exhibit a standard deviation greater than Y% amino acid difference. X and Y were both varied between 2 and 22. For each pair of X and Y, we estimated the number of genes that met both requirements (A), the estimated connectivity $\times$ divergence interaction effect (B), and the F statistic (C) and p value (D) associated with the statistical test. The $\times$ in each plot shows the analysis described in the main text, that required $\geq 16$ orthologs and $\geq 12\%$ amino acid divergence.

**Figure S4. Effect of divergence on transferability for orthologs of *E. coli* coding sequences.** Points in each plot show transferability, estimated as relative coverage ($\hat{c}_{ij}$), and divergence (% amino acid difference from the *E. coli* ortholog) of coding sequences in other bacterial genomes that were identified as likely orthologs of the *E. coli* gene indicated at the top of the plot (see *Methods: Effect of divergence on coverage*). For genes for which we identified likely orthologs from at least 16 of the genomes in our collection, dashed lines represent the mean transferability across all orthologs and solid lines represent the best fit linear model.

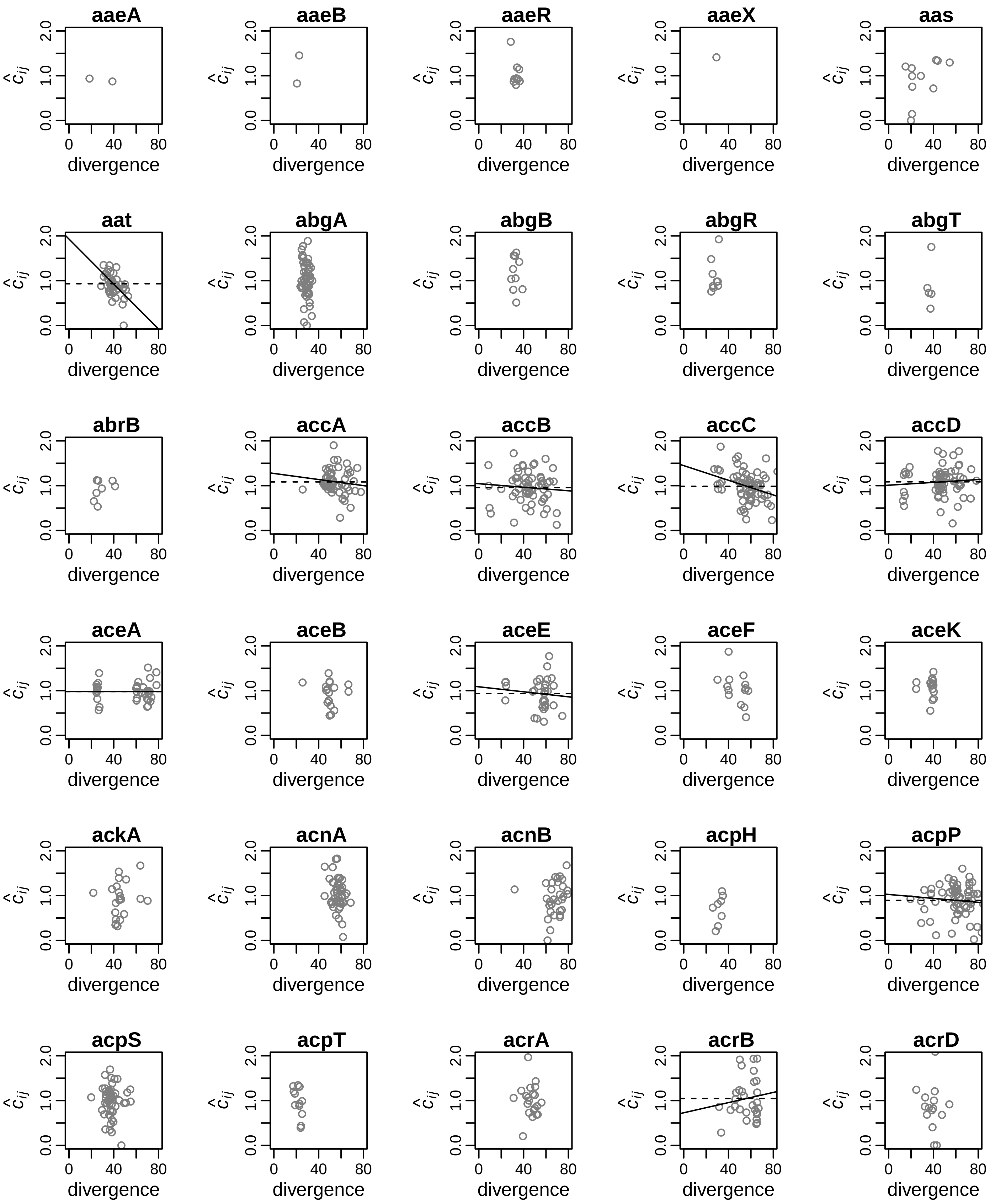
aaeA
aaeB
aaeR
aaeX
aas
aat
abgA
abgB
abgR
abgT
abrB
accA
accB
accC
accD
aceA
aceB
aceE
aceF
aceK
ackA
acnA
acnB
acpH
acpP
acpS
acpT
acrA
acrB
acrD
$\hat{c}_{ij}$
0.0
1.0
2.0
0
40
80
divergence

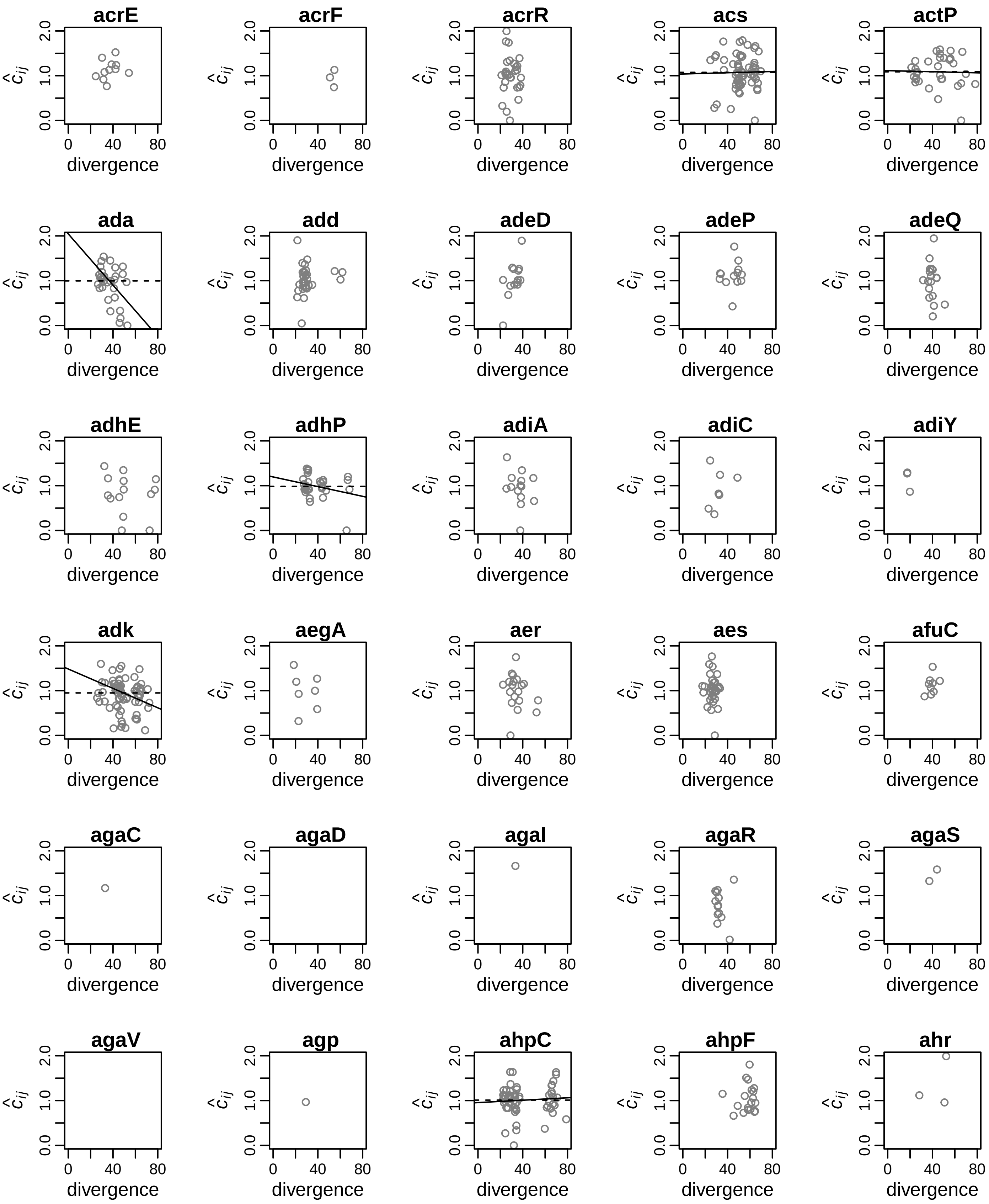
acrE
acrF
acrR
acs
actP
ada
add
adeD
adeP
adeQ
adhE
adhP
adiA
adiC
adiY
adk
aegA
aer
aes
afuC
agaC
agaD
agaI
agaR
agaS
agaV
agp
ahpC
ahpF
ahr
$\hat{c}_{ij}$
0.0
1.0
2.0
0
40
80
divergence

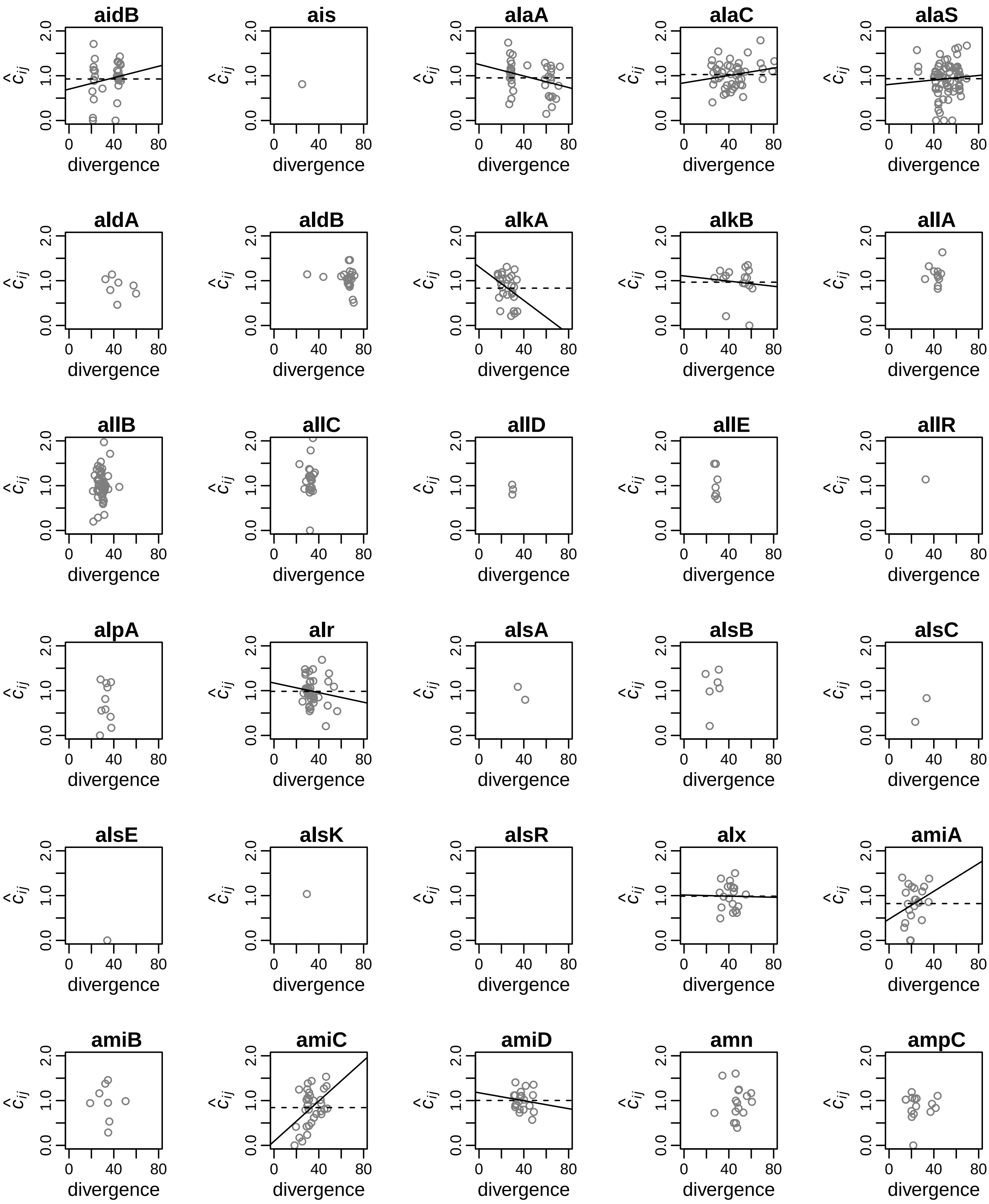
aidB
ais
alaA
alaC
alaS
aldA
aldB
alkA
alkB
allA
allB
allC
allD
allE
allR
alpA
alr
alsA
alsB
alsC
alsE
alsK
alsR
alx
amiA
amiB
amiC
amiD
amn
ampC
$\hat{c}_{ij}$
0.0
1.0
2.0
0
40
80
divergence

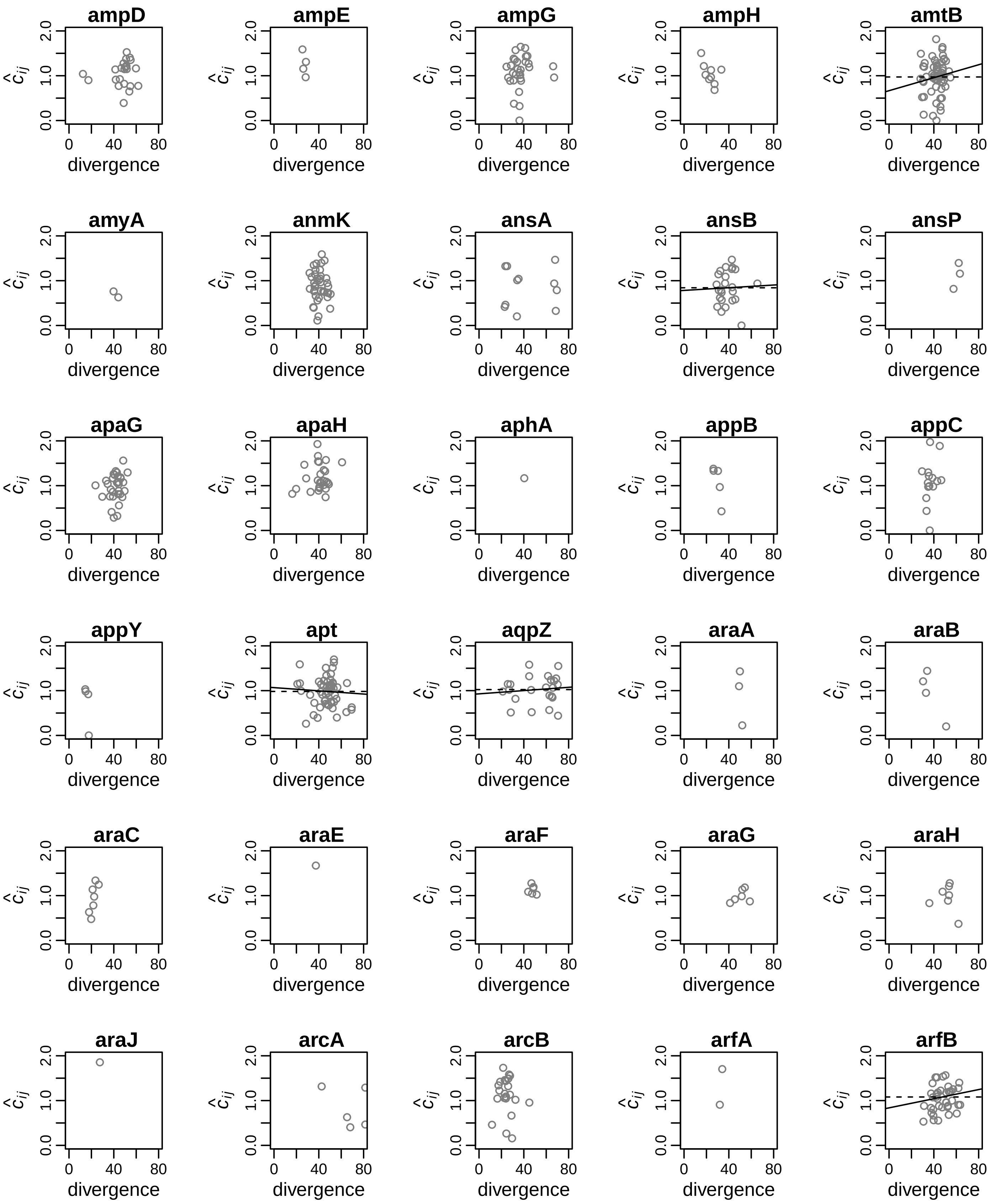
ampD
ampE
ampG
ampH
amtB
amyA
anmK
ansA
ansB
ansP
apaG
apaH
aphA
appB
appC
appY
apt
aqpZ
araA
araB
araC
araE
araF
araG
araH
araJ
arcA
arcB
arfA
arfB
$\hat{c}_{ij}$
0.0 1.0 2.0
0 40 80
divergence

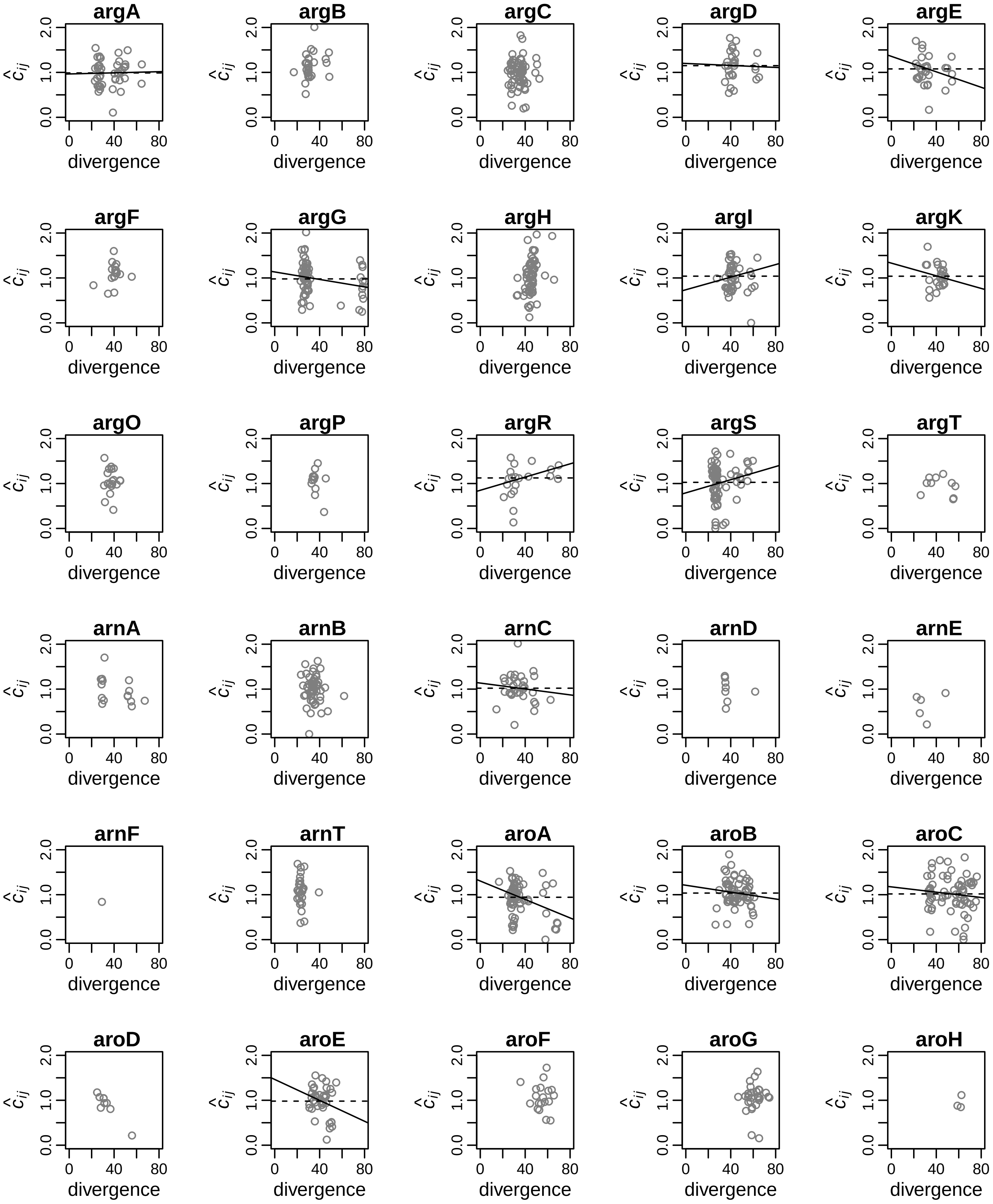
argA
argB
argC
argD
argE
argF
argG
argH
argI
argK
argO
argP
argR
argS
argT
arnA
arnB
arnC
arnD
arnE
arnF
arnT
aroA
aroB
aroC
aroD
aroE
aroF
aroG
aroH
$\hat{c}_{ij}$
0.0
1.0
2.0
divergence
0
40
80

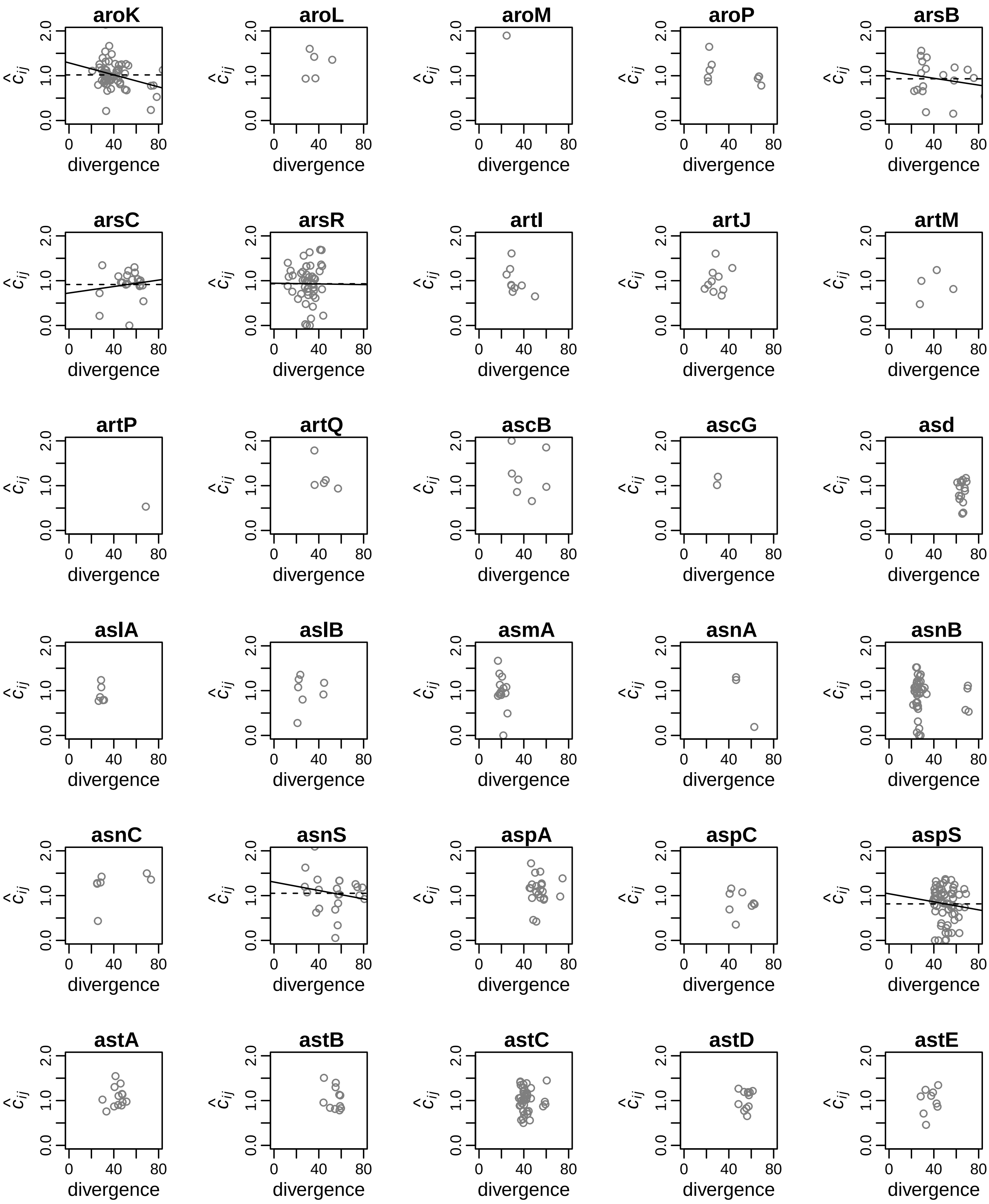
aroK
aroL
aroM
aroP
arsB
arsC
arsR
artI
artJ
artM
artP
artQ
ascB
ascG
asd
aslA
aslB
asmA
asnA
asnB
asnC
asnS
aspA
aspC
aspS
astA
astB
astC
astD
astE
$\hat{c}_{ij}$
divergence
0.0
1.0
2.0
0
40
80

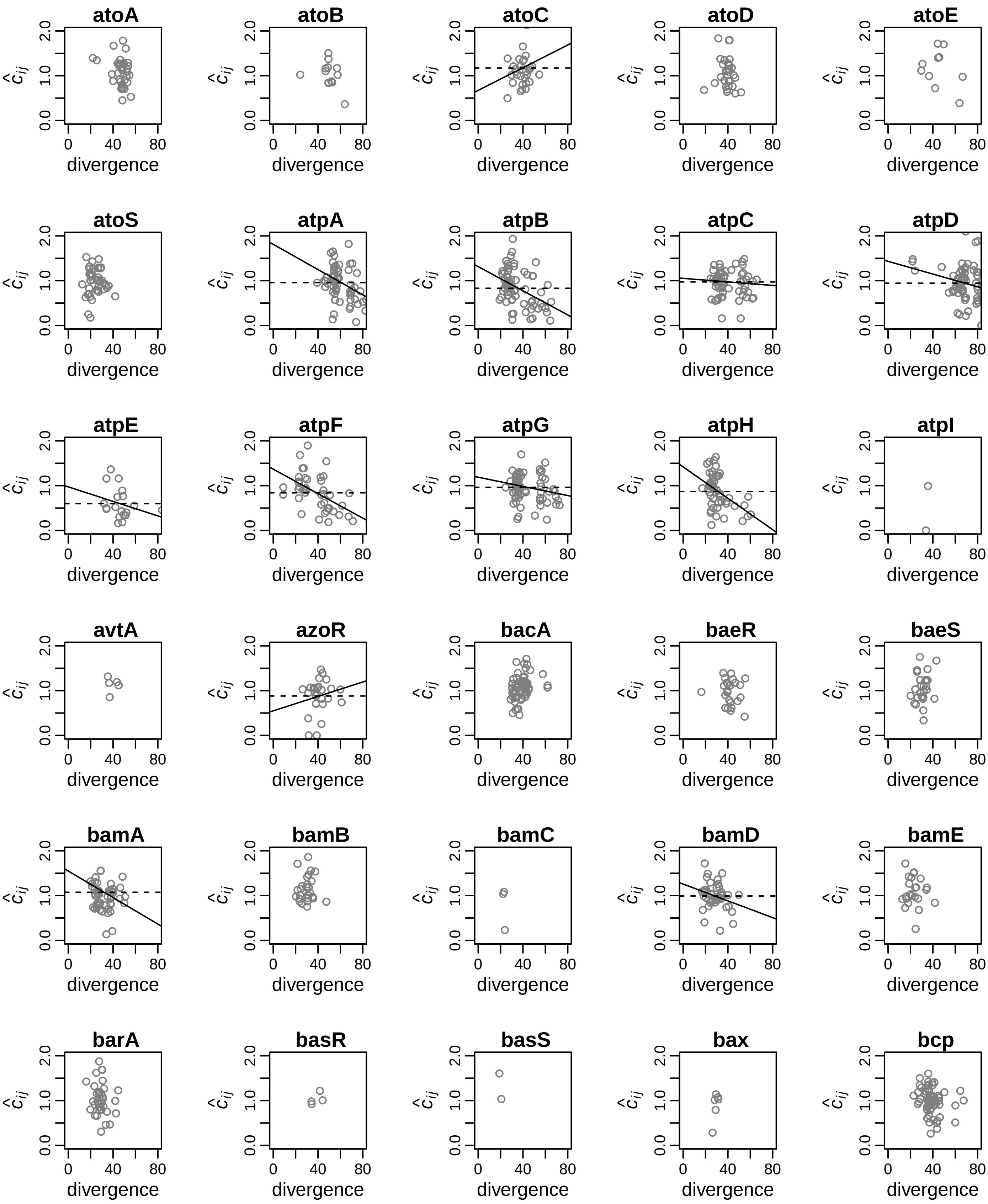
atoA
atoB
atoC
atoD
atoE
atoS
atpA
atpB
atpC
atpD
atpE
atpF
atpG
atpH
atpI
avtA
azoR
bacA
baeR
baeS
bamA
bamB
bamC
bamD
bamE
barA
basR
basS
bax
bcp
$\hat{c}_{ij}$
0.0
1.0
2.0
0
40
80
divergence

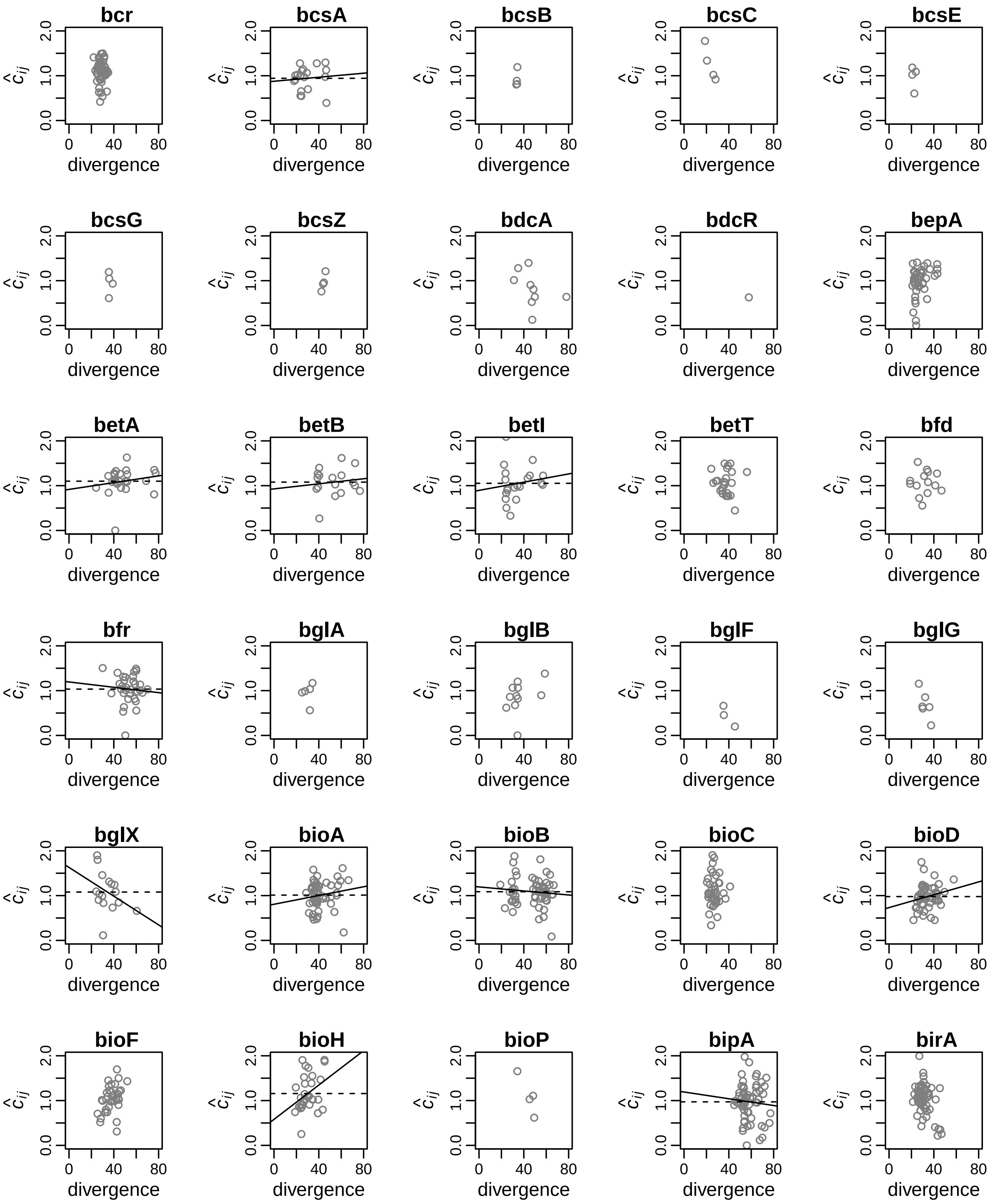
bcr
bcsA
bcsB
bcsC
bcsE
bcsG
bcsZ
bdcA
bdcR
bepA
betA
betB
betI
betT
bfd
bfr
bglA
bglB
bglF
bglG
bglX
bioA
bioB
bioC
bioD
bioF
bioH
bioP
bipA
birA
$\hat{c}_{ij}$
divergence
0.0
1.0
2.0
0
40
80

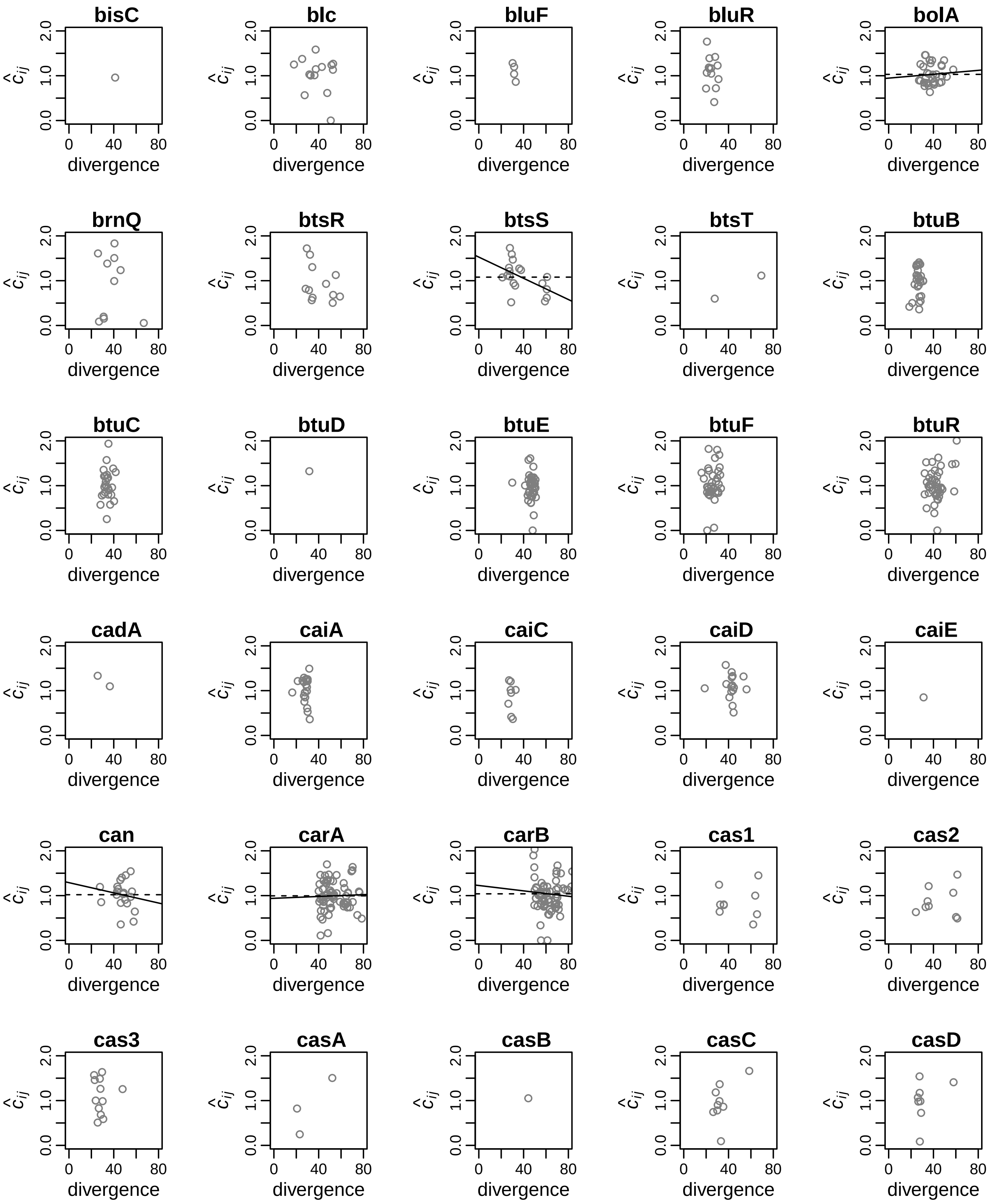
bisC
blc
bluF
bluR
bolA
brnQ
btsR
btsS
btsT
btuB
btuC
btuD
btuE
btuF
btuR
cadA
caiA
caiC
caiD
caiE
can
carA
carB
cas1
cas2
cas3
casA
casB
casC
casD
$\hat{c}_{ij}$
0.0
1.0
2.0
0
40
80
divergence

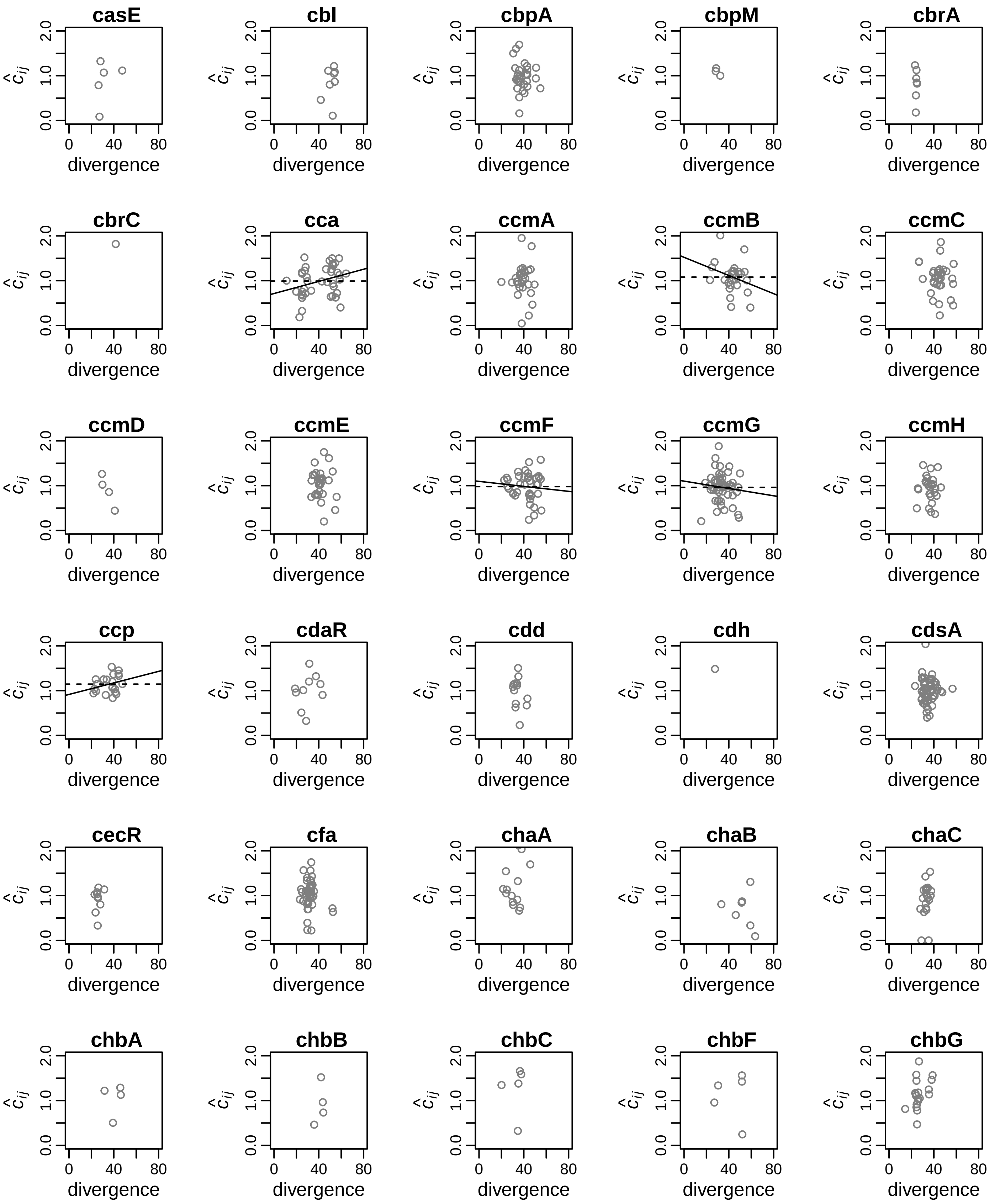
casE
cbl
cbpA
cbpM
cbrA
cbrC
cca
ccmA
ccmB
ccmC
ccmD
ccmE
ccmF
ccmG
ccmH
ccp
cdaR
cdd
cdh
cdsA
cecR
cfa
chaA
chaB
chaC
chbA
chbB
chbC
chbF
chbG
$\hat{c}_{ij}$
0.0
1.0
2.0
0
40
80
divergence

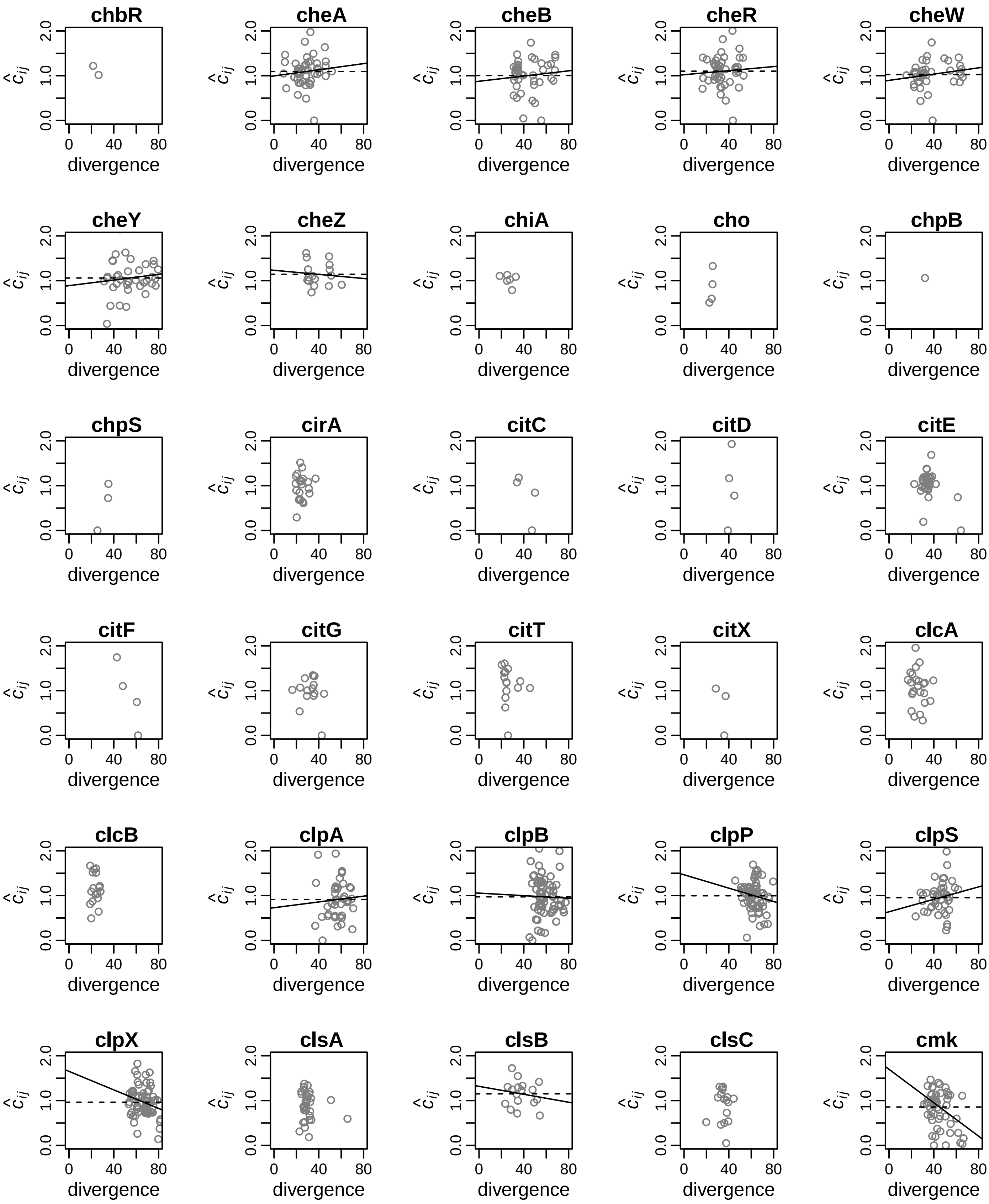
chbR
cheA
cheB
cheR
cheW
cheY
cheZ
chiA
cho
chpB
chpS
cirA
citC
citD
citE
citF
citG
citT
citX
clcA
clcB
clpA
clpB
clpP
clpS
clpX
clsA
clsB
clsC
cmk
$\hat{c}_{ij}$
0.0
1.0
2.0
0
40
80
divergence

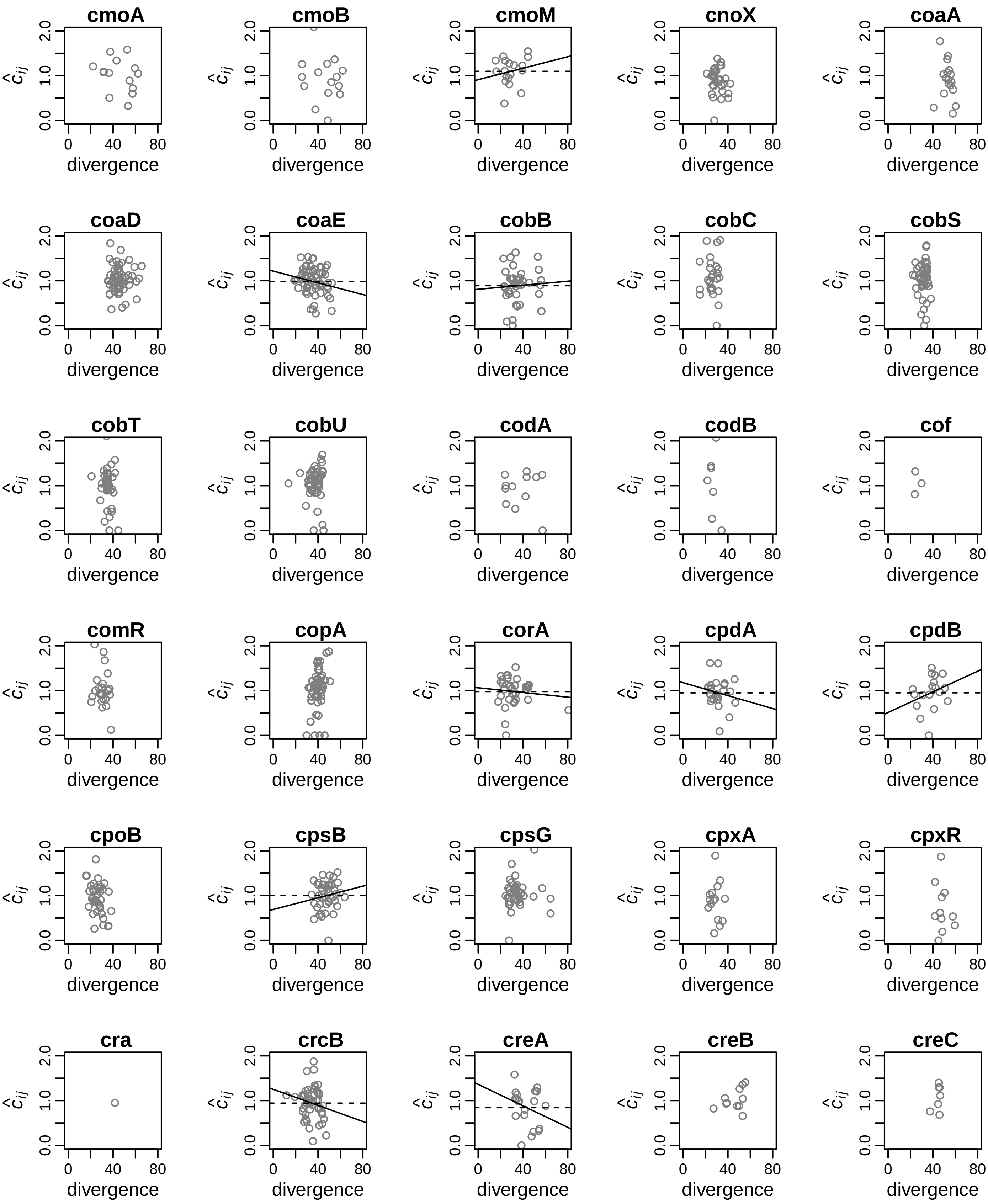
cmoA
cmoB
cmoM
cnoX
coaA
coaD
coaE
cobB
cobC
cobS
cobT
cobU
codA
codB
cof
comR
copA
corA
cpdA
cpdB
cpoB
cpsB
cpsG
cpxA
cpxR
cra
crcB
creA
creB
creC
$\hat{c}_{ij}$
0.0
1.0
2.0
divergence
0
40
80

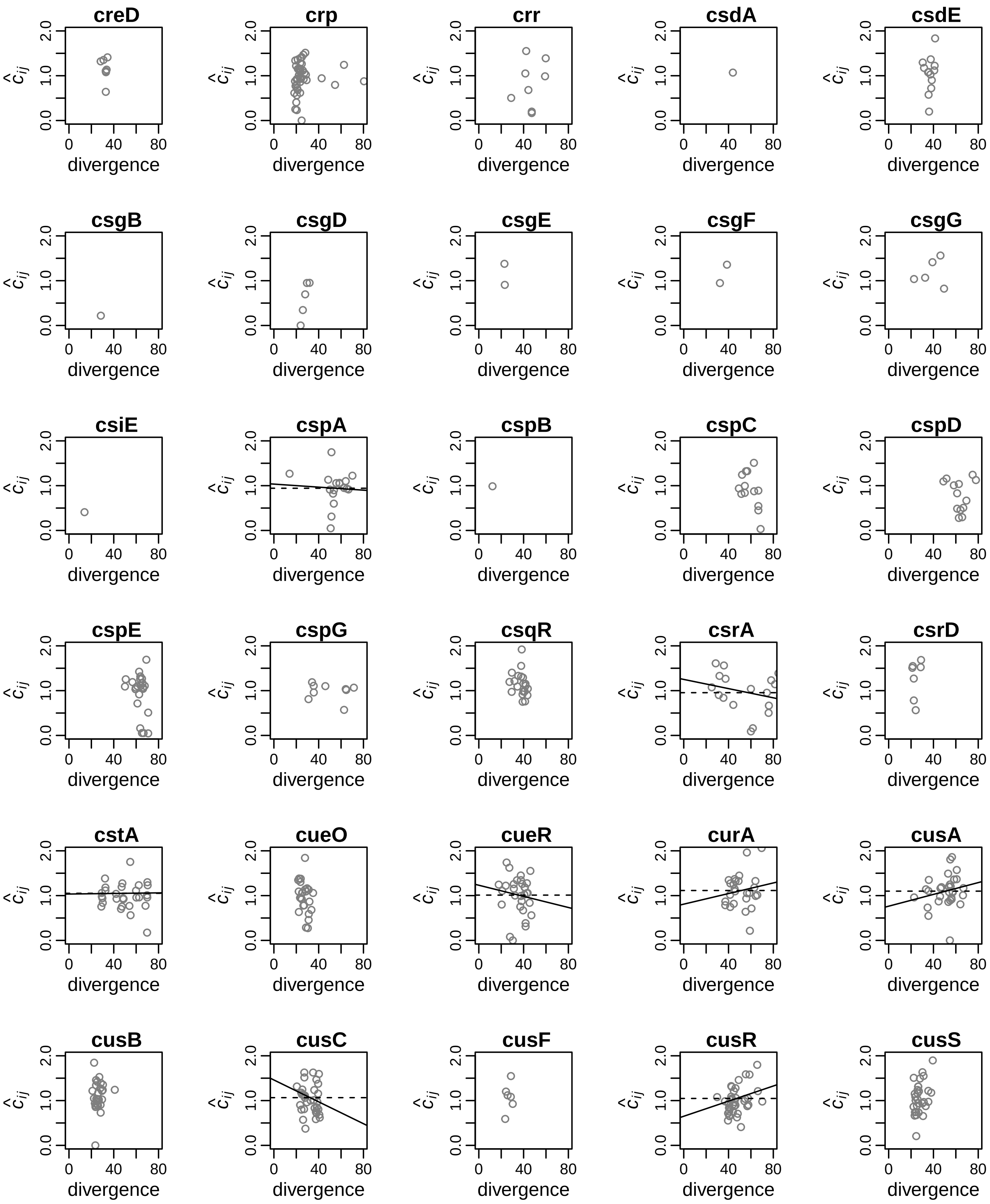
creD
crp
crr
csdA
csdE
csgB
csgD
csgE
csgF
csgG
csiE
cspA
cspB
cspC
cspD
cspE
cspG
csqR
csrA
csrD
cstA
cueO
cueR
curA
cusA
cusB
cusC
cusF
cusR
cusS
$\hat{c}_{ij}$
divergence
0.0
1.0
2.0
0
40
80

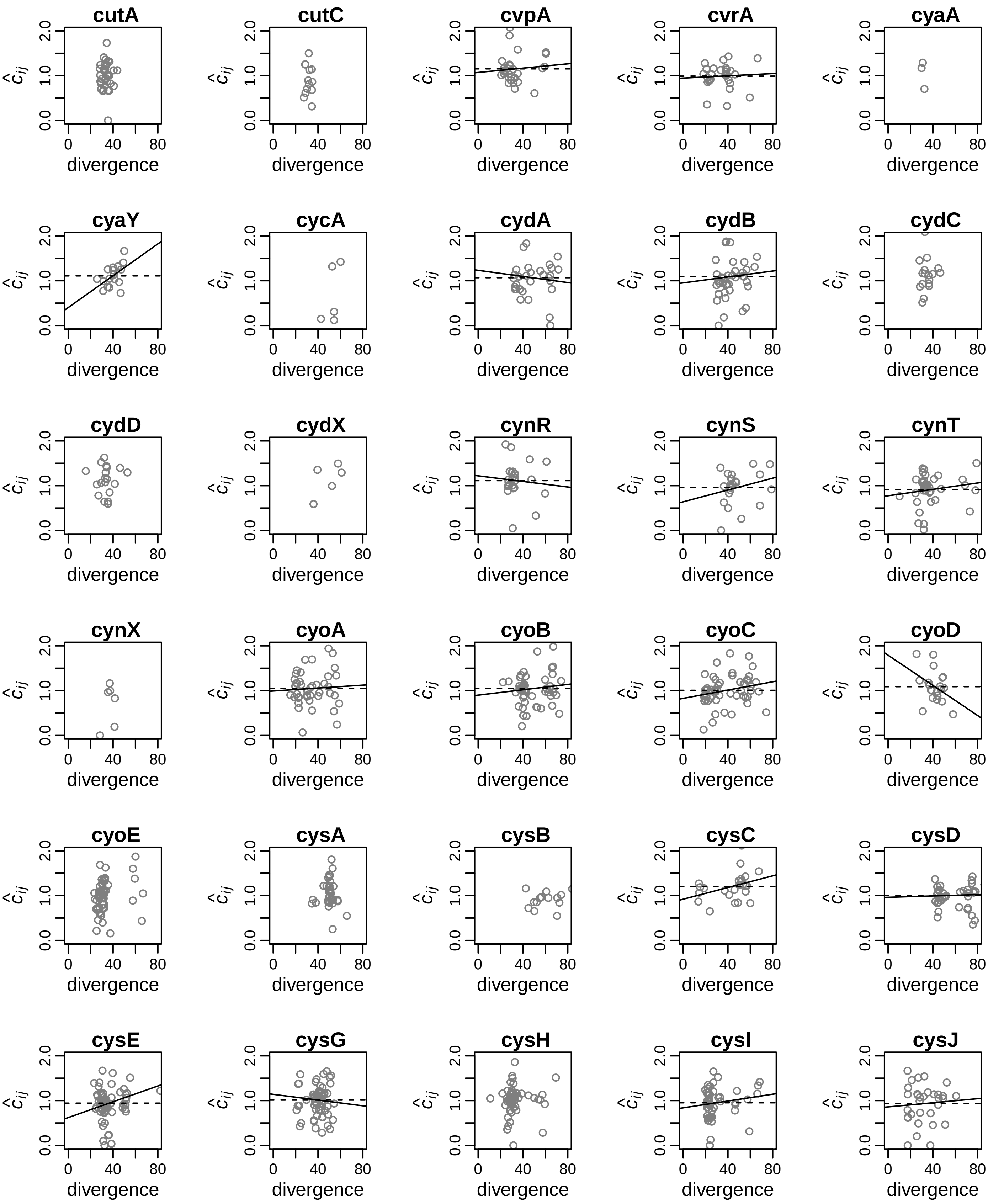
cutA
cutC
cvpA
cvrA
cyaA
cyaY
cycA
cydA
cydB
cydC
cydD
cydX
cynR
cynS
cynT
cynX
cyoA
cyoB
cyoC
cyoD
cyoE
cysA
cysB
cysC
cysD
cysE
cysG
cysH
cysI
cysJ
$\hat{c}_{ij}$
0.0
1.0
2.0
0
40
80
divergence

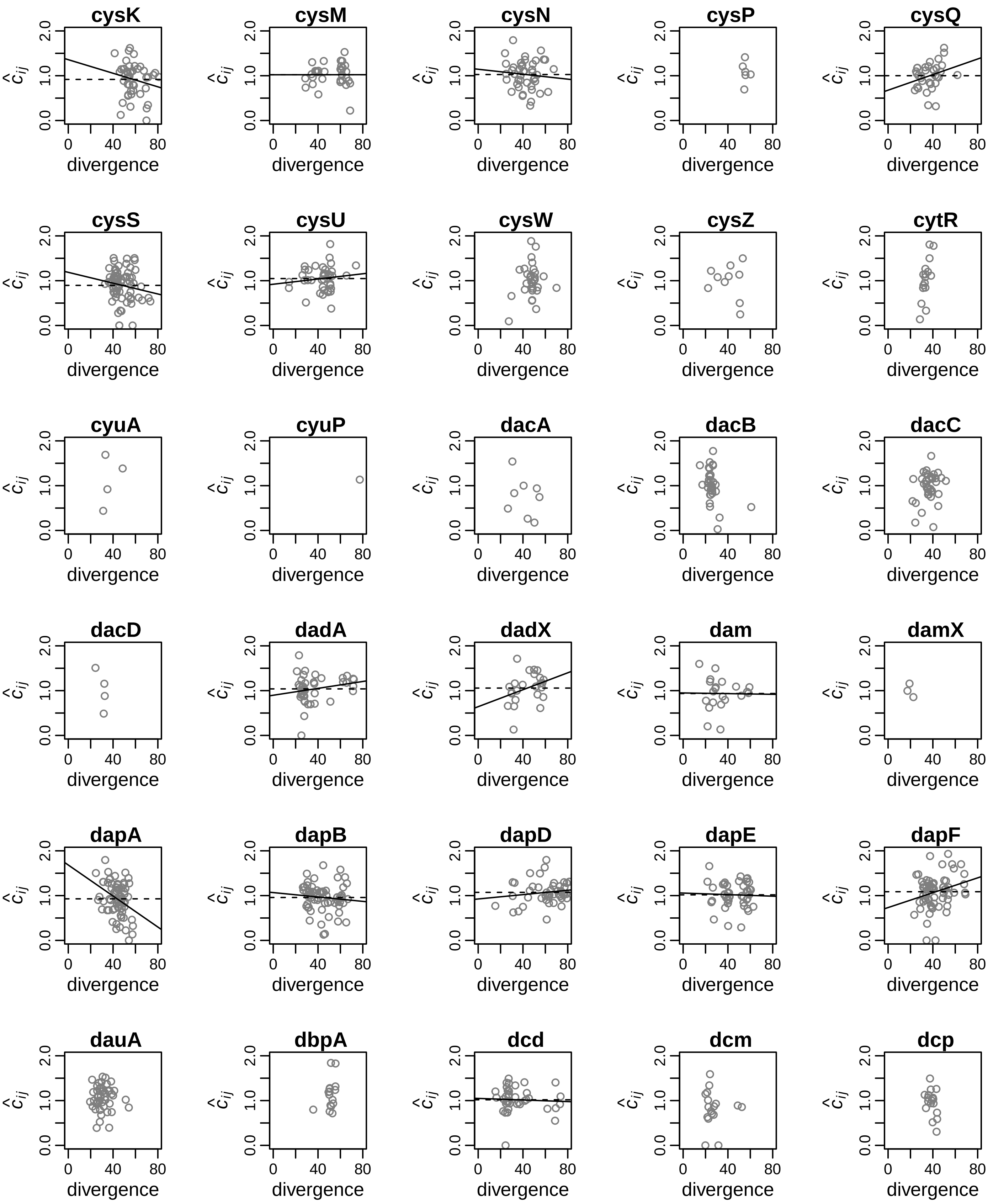
cysK
cysM
cysN
cysP
cysQ
cysS
cysU
cysW
cysZ
cytR
cyuA
cyuP
dacA
dacB
dacC
dacD
dadA
dadX
dam
damX
dapA
dapB
dapD
dapE
dapF
dauA
dbpA
dcd
dcm
dcp
$\hat{c}_{ij}$
0.0
1.0
2.0
0
40
80
divergence

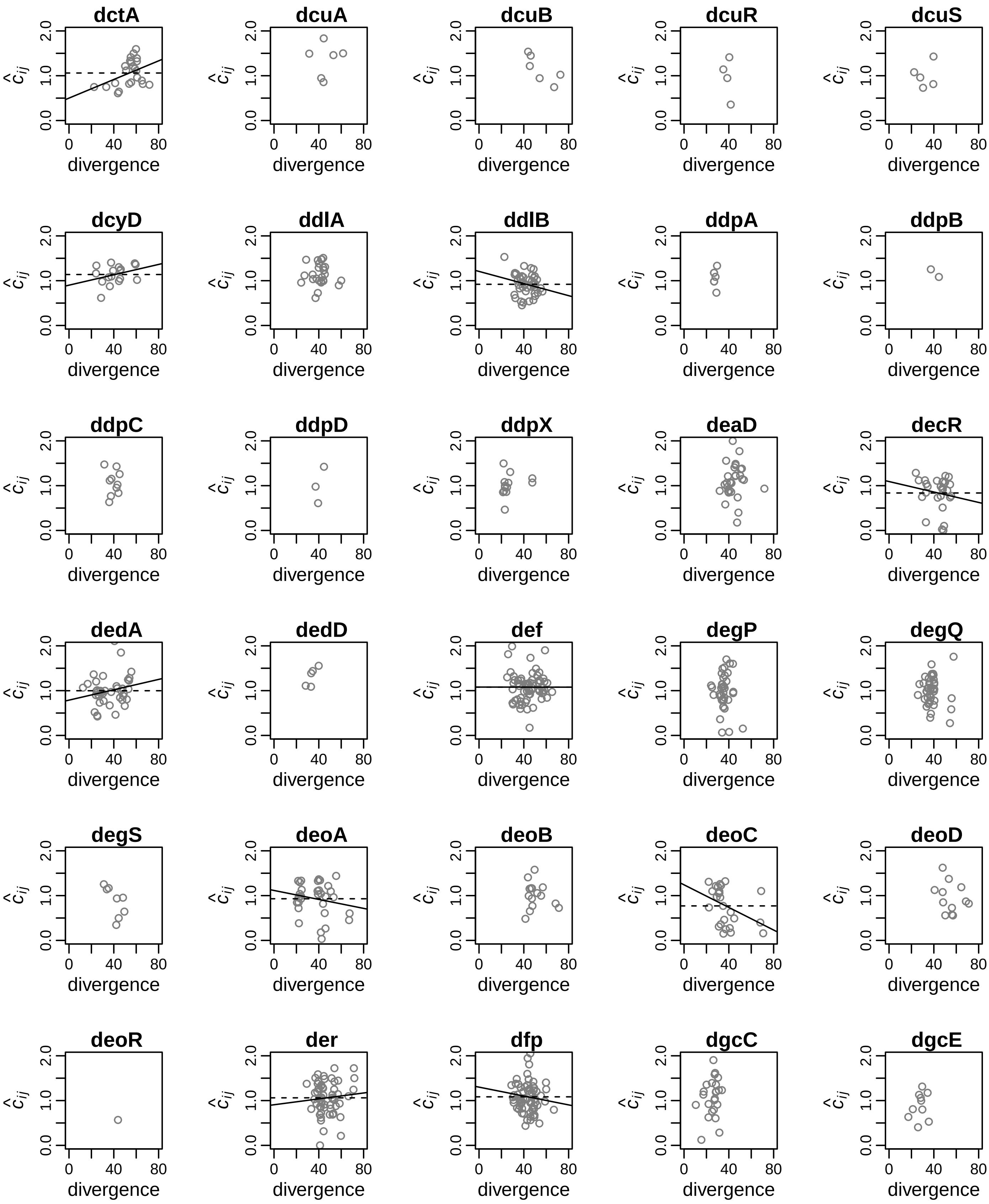
dctA
dcuA
dcuB
dcuR
dcuS
dcyD
ddlA
ddlB
ddpA
ddpB
ddpC
ddpD
ddpX
deaD
decR
dedA
dedD
def
degP
degQ
degS
deoA
deoB
deoC
deoD
deoR
der
dfp
dgcC
dgcE
$\hat{c}_{ij}$
0.0
1.0
2.0
0
40
80
divergence

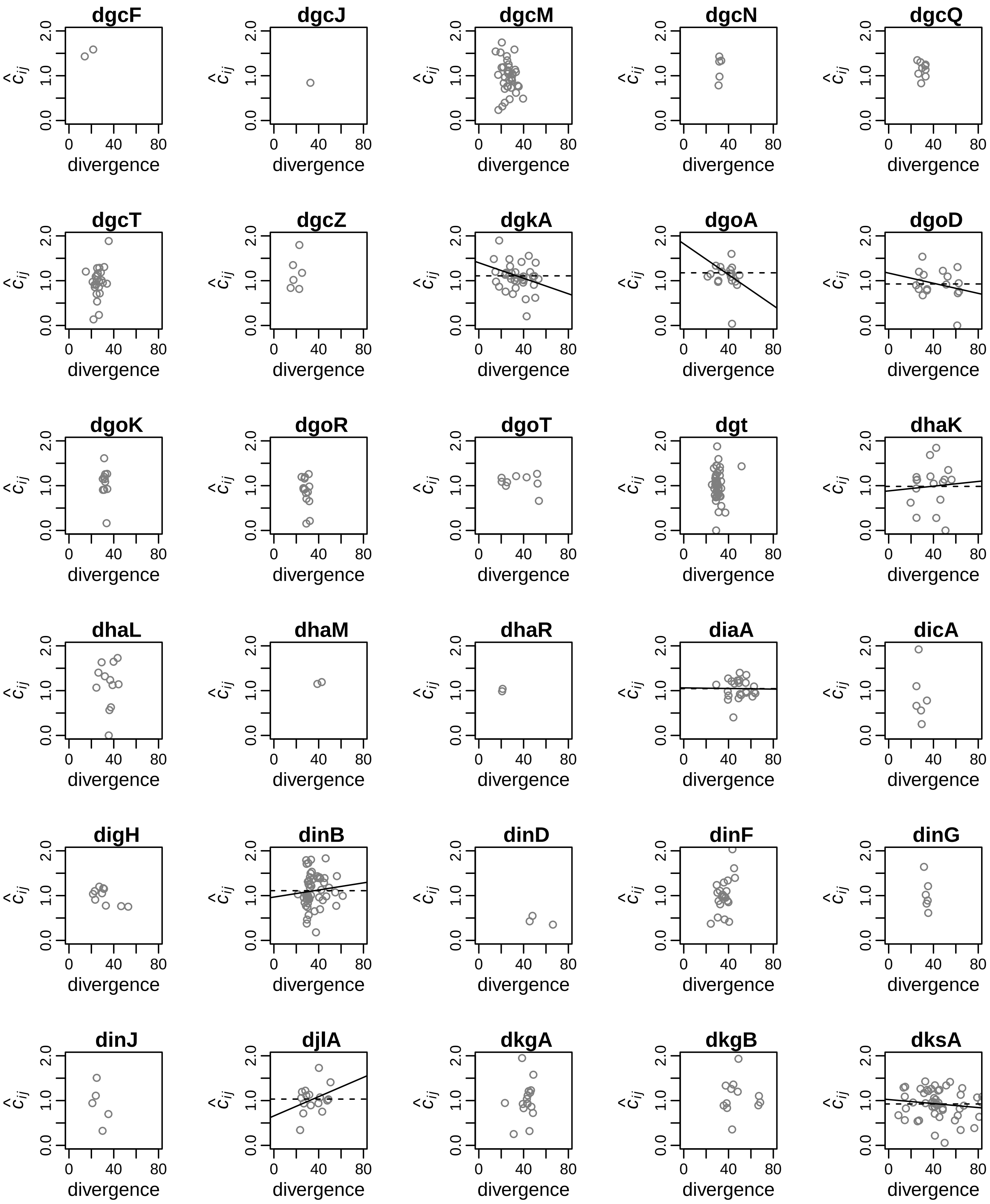

dgcF
dgcJ
dgcM
dgcN
dgcQ
dgcT
dgcZ
dgkA
dgoA
dgoD
dgoK
dgoR
dgoT
dgt
dhaK
dhaL
dhaM
dhaR
diaA
dicA
digH
dinB
dinD
dinF
dinG
dinJ
djlA
dkgA
dkgB
dksA
$\hat{c}_{ij}$
divergence
0.0
1.0
2.0
0
40
80

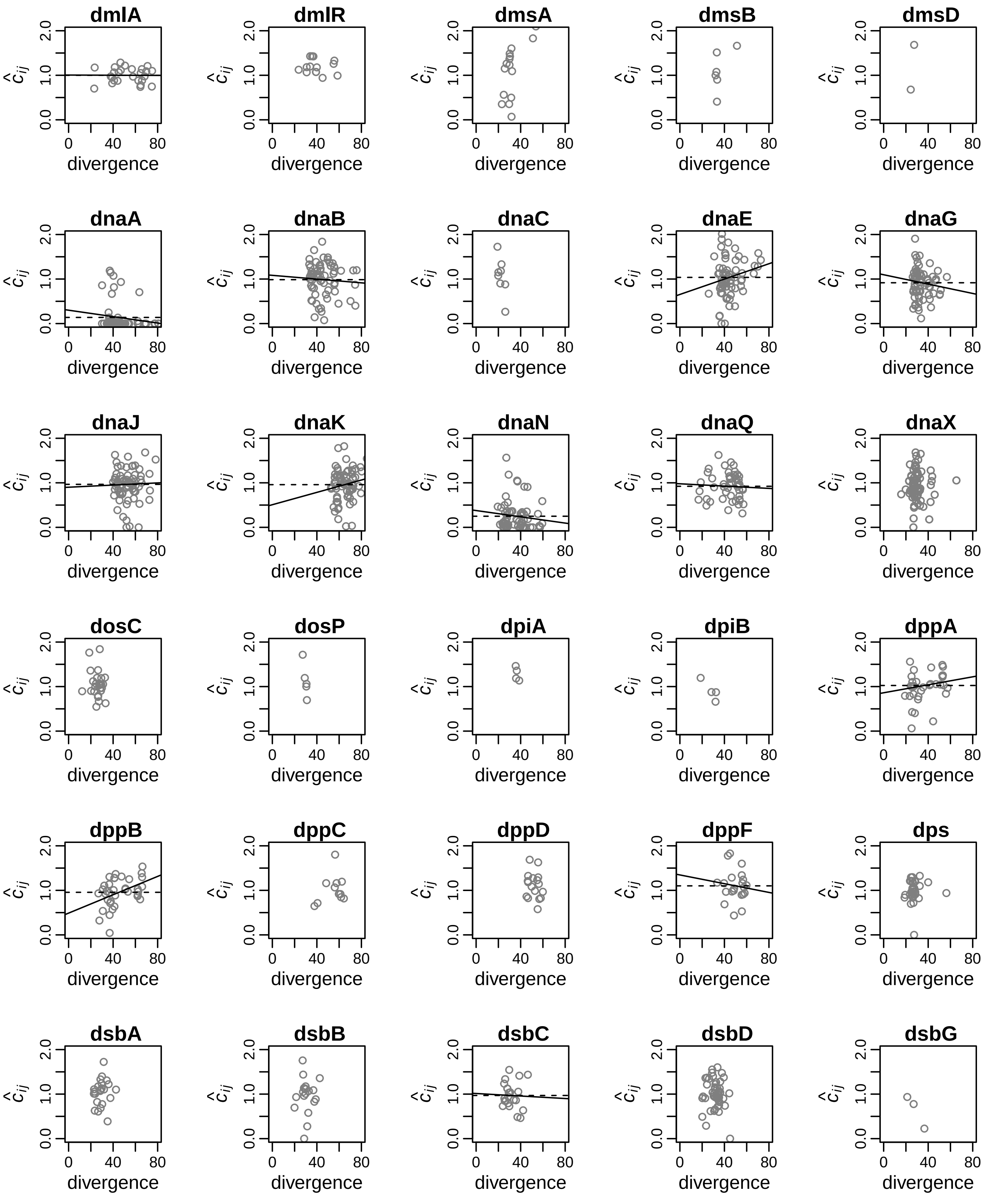
dmlA
dmlR
dmsA
dmsB
dmsD
dnaA
dnaB
dnaC
dnaE
dnaG
dnaJ
dnaK
dnaN
dnaQ
dnaX
dosC
dosP
dpiA
dpiB
dppA
dppB
dppC
dppD
dppF
dps
dsbA
dsbB
dsbC
dsbD
dsbG
$\hat{c}_{ij}$
divergence
0.0
1.0
2.0
0
40
80

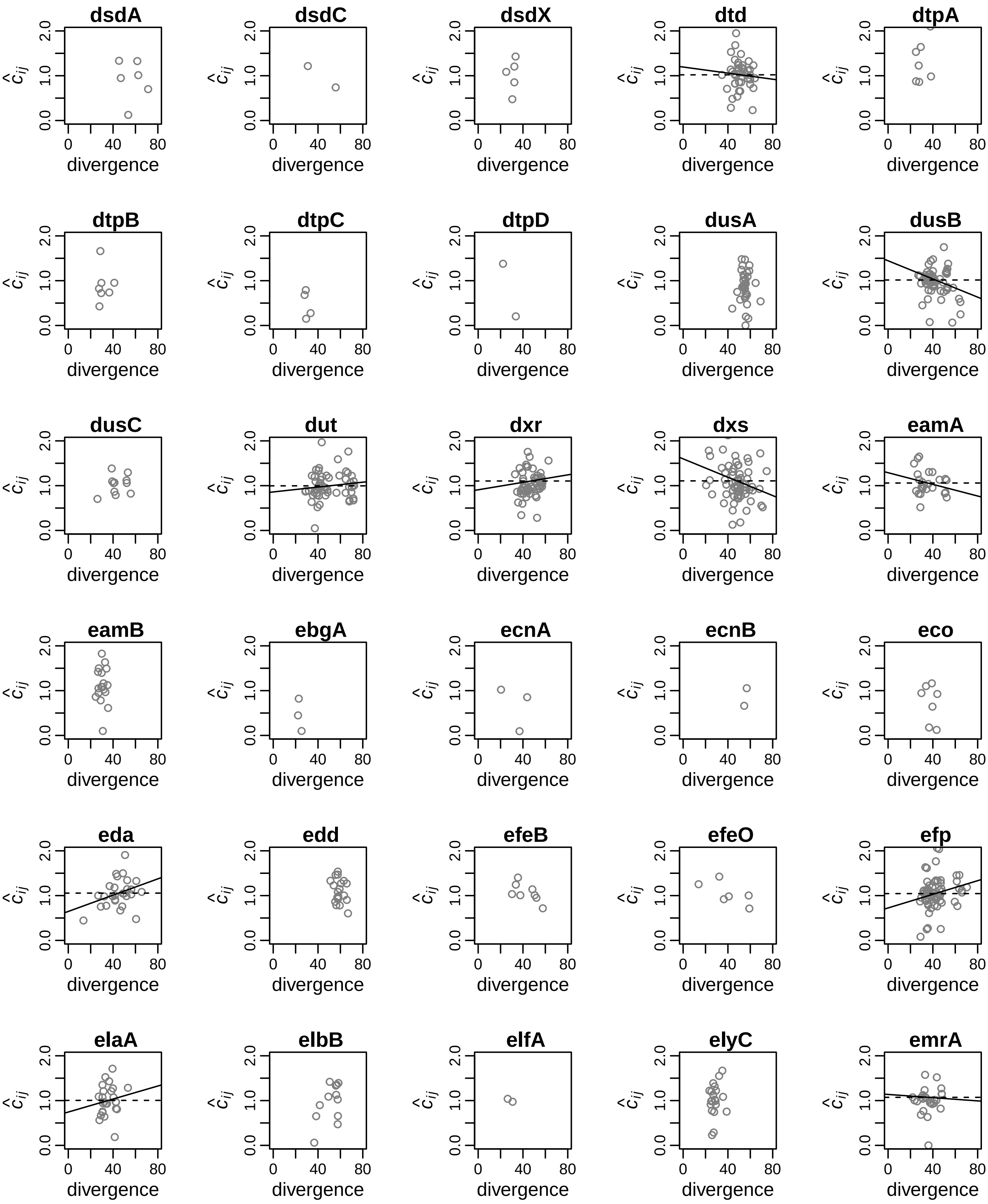
dsdA
dsdC
dsdX
dtd
dtpA
dtpB
dtpC
dtpD
dusA
dusB
dusC
dut
dxr
dxs
eamA
eamB
ebgA
ecnA
ecnB
eco
eda
edd
efeB
efeO
efp
elaA
elbB
elfA
elyC
emrA
$\hat{c}_{ij}$
0.0
1.0
2.0
0
40
80
divergence

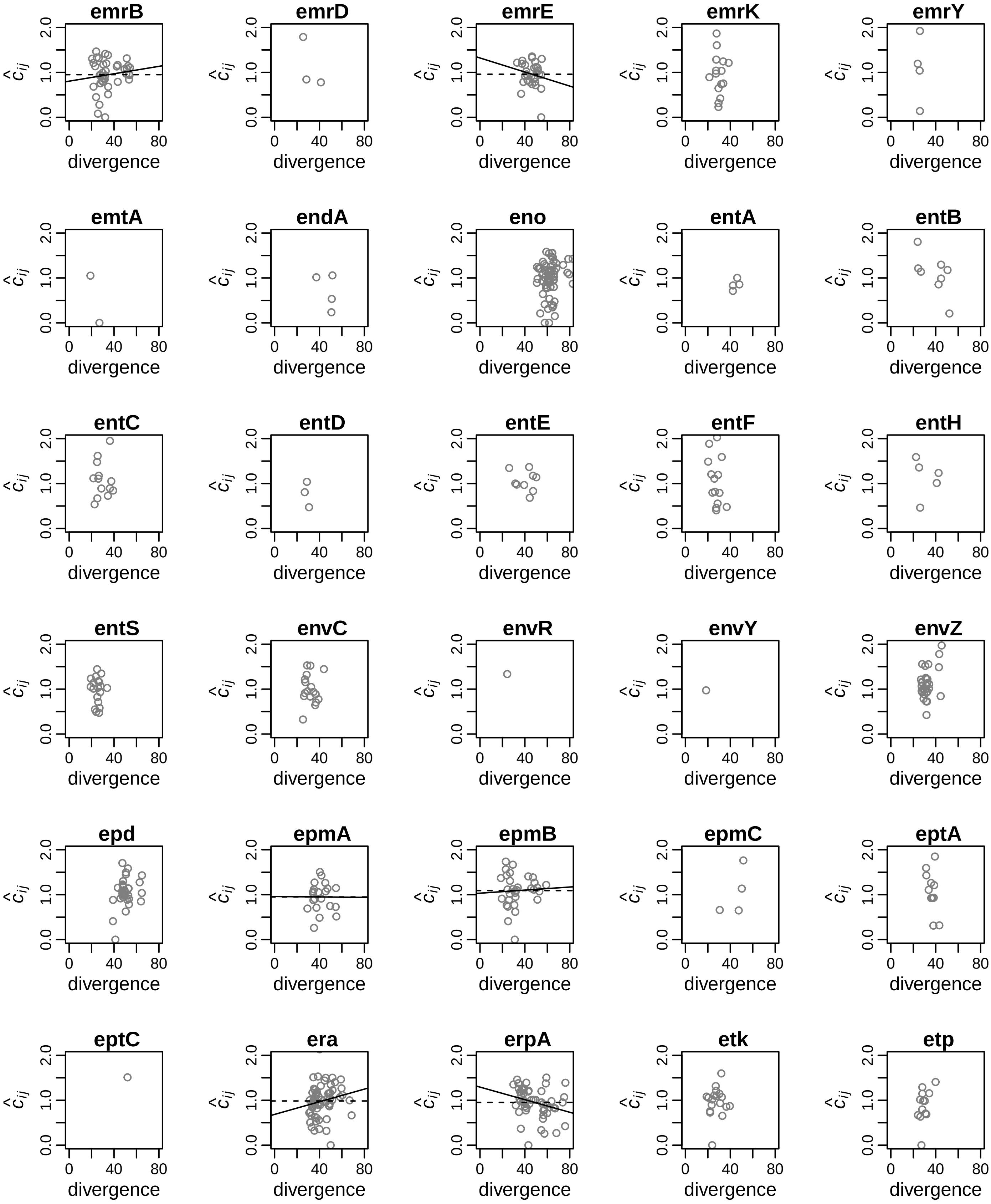
emrB
emrD
emrE
emrK
emrY
emtA
endA
eno
entA
entB
entC
entD
entE
entF
entH
entS
envC
envR
envY
envZ
epd
epmA
epmB
epmC
eptA
eptC
era
erpA
etk
etp
$\hat{c}_{ij}$
0.0
1.0
2.0
0
40
80
divergence

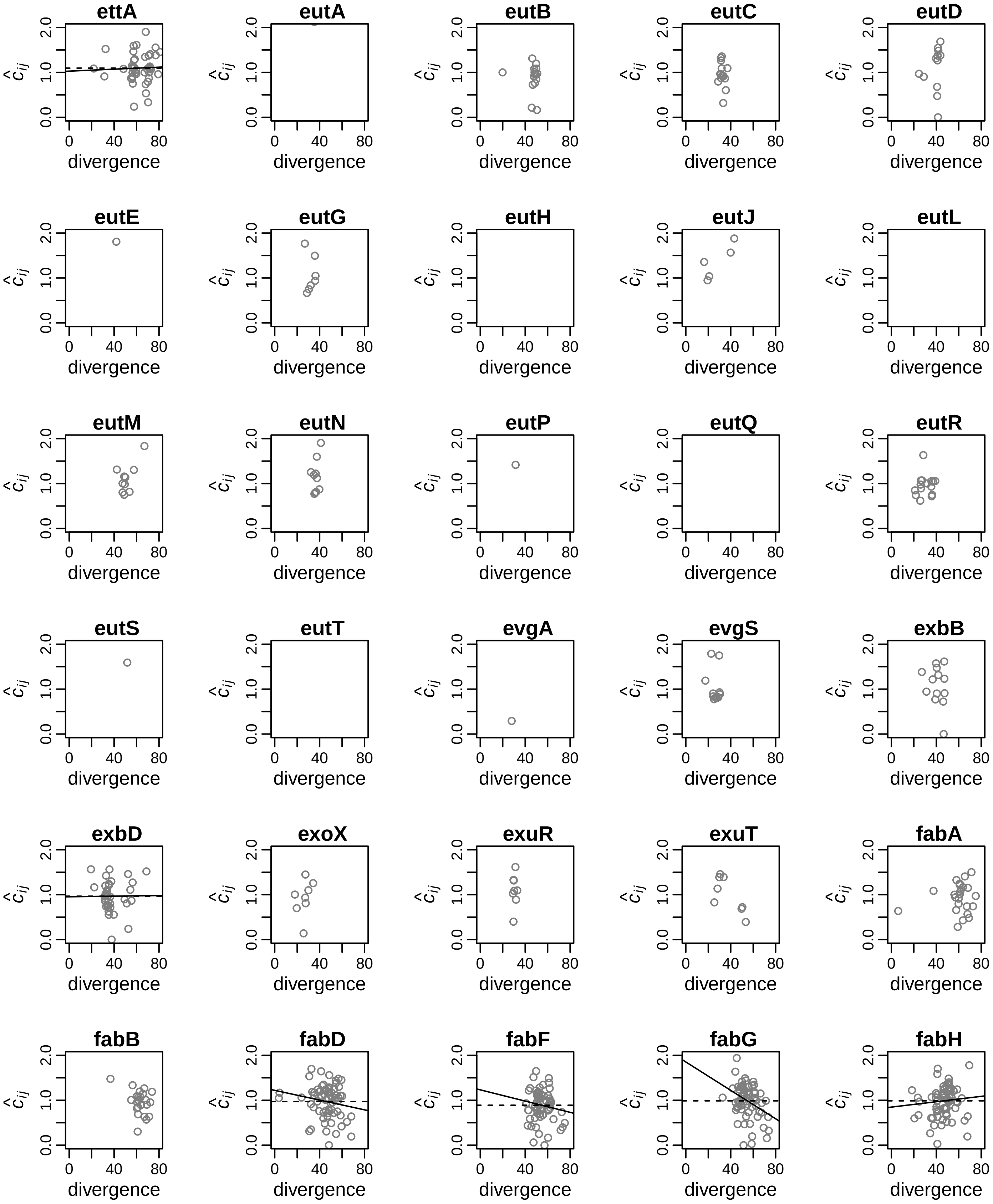
ettA
eutA
eutB
eutC
eutD
eutE
eutG
eutH
eutJ
eutL
eutM
eutN
eutP
eutQ
eutR
eutS
eutT
evgA
evgS
exbB
exbD
exoX
exuR
exuT
fabA
fabB
fabD
fabF
fabG
fabH
$\hat{c}_{ij}$
0.0 1.0 2.0
0 40 80
divergence

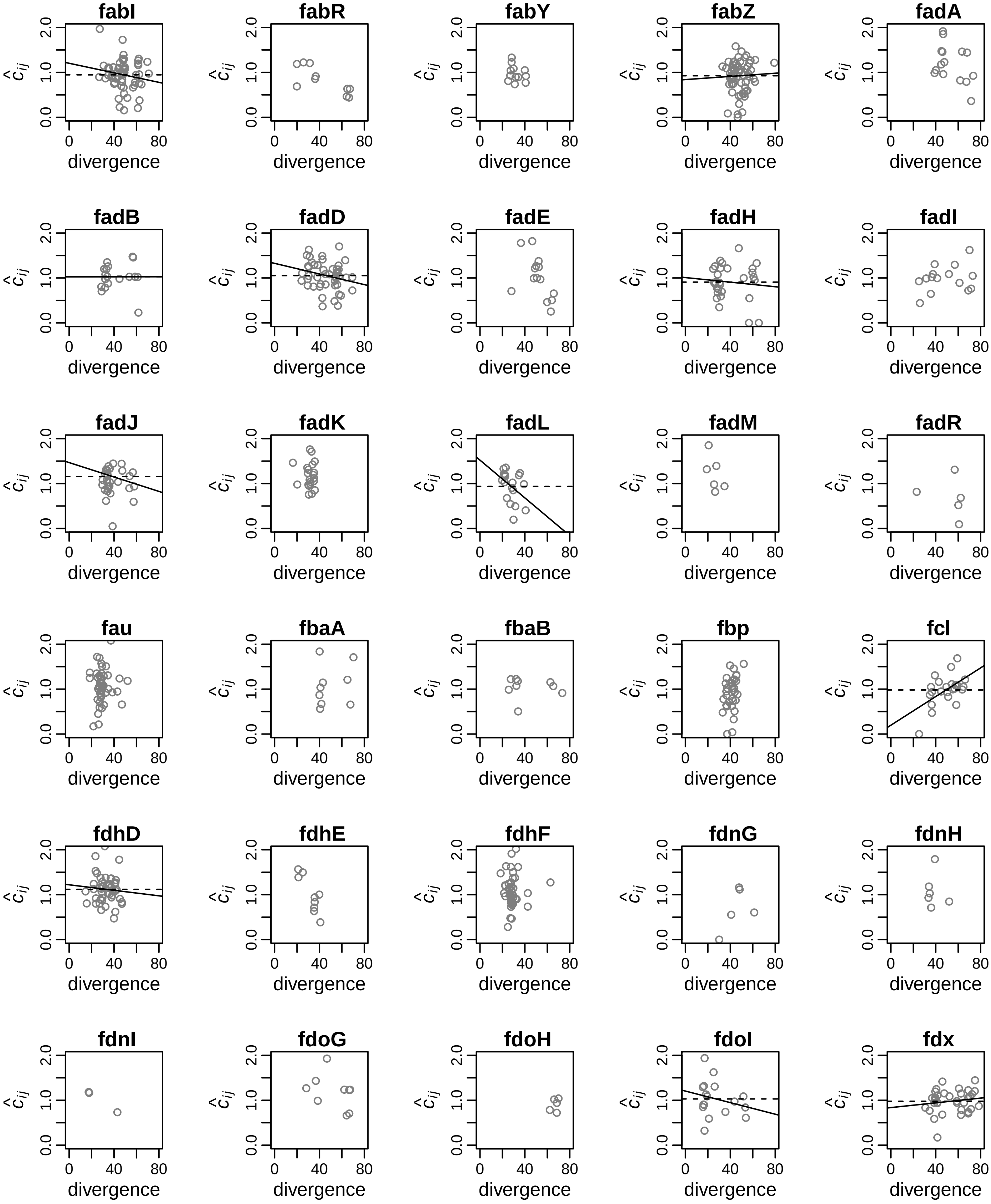
fabI
fabR
fabY
fabZ
fadA
fadB
fadD
fadE
fadH
fadI
fadJ
fadK
fadL
fadM
fadR
fau
fbaA
fbaB
fbp
fcl
fdhD
fdhE
fdhF
fdnG
fdnH
fdnI
fdoG
fdoH
fdoI
fdx
divergence
$\hat{c}_{ij}$
0.0
1.0
2.0
0
40
80

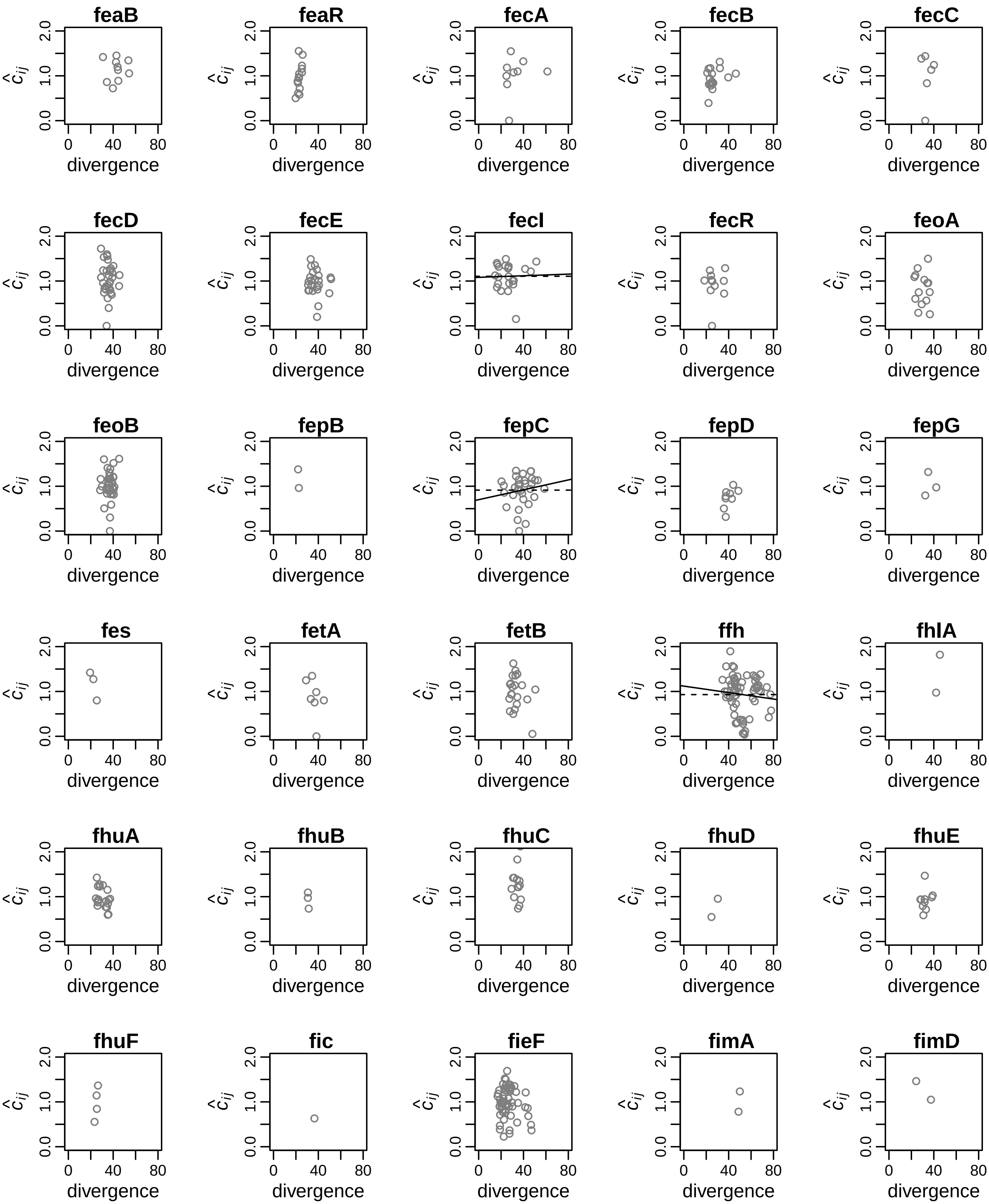
feaB
feaR
fecA
fecB
fecC
fecD
fecE
fecI
fecR
feoA
feoB
fepB
fepC
fepD
fepG
fes
fetA
fetB
ffh
fhlA
fhuA
fhuB
fhuC
fhuD
fhuE
fhuF
fic
fieF
fimA
fimD
$\hat{c}_{ij}$
0.0
1.0
2.0
0
40
80
divergence

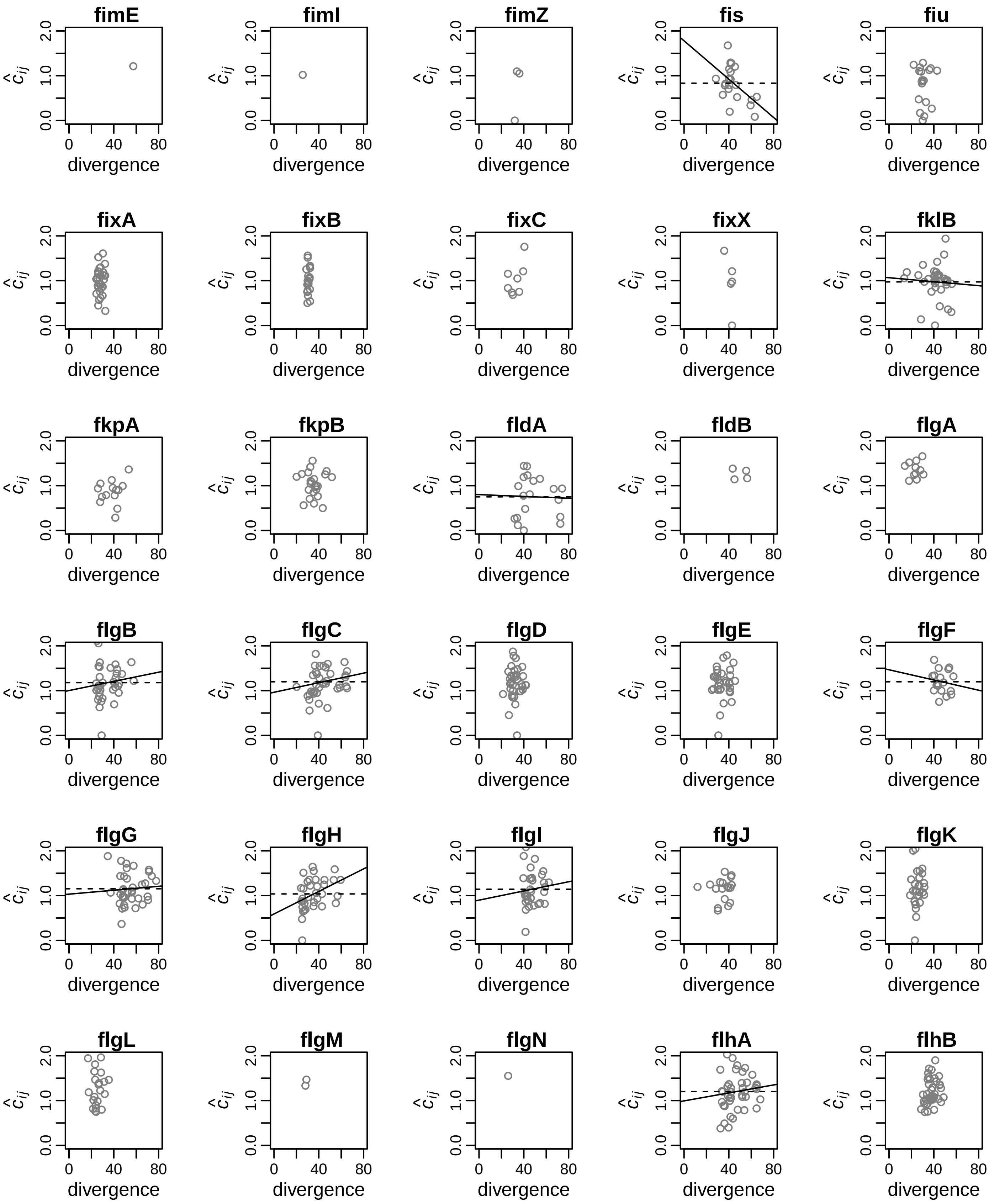

fimE
fimI
fimZ
fis
fiu
fixA
fixB
fixC
fixX
fklB
fkpA
fkpB
fldA
fldB
flgA
flgB
flgC
flgD
flgE
flgF
flgG
flgH
flgI
flgJ
flgK
flgL
flgM
flgN
flhA
flhB
$\hat{c}_{ij}$
0.0
1.0
2.0
0
40
80
divergence

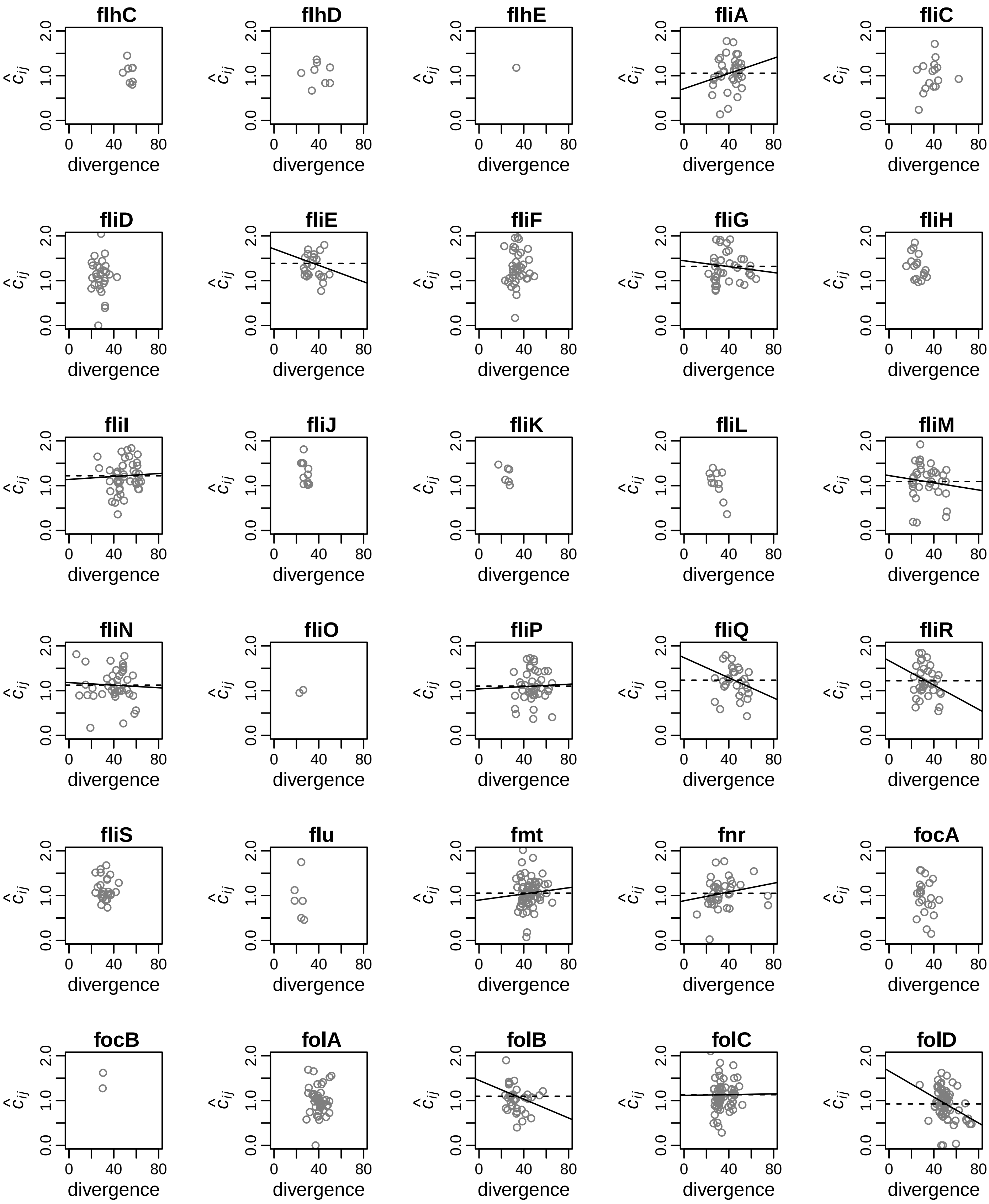
flhC
flhD
flhE
fliA
fliC
fliD
fliE
fliF
fliG
fliH
fliI
fliJ
fliK
fliL
fliM
fliN
fliO
fliP
fliQ
fliR
fliS
flu
fmt
fnr
focA
focB
folA
folB
folC
folD
$\hat{c}_{ij}$
0.0
1.0
2.0
0
40
80
divergence

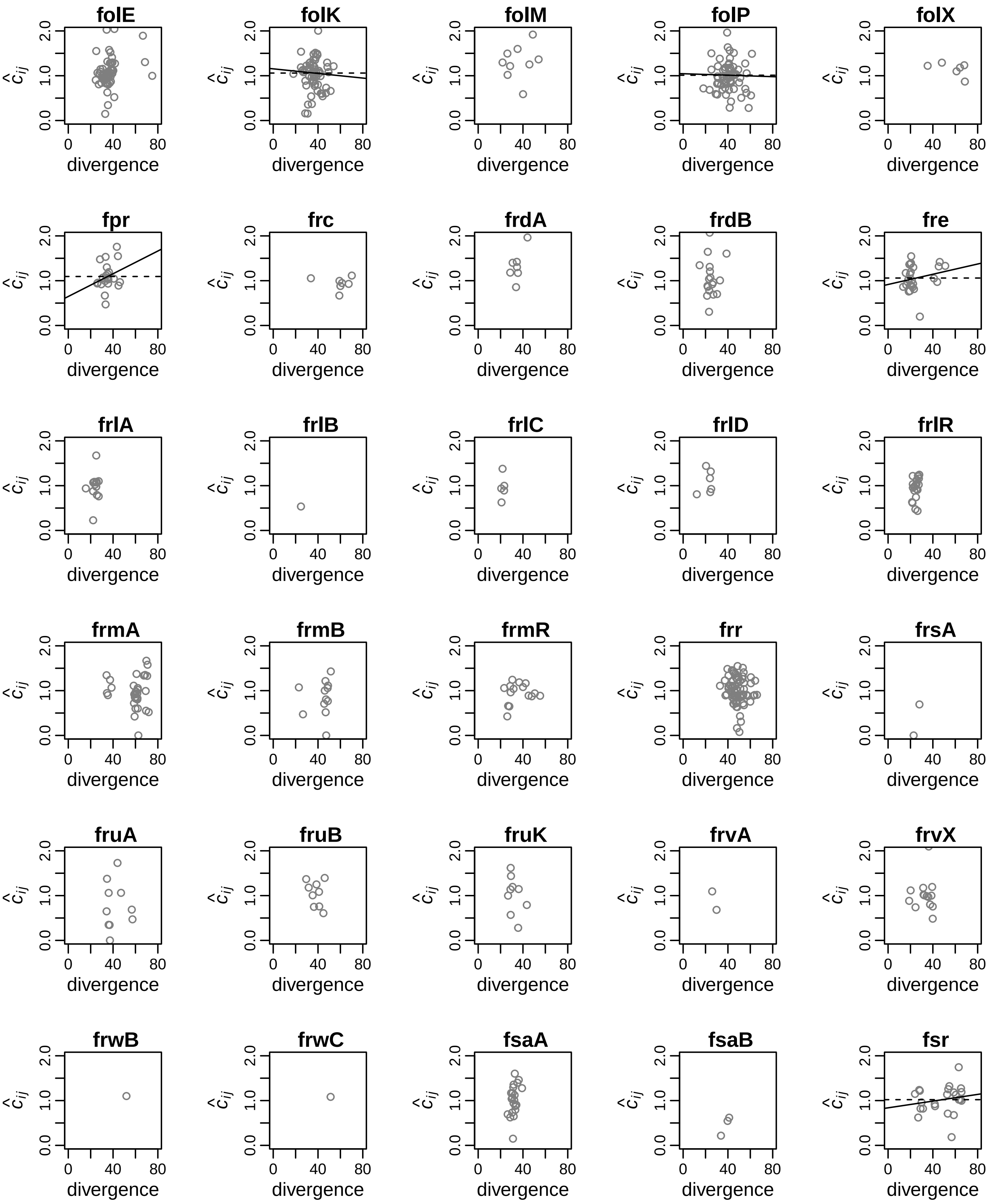
folE
folK
folM
folP
folX
fpr
frc
frdA
frdB
fre
frlA
frlB
frlC
frlD
frlR
frmA
frmB
frmR
frr
frsA
fruA
fruB
fruK
frvA
frvX
frwB
frwC
fsaA
fsaB
fsr
divergence
$\hat{c}_{ij}$

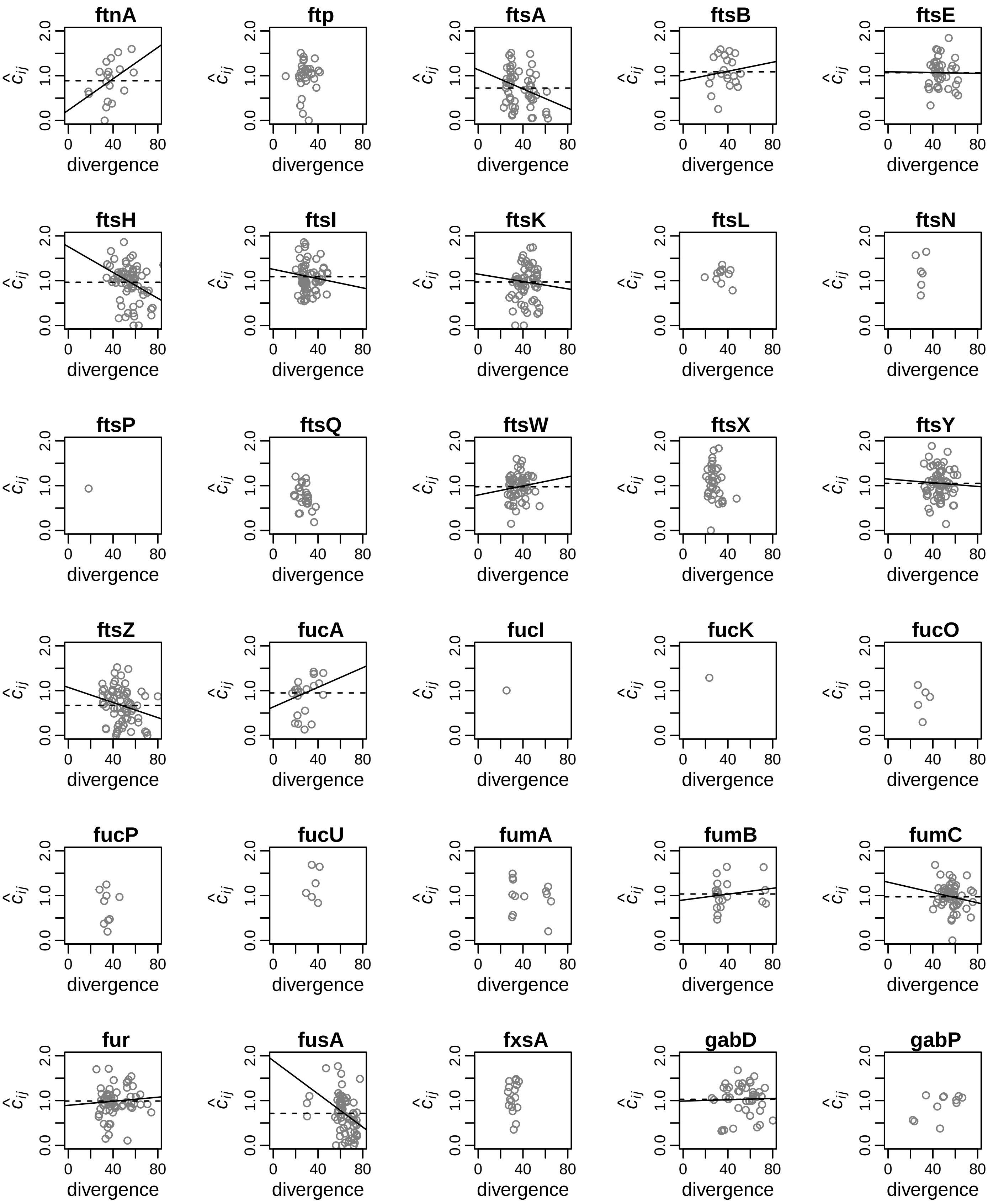
ftnA
ftp
ftsA
ftsB
ftsE
ftsH
ftsI
ftsK
ftsL
ftsN
ftsP
ftsQ
ftsW
ftsX
ftsY
ftsZ
fucA
fucI
fucK
fucO
fucP
fucU
fumA
fumB
fumC
fur
fusA
fxsA
gabD
gabP
$\hat{c}_{ij}$
0.0
1.0
2.0
divergence
0
40
80

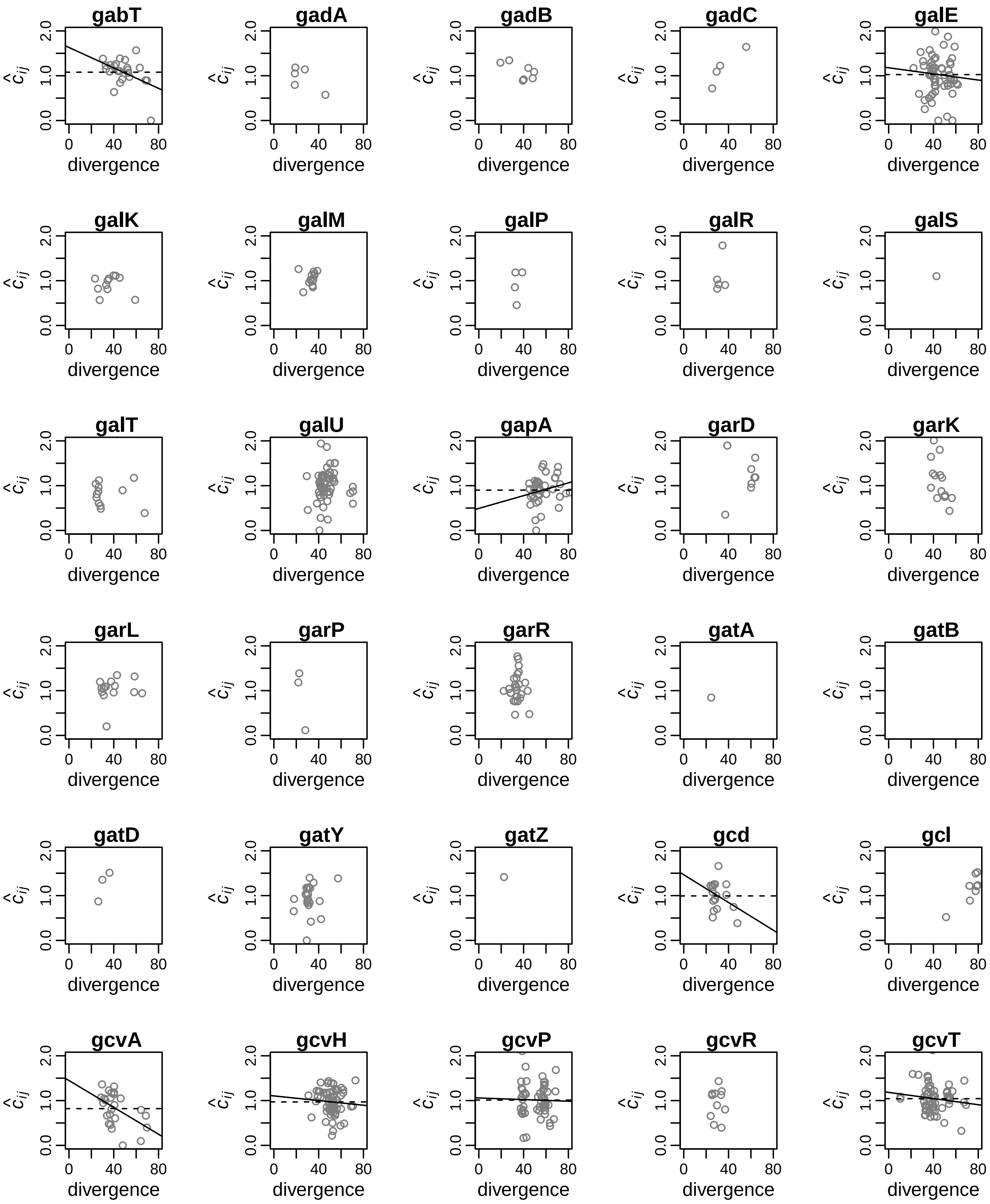
gabT
gadA
gadB
gadC
galE
galK
galM
galP
galR
galS
galT
galU
gapA
garD
garK
garL
garP
garR
gatA
gatB
gatD
gatY
gatZ
gcd
gcl
gcvA
gcvH
gcvP
gcvR
gcvT
$\hat{c}_{ij}$
0.0
1.0
2.0
0
40
80
divergence

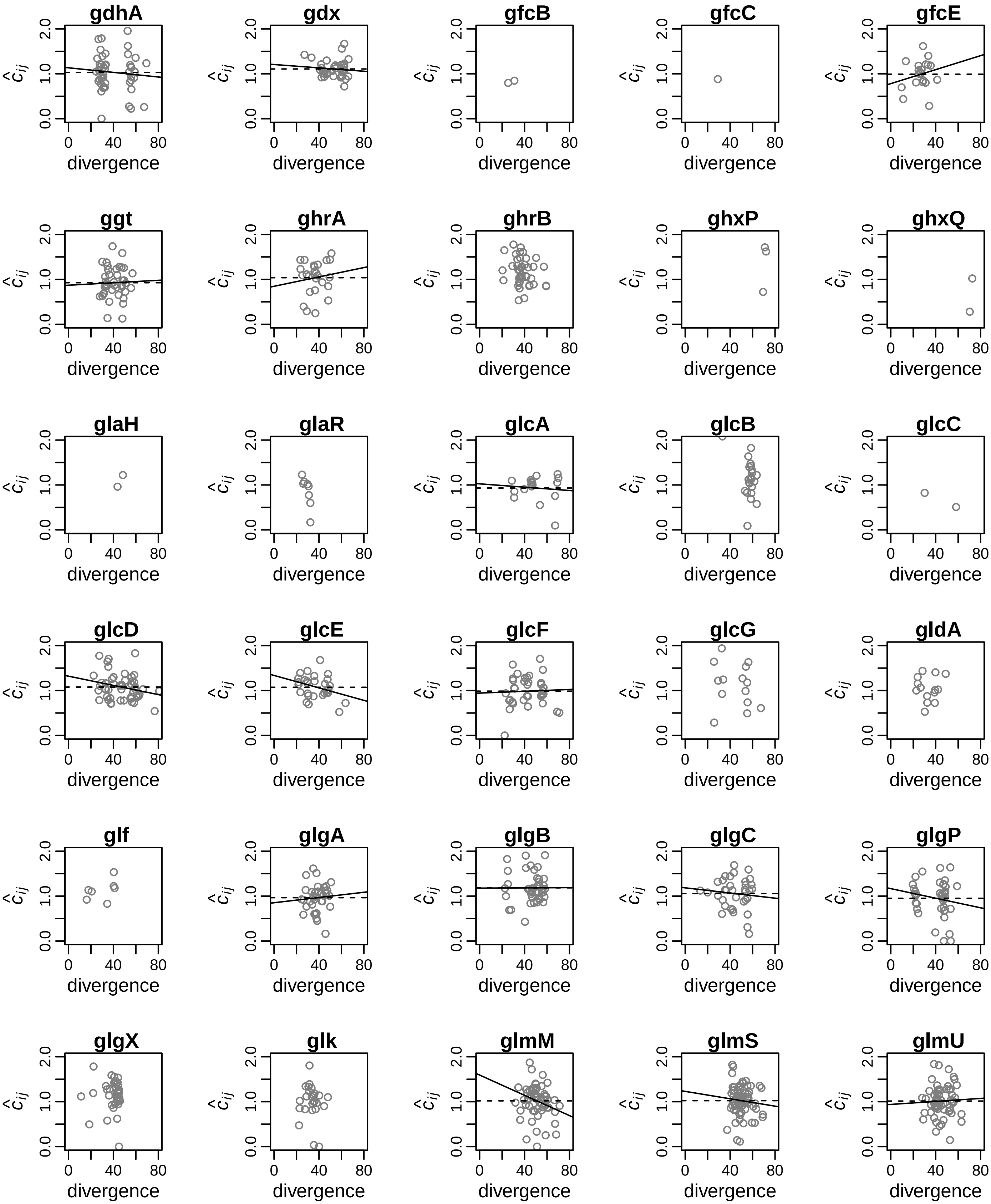
gdhA
gdx
gfcB
gfcC
gfcE
ggt
ghrA
ghrB
ghxP
ghxQ
glaH
glaR
glcA
glcB
glcC
glcD
glcE
glcF
glcG
gldA
glf
glgA
glgB
glgC
glgP
glgX
glk
glmM
glmS
glmU
$\hat{c}_{ij}$
0.0
1.0
2.0
0
40
80
divergence

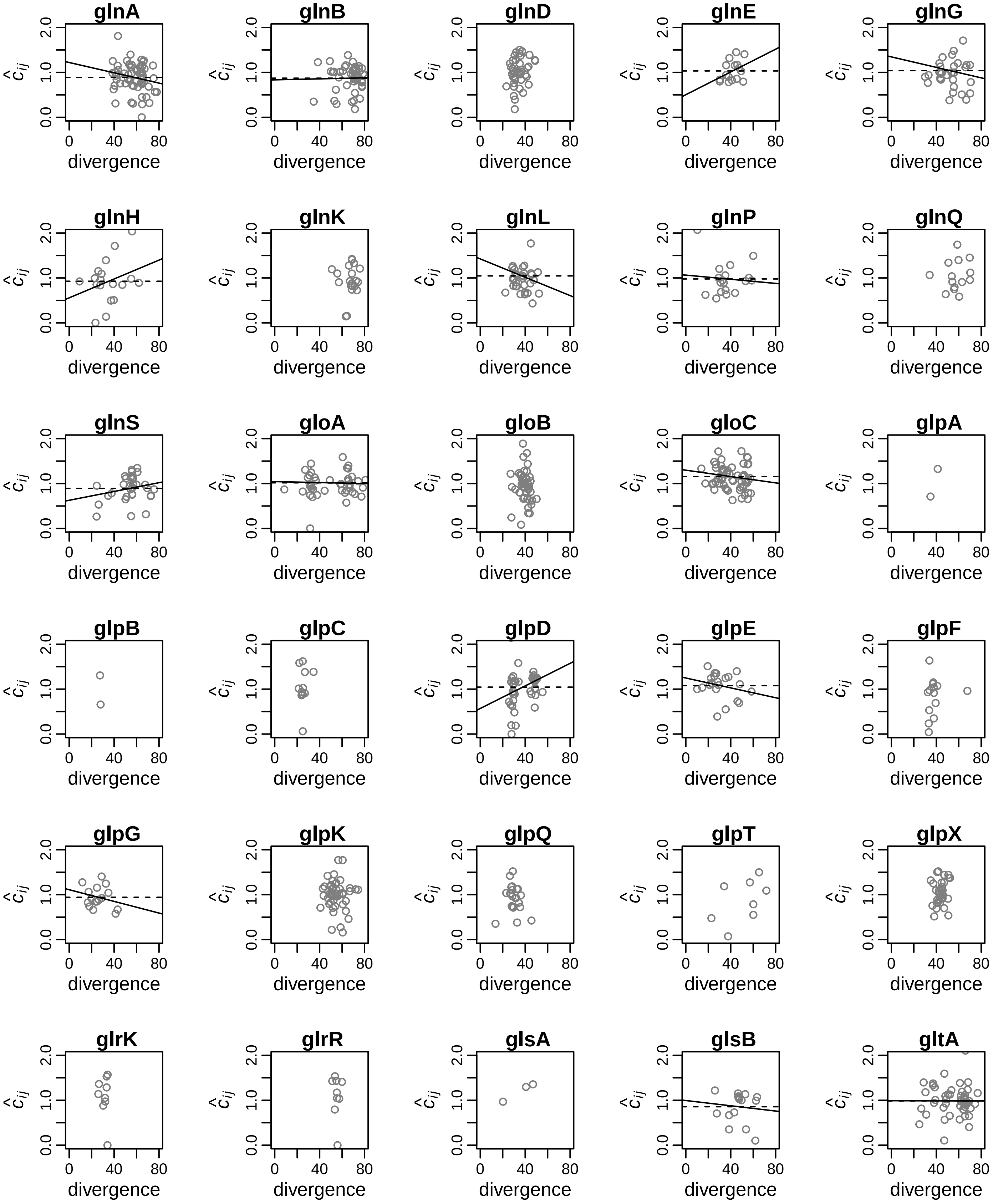
glnA
glnB
glnD
glnE
glnG
glnH
glnK
glnL
glnP
glnQ
glnS
gloA
gloB
gloC
glpA
glpB
glpC
glpD
glpE
glpF
glpG
glpK
glpQ
glpT
glpX
glrK
glrR
glsA
glsB
gltA
$\hat{c}_{ij}$
0.0 1.0 2.0
0 40 80
divergence

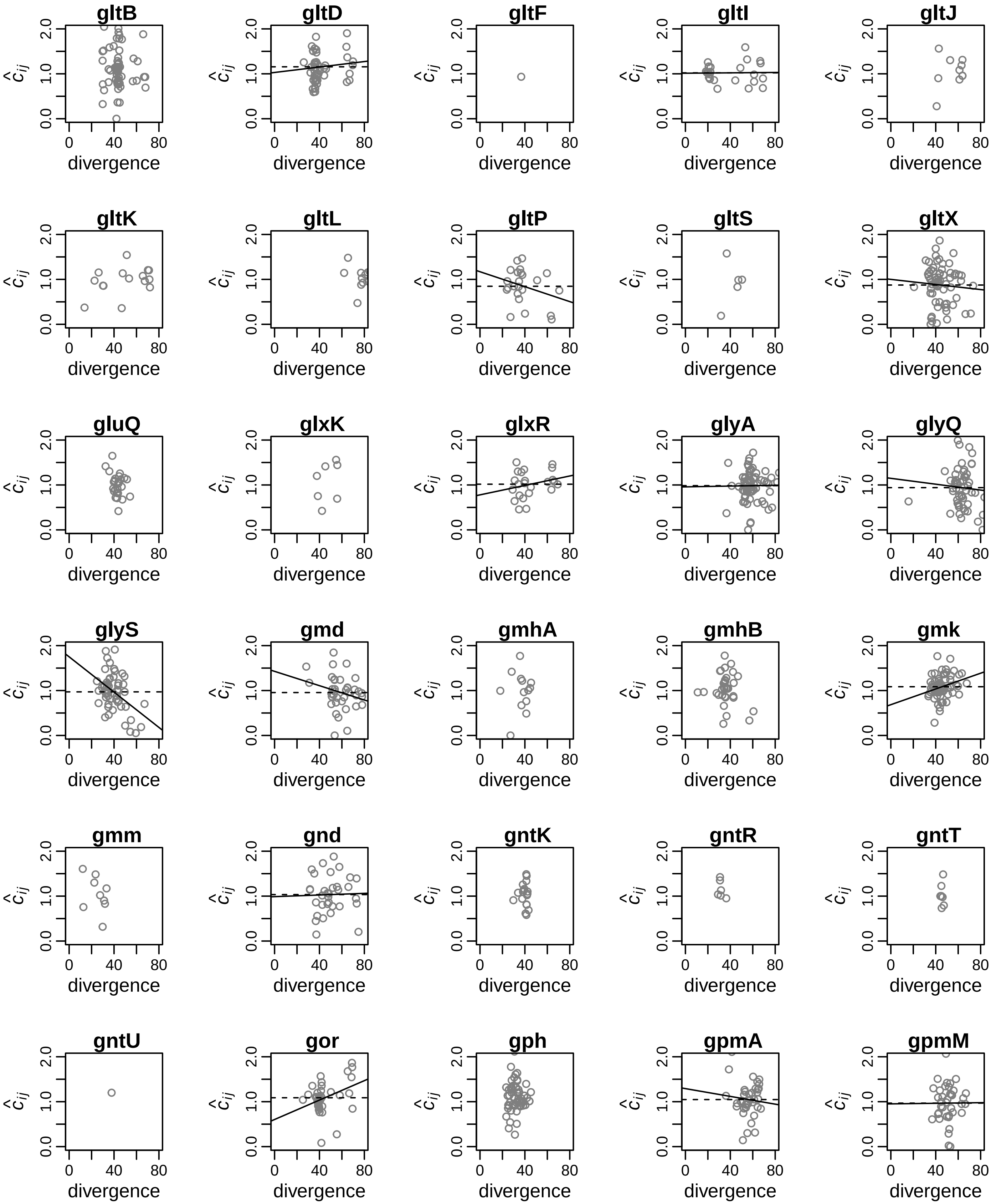
gltB
gltD
gltF
gltI
gltJ
gltK
gltL
gltP
gltS
gltX
gluQ
glxK
glxR
glyA
glyQ
glyS
gmd
gmhA
gmhB
gmk
gmm
gnd
gntK
gntR
gntT
gntU
gor
gph
gpmA
gpmM
$\hat{c}_{ij}$
0.0
1.0
2.0
0
40
80
divergence

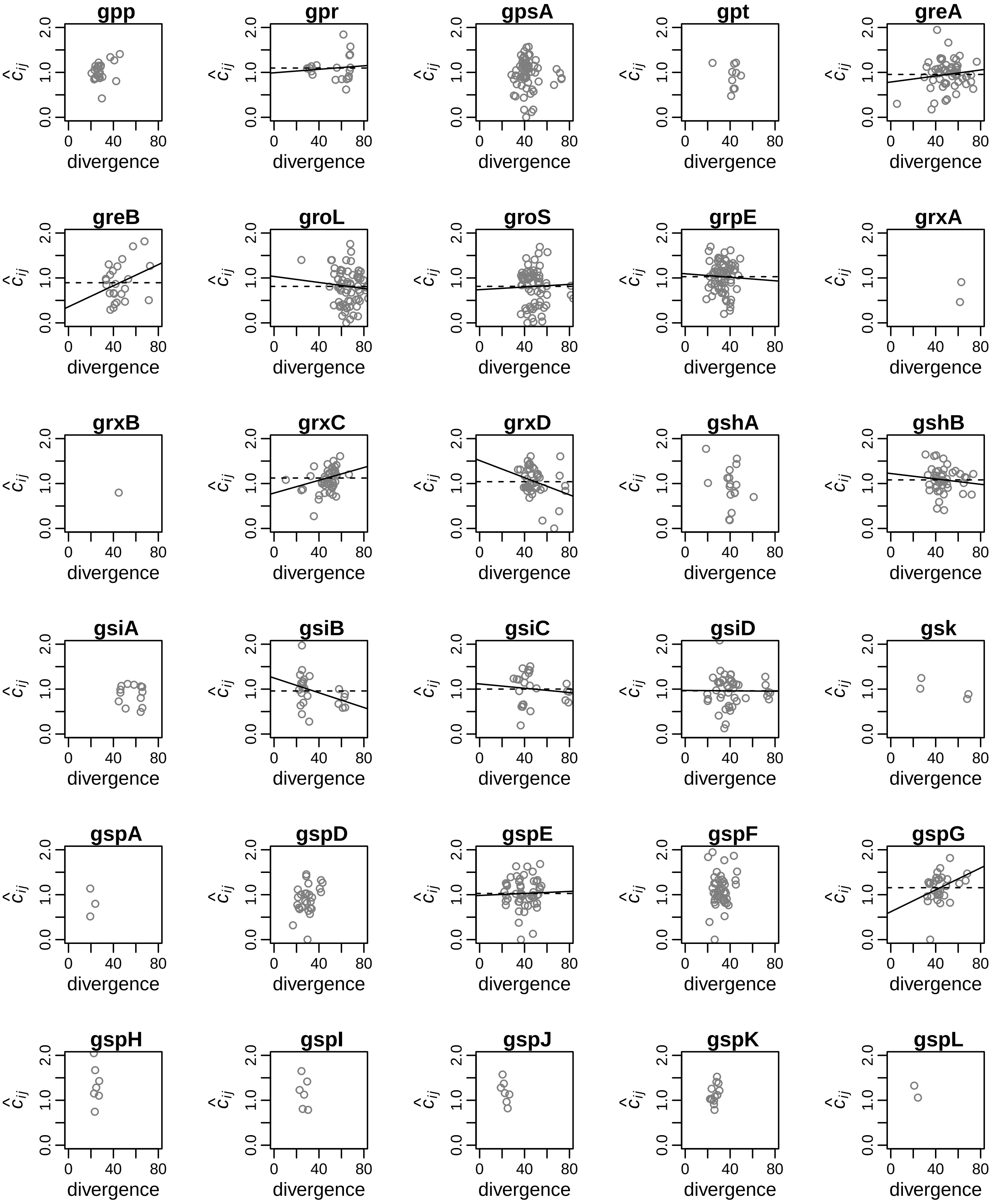
gpp
gpr
gpsA
gpt
greA
greB
groL
groS
grpE
grxA
grxB
grxC
grxD
gshA
gshB
gsiA
gsiB
gsiC
gsiD
gsk
gspA
gspD
gspE
gspF
gspG
gspH
gspI
gspJ
gspK
gspL
divergence
$\hat{c}_{ij}$
0.0
1.0
2.0
0
40
80

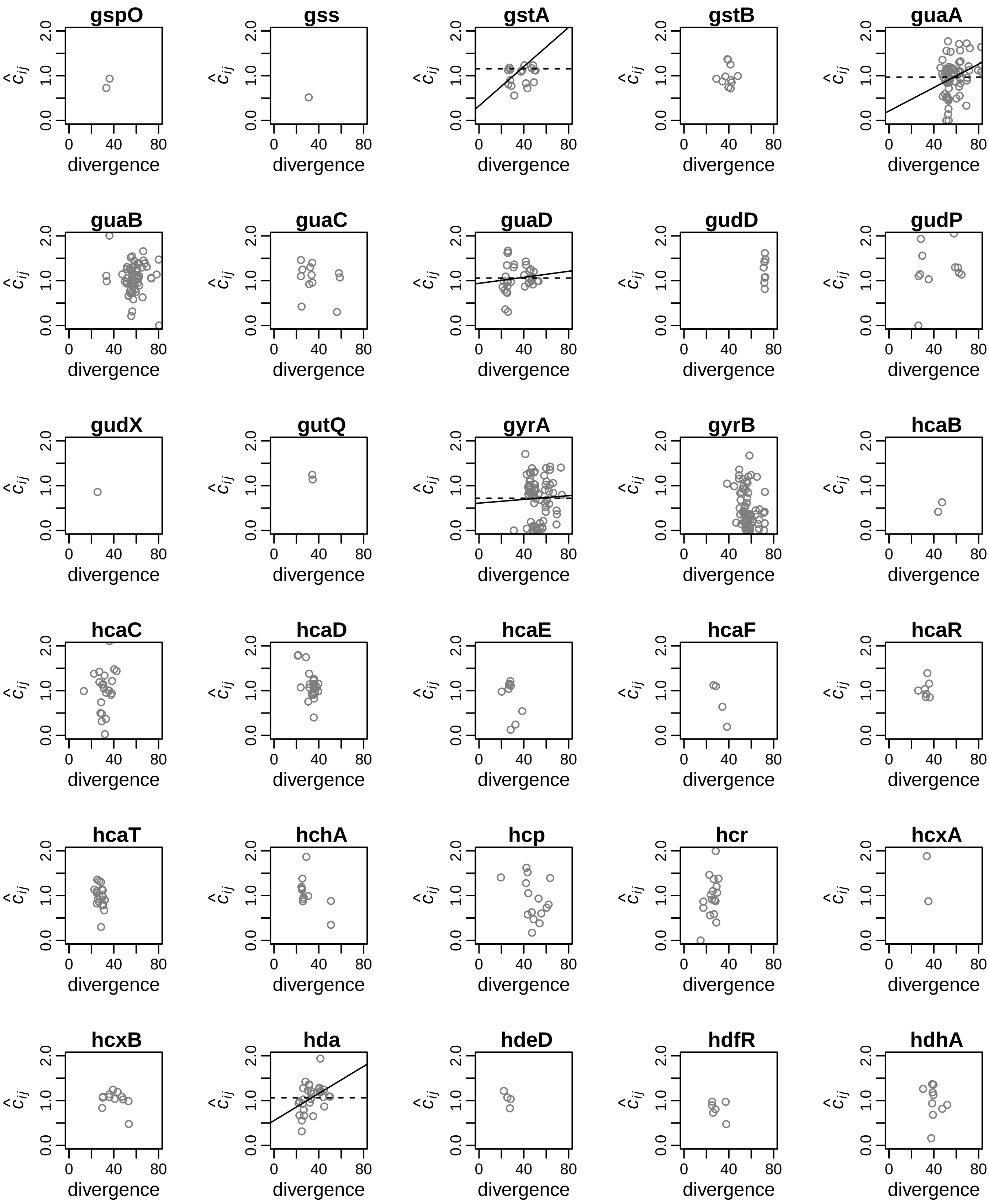
gspO
gss
gstA
gstB
guaA
guaB
guaC
guaD
gudD
gudP
gudX
gutQ
gyrA
gyrB
hcaB
hcaC
hcaD
hcaE
hcaF
hcaR
hcaT
hchA
hcp
hcr
hcxA
hcxB
hda
hdeD
hdfR
hdhA
$\hat{c}_{ij}$
0.0
1.0
2.0
0
40
80
divergence

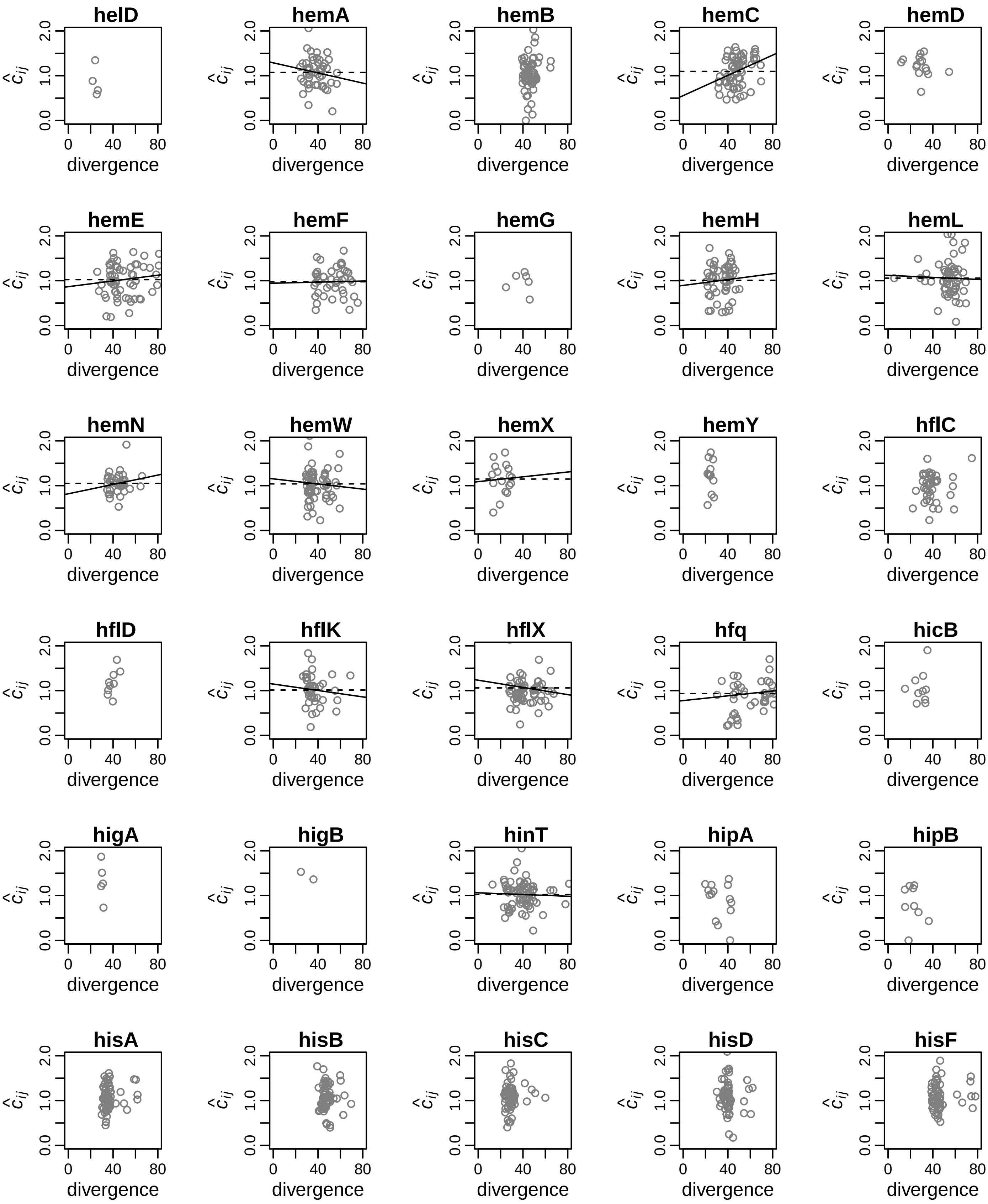
helD
hemA
hemB
hemC
hemD
hemE
hemF
hemG
hemH
hemL
hemN
hemW
hemX
hemY
hflC
hflD
hflK
hflX
hfq
hicB
higA
higB
hinT
hipA
hipB
hisA
hisB
hisC
hisD
hisF
$\hat{c}_{ij}$
0.0
1.0
2.0
0
40
80
divergence

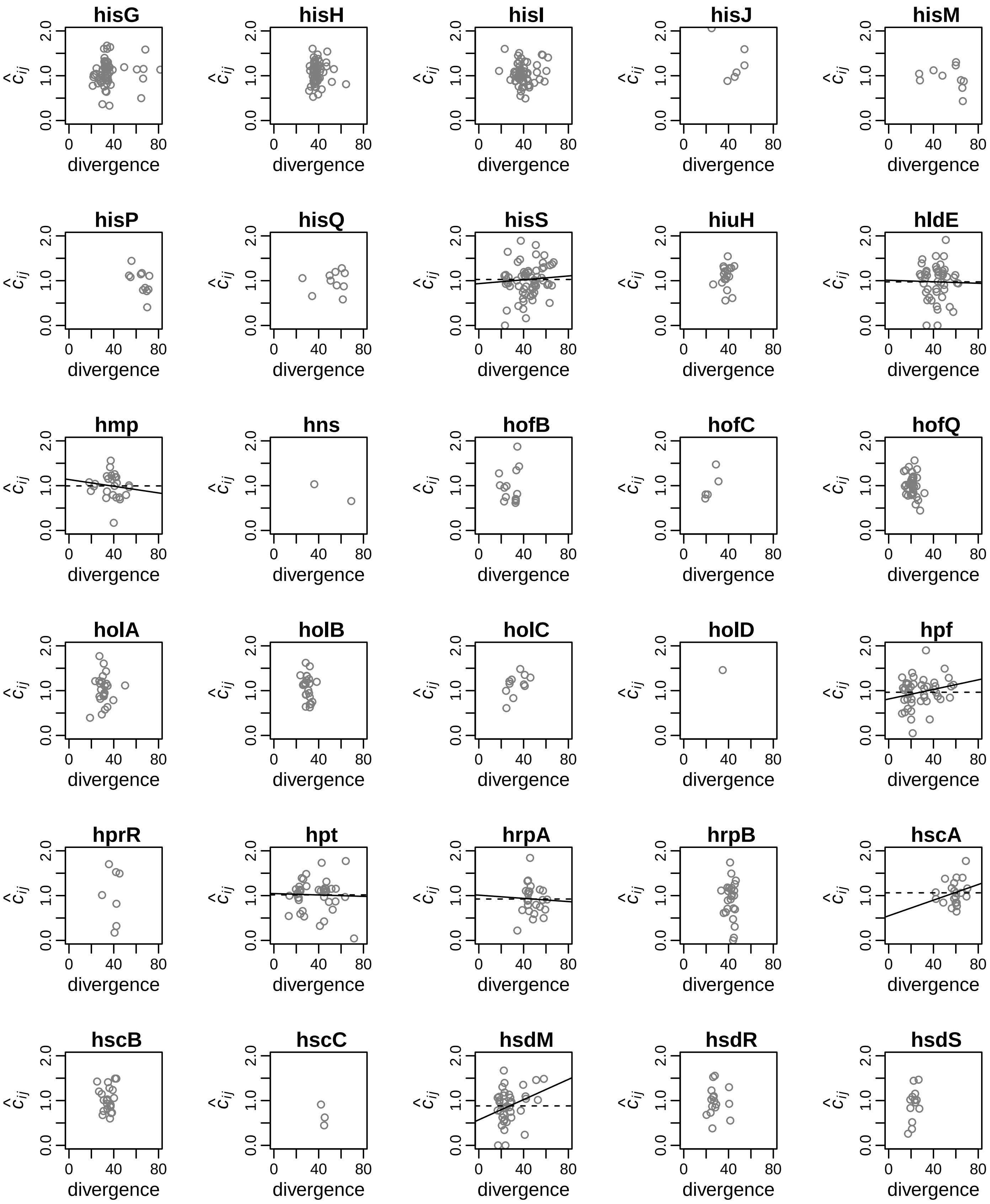
hisG
hisH
hisI
hisJ
hisM
hisP
hisQ
hisS
hiuH
hldE
hmp
hns
hofB
hofC
hofQ
holA
holB
holC
holD
hpf
hprR
hpt
hrpA
hrpB
hscA
hscB
hscC
hsdM
hsdR
hsdS
$\hat{c}_{ij}$
0.0
1.0
2.0
divergence
0
40
80

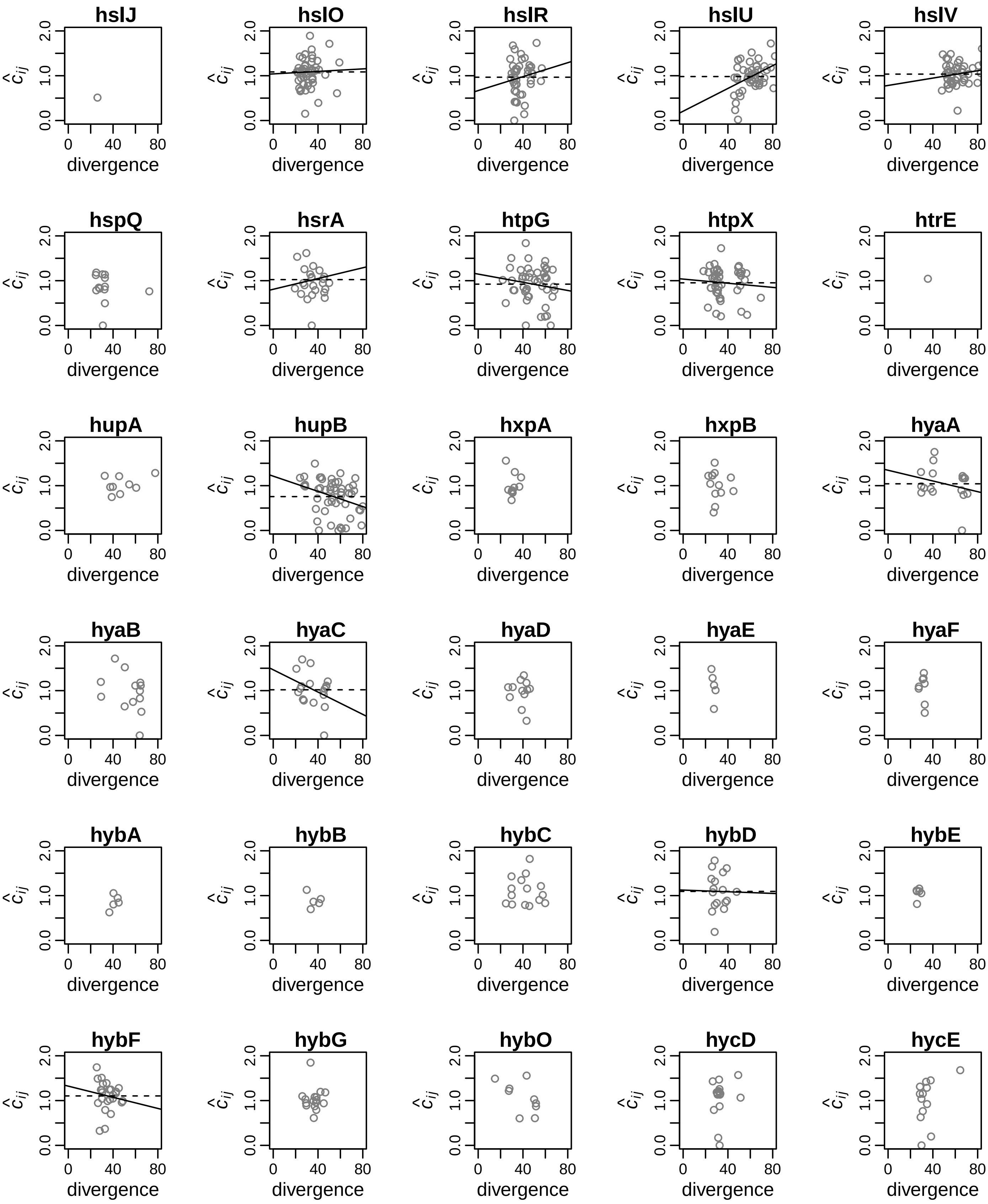
hslJ
hslO
hslR
hslU
hslV
hspQ
hsrA
htpG
htpX
htrE
hupA
hupB
hxpA
hxpB
hyaA
hyaB
hyaC
hyaD
hyaE
hyaF
hybA
hybB
hybC
hybD
hybE
hybF
hybG
hybO
hycD
hycE
$\hat{c}_{ij}$
0.0
1.0
2.0
0
40
80
divergence

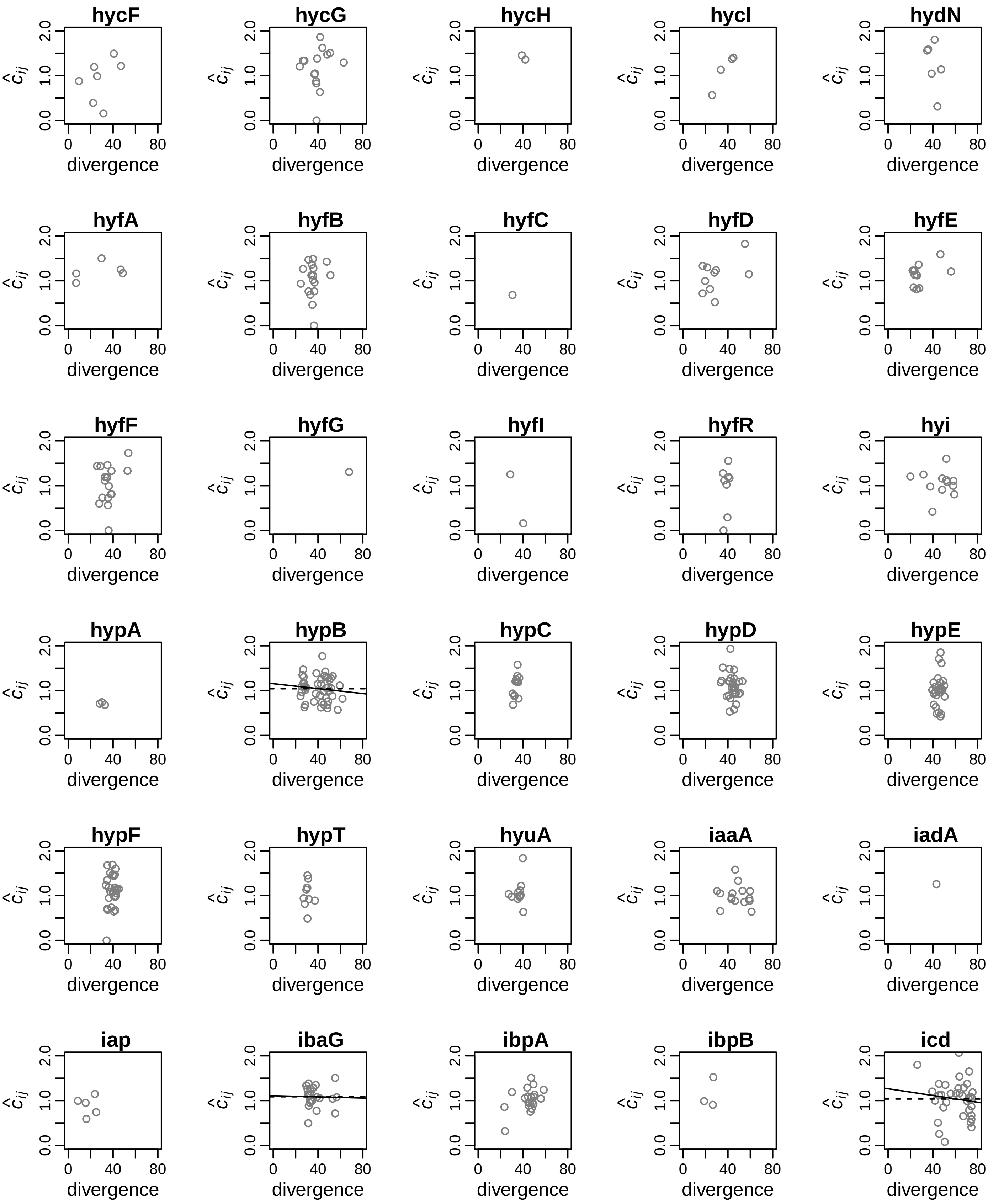
hycF
hycG
hycH
hycI
hydN
hyfA
hyfB
hyfC
hyfD
hyfE
hyfF
hyfG
hyfI
hyfR
hyi
hypA
hypB
hypC
hypD
hypE
hypF
hypT
hyuA
iaaA
iadA
iap
ibaG
ibpA
ibpB
icd
$\hat{c}_{ij}$
0.0
1.0
2.0
0
40
80
divergence

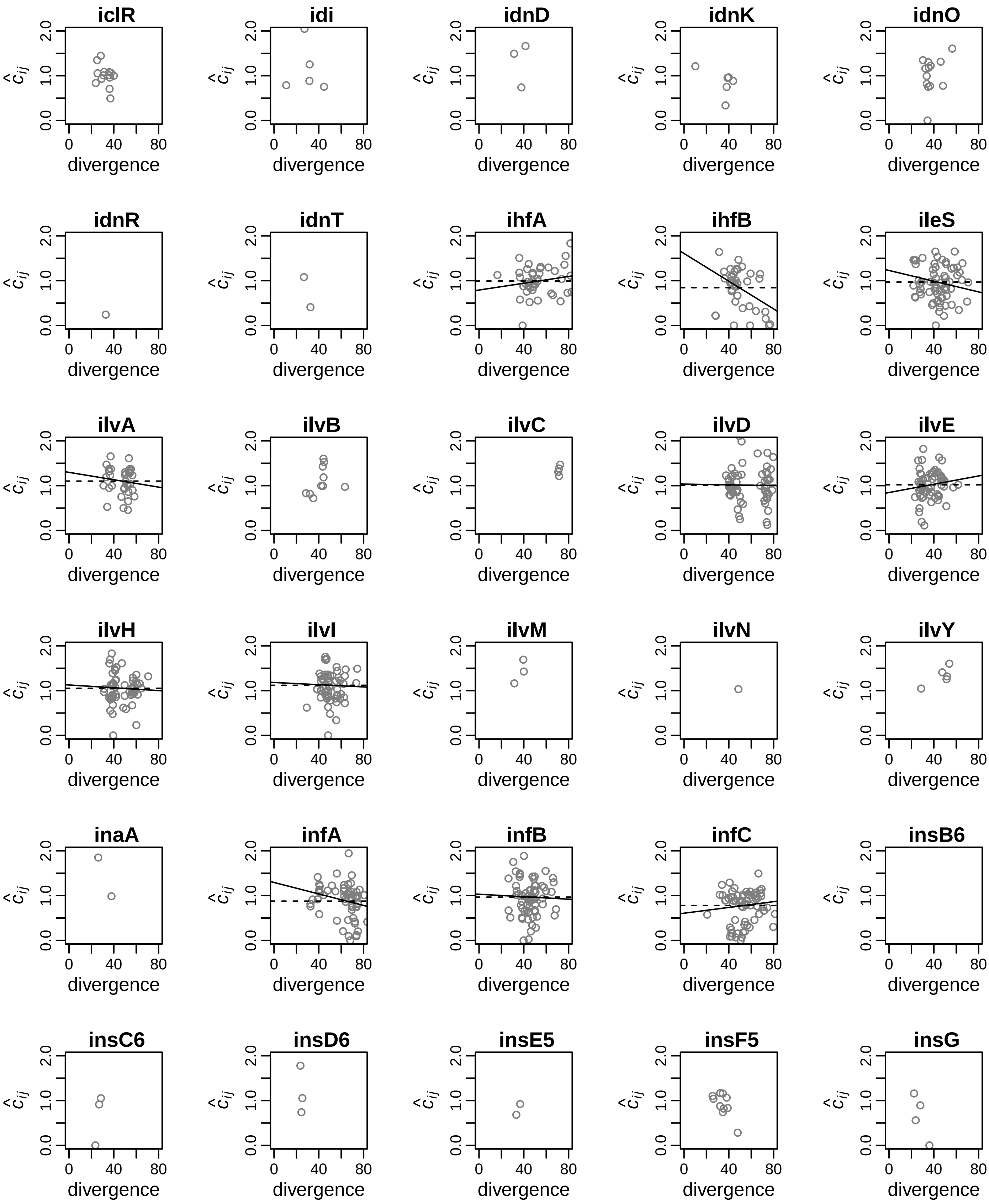
iclR
idi
idnD
idnK
idnO
idnR
idnT
ihfA
ihfB
ileS
ilvA
ilvB
ilvC
ilvD
ilvE
ilvH
ilvI
ilvM
ilvN
ilvY
inaA
infA
infB
infC
insB6
insC6
insD6
insE5
insF5
insG
$\hat{c}_{ij}$
0.0
1.0
2.0
0
40
80
divergence

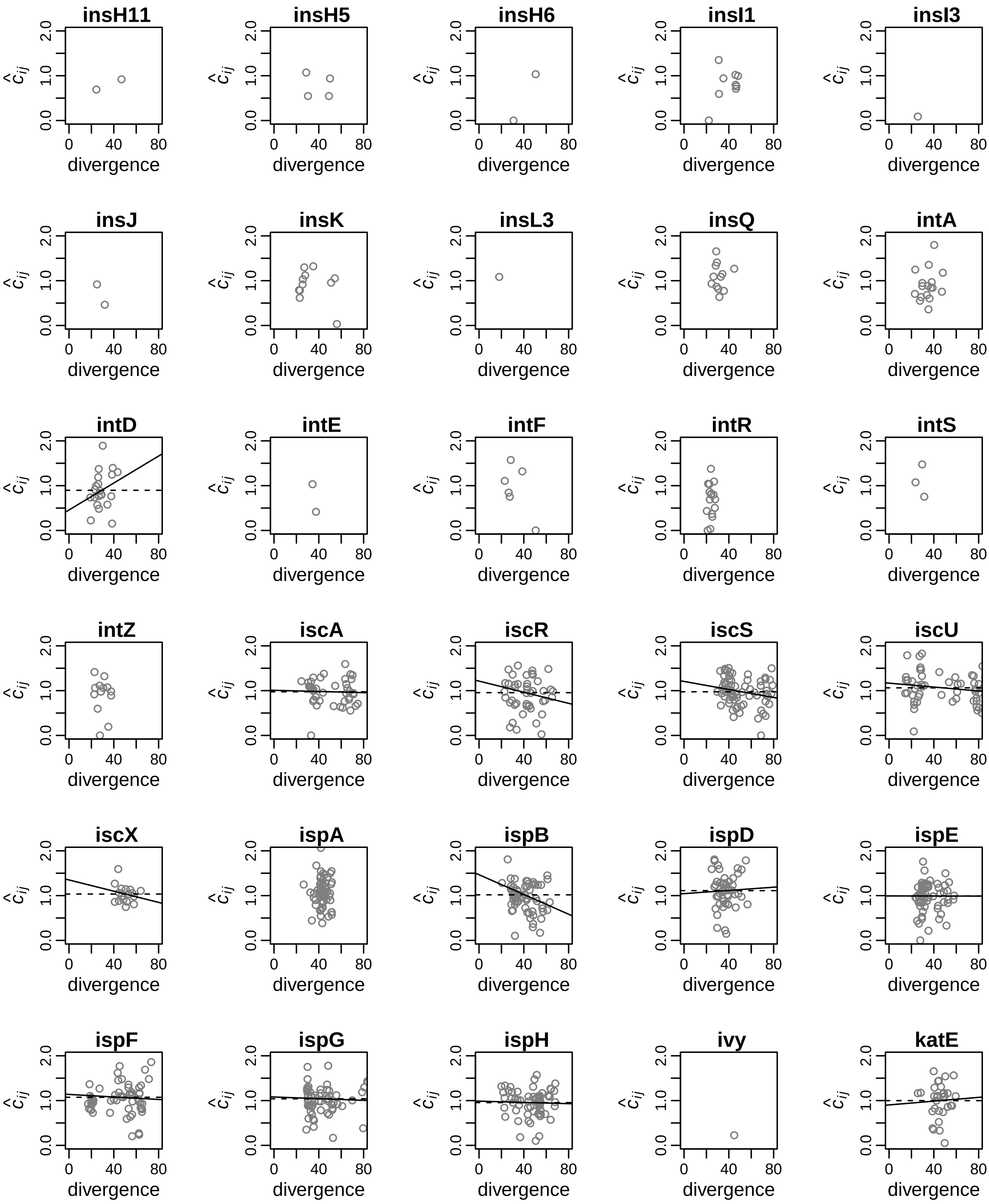
insH11
insH5
insH6
insI1
insI3
insJ
insK
insL3
insQ
intA
intD
intE
intF
intR
intS
intZ
iscA
iscR
iscS
iscU
iscX
ispA
ispB
ispD
ispE
ispF
ispG
ispH
ivy
katE
$\hat{c}_{ij}$
0.0
1.0
2.0
0
40
80
divergence

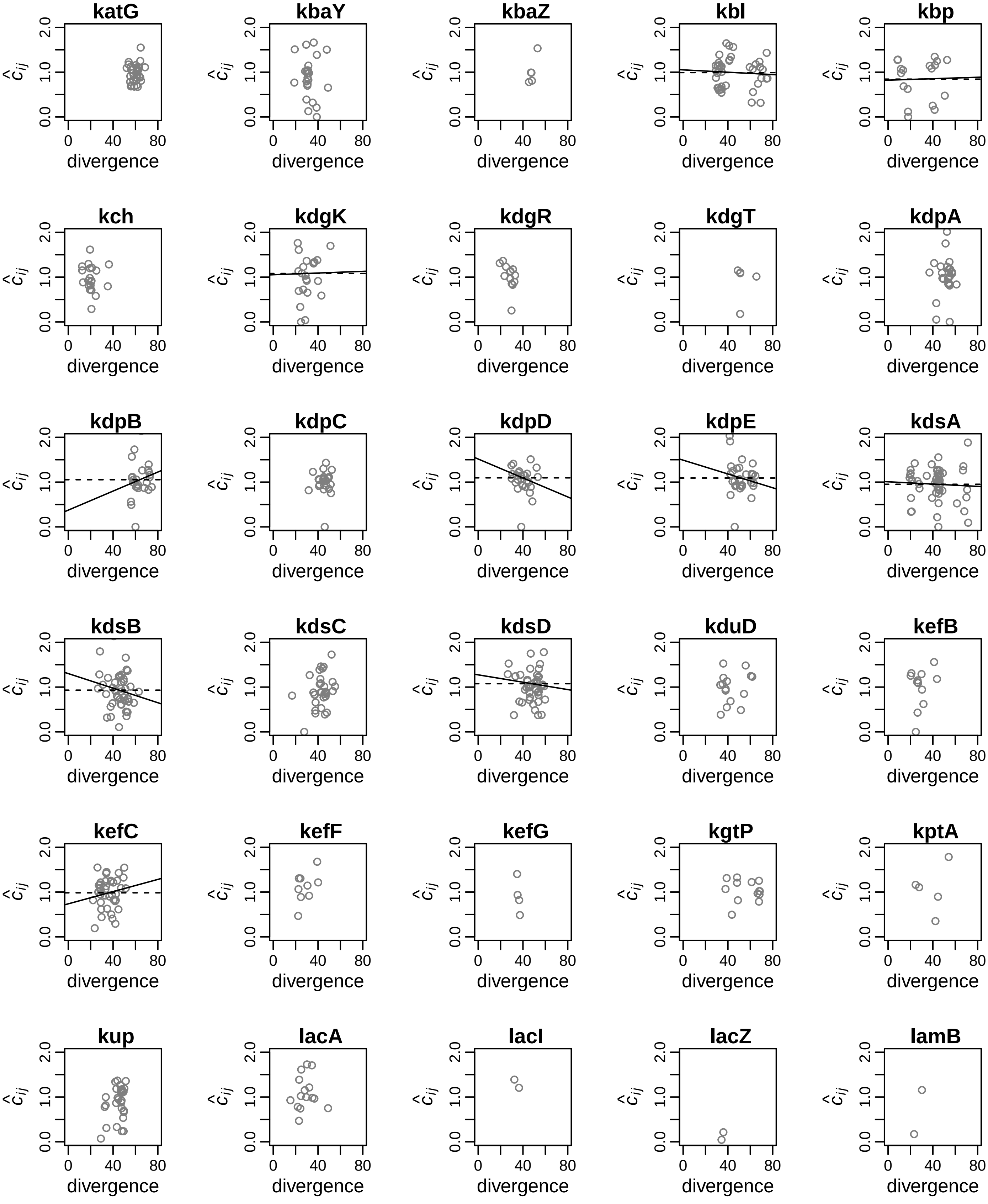
katG
kbaY
kbaZ
kbl
kbp
kch
kdgK
kdgR
kdgT
kdpA
kdpB
kdpC
kdpD
kdpE
kdsA
kdsB
kdsC
kdsD
kduD
kefB
kefC
kefF
kefG
kgtP
kptA
kup
lacA
lacI
lacZ
lamB
$\hat{c}_{ij}$
divergence
0.0
1.0
2.0
0
40
80

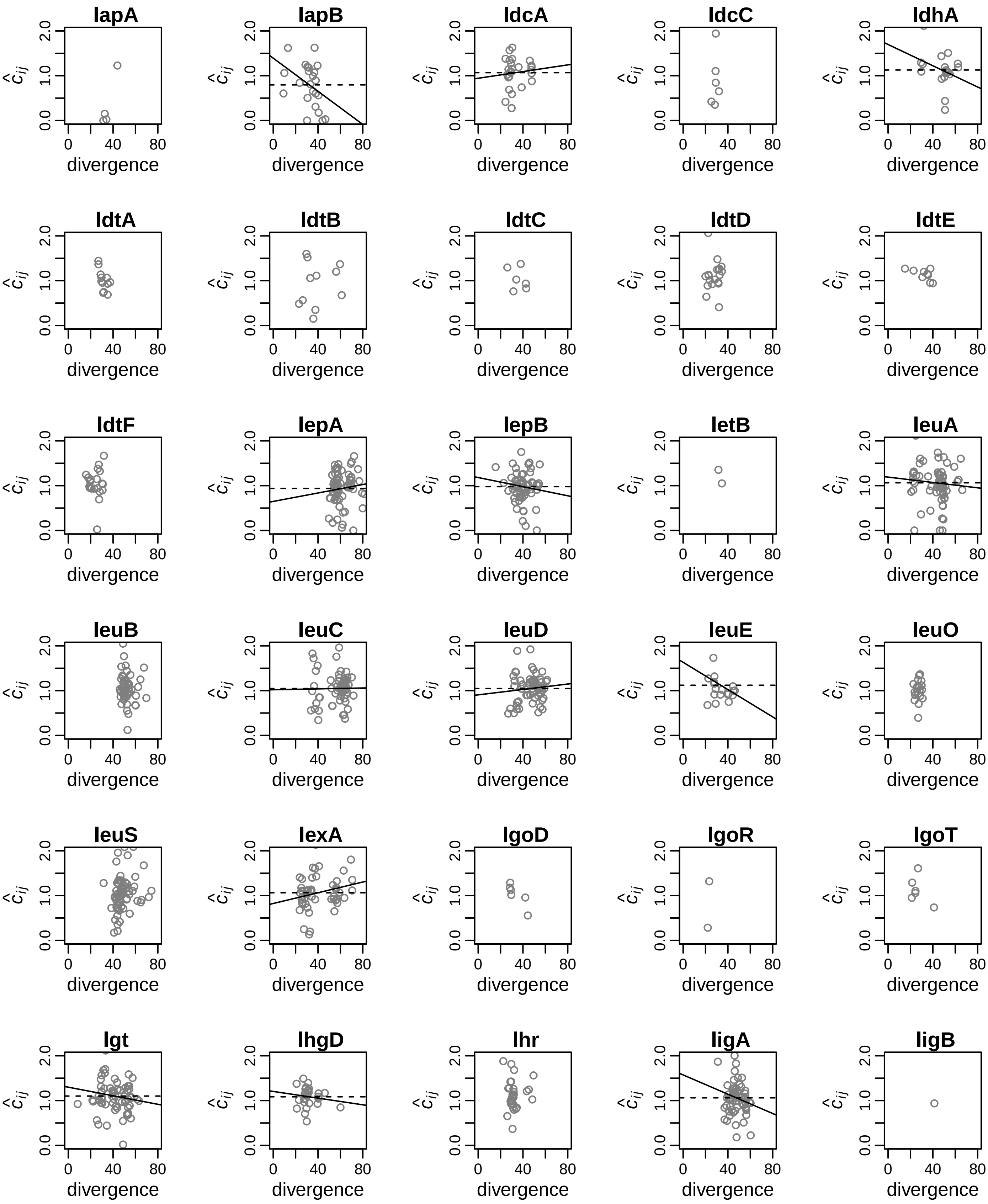

lapA
lapB
ldcA
ldcC
ldhA
ldtA
ldtB
ldtC
ldtD
ldtE
ldtF
lepA
lepB
letB
leuA
leuB
leuC
leuD
leuE
leuO
leuS
lexA
lgoD
lgoR
lgoT
lgt
lhgD
lhr
ligA
ligB
$\hat{c}_{ij}$
divergence
0.0
1.0
2.0
0
40
80

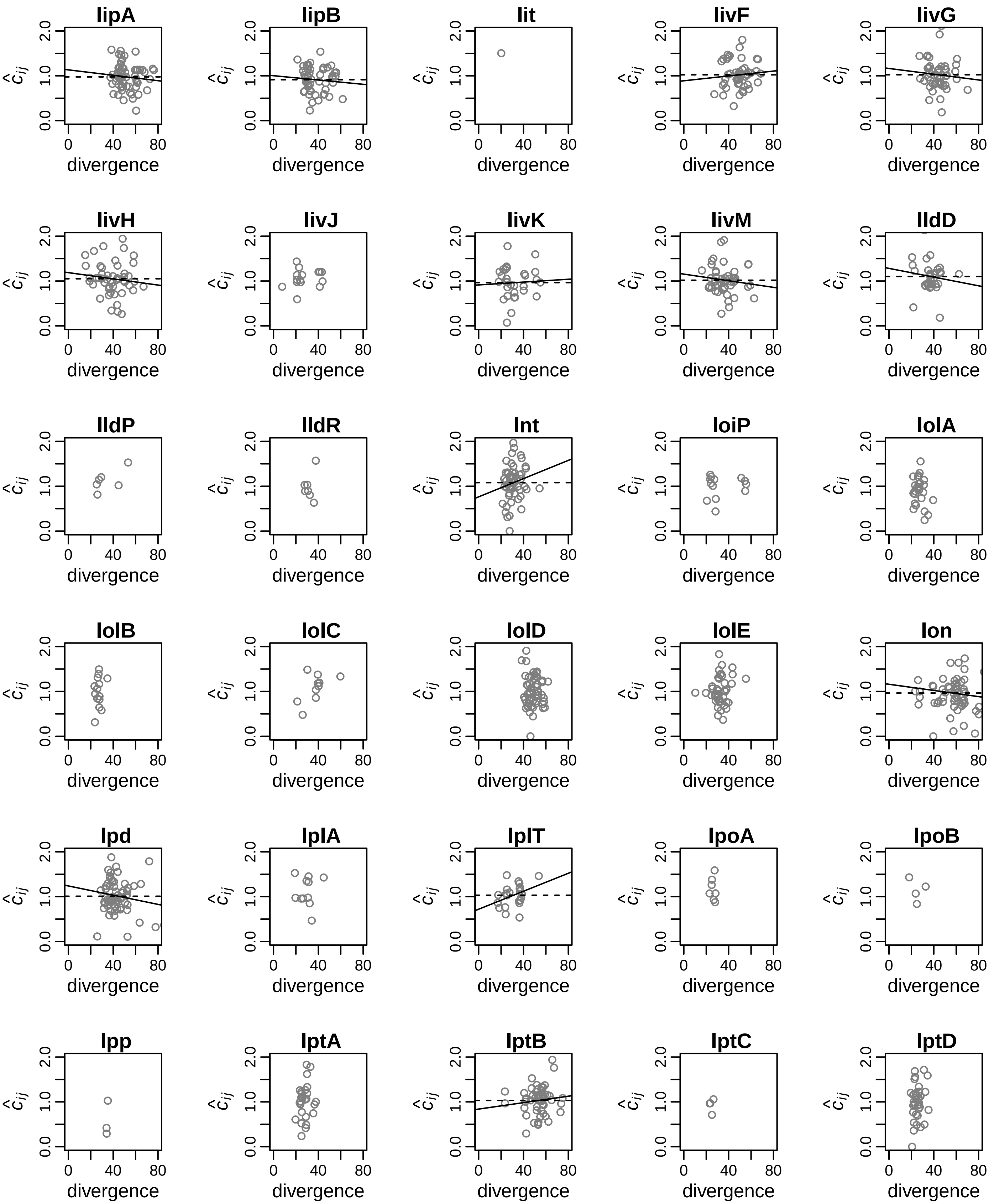
lipA
lipB
lit
livF
livG
livH
livJ
livK
livM
lldD
lldP
lldR
lnt
loiP
lolA
lolB
lolC
lolD
lolE
lon
lpd
lplA
lplT
lpoA
lpoB
lpp
lptA
lptB
lptC
lptD
$\hat{c}_{ij}$
0.0
1.0
2.0
0
40
80
divergence

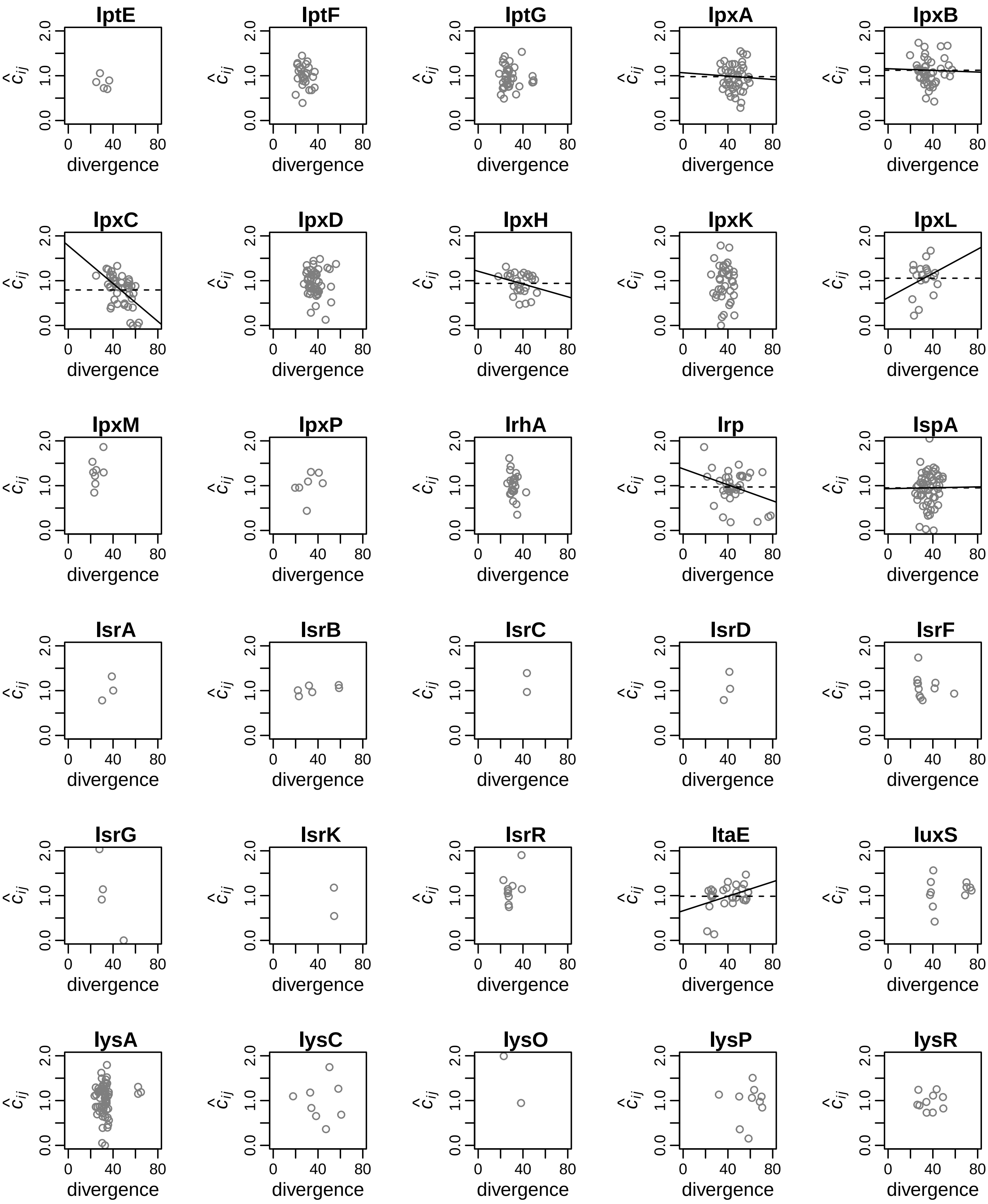

lptE
lptF
lptG
lpxA
lpxB
lpxC
lpxD
lpxH
lpxK
lpxL
lpxM
lpxP
lrhA
lrp
lspA
lsrA
lsrB
lsrC
lsrD
lsrF
lsrG
lsrK
lsrR
ltaE
luxS
lysA
lysC
lysO
lysP
lysR
$\hat{c}_{ij}$
0.0
1.0
2.0
0
40
80
divergence

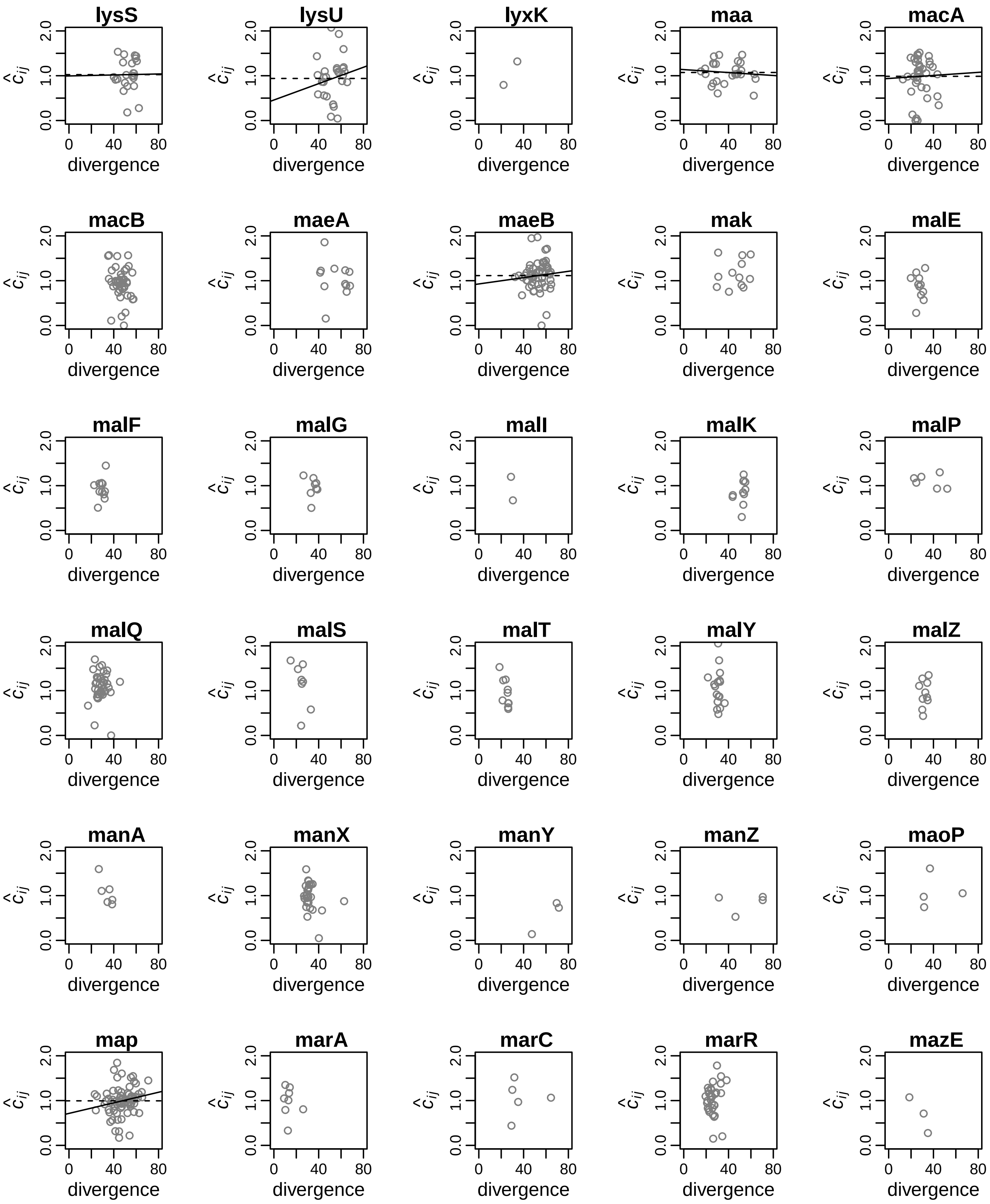

lysS
lysU
lyxK
maa
macA
macB
maeA
maeB
mak
malE
malF
malG
malI
malK
malP
malQ
malS
malT
malY
malZ
manA
manX
manY
manZ
maoP
map
marA
marC
marR
mazE
$\hat{c}_{ij}$
0.0
1.0
2.0
0
40
80
divergence

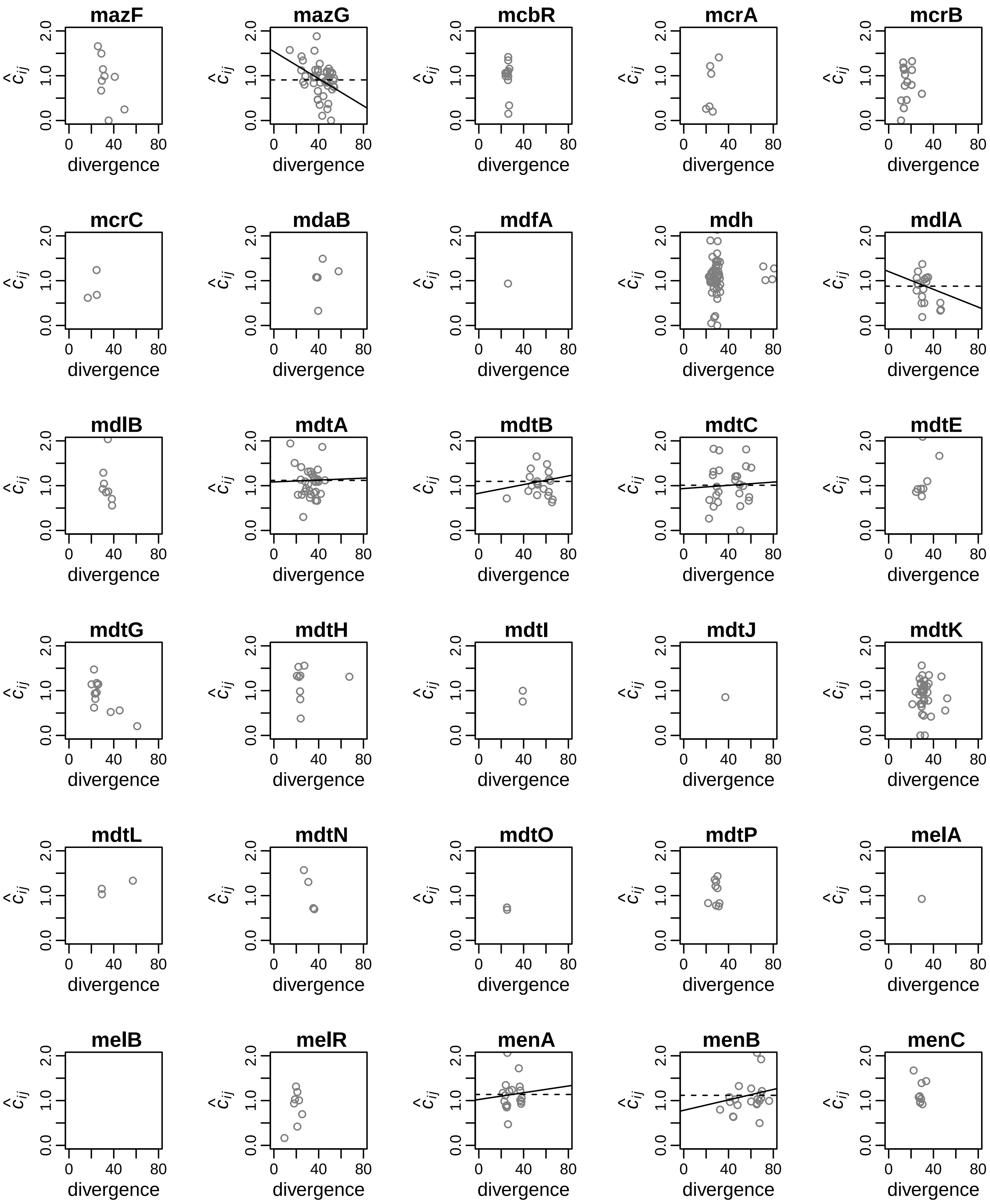
mazF
mazG
mcbR
mcrA
mcrB
mcrC
mdaB
mdfA
mdh
mdlA
mdlB
mdtA
mdtB
mdtC
mdtE
mdtG
mdtH
mdtI
mdtJ
mdtK
mdtL
mdtN
mdtO
mdtP
melA
melB
melR
menA
menB
menC
$\hat{c}_{ij}$
divergence
0.0
1.0
2.0
0
40
80

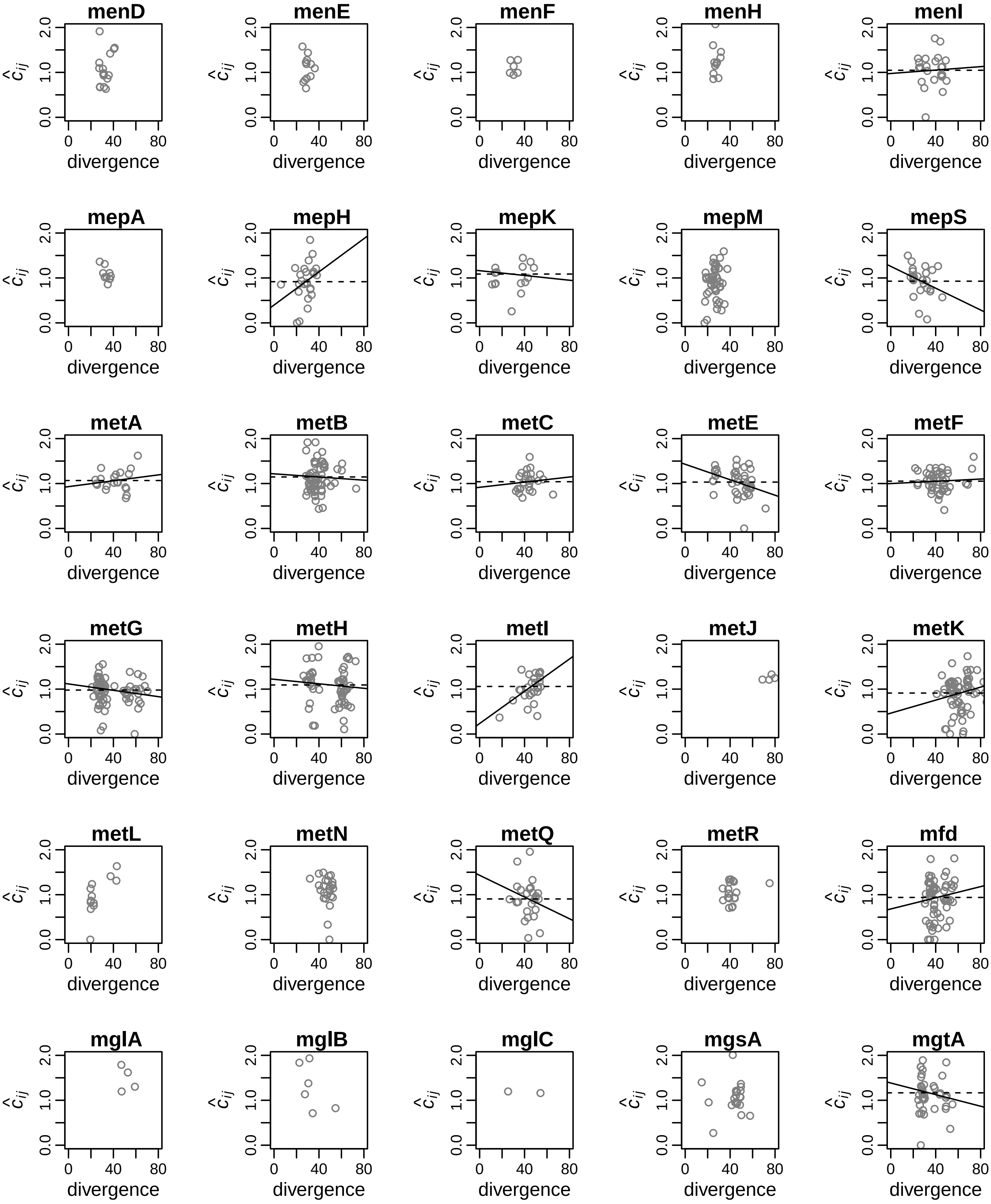
menD
menE
menF
menH
menI
mepA
mepH
mepK
mepM
mepS
metA
metB
metC
metE
metF
metG
metH
metI
metJ
metK
metL
metN
metQ
metR
mfd
mglA
mglB
mglC
mgsA
mgtA
divergence
$\hat{c}_{ij}$
0
40
80
0.0
1.0
2.0

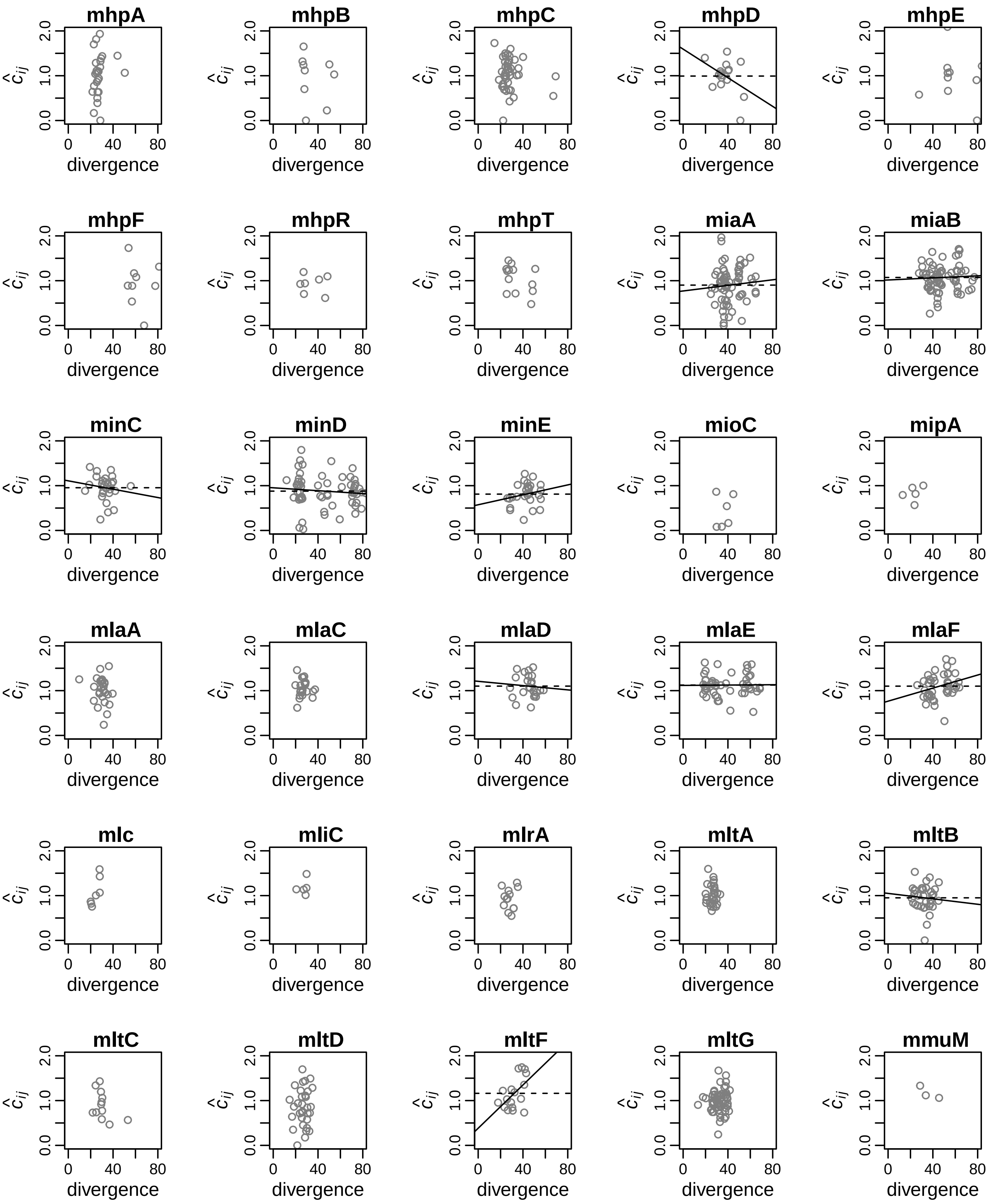
mhpA
mhpB
mhpC
mhpD
mhpE
mhpF
mhpR
mhpT
miaA
miaB
minC
minD
minE
mioC
mipA
mlaA
mlaC
mlaD
mlaE
mlaF
mlc
mliC
mlrA
mltA
mltB
mltC
mltD
mltF
mltG
mmuM
$\hat{c}_{ij}$
0.0
1.0
2.0
divergence
0
40
80

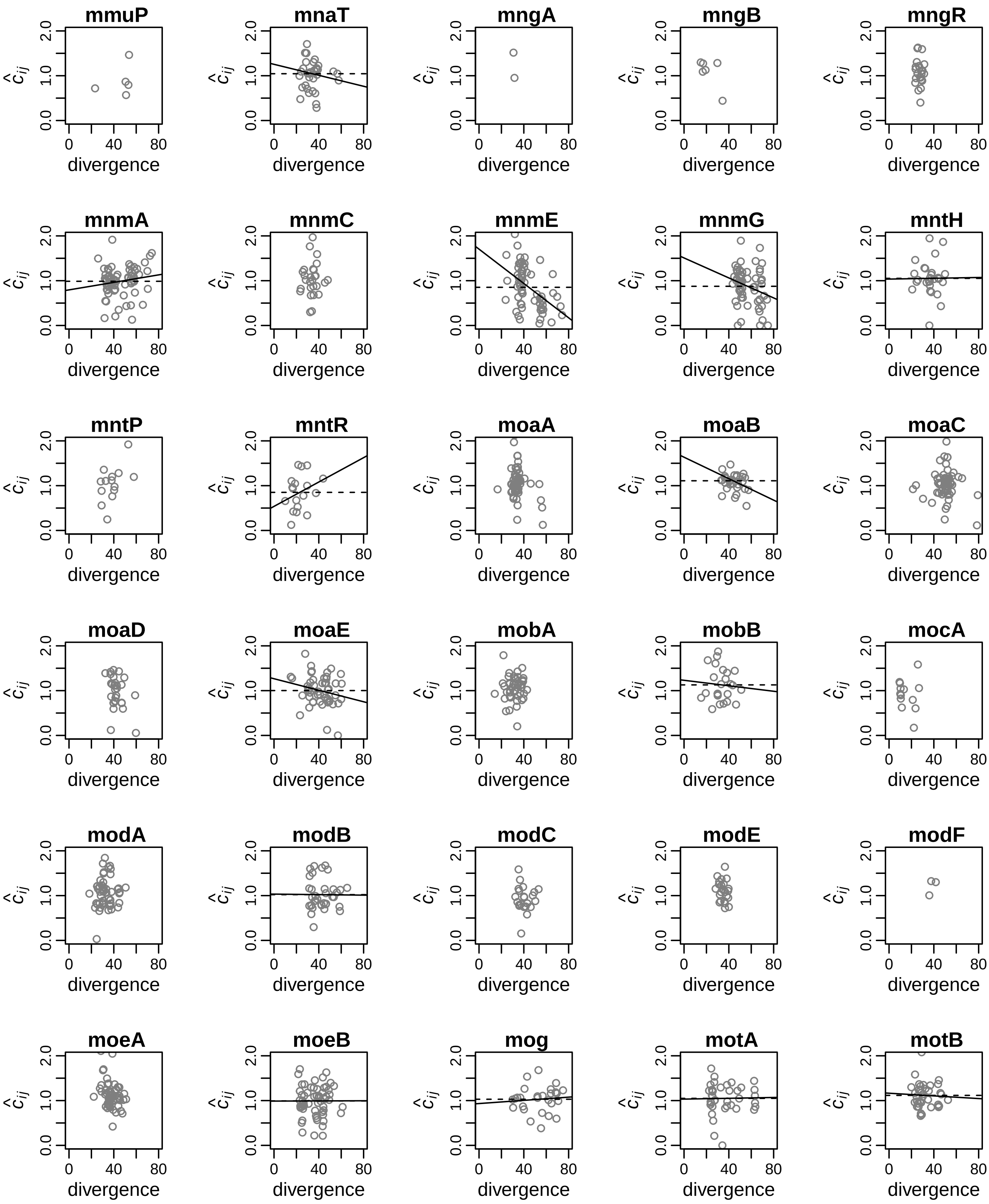
mmuP
mnaT
mngA
mngB
mngR
mnmA
mnmC
mnmE
mnmG
mntH
mntP
mntR
moaA
moaB
moaC
moaD
moaE
mobA
mobB
mocA
modA
modB
modC
modE
modF
moeA
moeB
mog
motA
motB
$\hat{c}_{ij}$
divergence
0.0
1.0
2.0
0
40
80

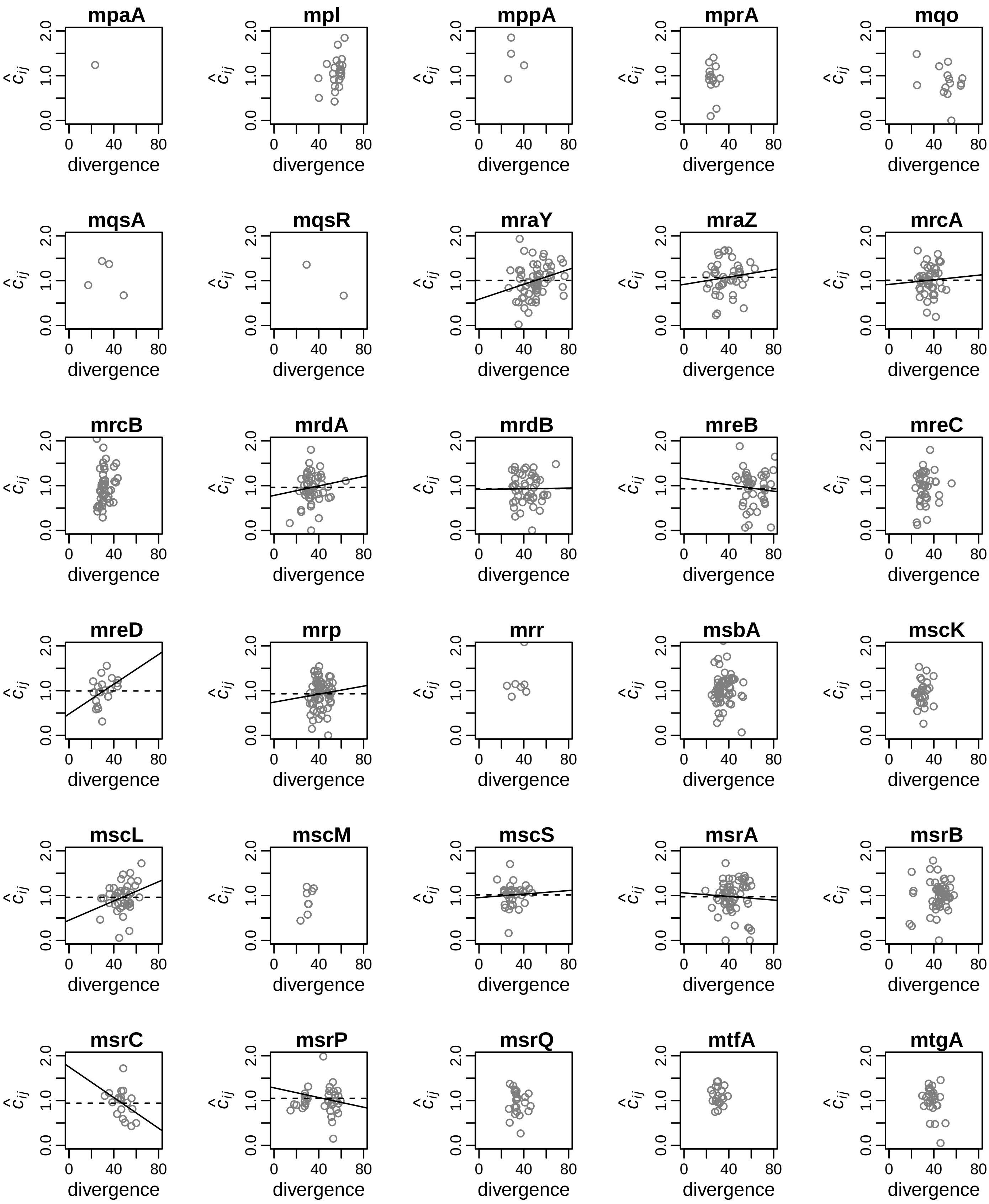
mpaA
mpl
mppA
mprA
mqo
mqsA
mqsR
mraY
mraZ
mrcA
mrcB
mrdA
mrdB
mreB
mreC
mreD
mrp
mrr
msbA
mscK
mscL
mscM
mscS
msrA
msrB
msrC
msrP
msrQ
mtfA
mtgA
$\hat{c}_{ij}$
divergence
0.0
1.0
2.0
0
40
80

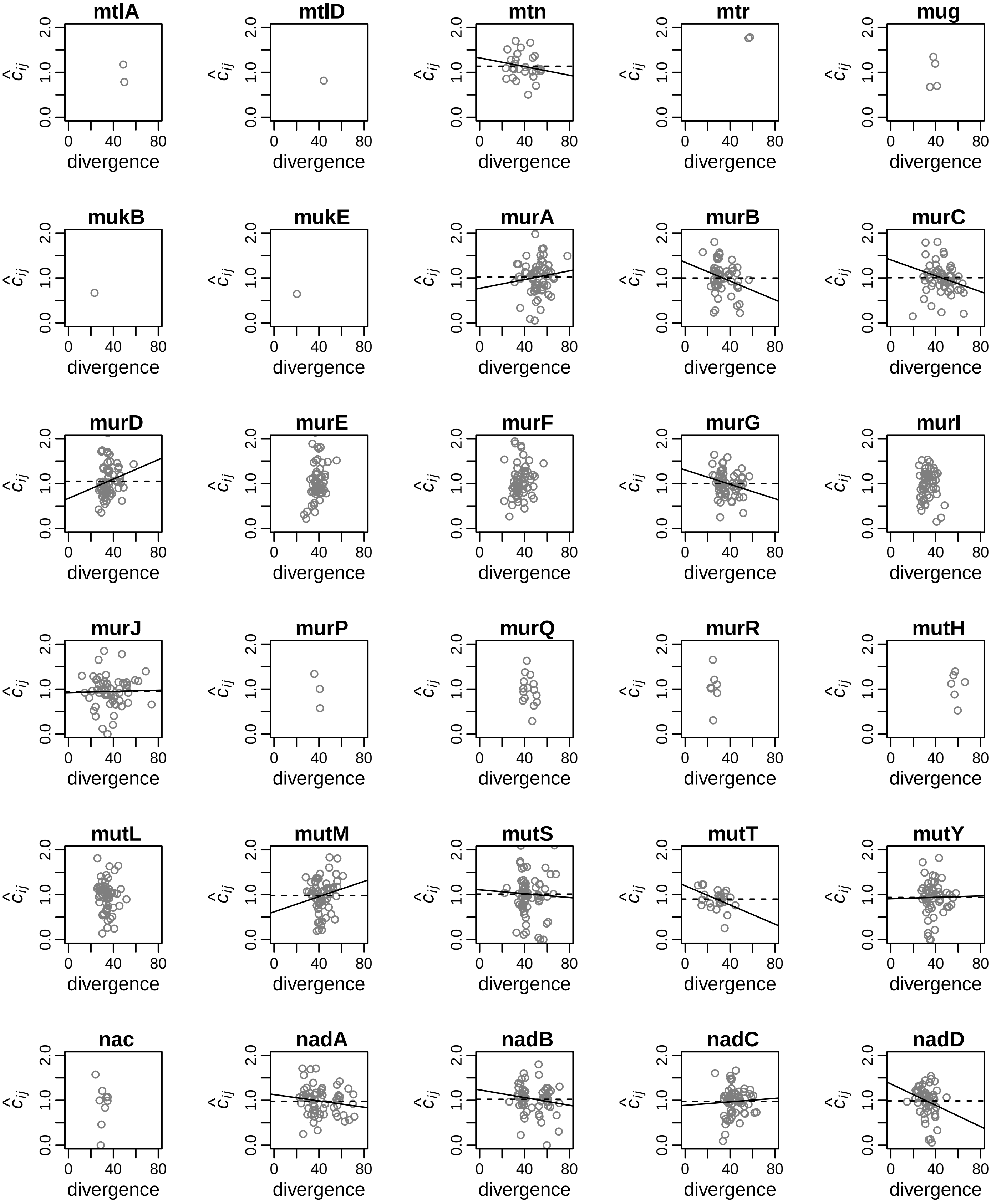
mtlA
mtlD
mtn
mtr
mug
mukB
mukE
murA
murB
murC
murD
murE
murF
murG
murI
murJ
murP
murQ
murR
mutH
mutL
mutM
mutS
mutT
mutY
nac
nadA
nadB
nadC
nadD
$\hat{c}_{ij}$
0.0
1.0
2.0
0
40
80
divergence

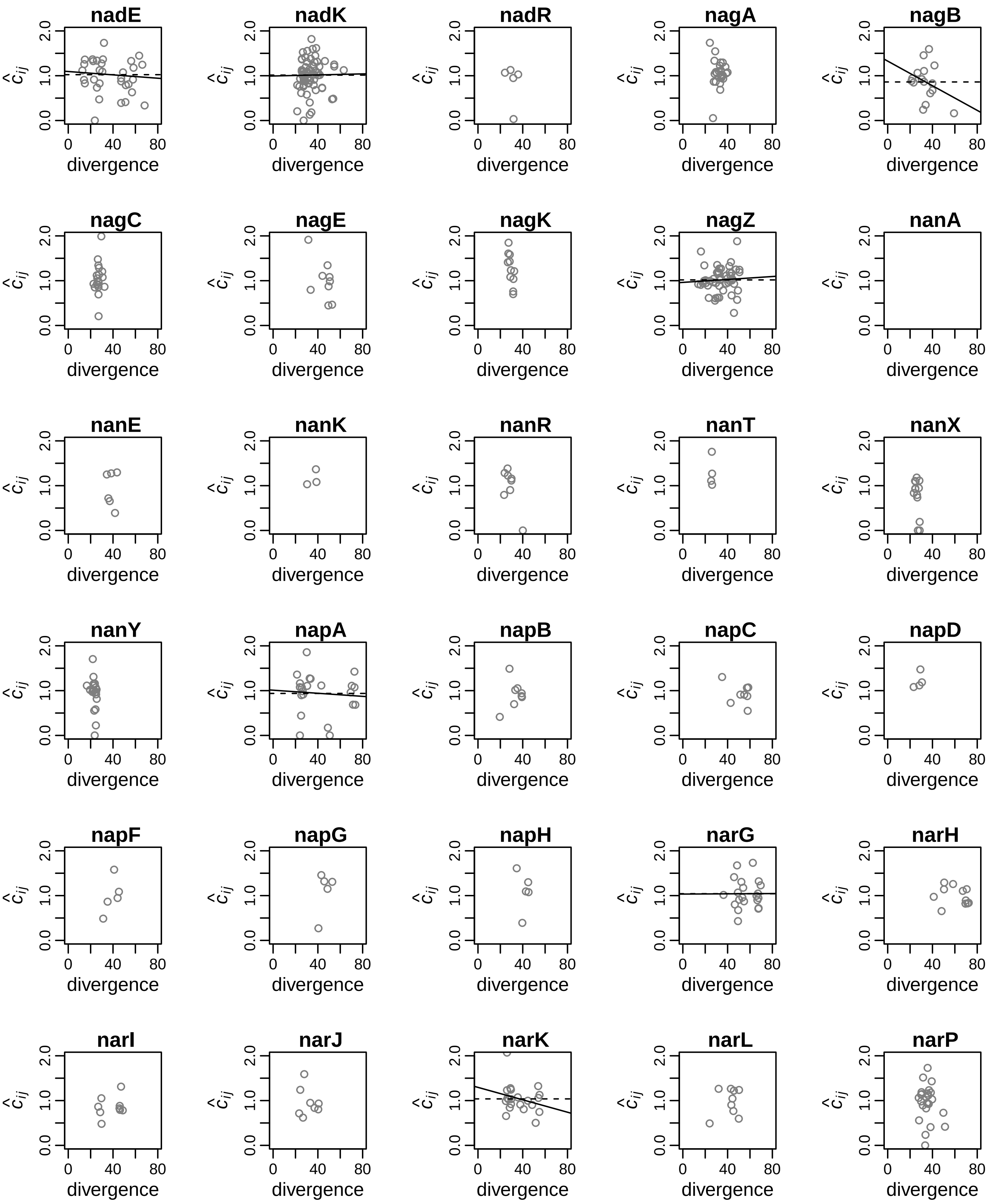
nadE
nadK
nadR
nagA
nagB
nagC
nagE
nagK
nagZ
nanA
nanE
nanK
nanR
nanT
nanX
nanY
napA
napB
napC
napD
napF
napG
napH
narG
narH
narI
narJ
narK
narL
narP
$\hat{c}_{ij}$
0.0
1.0
2.0
0
40
80
divergence

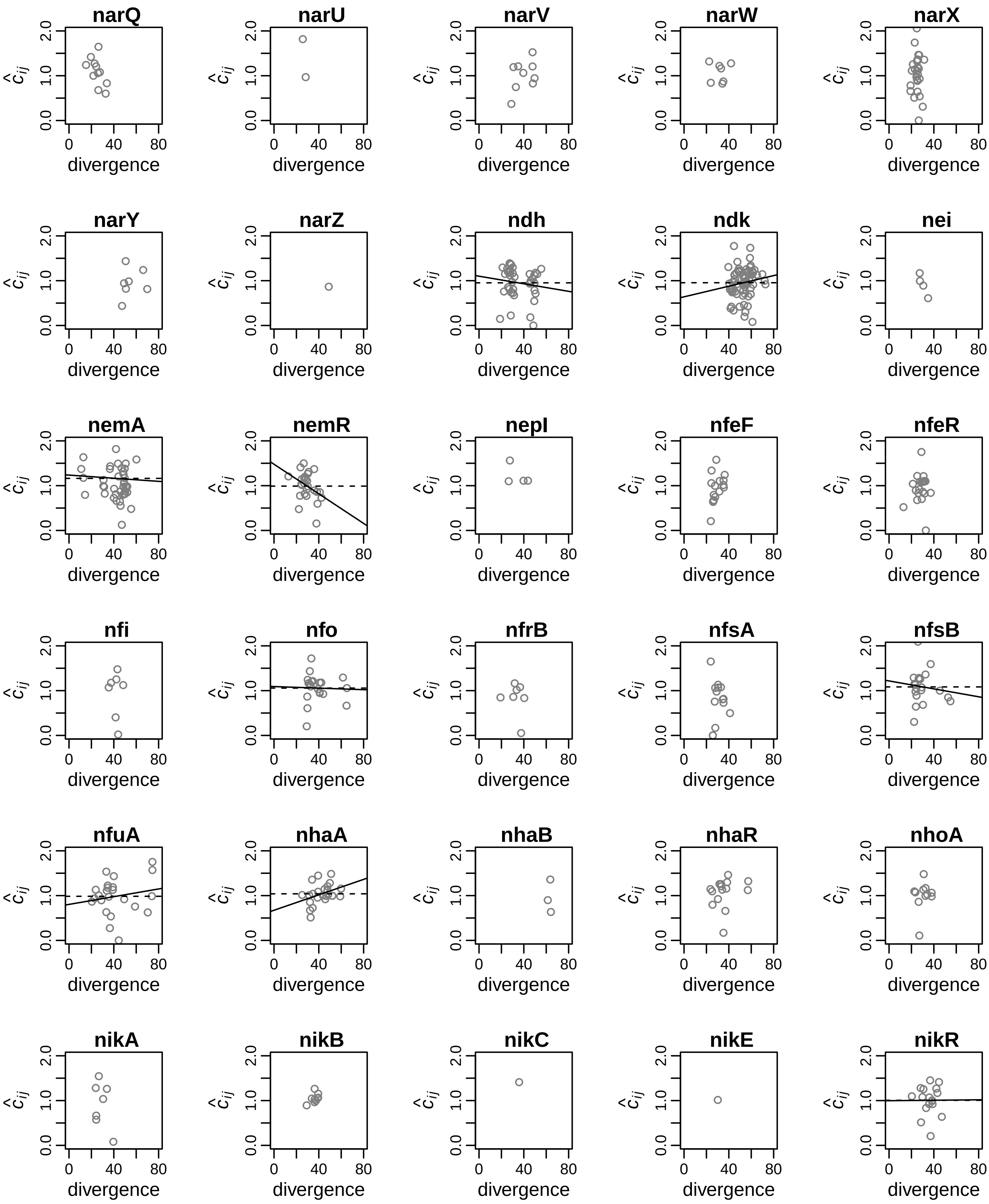
narQ
narU
narV
narW
narX
narY
narZ
ndh
ndk
nei
nemA
nemR
nepI
nfeF
nfeR
nfi
nfo
nfrB
nfsA
nfsB
nfuA
nhaA
nhaB
nhaR
nhoA
nikA
nikB
nikC
nikE
nikR
divergence
$\hat{c}_{ij}$
0.0
1.0
2.0
0
40
80

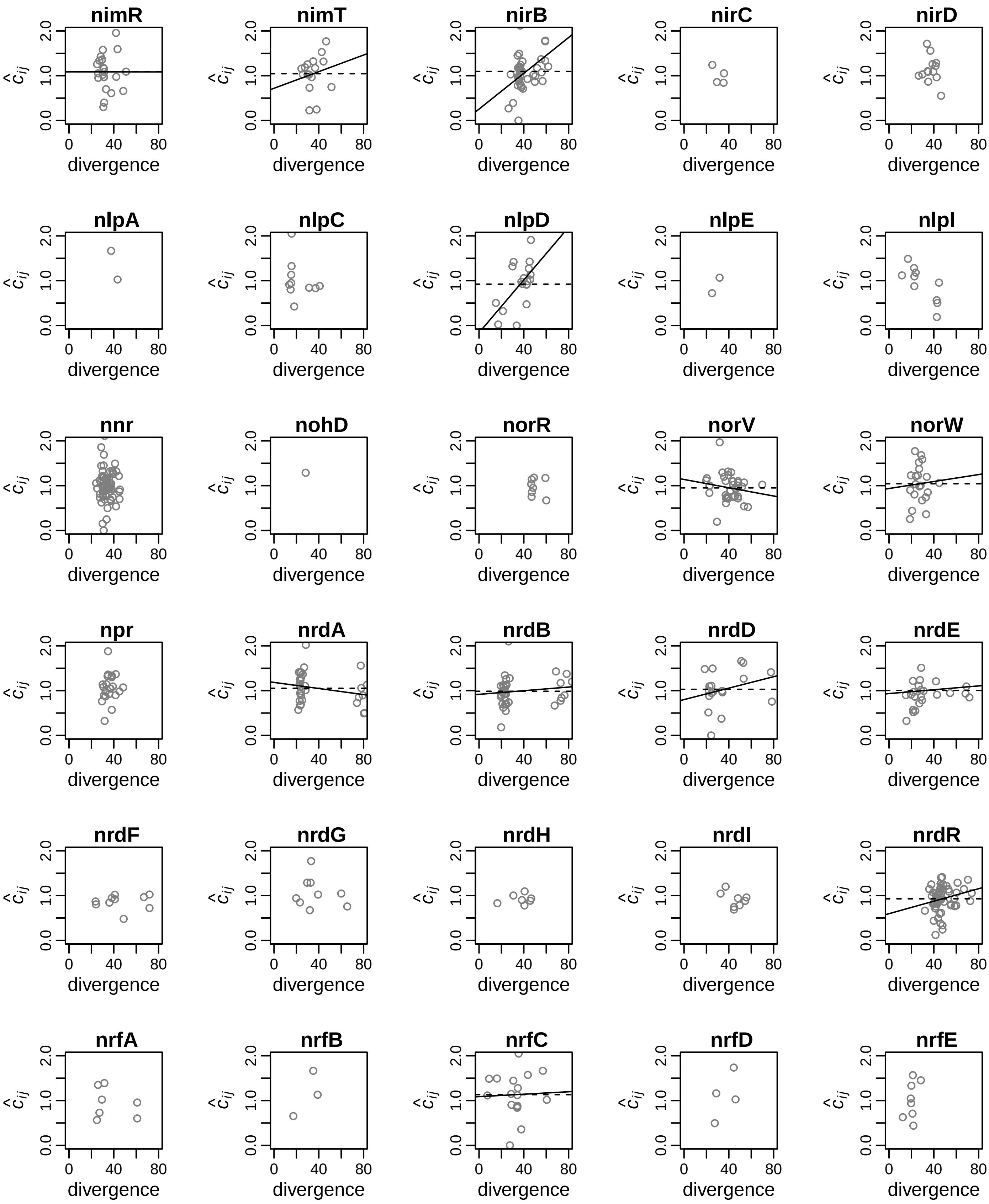
nimR
nimT
nirB
nirC
nirD
nlpA
nlpC
nlpD
nlpE
nlpI
nnr
nohD
norR
norV
norW
npr
nrdA
nrdB
nrdD
nrdE
nrdF
nrdG
nrdH
nrdI
nrdR
nrfA
nrfB
nrfC
nrfD
nrfE
$\hat{c}_{ij}$
divergence
0.0
1.0
2.0
0
40
80

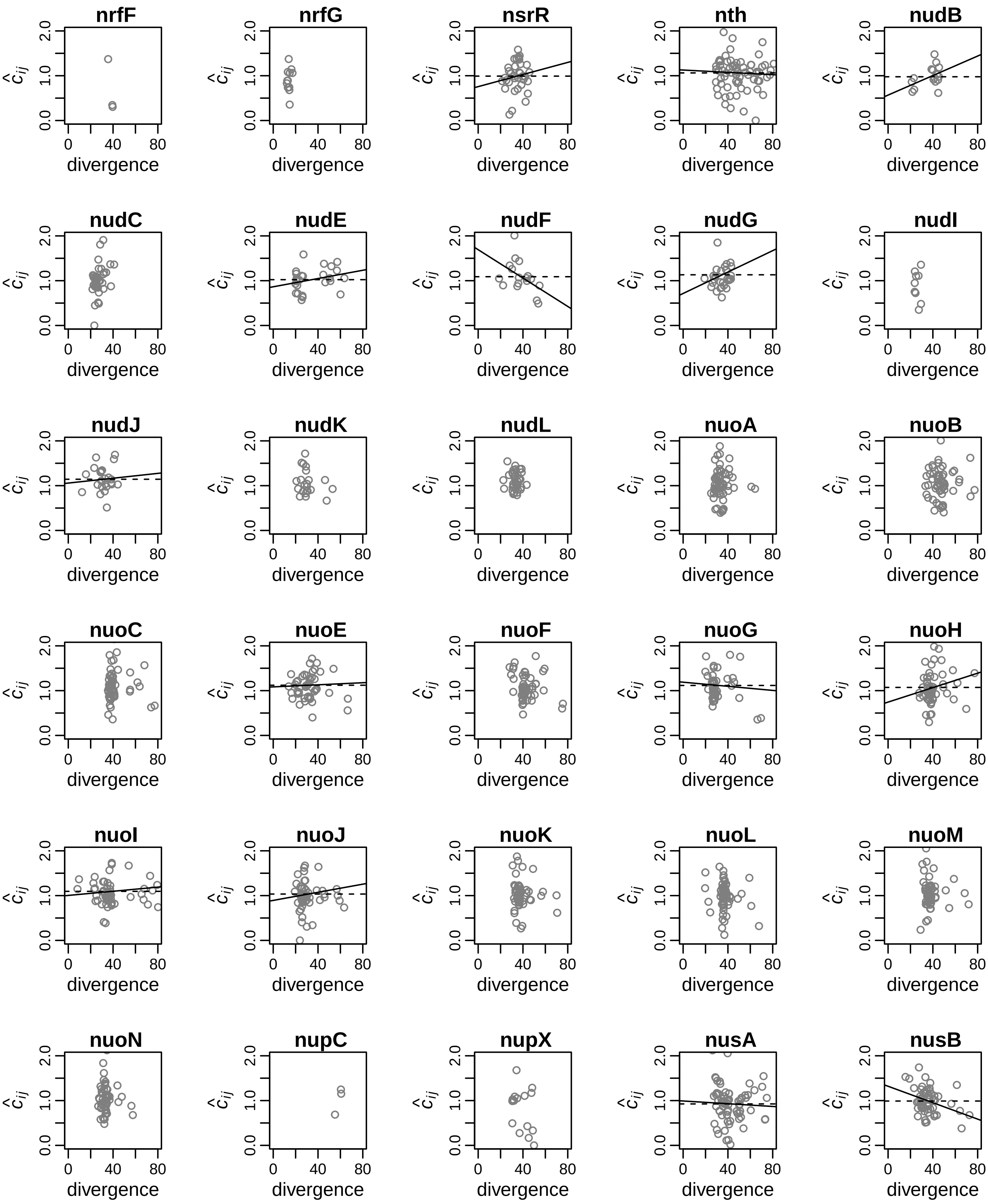
nrfF
nrfG
nsrR
nth
nudB
nudC
nudE
nudF
nudG
nudI
nudJ
nudK
nudL
nuoA
nuoB
nuoC
nuoE
nuoF
nuoG
nuoH
nuoI
nuoJ
nuoK
nuoL
nuoM
nuoN
nupC
nupX
nusA
nusB
$\hat{c}_{ij}$
0.0
1.0
2.0
0
40
80
divergence

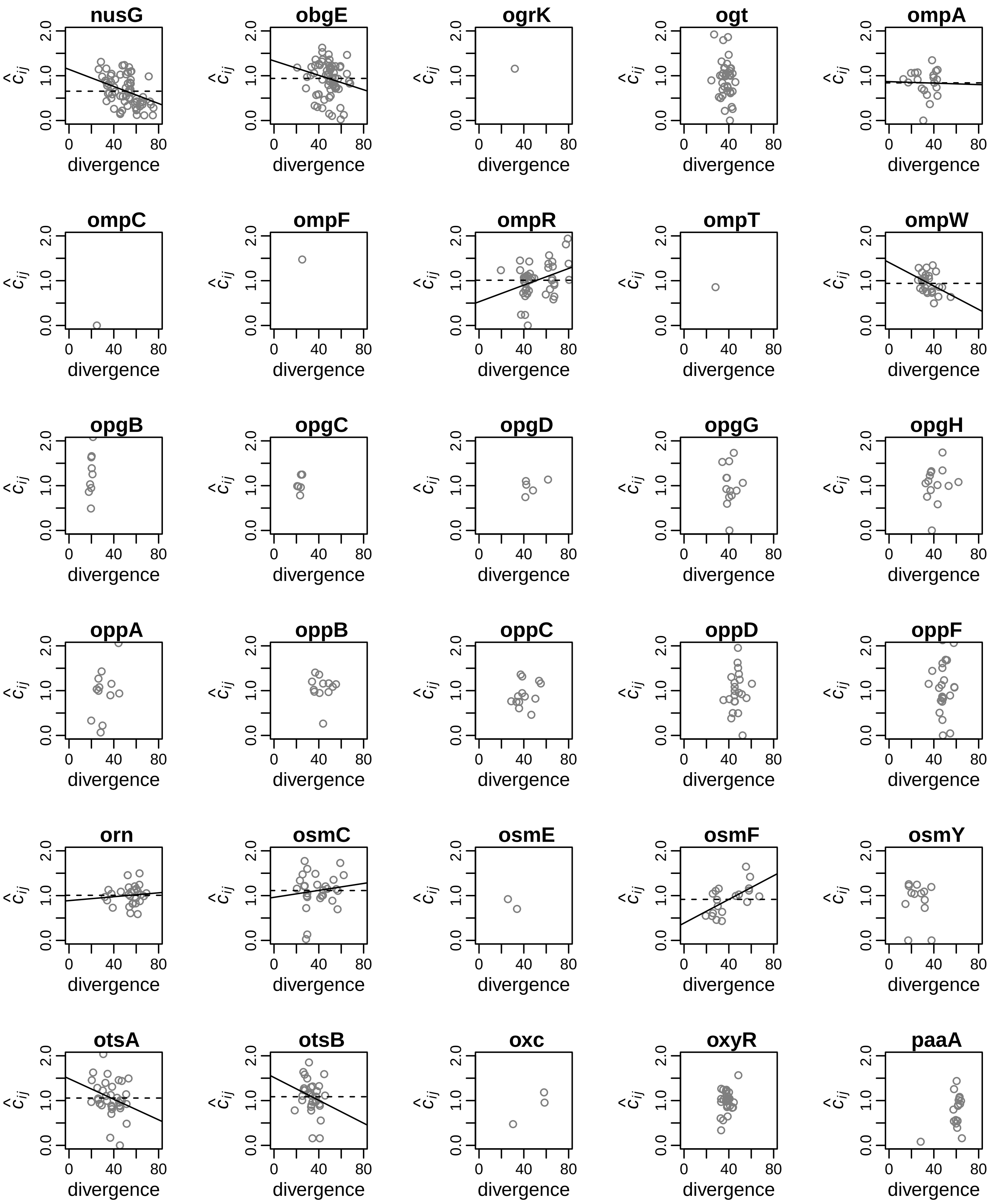
nusG
obgE
ogrK
ogt
ompA
ompC
ompF
ompR
ompT
ompW
opgB
opgC
opgD
opgG
opgH
oppA
oppB
oppC
oppD
oppF
orn
osmC
osmE
osmF
osmY
otsA
otsB
oxc
oxyR
paaA
$\hat{c}_{ij}$
0.0
1.0
2.0
0
40
80
divergence

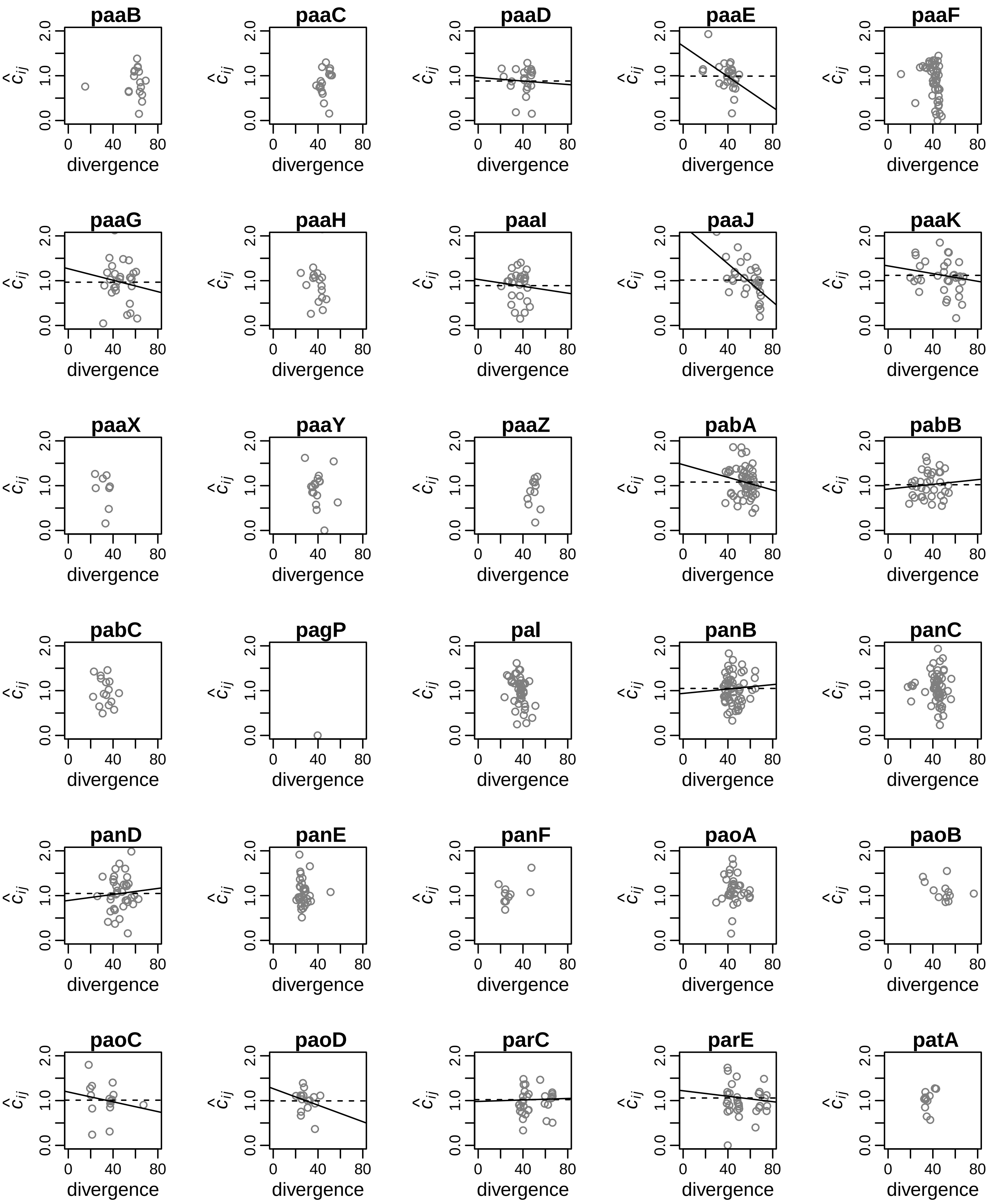

paaB
paaC
paaD
paaE
paaF
paaG
paaH
paaI
paaJ
paaK
paaX
paaY
paaZ
pabA
pabB
pabC
pagP
pal
panB
panC
panD
panE
panF
paoA
paoB
paoC
paoD
parC
parE
patA
$\hat{c}_{ij}$
0.0
1.0
2.0
0
40
80
divergence

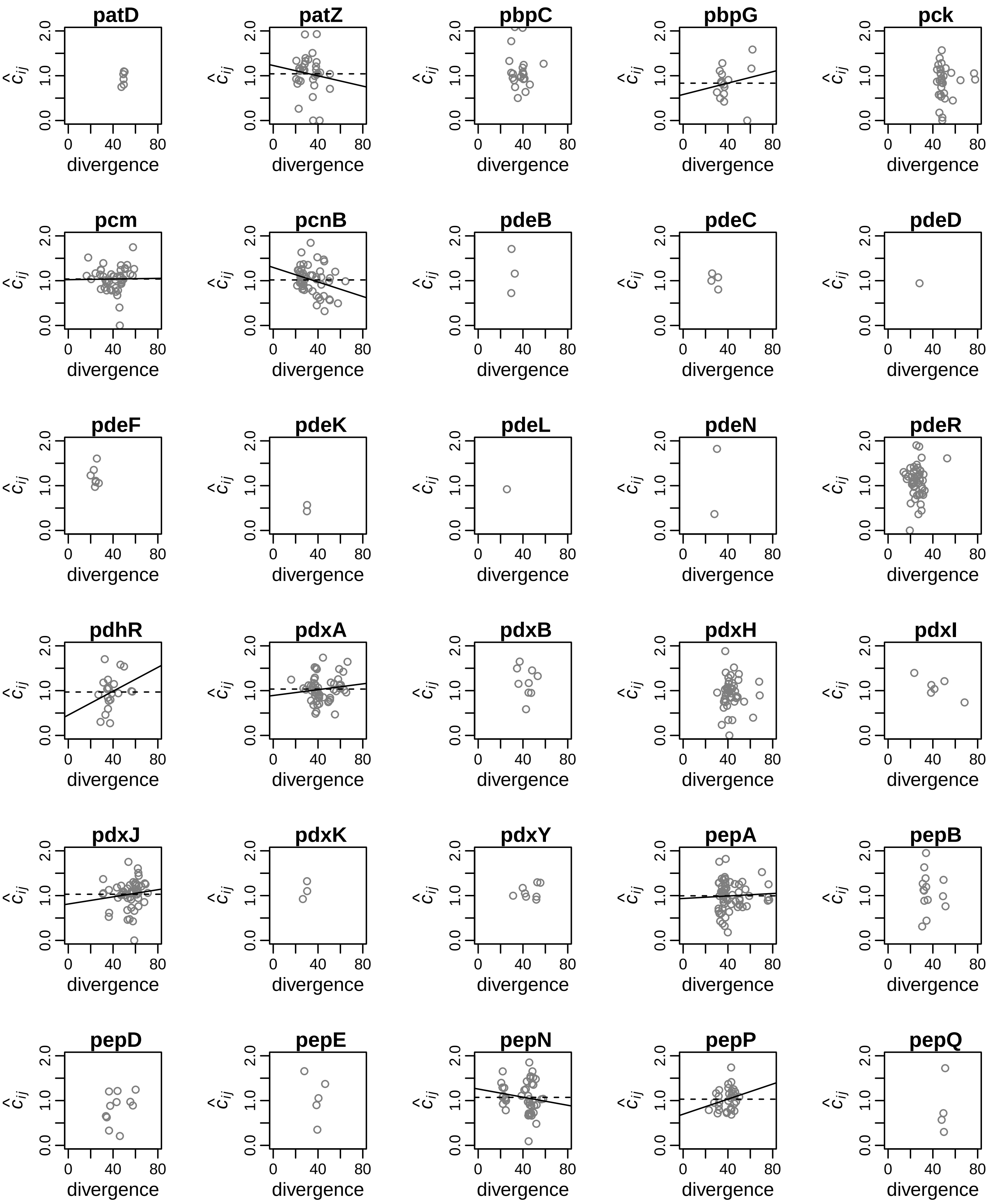
patD
patZ
pbpC
pbpG
pck
pcm
pcnB
pdeB
pdeC
pdeD
pdeF
pdeK
pdeL
pdeN
pdeR
pdhR
pdxA
pdxB
pdxH
pdxI
pdxJ
pdxK
pdxY
pepA
pepB
pepD
pepE
pepN
pepP
pepQ
$\hat{c}_{ij}$
0.0 1.0 2.0
0 40 80
divergence

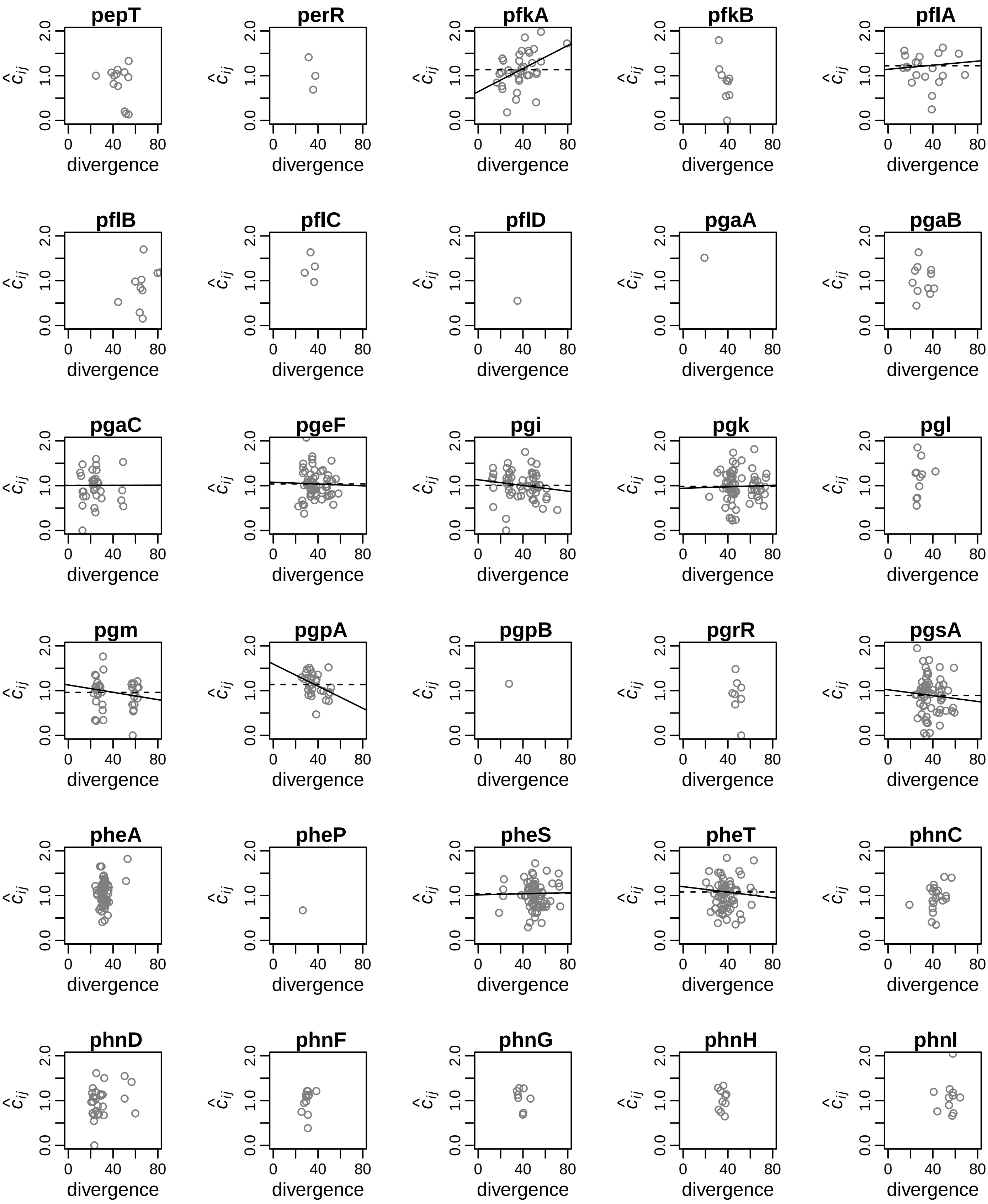

pepT
perR
pfkA
pfkB
pflA
pflB
pflC
pflD
pgaA
pgaB
pgaC
pgeF
pgi
pgk
pgl
pgm
pgpA
pgpB
pgrR
pgsA
pheA
pheP
pheS
pheT
phnC
phnD
phnF
phnG
phnH
phnI
$\hat{c}_{ij}$
0.0
1.0
2.0
divergence
0
40
80

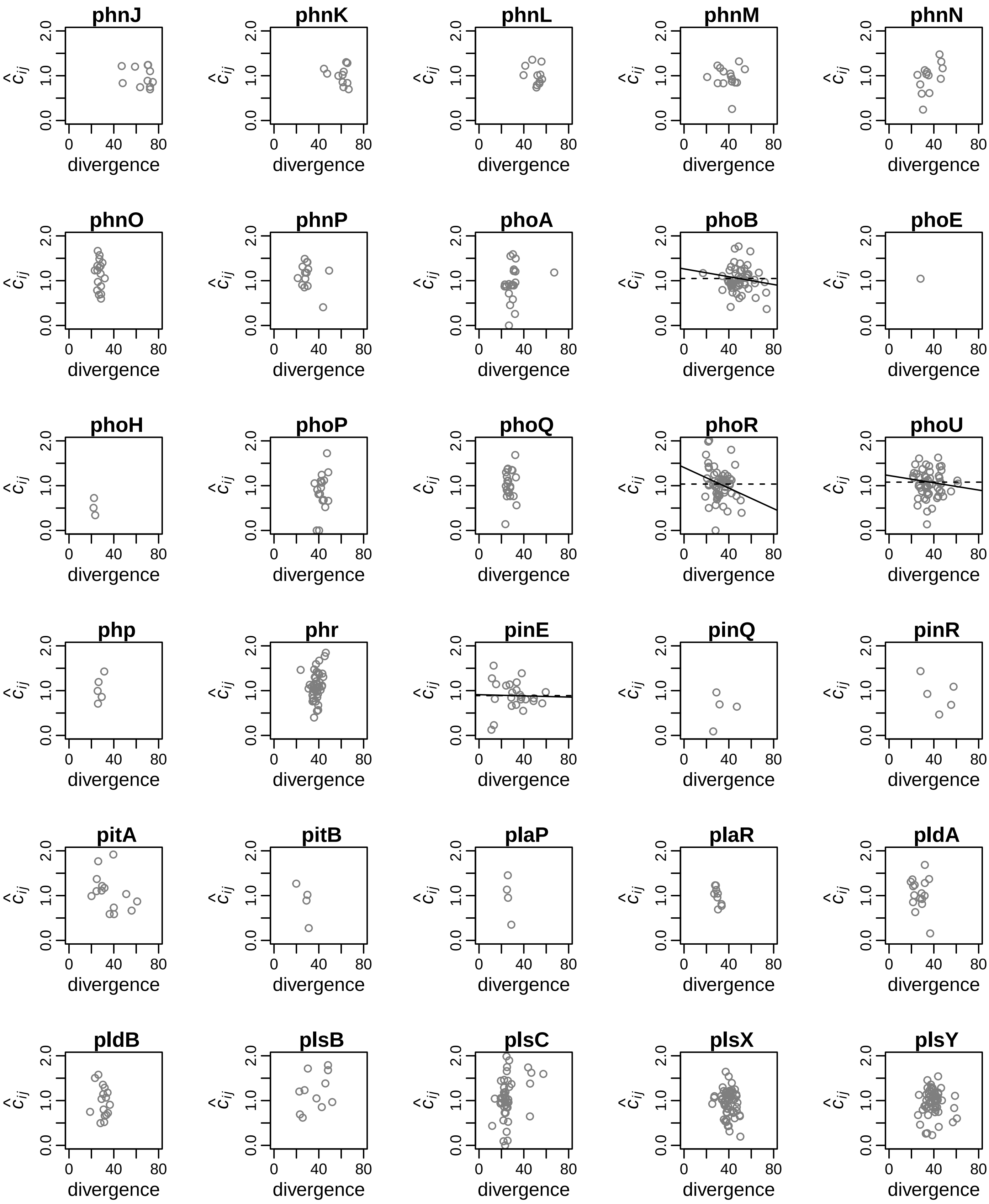

phnJ
phnK
phnL
phnM
phnN
phnO
phnP
phoA
phoB
phoE
phoH
phoP
phoQ
phoR
phoU
php
phr
pinE
pinQ
pinR
pitA
pitB
plaP
plaR
pldA
pldB
plsB
plsC
plsX
plsY
$\hat{c}_{ij}$
divergence
0.0
1.0
2.0
0
40
80

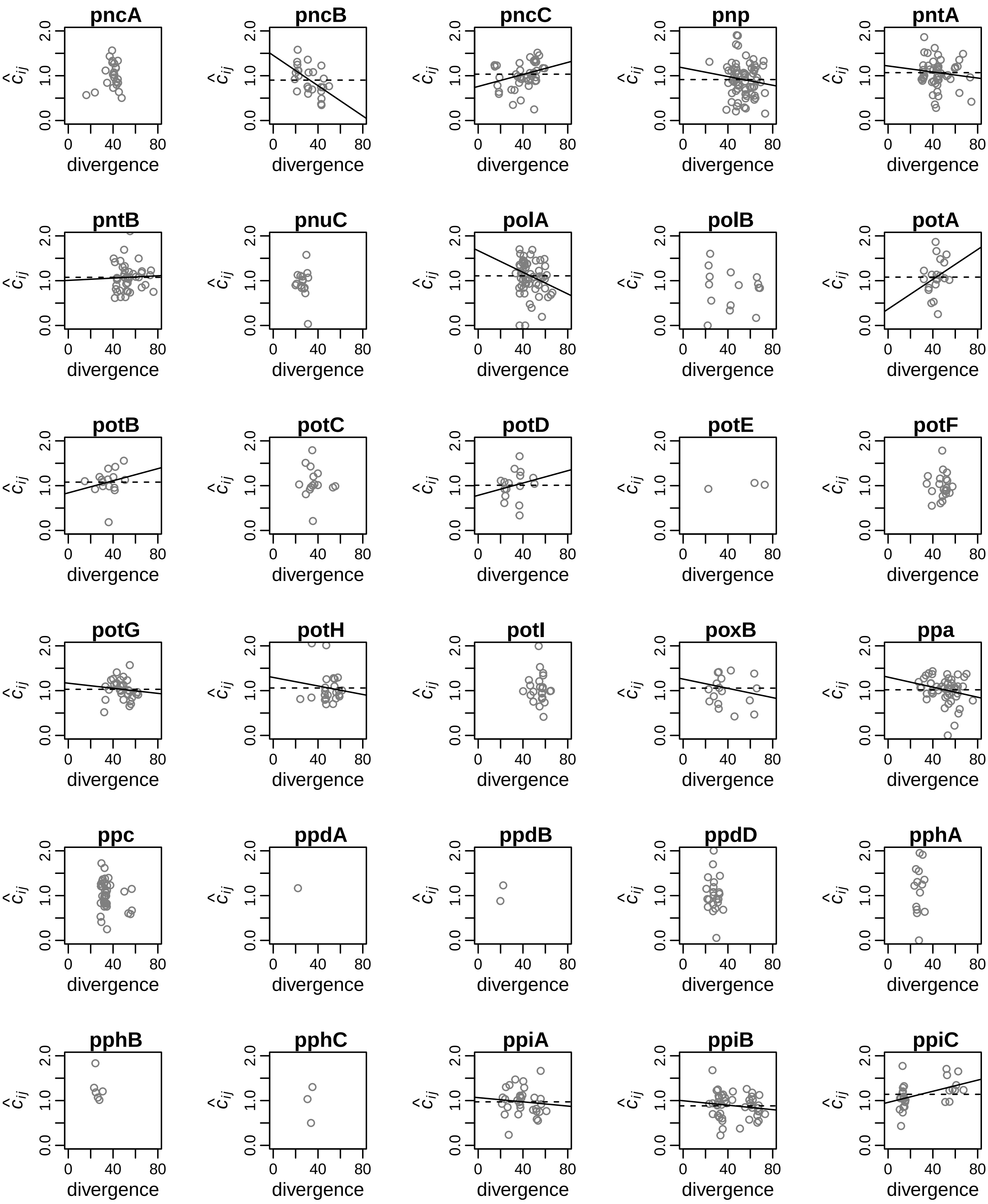

pncA
pncB
pncC
pnp
pntA
pntB
pnuC
polA
polB
potA
potB
potC
potD
potE
potF
potG
potH
potI
poxB
ppa
ppc
ppdA
ppdB
ppdD
pphA
pphB
pphC
ppiA
ppiB
ppiC
$\hat{c}_{ij}$
0.0
1.0
2.0
0
40
80
divergence

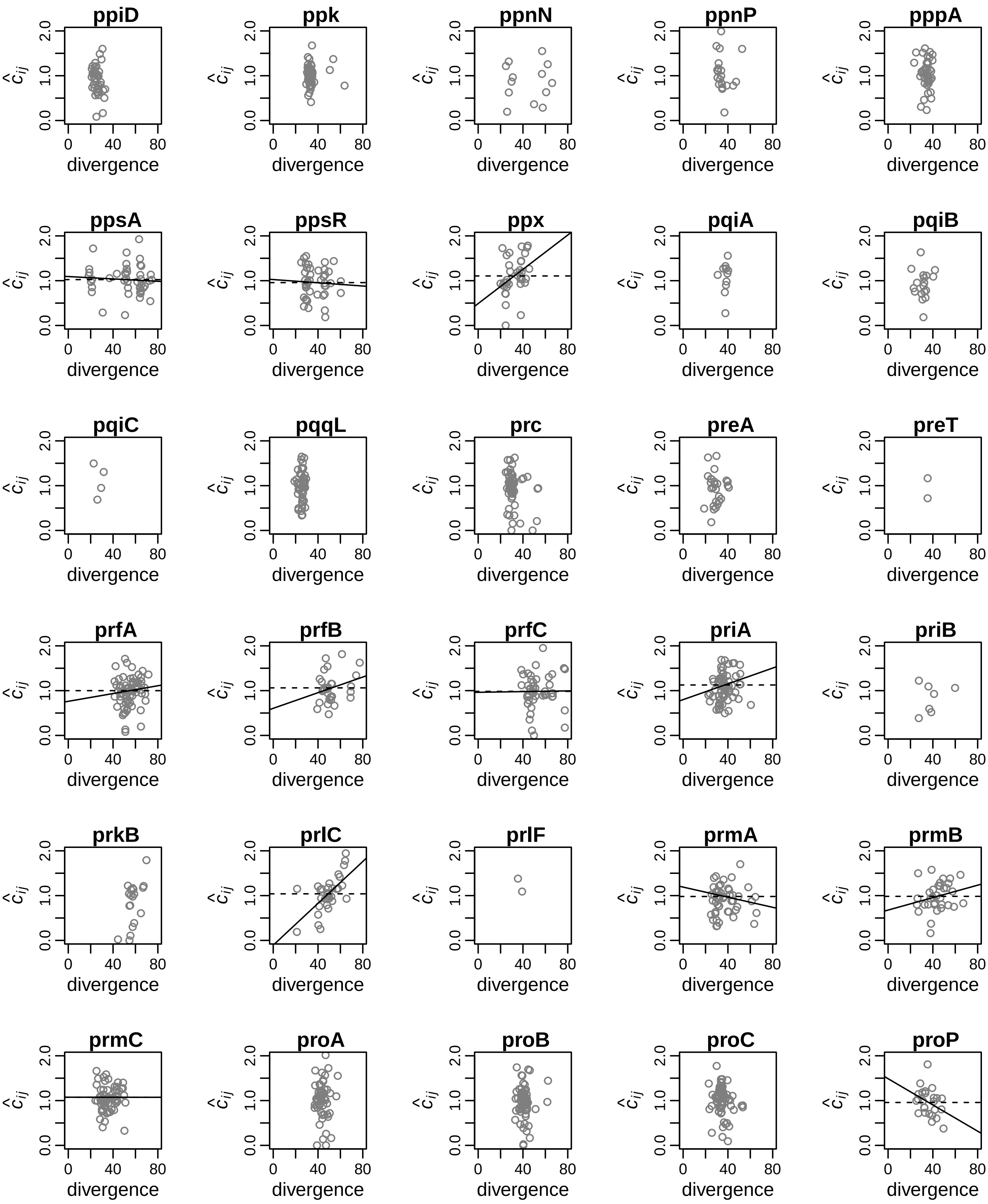

ppiD
ppk
ppnN
ppnP
pppA
ppsA
ppsR
ppx
pqiA
pqiB
pqiC
pqqL
prc
preA
preT
prfA
prfB
prfC
priA
priB
prkB
prlC
prlF
prmA
prmB
prmC
proA
proB
proC
proP
$\hat{c}_{ij}$
0.0
1.0
2.0
0
40
80
divergence

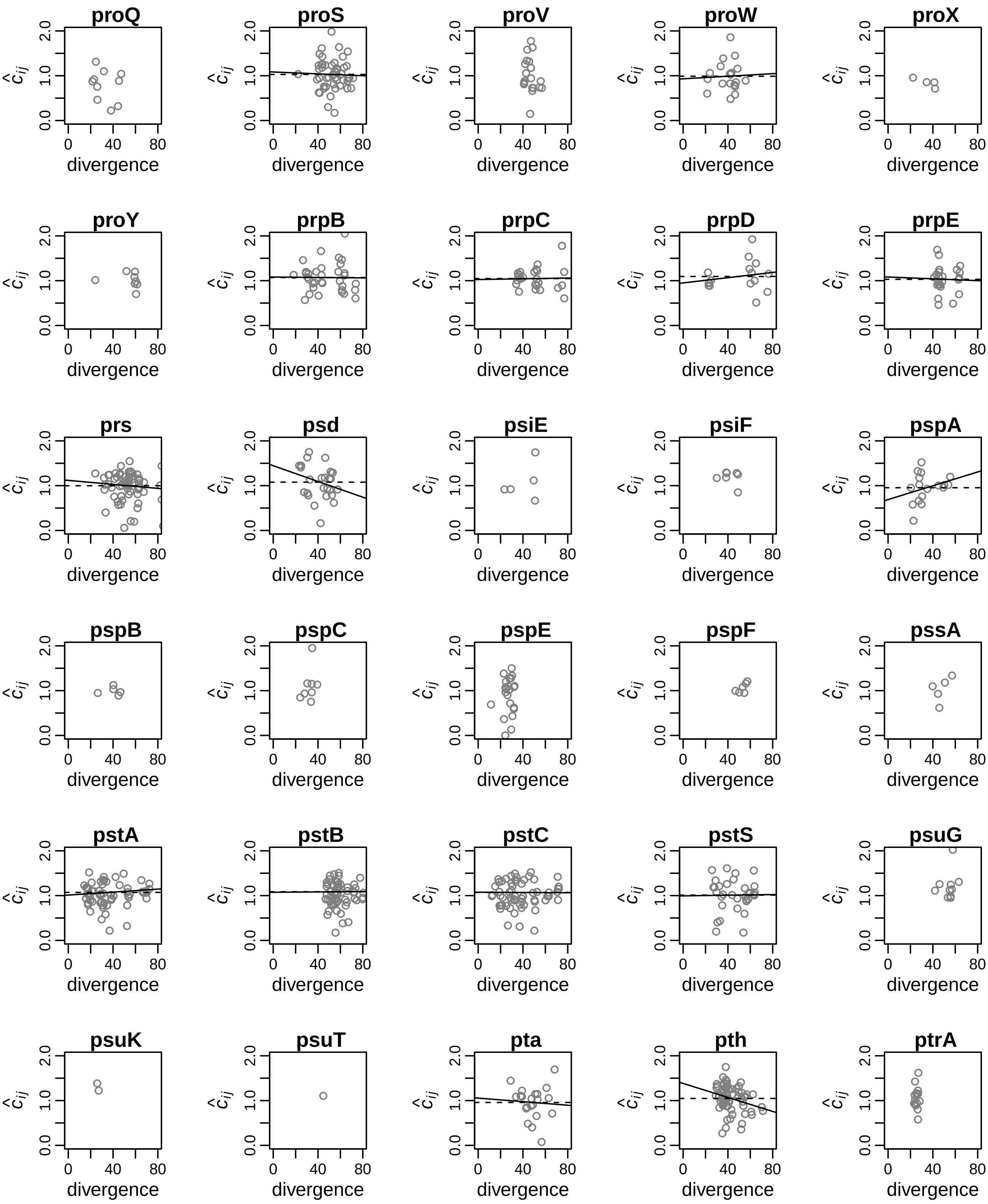
proQ
proS
proV
proW
proX
proY
prpB
prpC
prpD
prpE
prs
psd
psiE
psiF
pspA
pspB
pspC
pspE
pspF
pssA
pstA
pstB
pstC
pstS
psuG
psuK
psuT
pta
pth
ptrA
divergence
$\hat{c}_{ij}$
0
40
80
0.0
1.0
2.0

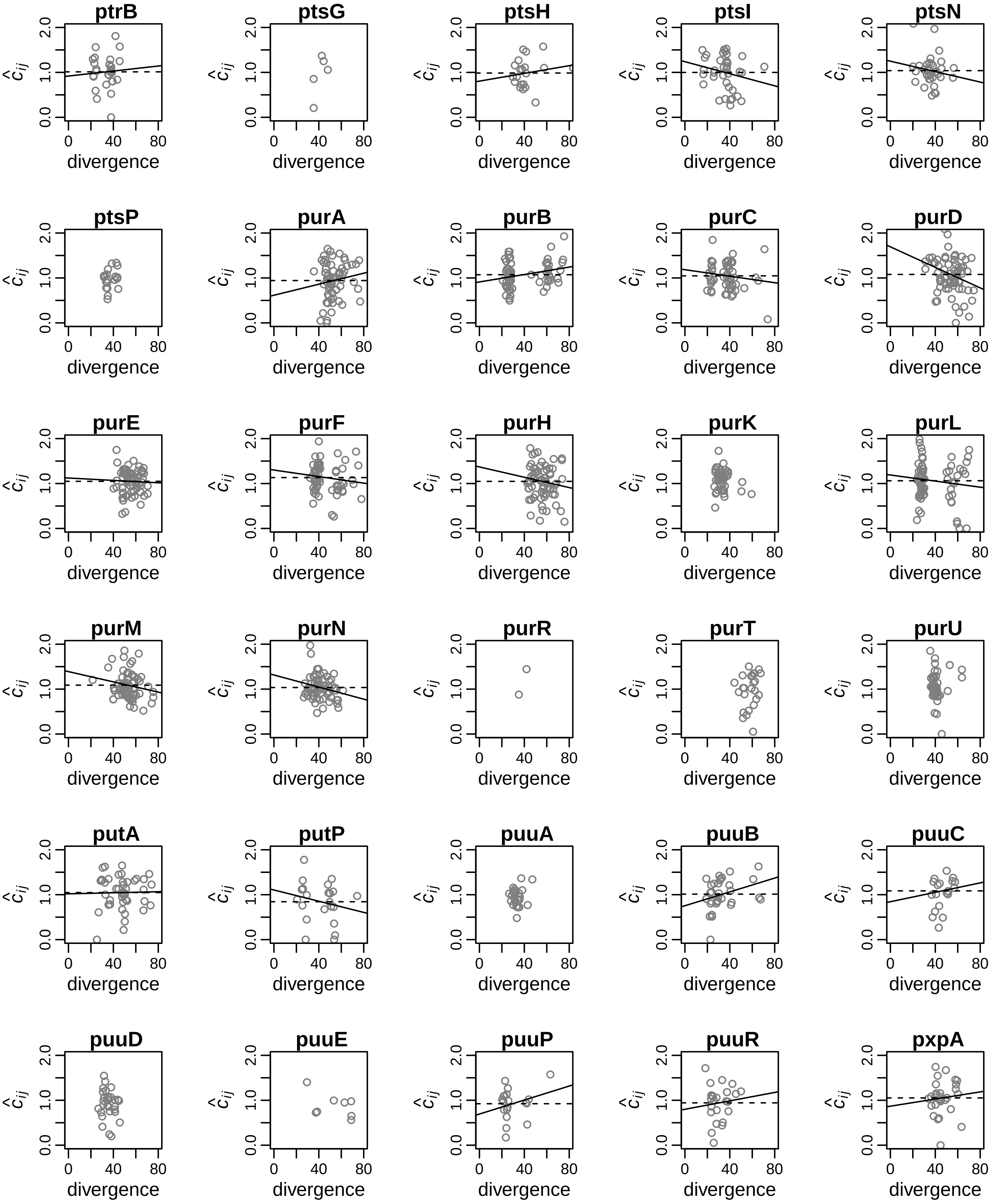
ptrB
ptsG
ptsH
ptsI
ptsN
ptsP
purA
purB
purC
purD
purE
purF
purH
purK
purL
purM
purN
purR
purT
purU
putA
putP
puuA
puuB
puuC
puuD
puuE
puuP
puuR
pxpA
$\hat{c}_{ij}$
0.0
1.0
2.0
0
40
80
divergence

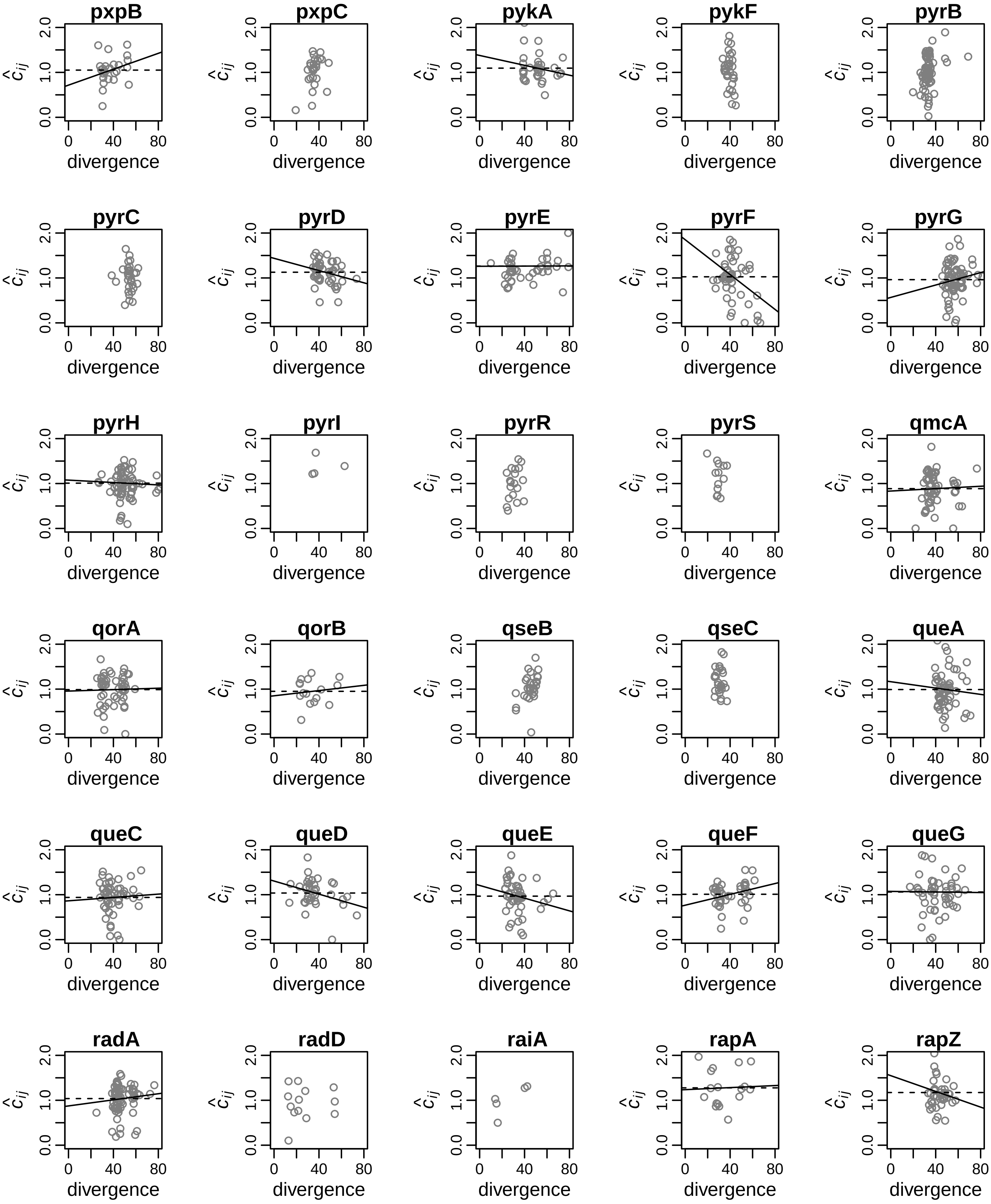
pxpB
pxpC
pykA
pykF
pyrB
pyrC
pyrD
pyrE
pyrF
pyrG
pyrH
pyrI
pyrR
pyrS
qmcA
qorA
qorB
qseB
qseC
queA
queC
queD
queE
queF
queG
radA
radD
raiA
rapA
rapZ
$\hat{c}_{ij}$
0.0
1.0
2.0
0
40
80
divergence

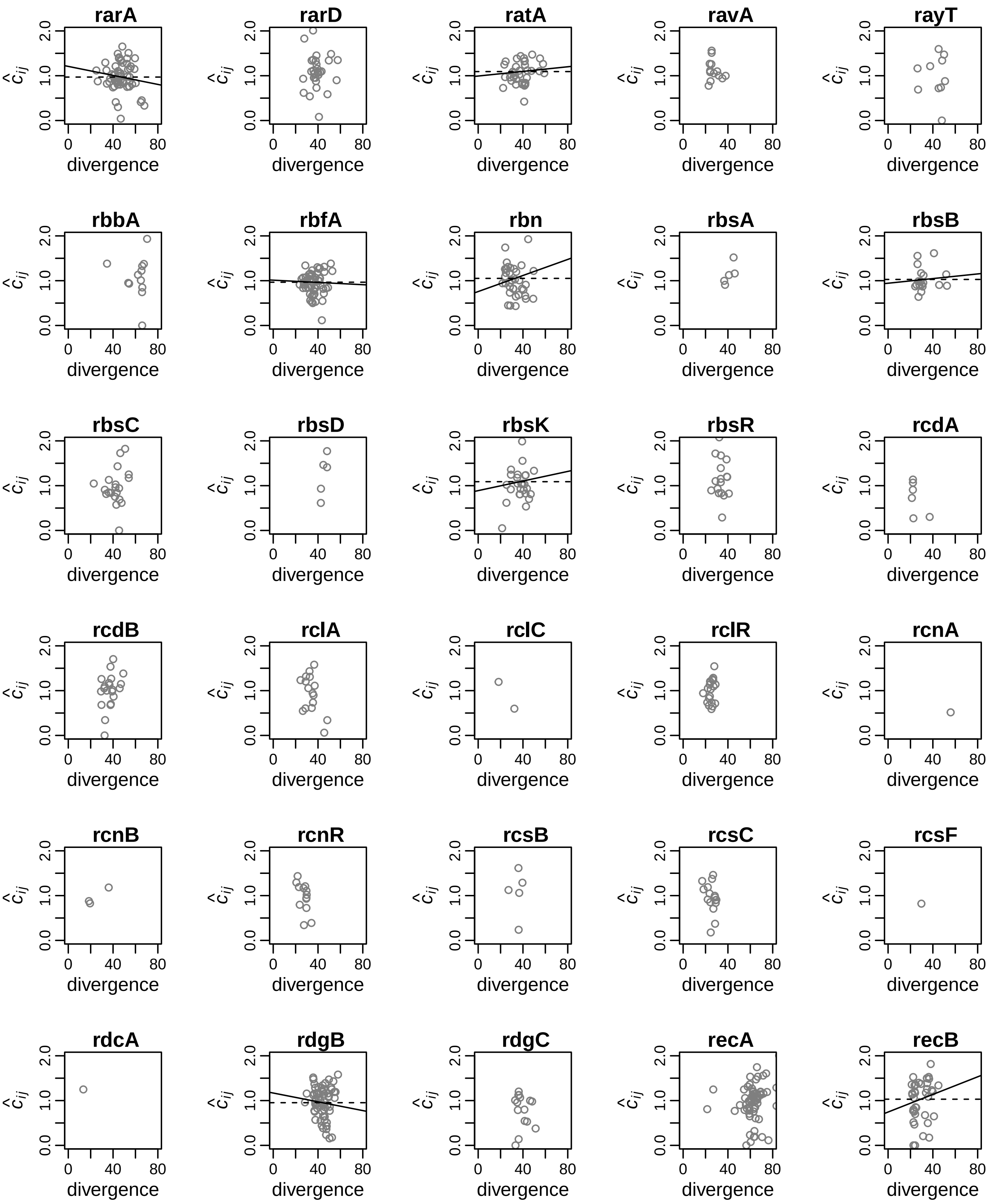
rarA
rarD
ratA
ravA
rayT
rbbA
rbfA
rbn
rbsA
rbsB
rbsC
rbsD
rbsK
rbsR
rcdA
rcdB
rclA
rclC
rclR
rcnA
rcnB
rcnR
rcsB
rcsC
rcsF
rdcA
rdgB
rdgC
recA
recB
$\hat{c}_{ij}$
0.0
1.0
2.0
0
40
80
divergence

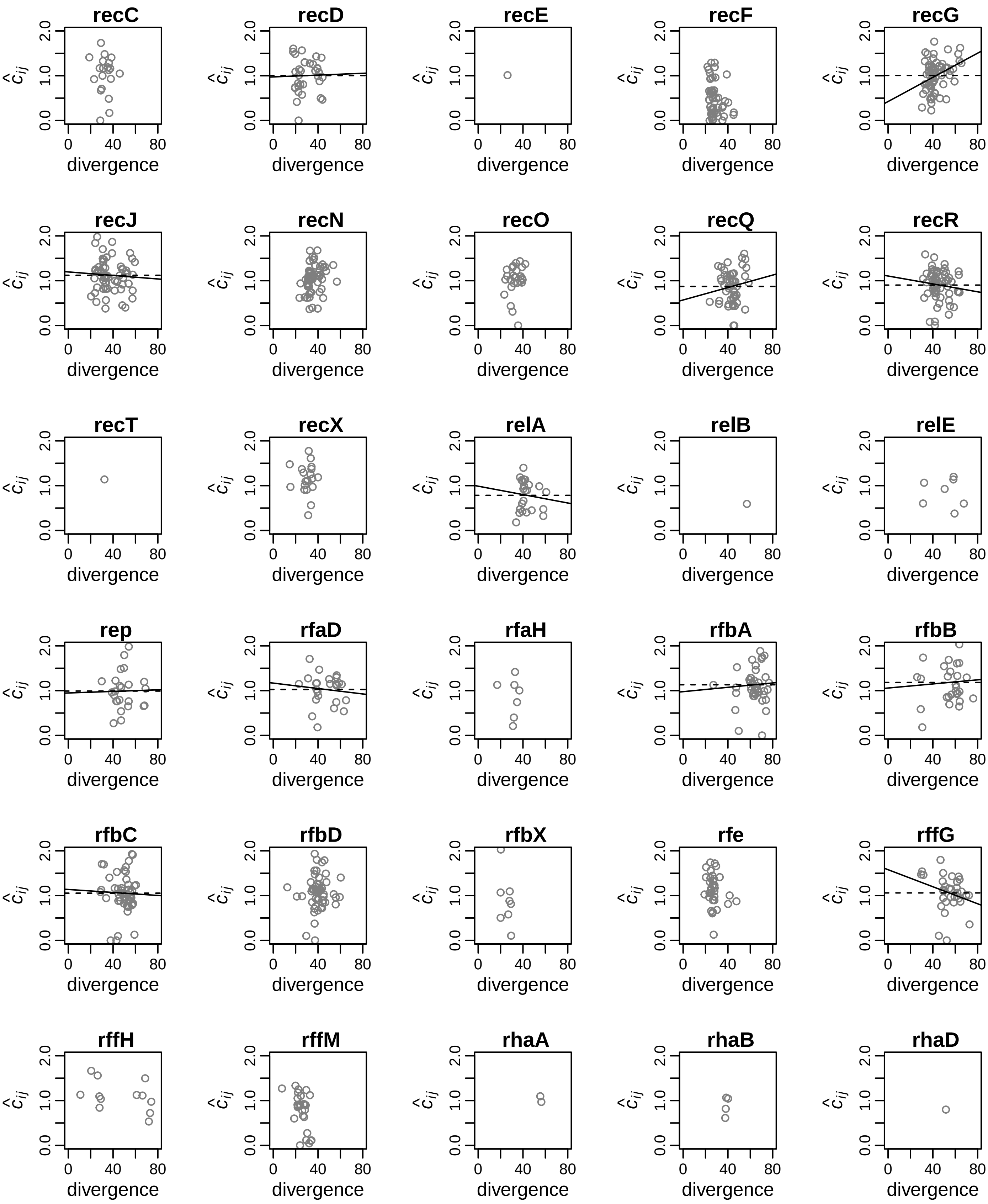
recC
recD
recE
recF
recG
recJ
recN
recO
recQ
recR
recT
recX
relA
relB
relE
rep
rfaD
rfaH
rfbA
rfbB
rfbC
rfbD
rfbX
rfe
rffG
rffH
rffM
rhaA
rhaB
rhaD
$\hat{c}_{ij}$
divergence
0.0
1.0
2.0
0
40
80

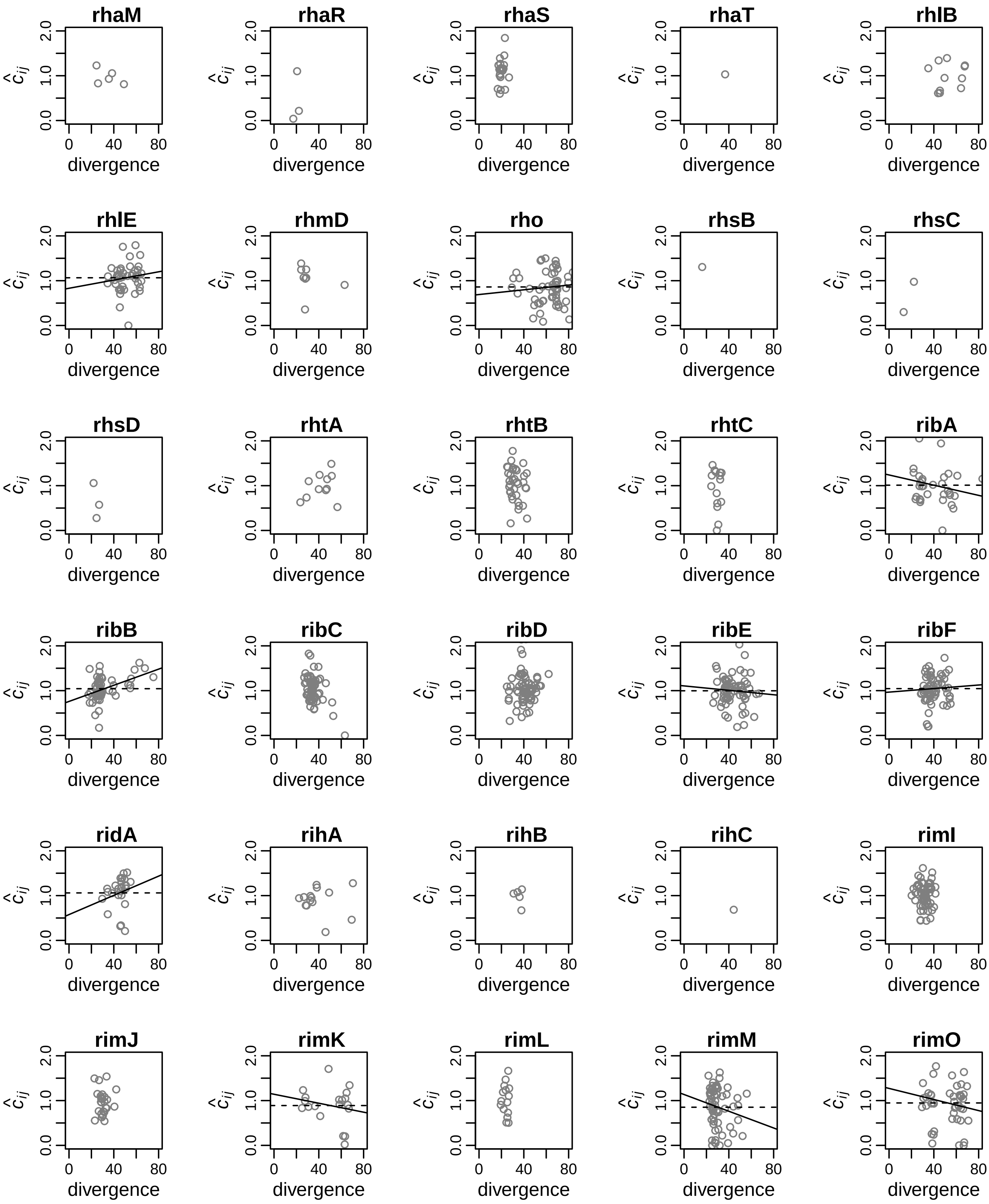

rhaM
rhaR
rhaS
rhaT
rhlB
rhlE
rhmD
rho
rhsB
rhsC
rhsD
rhtA
rhtB
rhtC
ribA
ribB
ribC
ribD
ribE
ribF
ridA
rihA
rihB
rihC
rimI
rimJ
rimK
rimL
rimM
rimO
$\hat{c}_{ij}$
0.0
1.0
2.0
0
40
80
divergence

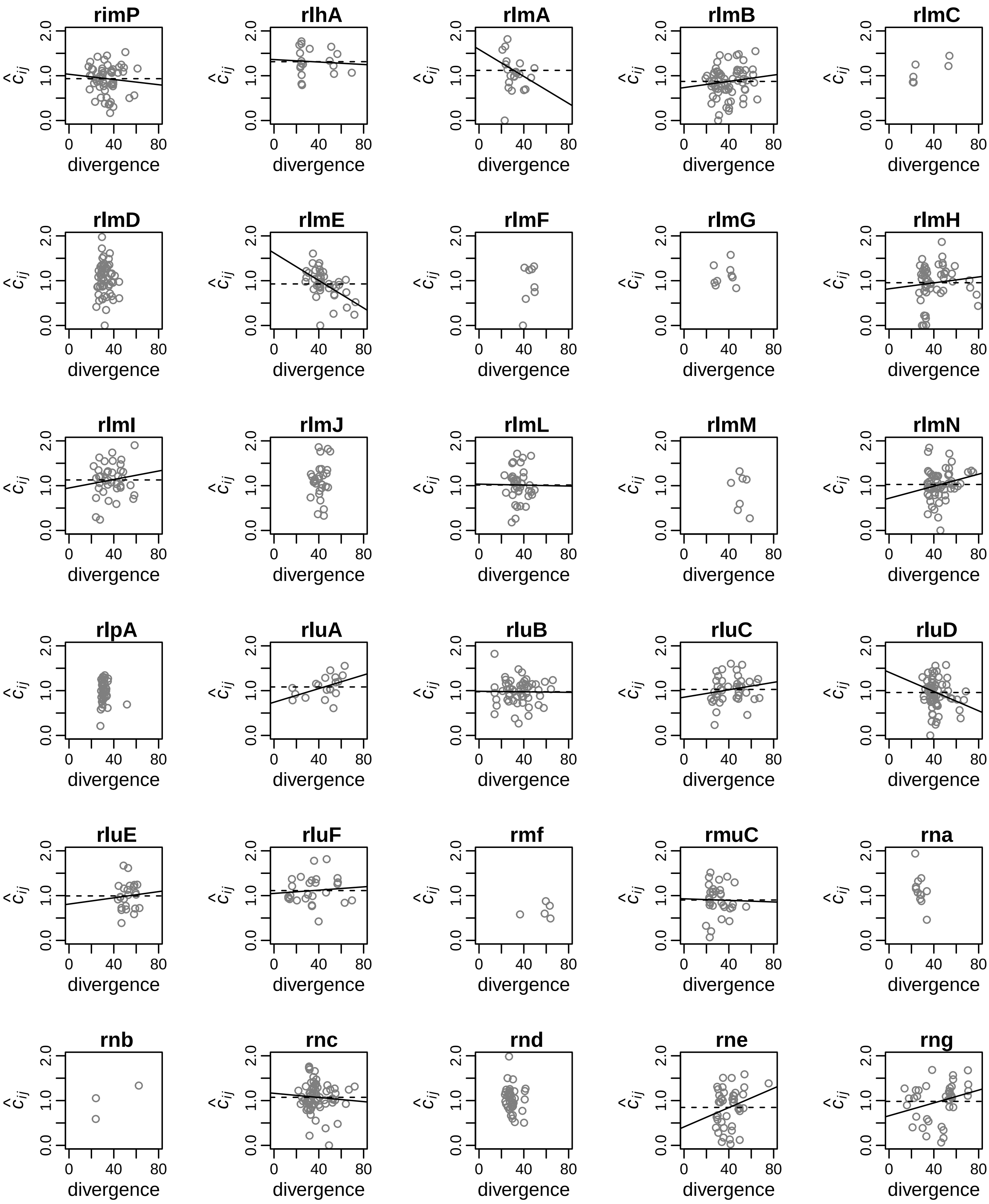
rimP
rlhA
rlmA
rlmB
rlmC
rlmD
rlmE
rlmF
rlmG
rlmH
rlmI
rlmJ
rlmL
rlmM
rlmN
rlpA
rluA
rluB
rluC
rluD
rluE
rluF
rmf
rmuC
rna
rnb
rnc
rnd
rne
rng
$\hat{c}_{ij}$
0.0
1.0
2.0
0
40
80
divergence

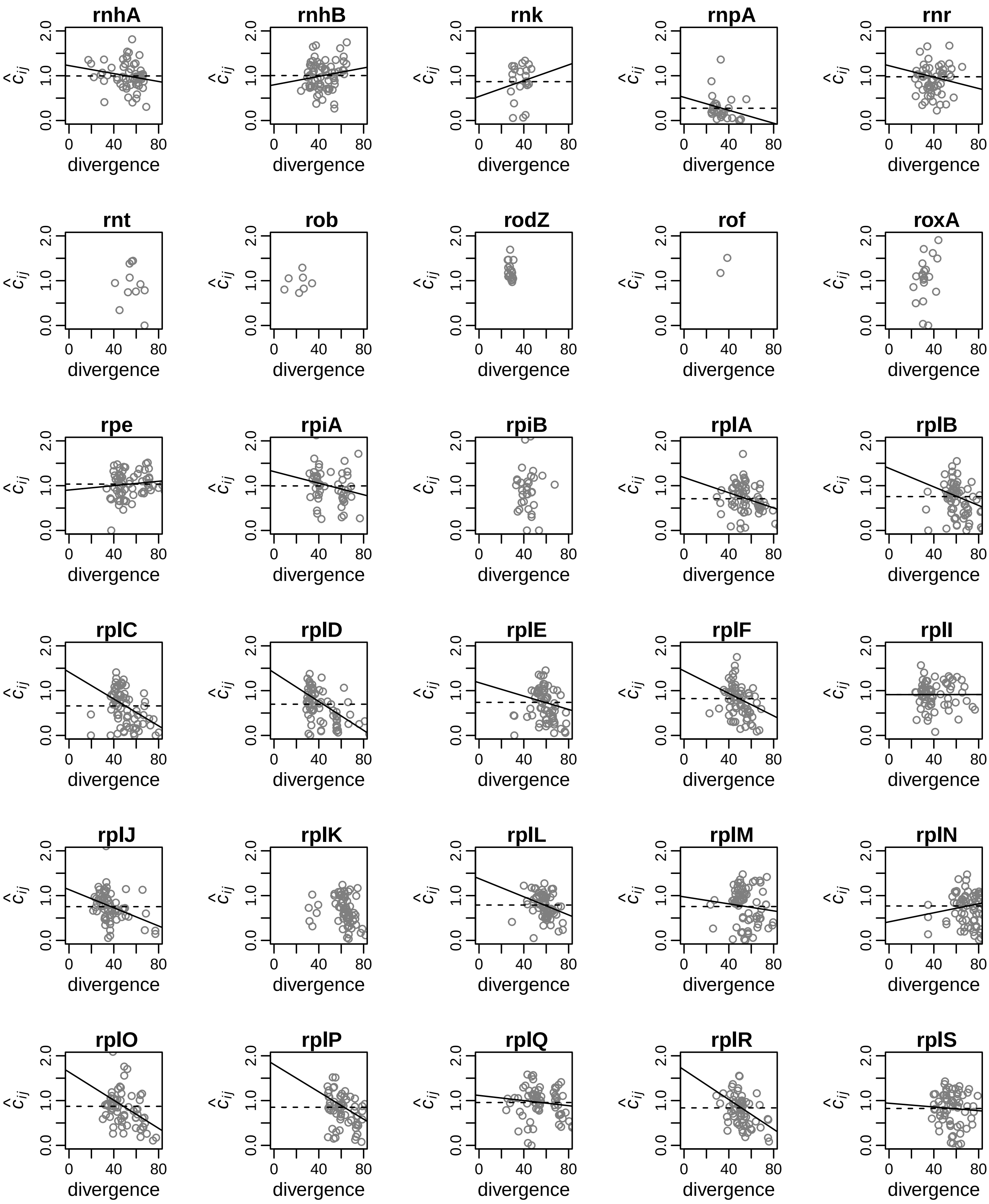

rnhA
rnhB
rnk
rnpA
rnr
rnt
rob
rodZ
rof
roxA
rpe
rpiA
rpiB
rplA
rplB
rplC
rplD
rplE
rplF
rplI
rplJ
rplK
rplL
rplM
rplN
rplO
rplP
rplQ
rplR
rplS
$\hat{c}_{ij}$
0.0
1.0
2.0
0
40
80
divergence

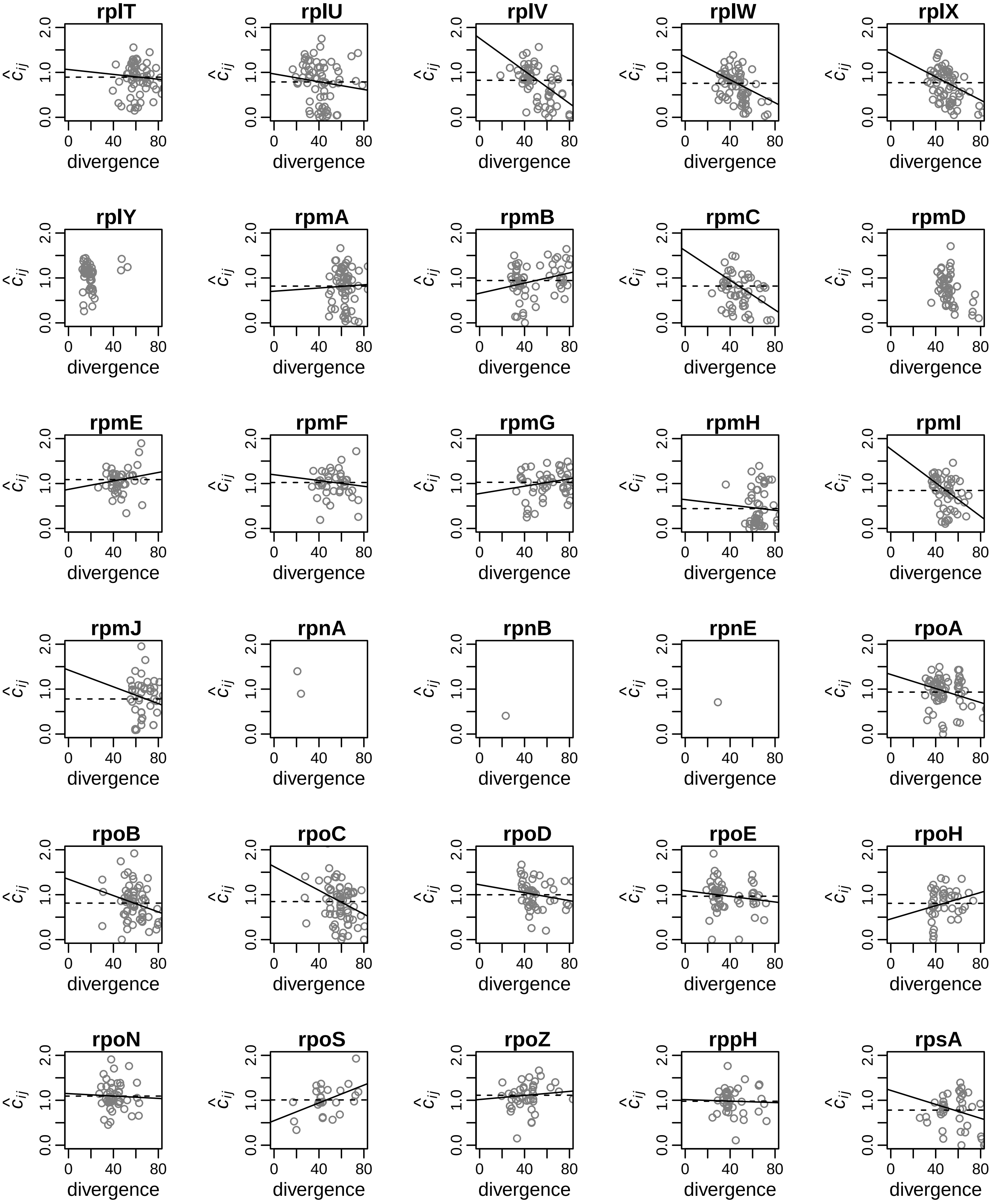

rplT
rplU
rplV
rplW
rplX
rplY
rpmA
rpmB
rpmC
rpmD
rpmE
rpmF
rpmG
rpmH
rpmI
rpmJ
rpnA
rpnB
rpnE
rpoA
rpoB
rpoC
rpoD
rpoE
rpoH
rpoN
rpoS
rpoZ
rppH
rpsA
$\hat{c}_{ij}$
divergence
0.0 1.0 2.0
0 40 80

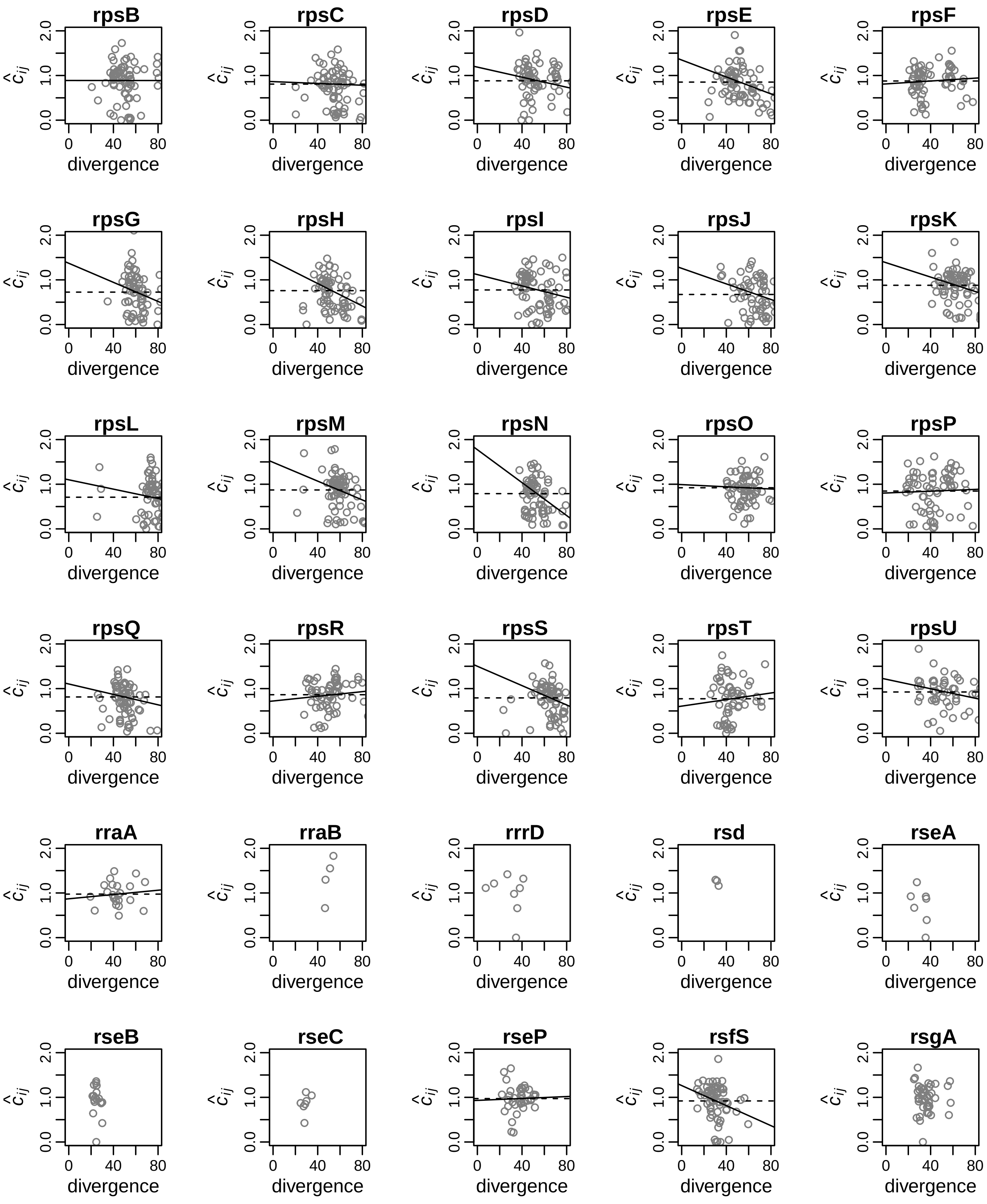
rpsB
rpsC
rpsD
rpsE
rpsF
rpsG
rpsH
rpsI
rpsJ
rpsK
rpsL
rpsM
rpsN
rpsO
rpsP
rpsQ
rpsR
rpsS
rpsT
rpsU
rraA
rraB
rrrD
rsd
rseA
rseB
rseC
rseP
rsfS
rsgA
$\hat{c}_{ij}$
0.0
1.0
2.0
0
40
80
divergence

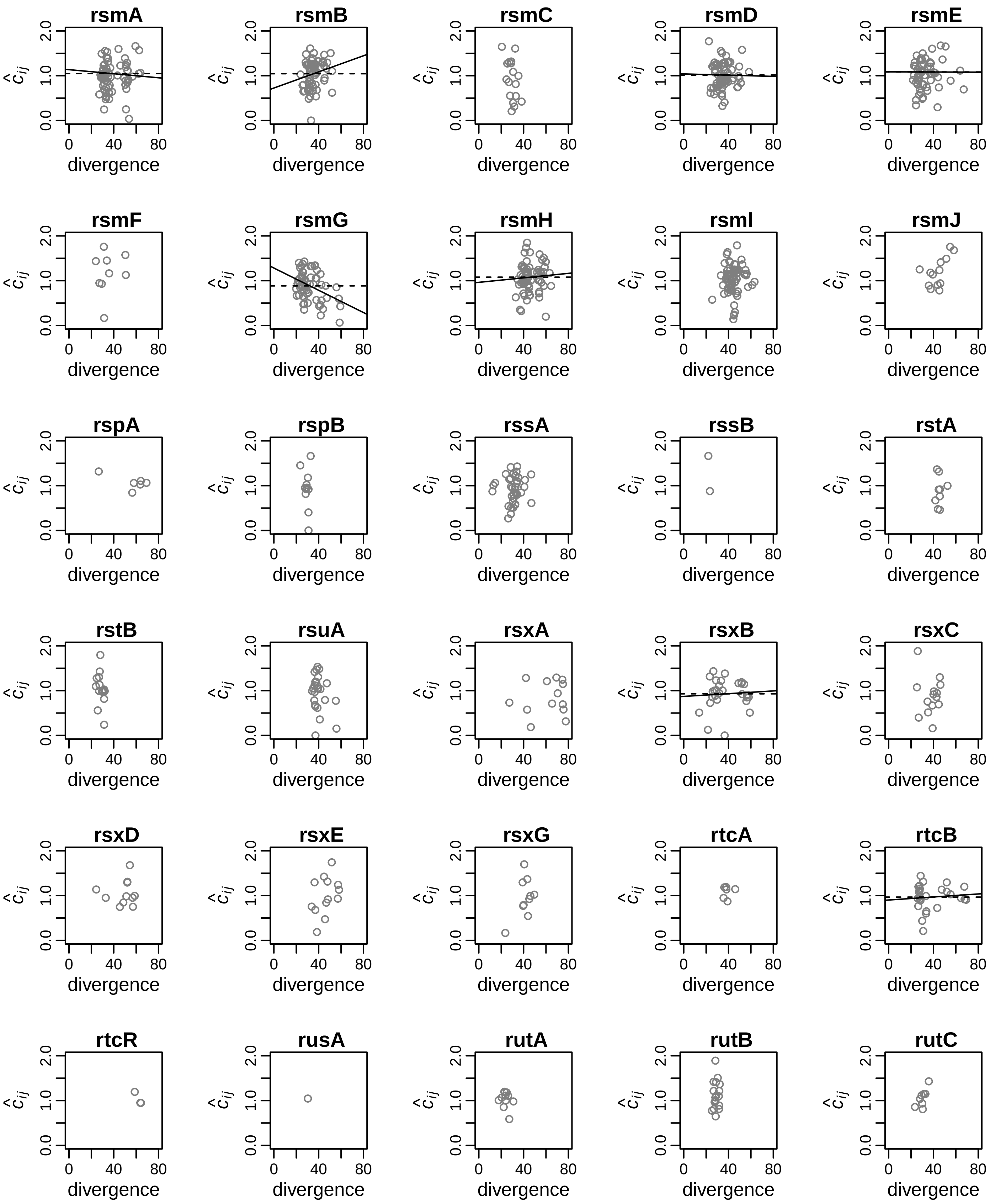

rsmA
rsmB
rsmC
rsmD
rsmE
rsmF
rsmG
rsmH
rsmI
rsmJ
rspA
rspB
rssA
rssB
rstA
rstB
rsuA
rsxA
rsxB
rsxC
rsxD
rsxE
rsxG
rtcA
rtcB
rtcR
rusA
rutA
rutB
rutC
$\hat{c}_{ij}$
divergence
0.0
1.0
2.0
0
40
80

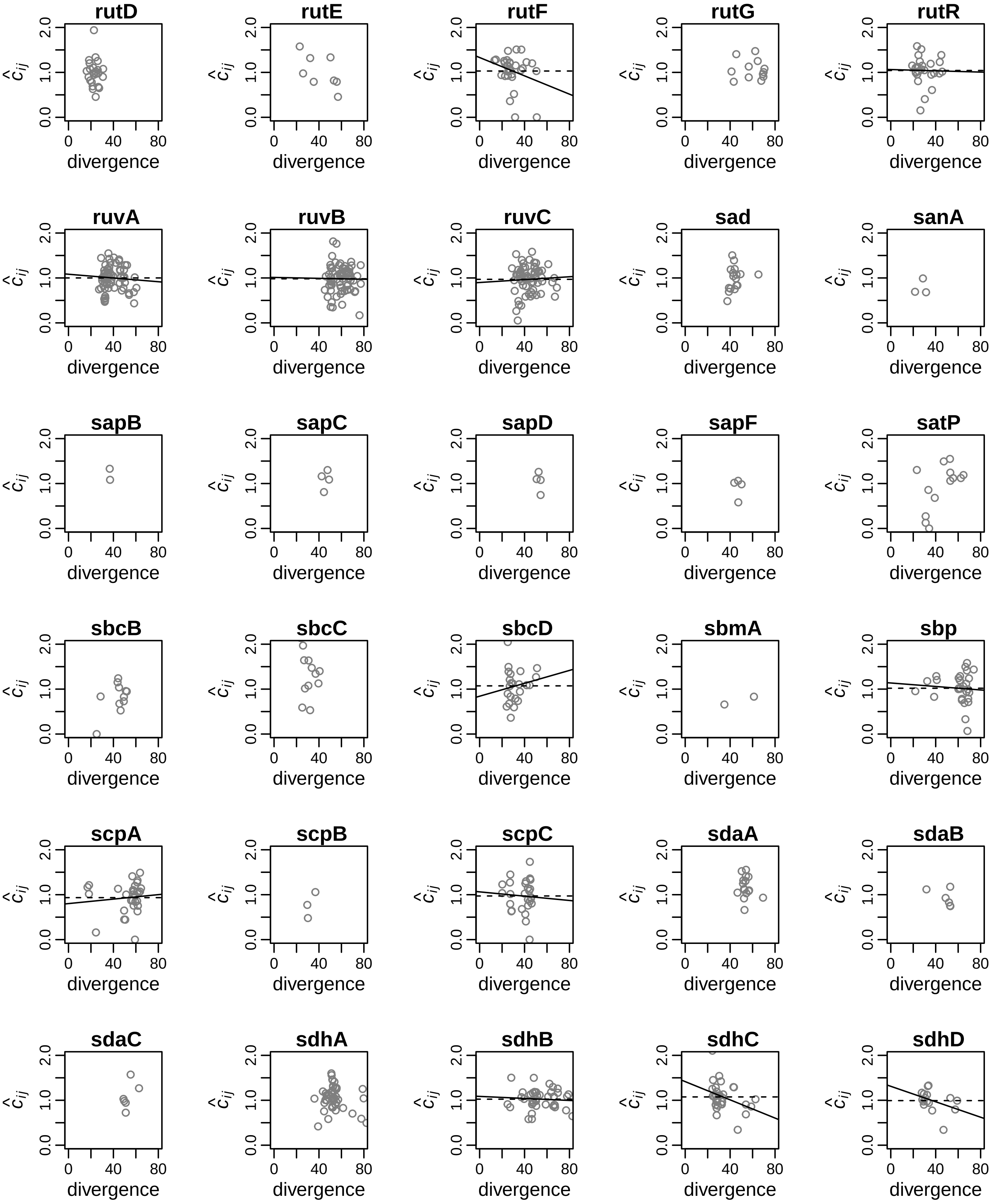
rutD
rutE
rutF
rutG
rutR
ruvA
ruvB
ruvC
sad
sanA
sapB
sapC
sapD
sapF
satP
sbcB
sbcC
sbcD
sbmA
sbp
scpA
scpB
scpC
sdaA
sdaB
sdaC
sdhA
sdhB
sdhC
sdhD
divergence
$\hat{c}_{ij}$
0
40
80
0.0
1.0
2.0

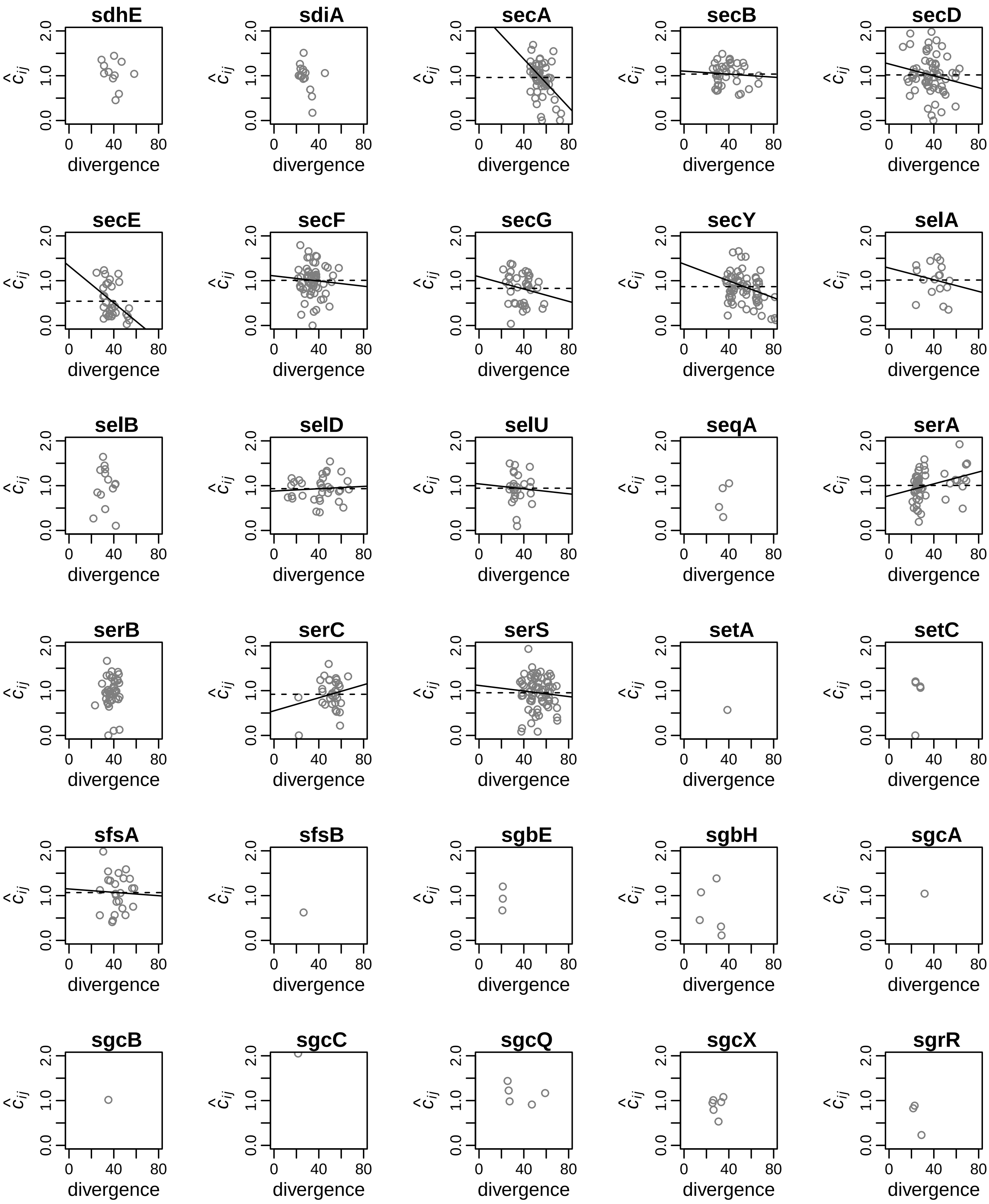
sdhE
sdiA
secA
secB
secD
secE
secF
secG
secY
selA
selB
selD
selU
seqA
serA
serB
serC
serS
setA
setC
sfsA
sfsB
sgbE
sgbH
sgcA
sgcB
sgcC
sgcQ
sgcX
sgrR
$\hat{c}_{ij}$
divergence
0.0
1.0
2.0
0
40
80

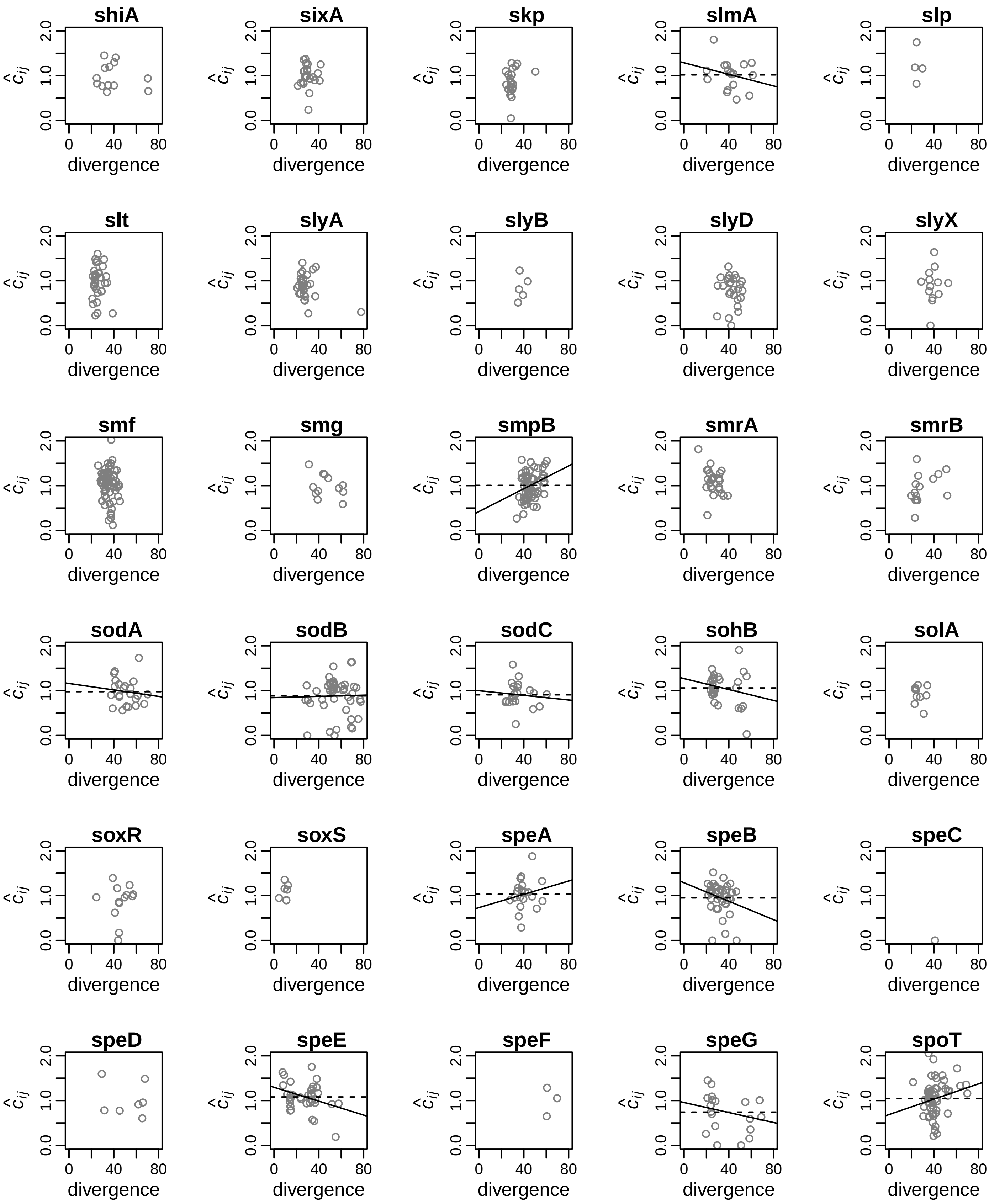

shiA
sixA
skp
slmA
slp
slt
slyA
slyB
slyD
slyX
smf
smg
smpB
smrA
smrB
sodA
sodB
sodC
sohB
solA
soxR
soxS
speA
speB
speC
speD
speE
speF
speG
spoT
$\hat{c}_{ij}$
0.0
1.0
2.0
0
40
80
divergence

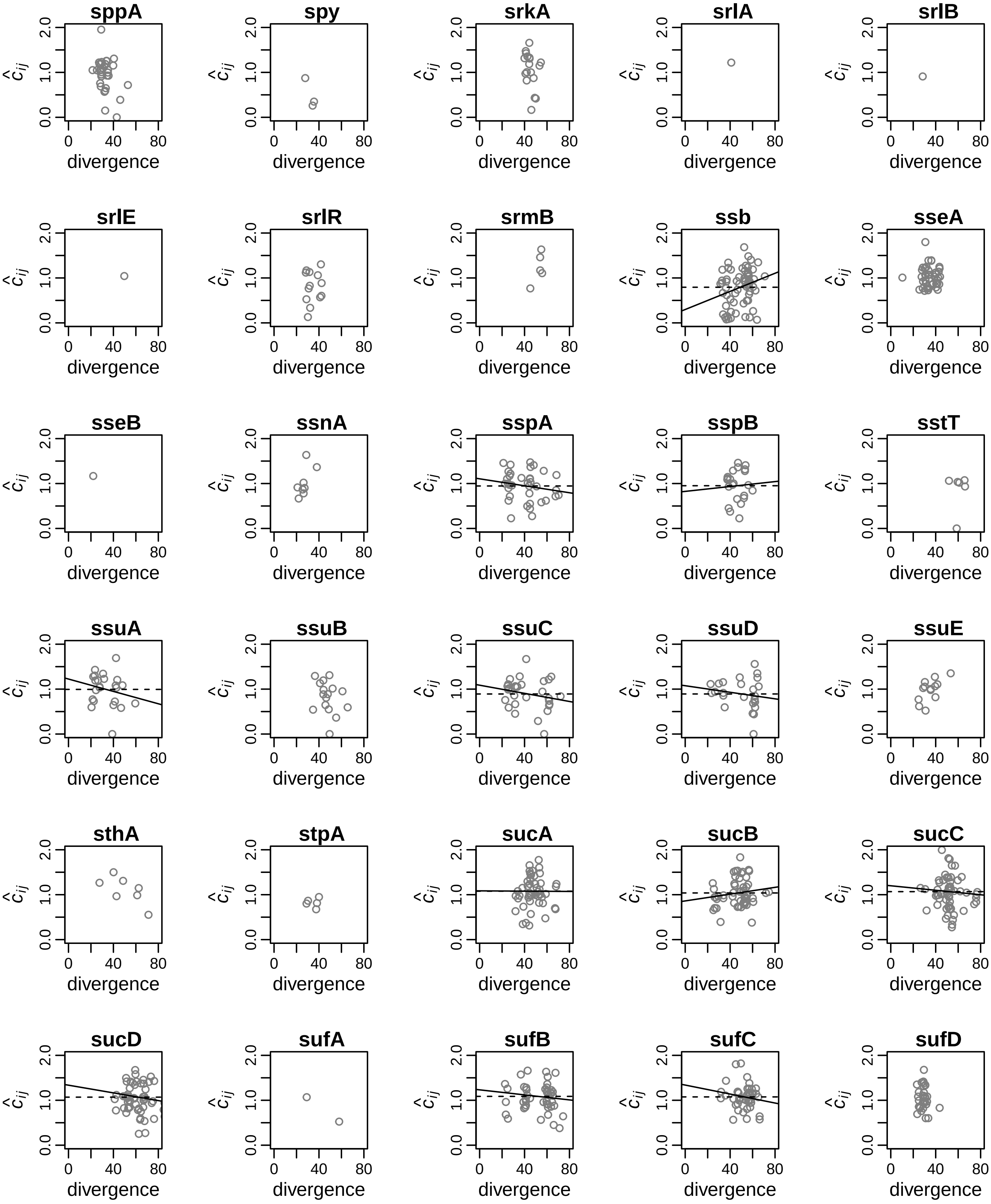
sppA
spy
srkA
srlA
srlB
srlE
srlR
srmB
ssb
sseA
sseB
ssnA
sspA
sspB
sstT
ssuA
ssuB
ssuC
ssuD
ssuE
sthA
stpA
sucA
sucB
sucC
sucD
sufA
sufB
sufC
sufD
$\hat{c}_{ij}$
0.0
1.0
2.0
0
40
80
divergence

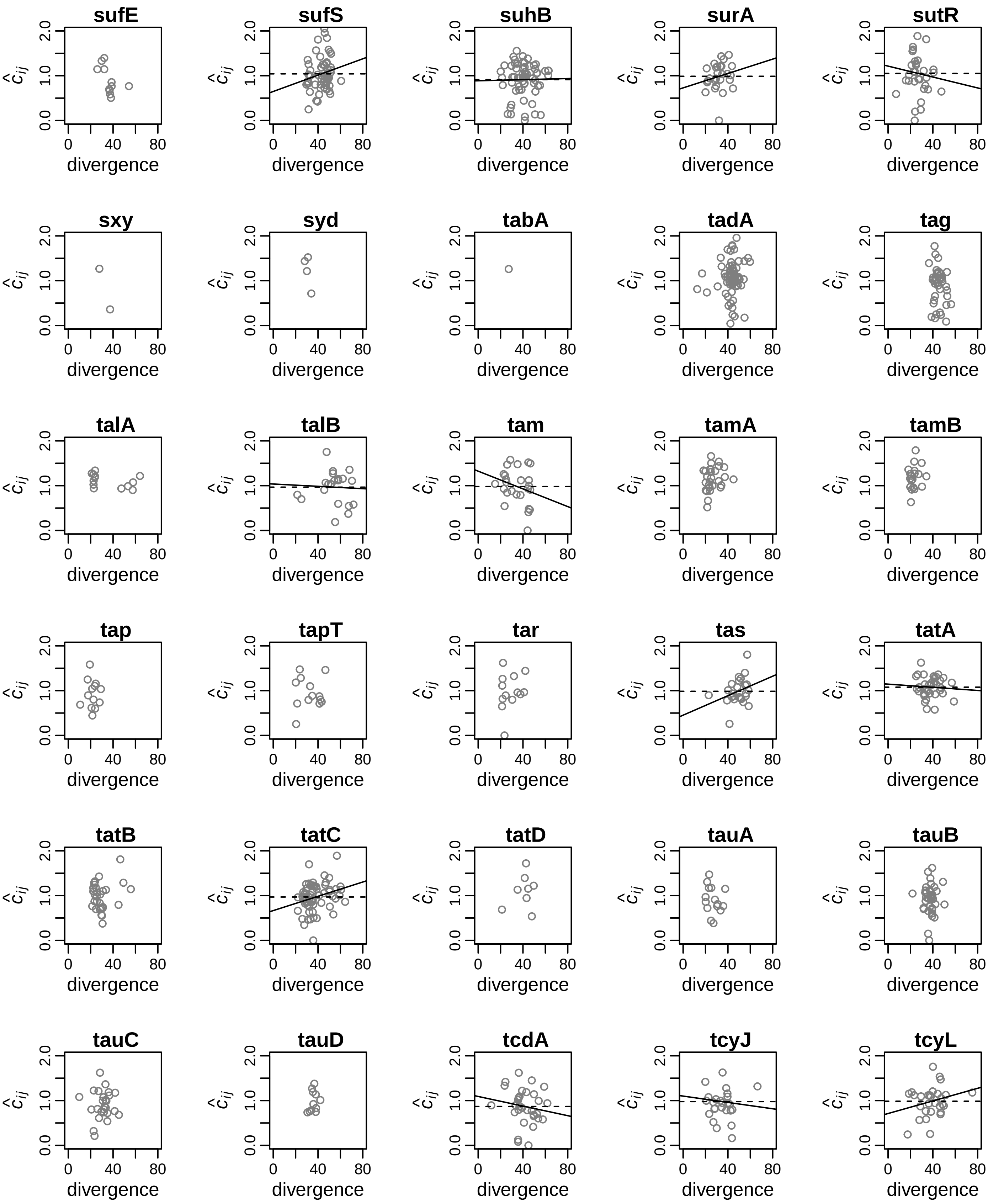
sufE
sufS
suhB
surA
sutR
sxy
syd
tabA
tadA
tag
talA
talB
tam
tamA
tamB
tap
tapT
tar
tas
tatA
tatB
tatC
tatD
tauA
tauB
tauC
tauD
tcdA
tcyJ
tcyL
$\hat{c}_{ij}$
0.0
1.0
2.0
0
40
80
divergence

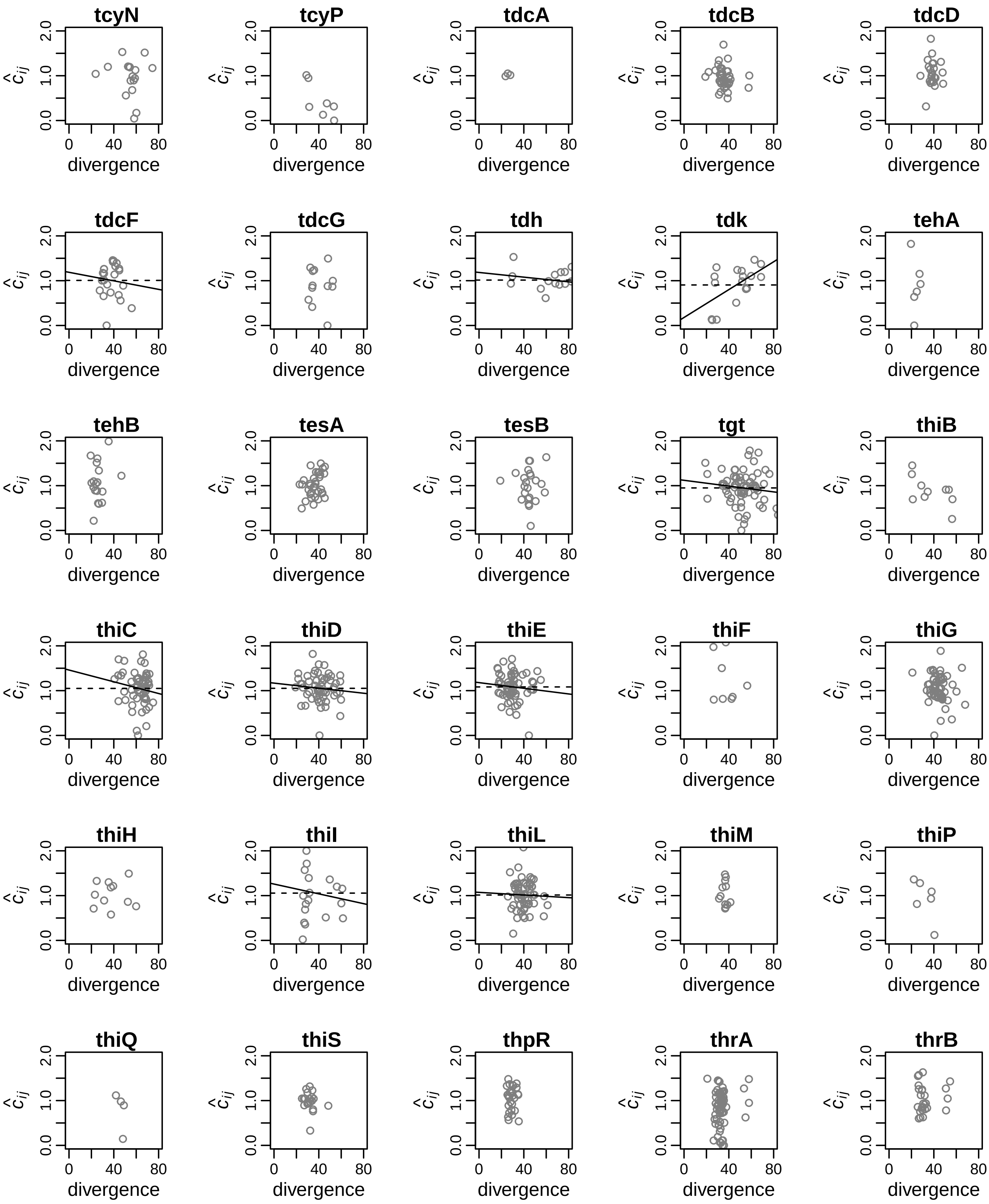
tcyN
tcyP
tdcA
tdcB
tdcD
tdcF
tdcG
tdh
tdk
tehA
tehB
tesA
tesB
tgt
thiB
thiC
thiD
thiE
thiF
thiG
thiH
thiI
thiL
thiM
thiP
thiQ
thiS
thpR
thrA
thrB
$\hat{c}_{ij}$
0.0
1.0
2.0
0
40
80
divergence

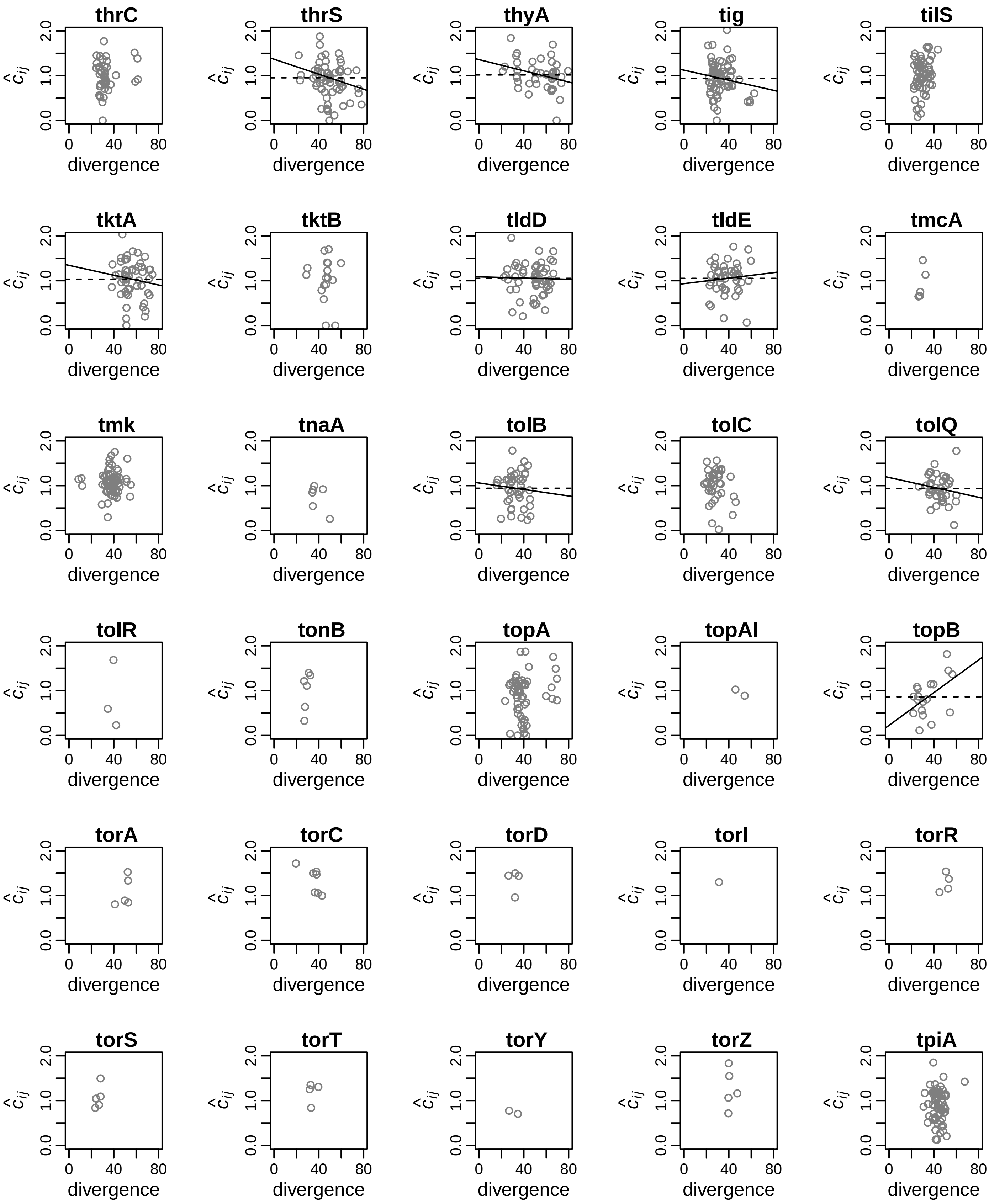
thrC
thrS
thyA
tig
tilS
tktA
tktB
tldD
tldE
tmcA
tmk
tnaA
tolB
tolC
tolQ
tolR
tonB
topA
topAI
topB
torA
torC
torD
torI
torR
torS
torT
torY
torZ
tpiA
$\hat{c}_{ij}$
0.0
1.0
2.0
0
40
80
divergence

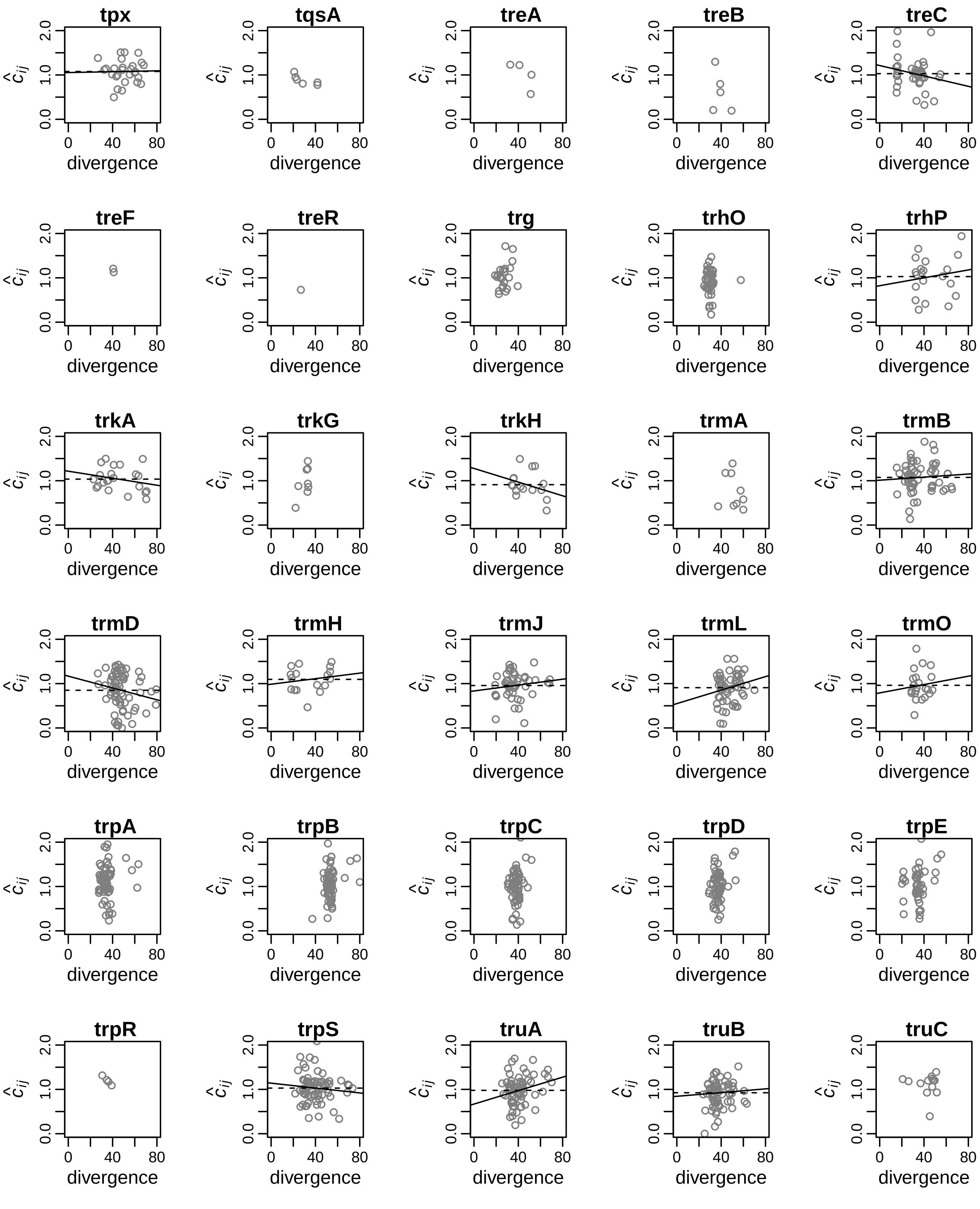
tpx
tqsA
treA
treB
treC
treF
treR
trg
trhO
trhP
trkA
trkG
trkH
trmA
trmB
trmD
trmH
trmJ
trmL
trmO
trpA
trpB
trpC
trpD
trpE
trpR
trpS
truA
truB
truC
$\hat{c}_{ij}$
0.0 1.0 2.0
0 40 80
divergence

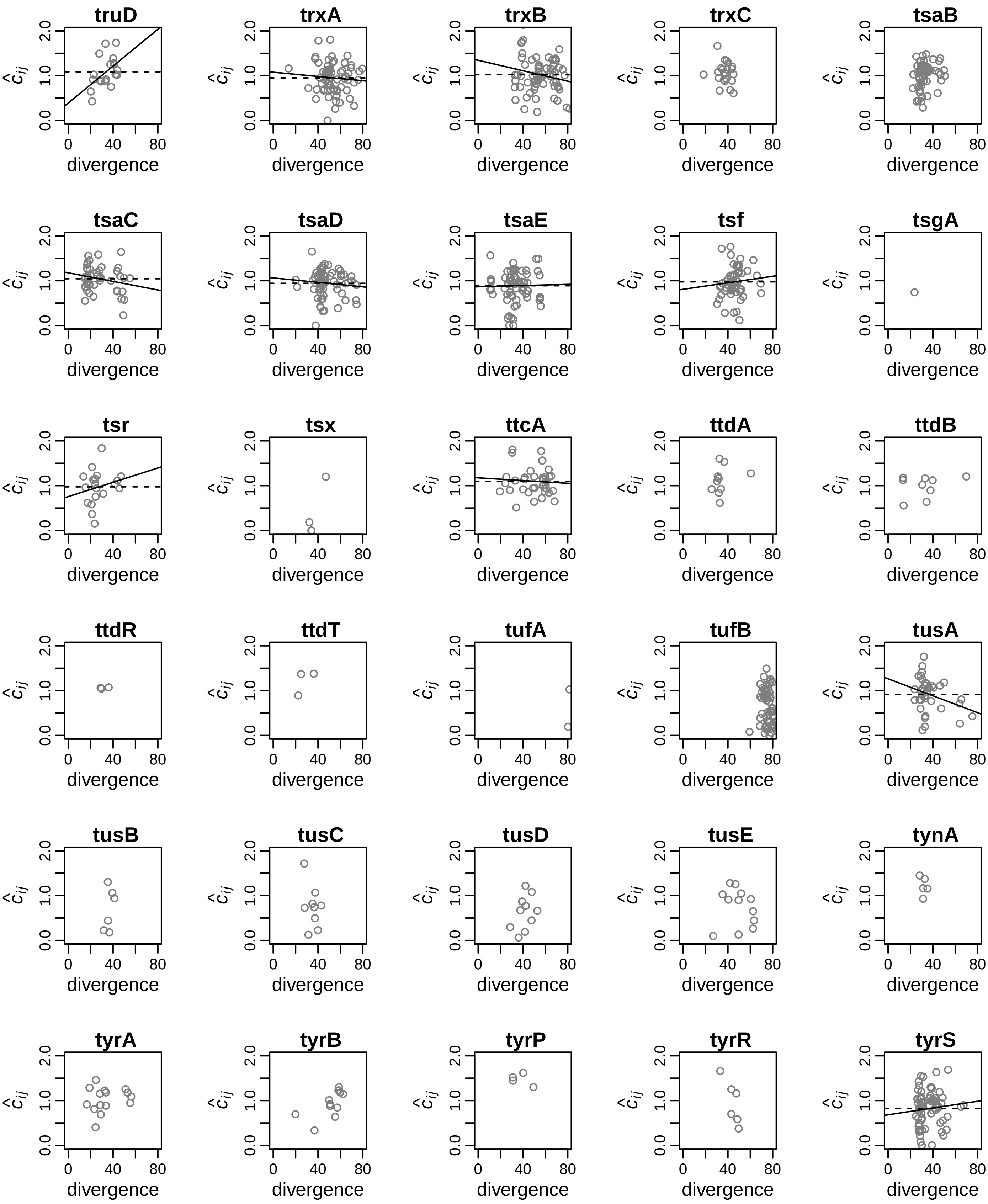
truD
trxA
trxB
trxC
tsaB
tsaC
tsaD
tsaE
tsf
tsgA
tsr
tsx
ttcA
ttdA
ttdB
ttdR
ttdT
tufA
tufB
tusA
tusB
tusC
tusD
tusE
tynA
tyrA
tyrB
tyrP
tyrR
tyrS
$\hat{c}_{ij}$
0.0
1.0
2.0
0
40
80
divergence

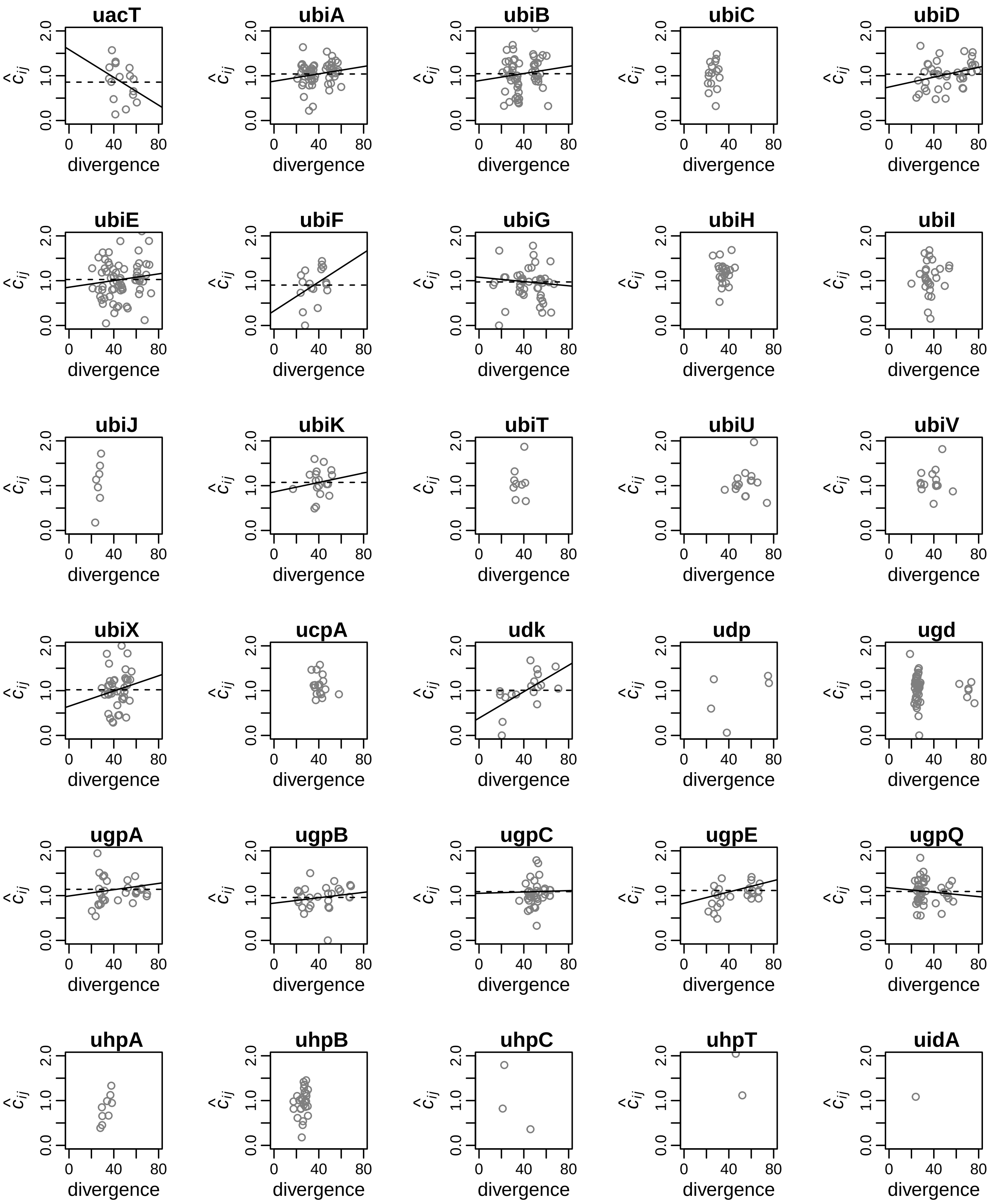
uacT
ubiA
ubiB
ubiC
ubiD
ubiE
ubiF
ubiG
ubiH
ubiI
ubiJ
ubiK
ubiT
ubiU
ubiV
ubiX
ucpA
udk
udp
ugd
ugpA
ugpB
ugpC
ugpE
ugpQ
uhpA
uhpB
uhpC
uhpT
uidA
$\hat{c}_{ij}$
0.0
1.0
2.0
0
40
80
divergence

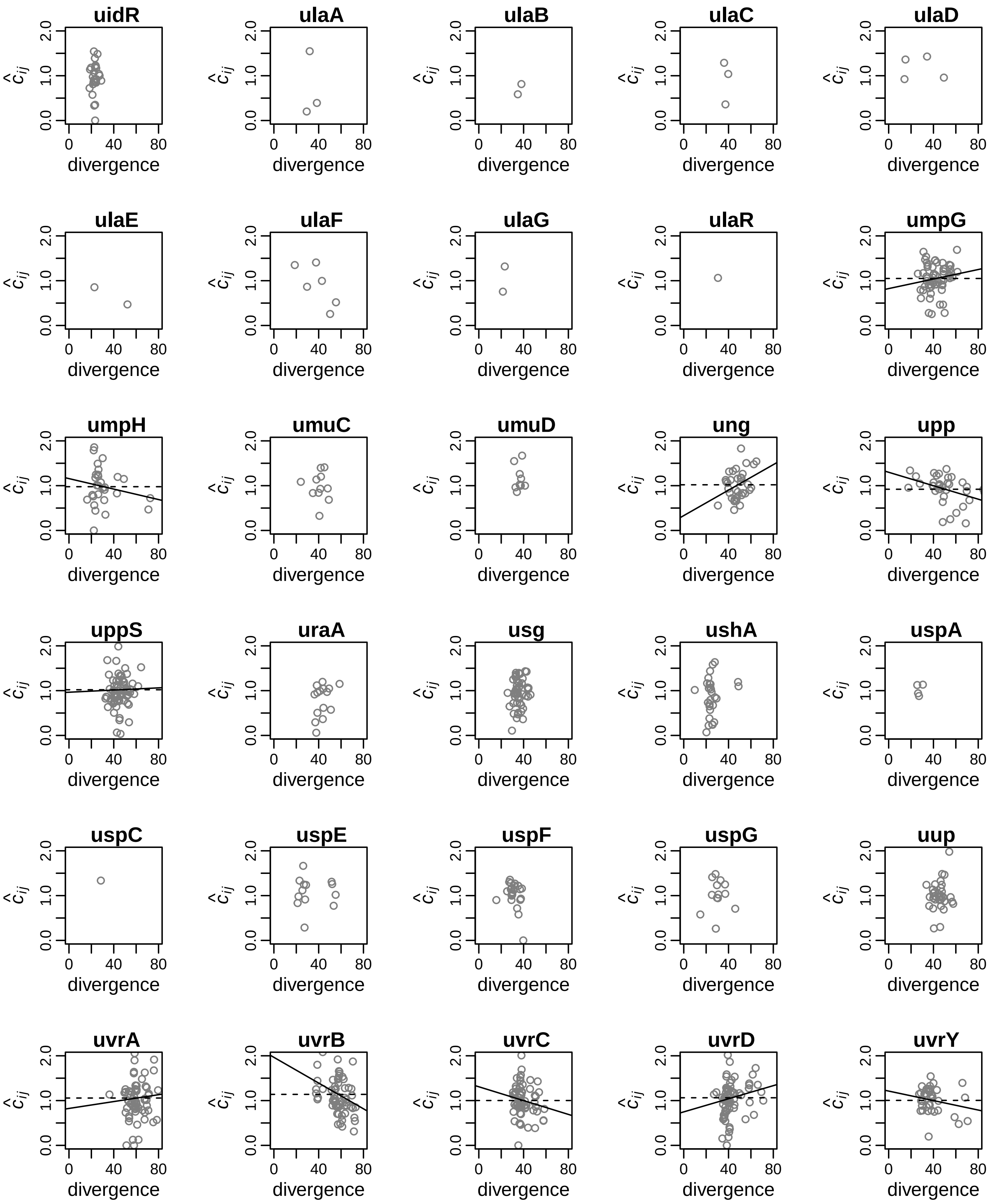
uidR
ulaA
ulaB
ulaC
ulaD
ulaE
ulaF
ulaG
ulaR
umpG
umpH
umuC
umuD
ung
upp
uppS
uraA
usg
ushA
uspA
uspC
uspE
uspF
uspG
uup
uvrA
uvrB
uvrC
uvrD
uvrY
$\hat{c}_{ij}$
0.0
1.0
2.0
0
40
80
divergence

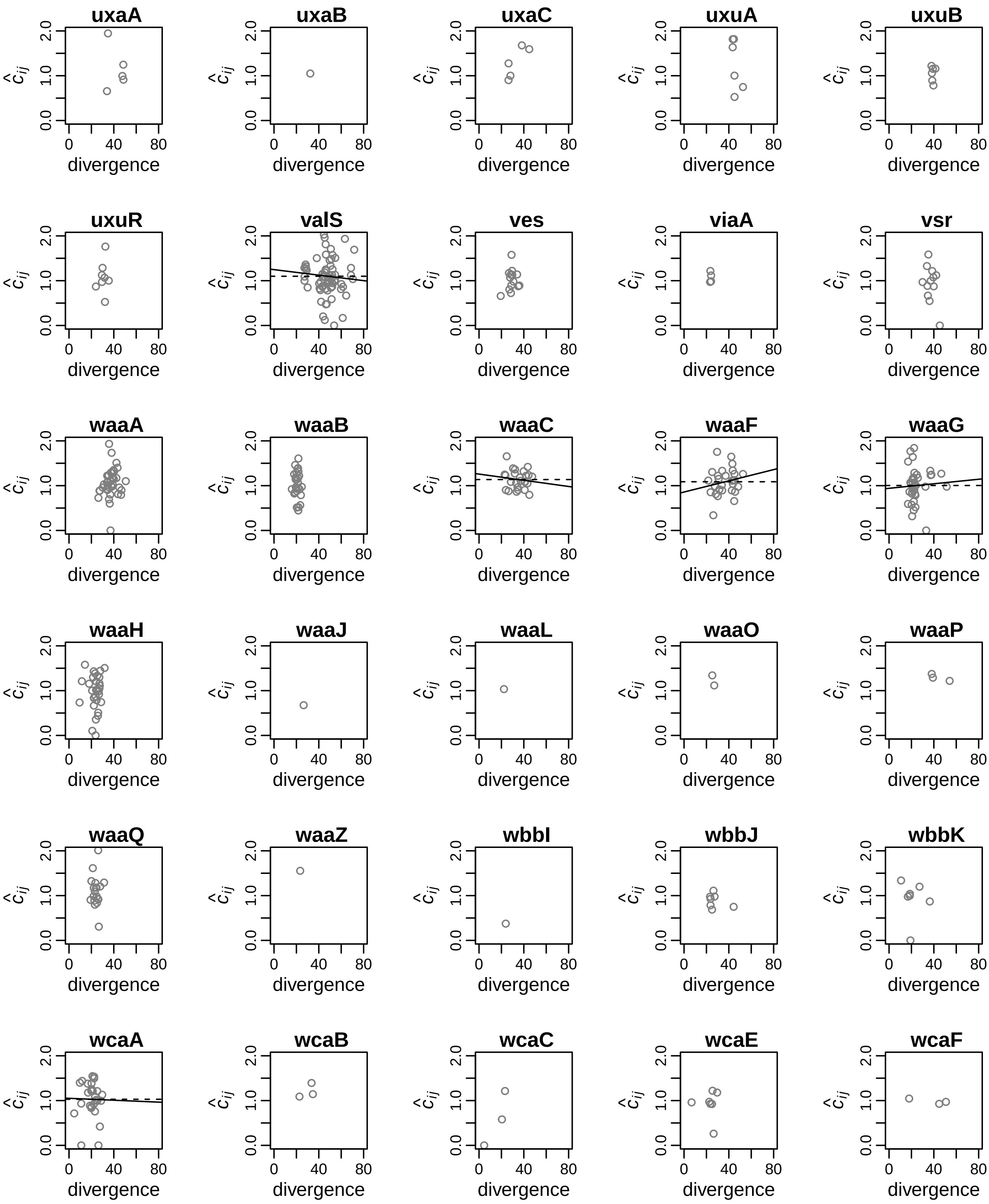
uxaA
uxaB
uxaC
uxuA
uxuB
uxuR
valS
ves
viaA
vsr
waaA
waaB
waaC
waaF
waaG
waaH
waaJ
waaL
waaO
waaP
waaQ
waaZ
wbbI
wbbJ
wbbK
wcaA
wcaB
wcaC
wcaE
wcaF
$\hat{c}_{ij}$
0.0 1.0 2.0
0 40 80
divergence

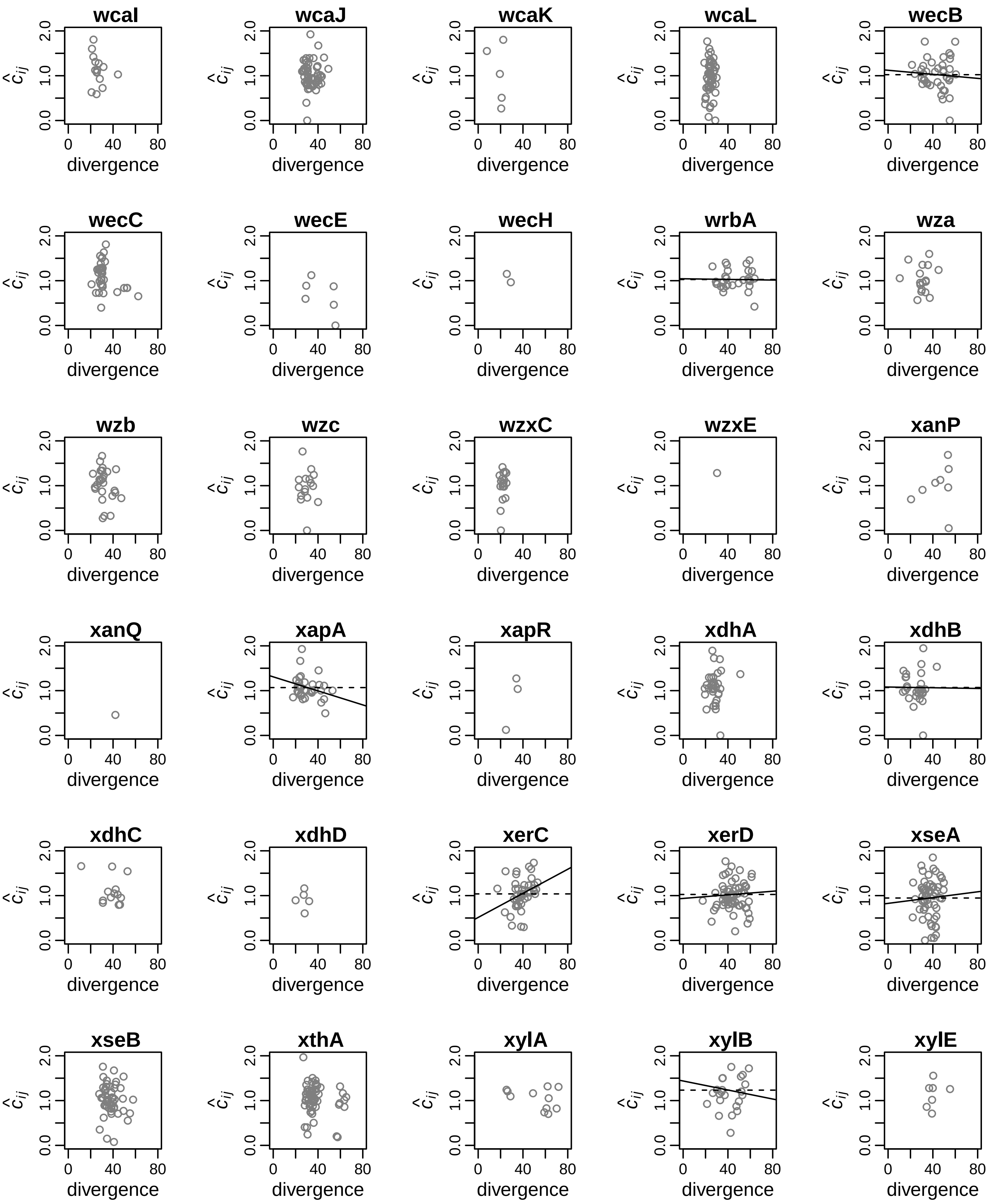
wcaI
wcaJ
wcaK
wcaL
wecB
wecC
wecE
wecH
wrbA
wza
wzb
wzc
wzxC
wzxE
xanP
xanQ
xapA
xapR
xdhA
xdhB
xdhC
xdhD
xerC
xerD
xseA
xseB
xthA
xylA
xylB
xylE
$\hat{c}_{ij}$
0.0
1.0
2.0
0
40
80
divergence

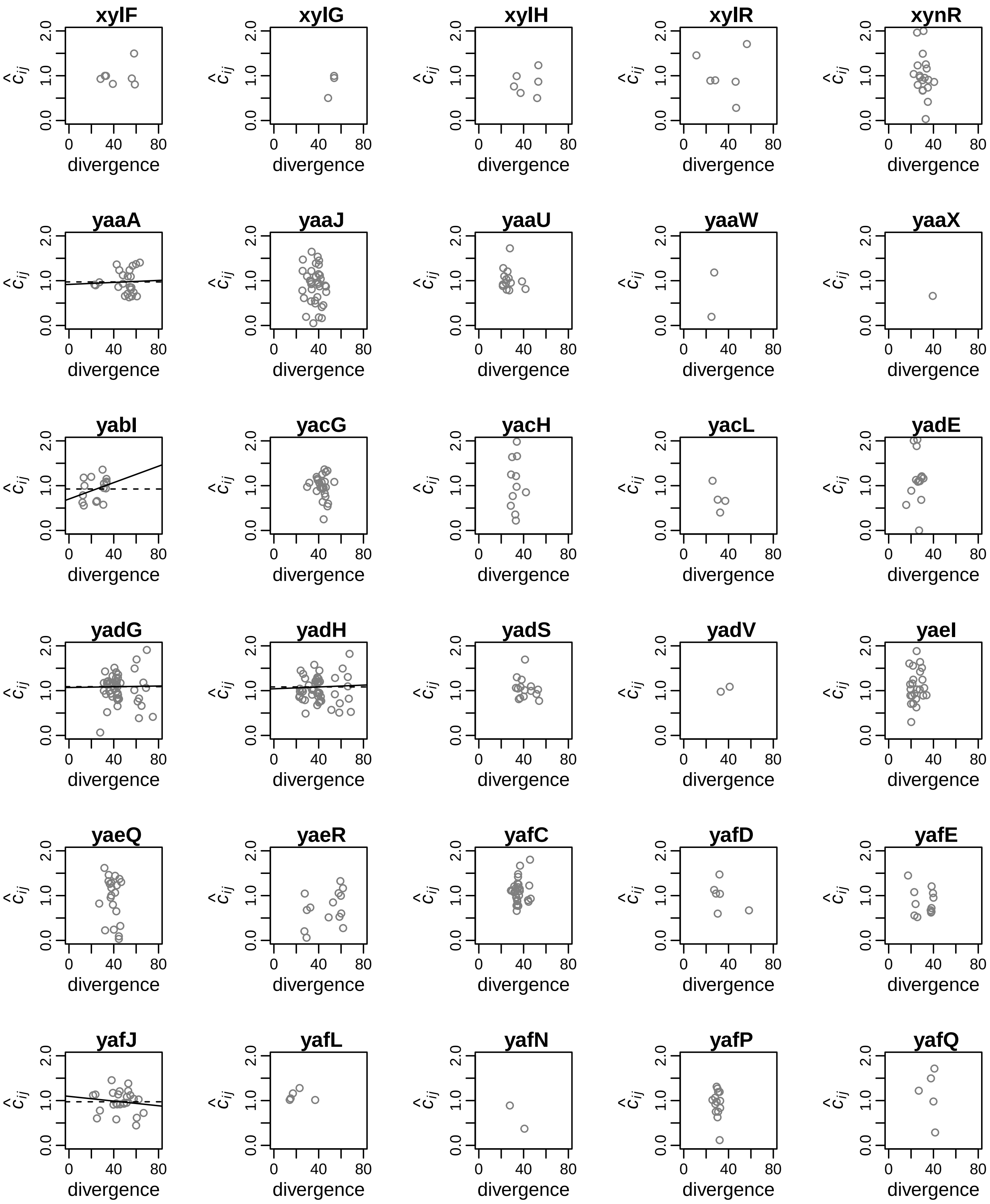
xylF
xylG
xylH
xylR
xynR
yaaA
yaaJ
yaaU
yaaW
yaaX
yabI
yacG
yacH
yacL
yadE
yadG
yadH
yadS
yadV
yaeI
yaeQ
yaeR
yafC
yafD
yafE
yafJ
yafL
yafN
yafP
yafQ
$\hat{c}_{ij}$
divergence
0.0
1.0
2.0
0
40
80

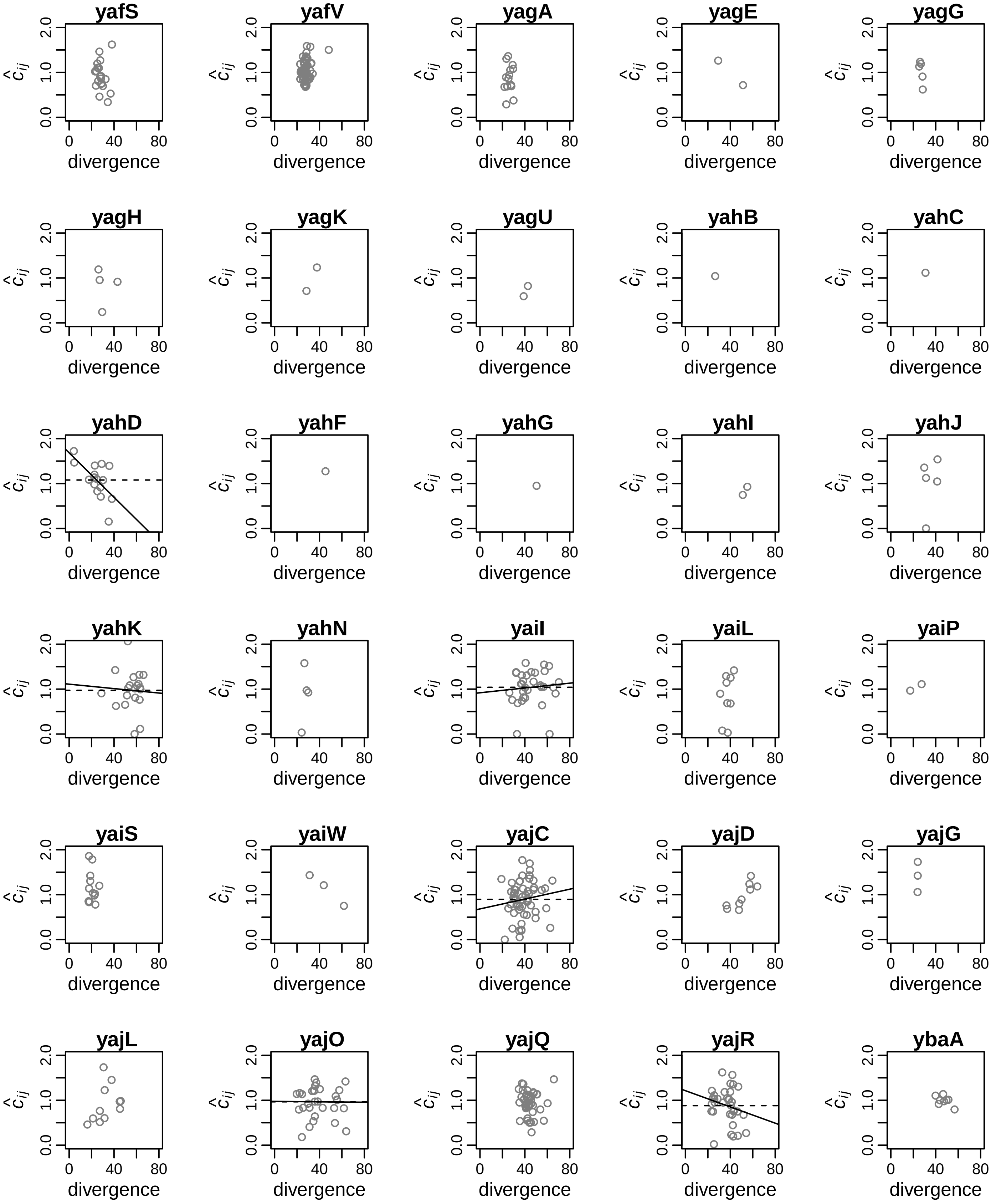

yafS
yafV
yagA
yagE
yagG
yagH
yagK
yagU
yahB
yahC
yahD
yahF
yahG
yahI
yahJ
yahK
yahN
yaiI
yaiL
yaiP
yaiS
yaiW
yajC
yajD
yajG
yajL
yajO
yajQ
yajR
ybaA
$\hat{c}_{ij}$
0.0
1.0
2.0
0
40
80
divergence

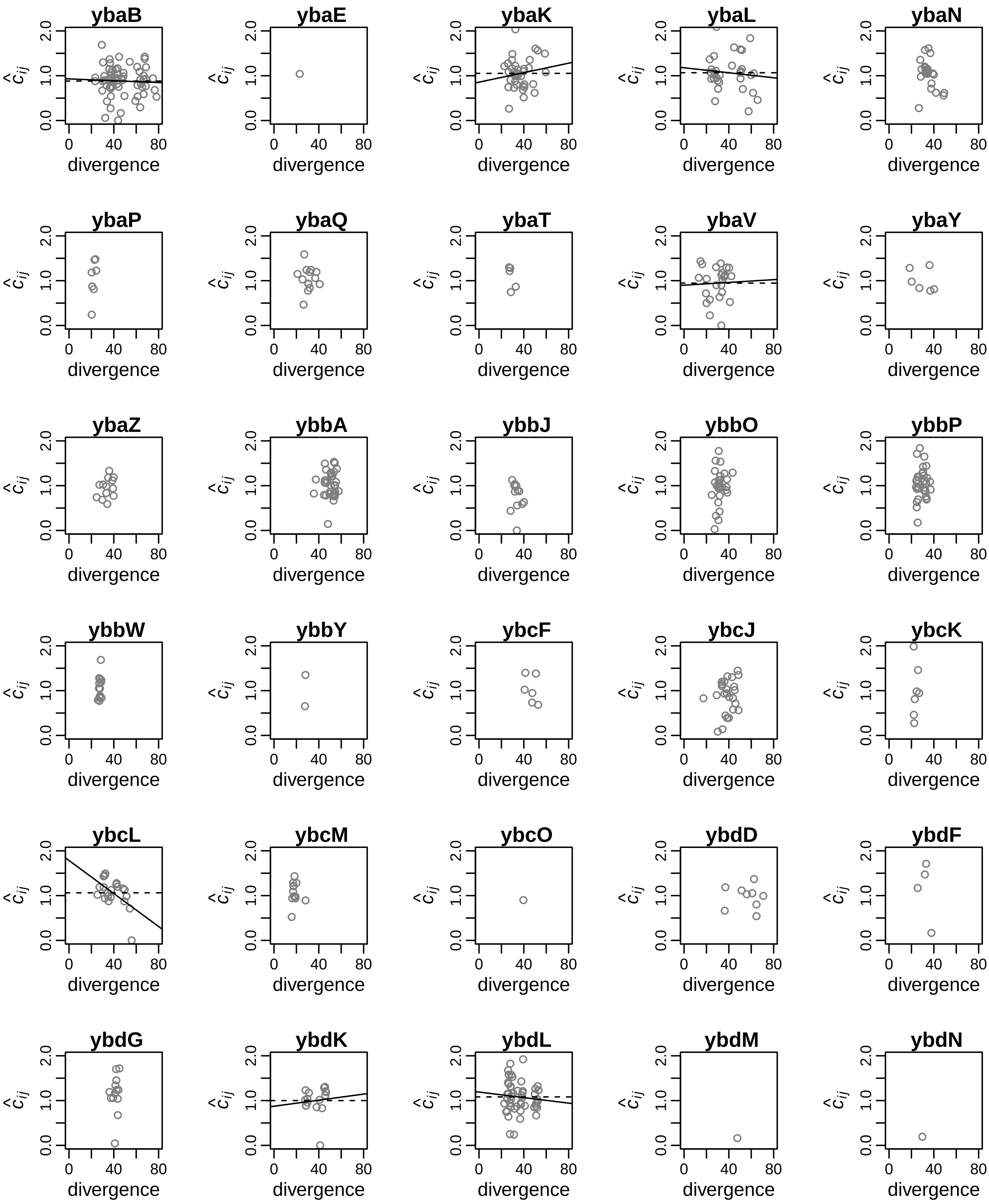
ybaB
ybaE
ybaK
ybaL
ybaN
ybaP
ybaQ
ybaT
ybaV
ybaY
ybaZ
ybbA
ybbJ
ybbO
ybbP
ybbW
ybbY
ybcF
ybcJ
ybcK
ybcL
ybcM
ybcO
ybdD
ybdF
ybdG
ybdK
ybdL
ybdM
ybdN
divergence
$\hat{c}_{ij}$
0.0
1.0
2.0
0
40
80

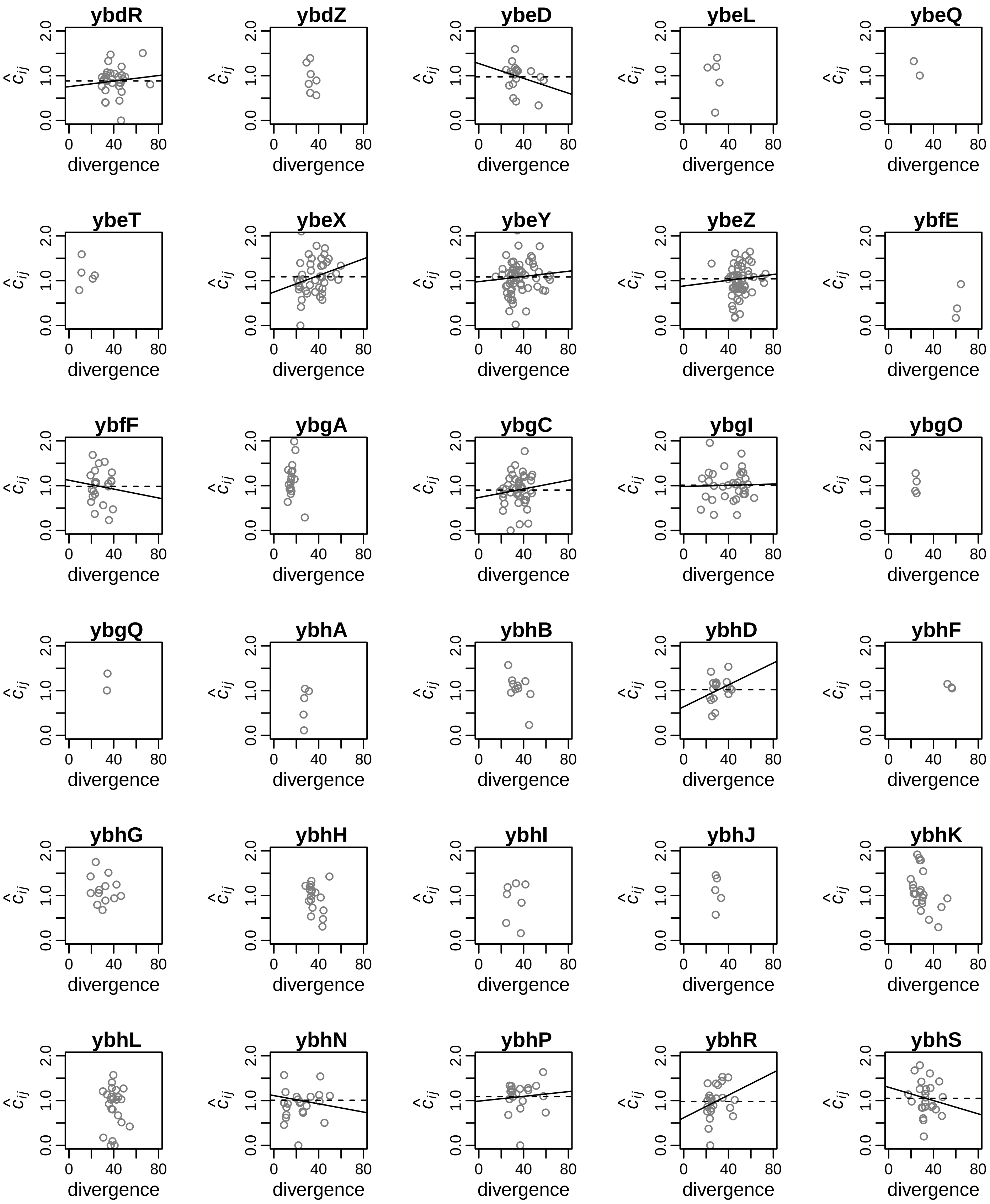
ybdR
ybdZ
ybeD
ybeL
ybeQ
ybeT
ybeX
ybeY
ybeZ
ybfE
ybfF
ybgA
ybgC
ybgI
ybgO
ybgQ
ybhA
ybhB
ybhD
ybhF
ybhG
ybhH
ybhI
ybhJ
ybhK
ybhL
ybhN
ybhP
ybhR
ybhS
$\hat{c}_{ij}$
0.0
1.0
2.0
0
40
80
divergence

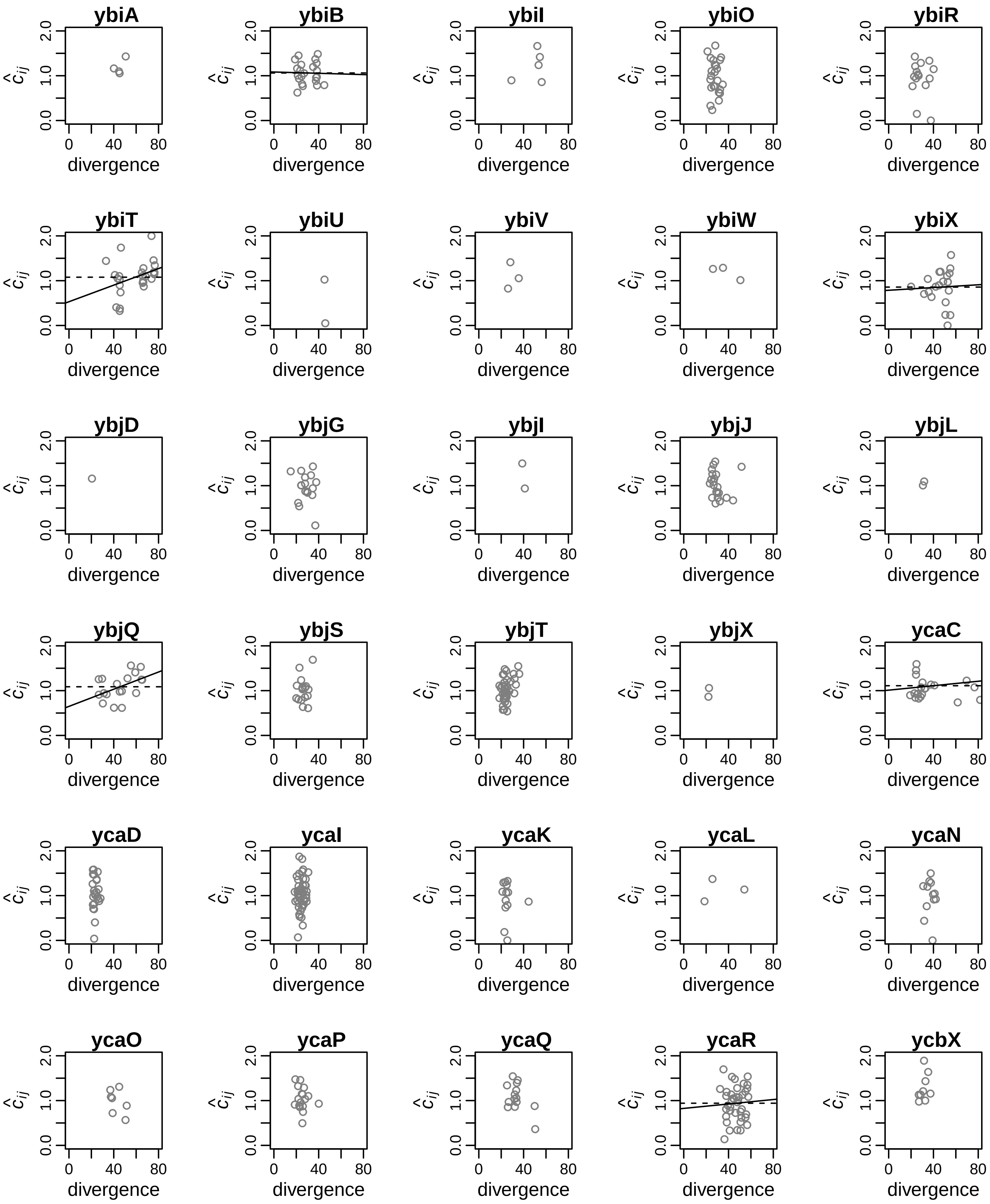
ybiA
ybiB
ybiI
ybiO
ybiR
ybiT
ybiU
ybiV
ybiW
ybiX
ybjD
ybjG
ybjI
ybjJ
ybjL
ybjQ
ybjS
ybjT
ybjX
ycaC
ycaD
ycaI
ycaK
ycaL
ycaN
ycaO
ycaP
ycaQ
ycaR
ycbX
$\hat{c}_{ij}$
divergence
0.0
1.0
2.0
0
40
80

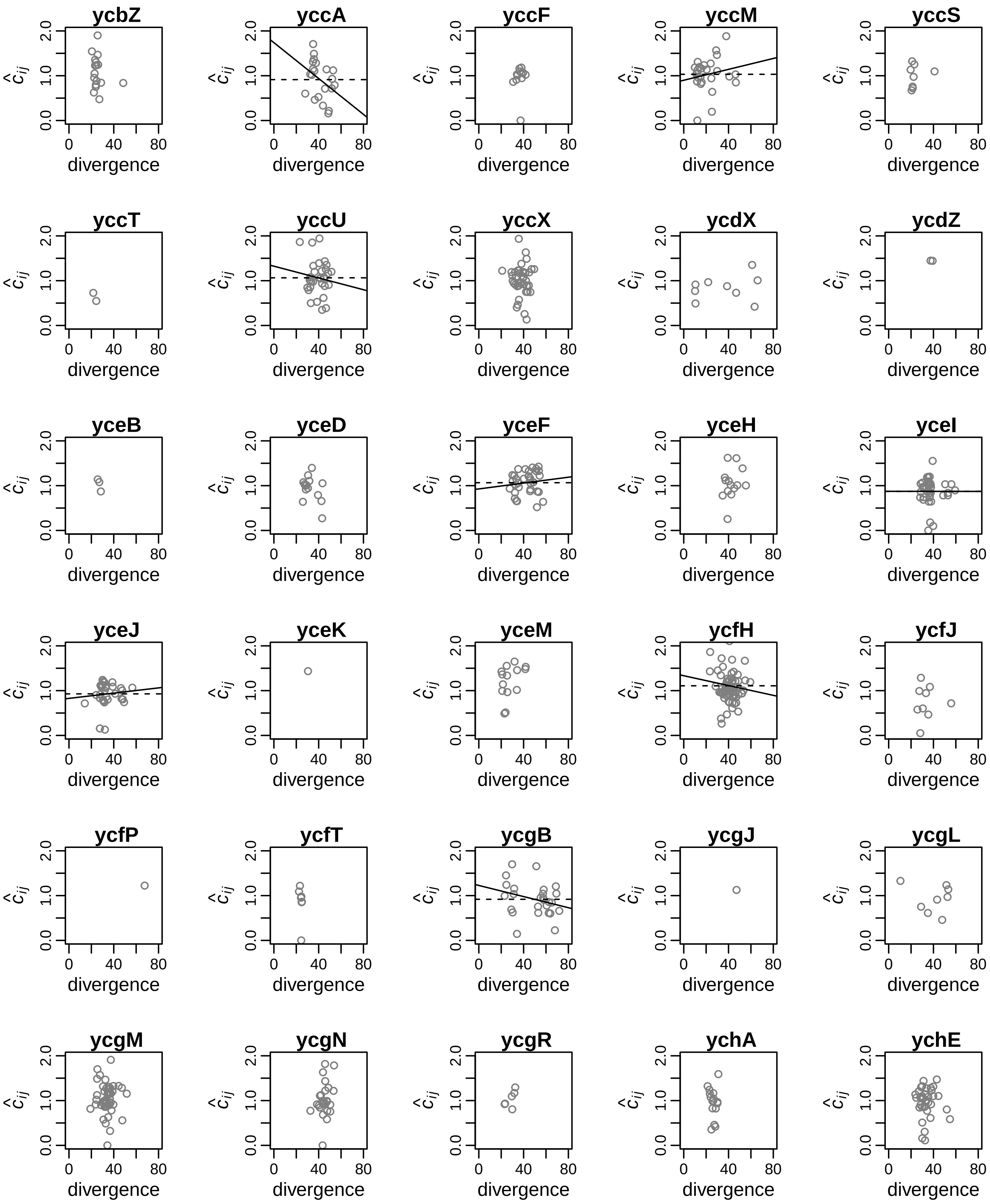
ycbZ
yccA
yccF
yccM
yccS
yccT
yccU
yccX
ycdX
ycdZ
yceB
yceD
yceF
yceH
yceI
yceJ
yceK
yceM
ycfH
ycfJ
ycfP
ycfT
ycgB
ycgJ
ycgL
ycgM
ycgN
ycgR
ychA
ychE
$\hat{c}_{ij}$
0.0
1.0
2.0
0
40
80
divergence

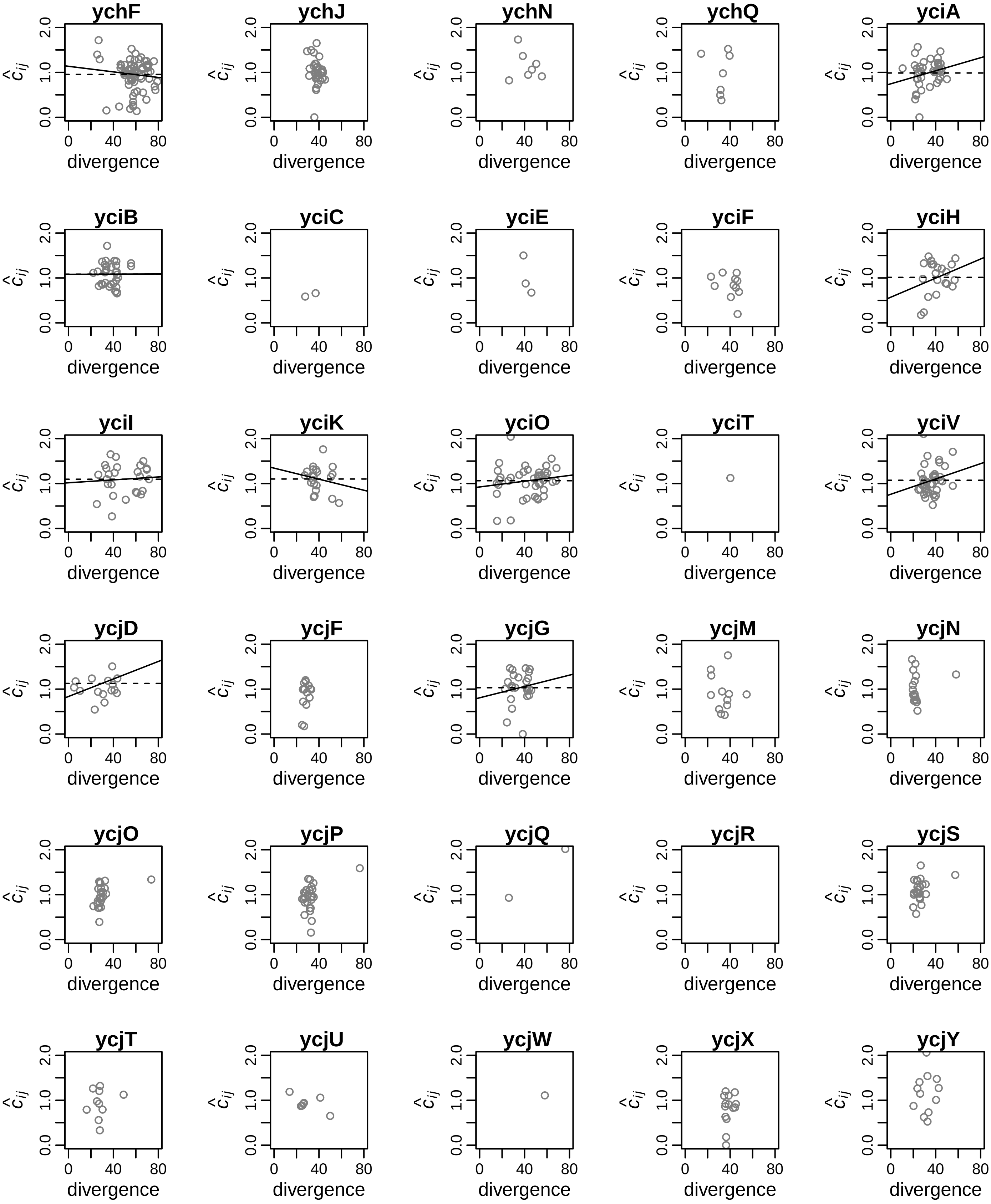

ychF
ychJ
ychN
ychQ
yciA
yciB
yciC
yciE
yciF
yciH
yciI
yciK
yciO
yciT
yciV
ycjD
ycjF
ycjG
ycjM
ycjN
ycjO
ycjP
ycjQ
ycjR
ycjS
ycjT
ycjU
ycjW
ycjX
ycjY
$\hat{c}_{ij}$
0.0
1.0
2.0
0
40
80
divergence

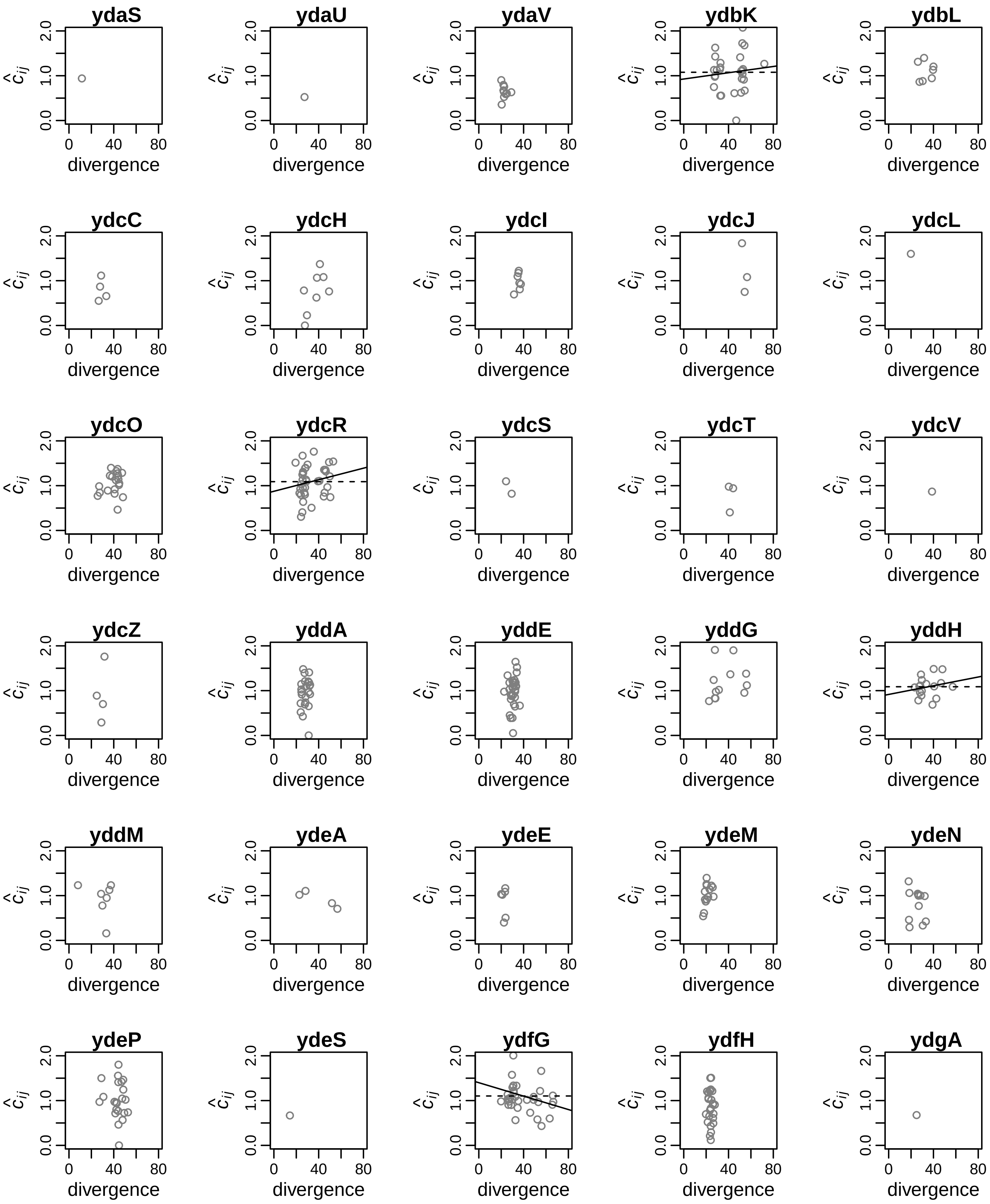
ydaS
ydaU
ydaV
ydbK
ydbL
ydcC
ydcH
ydcI
ydcJ
ydcL
ydcO
ydcR
ydcS
ydcT
ydcV
ydcZ
yddA
yddE
yddG
yddH
yddM
ydeA
ydeE
ydeM
ydeN
ydeP
ydeS
ydfG
ydfH
ydgA
$\hat{c}_{ij}$
0.0
1.0
2.0
0
40
80
divergence

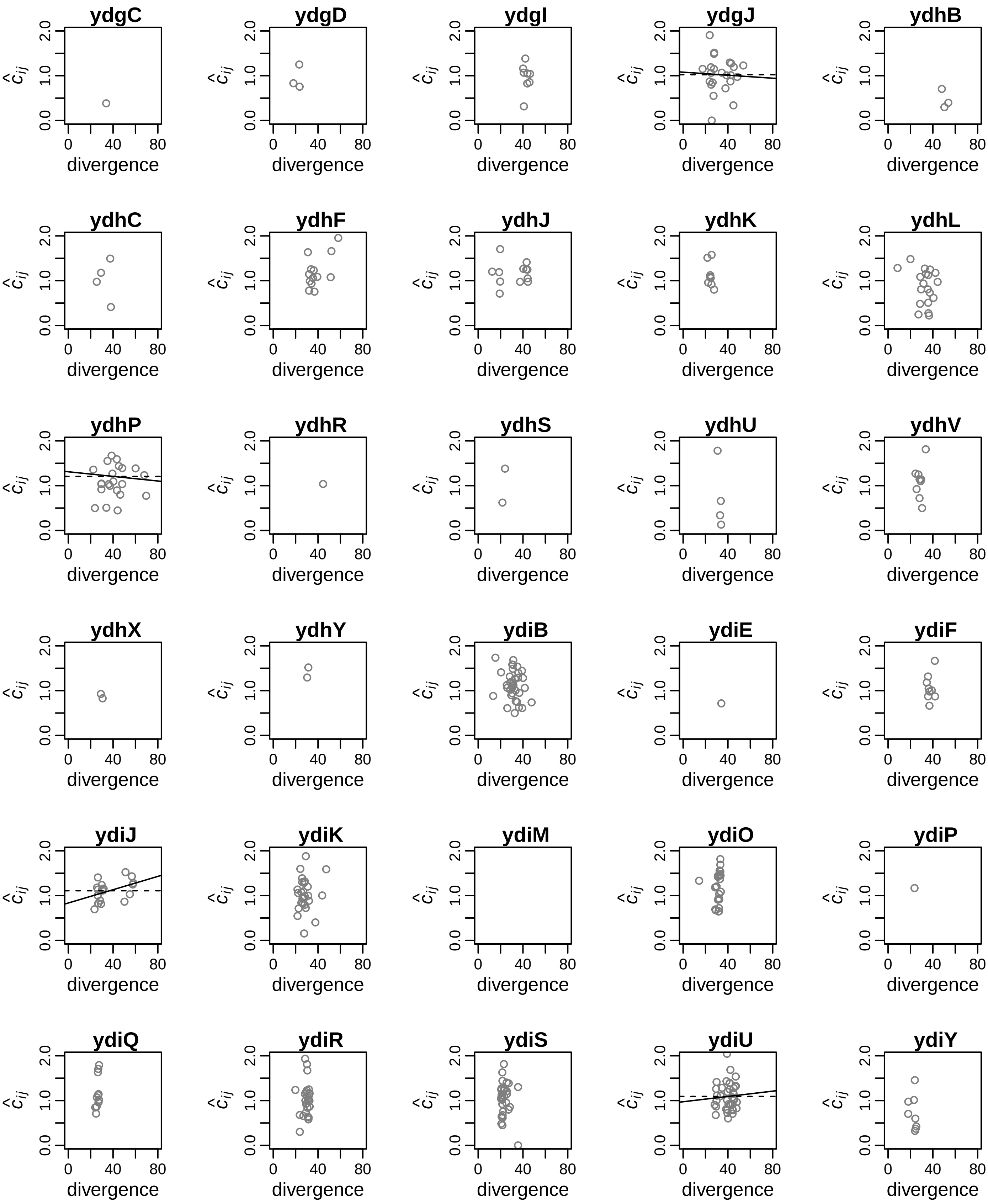
ydgC
ydgD
ydgI
ydgJ
ydhB
ydhC
ydhF
ydhJ
ydhK
ydhL
ydhP
ydhR
ydhS
ydhU
ydhV
ydhX
ydhY
ydiB
ydiE
ydiF
ydiJ
ydiK
ydiM
ydiO
ydiP
ydiQ
ydiR
ydiS
ydiU
ydiY
$\hat{c}_{ij}$
0.0
1.0
2.0
0
40
80
divergence

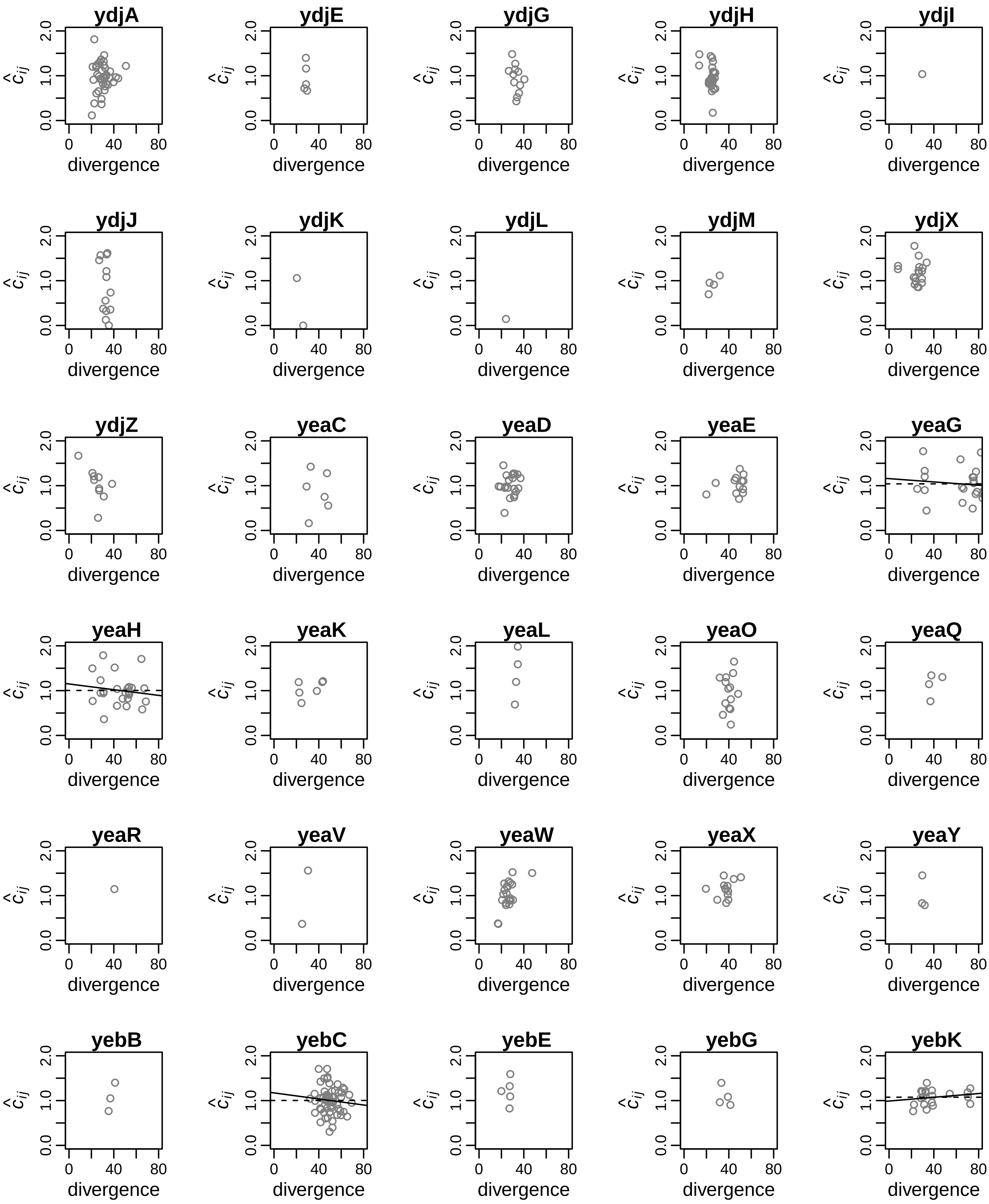
ydjA
ydjE
ydjG
ydjH
ydjI
ydjJ
ydjK
ydjL
ydjM
ydjX
ydjZ
yeaC
yeaD
yeaE
yeaG
yeaH
yeaK
yeaL
yeaO
yeaQ
yeaR
yeaV
yeaW
yeaX
yeaY
yebB
yebC
yebE
yebG
yebK
$\hat{c}_{ij}$
divergence
0.0
1.0
2.0
0
40
80

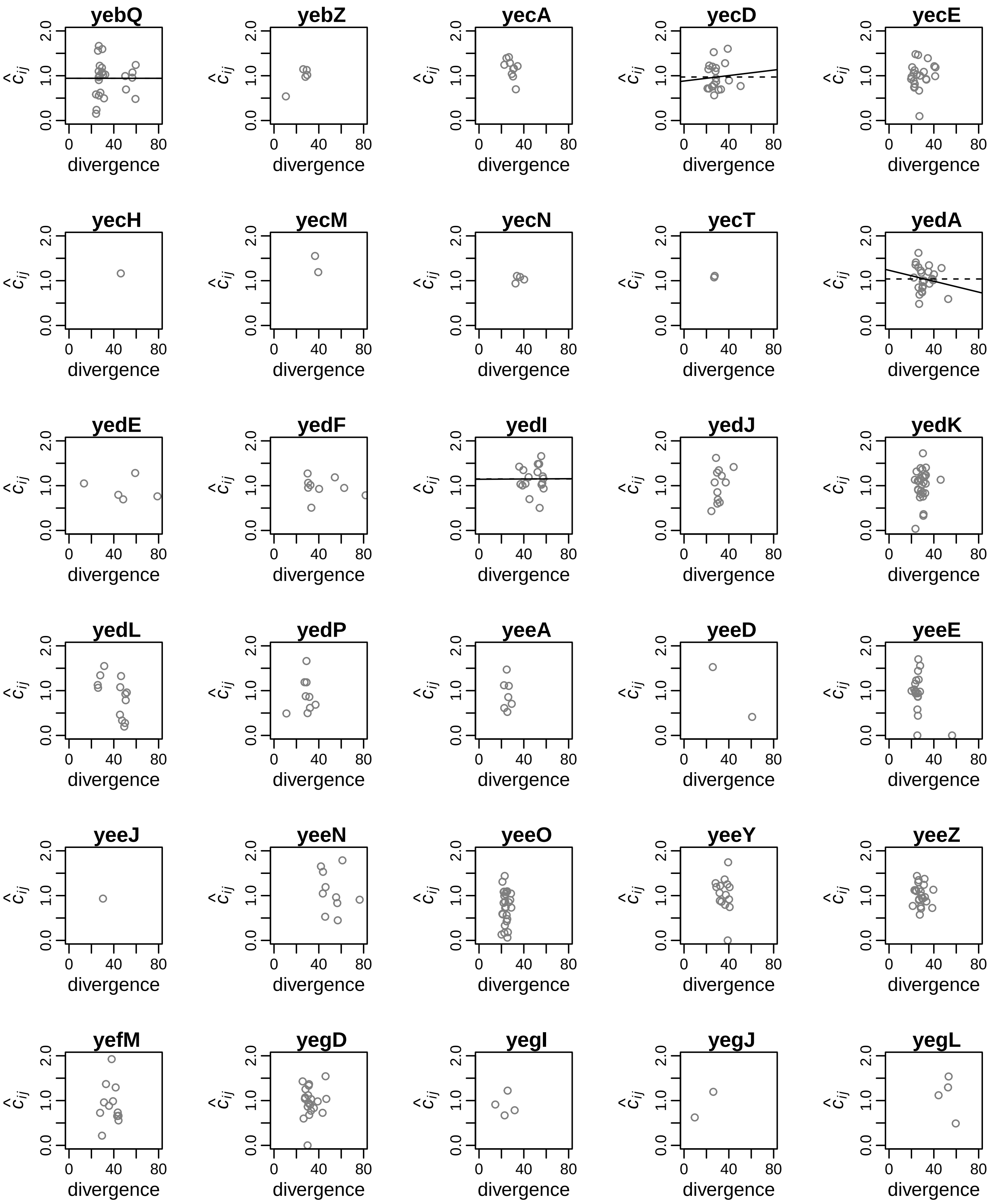
yebQ
yebZ
yecA
yecD
yecE
yecH
yecM
yecN
yecT
yedA
yedE
yedF
yedI
yedJ
yedK
yedL
yedP
yeeA
yeeD
yeeE
yeeJ
yeeN
yeeO
yeeY
yeeZ
yefM
yegD
yegI
yegJ
yegL
$\hat{c}_{ij}$
0.0
1.0
2.0
0
40
80
divergence

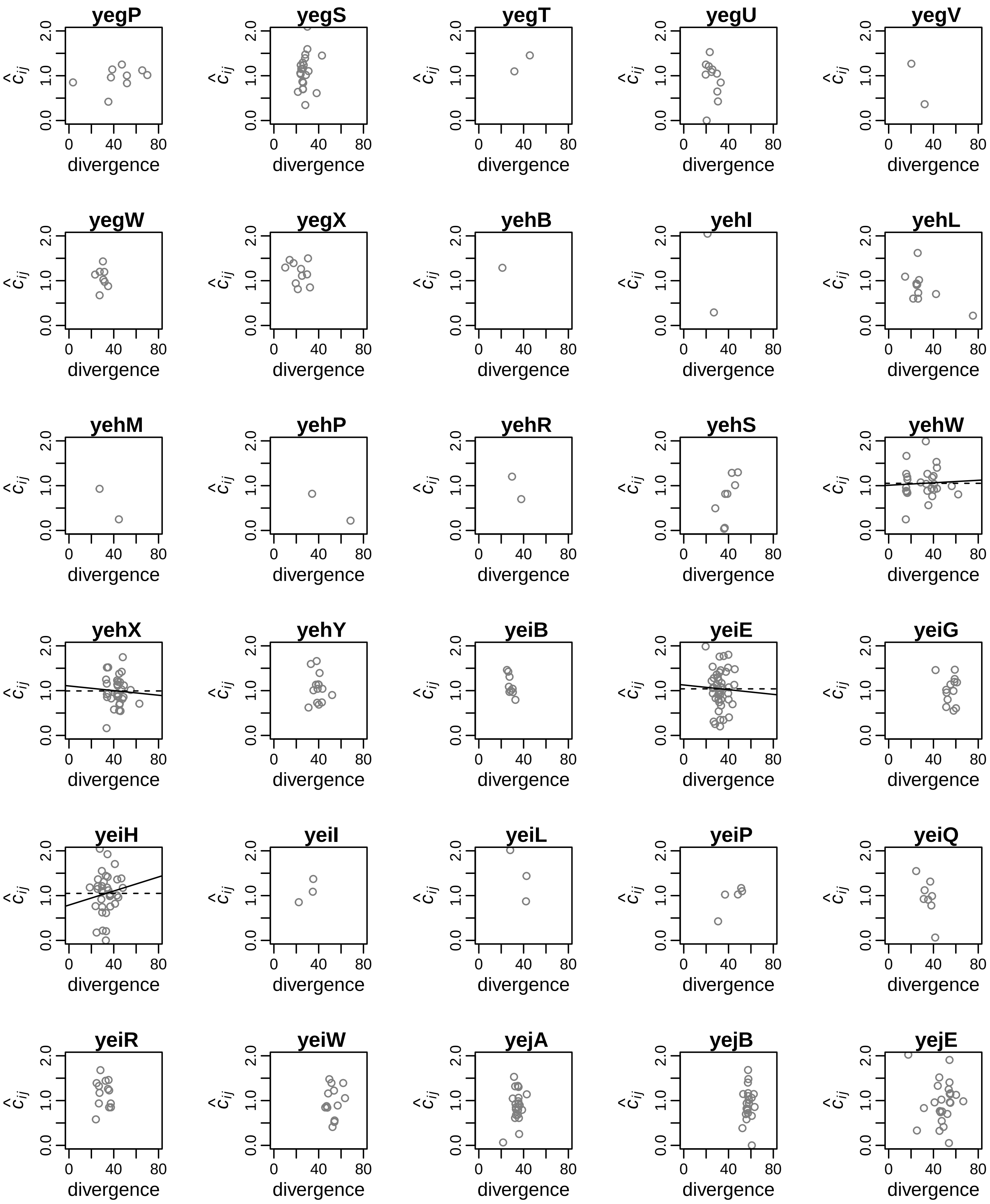
yegP
yegS
yegT
yegU
yegV
yegW
yegX
yehB
yehI
yehL
yehM
yehP
yehR
yehS
yehW
yehX
yehY
yeiB
yeiE
yeiG
yeiH
yeiI
yeiL
yeiP
yeiQ
yeiR
yeiW
yejA
yejB
yejE
$\hat{c}_{ij}$
0.0
1.0
2.0
0
40
80
divergence

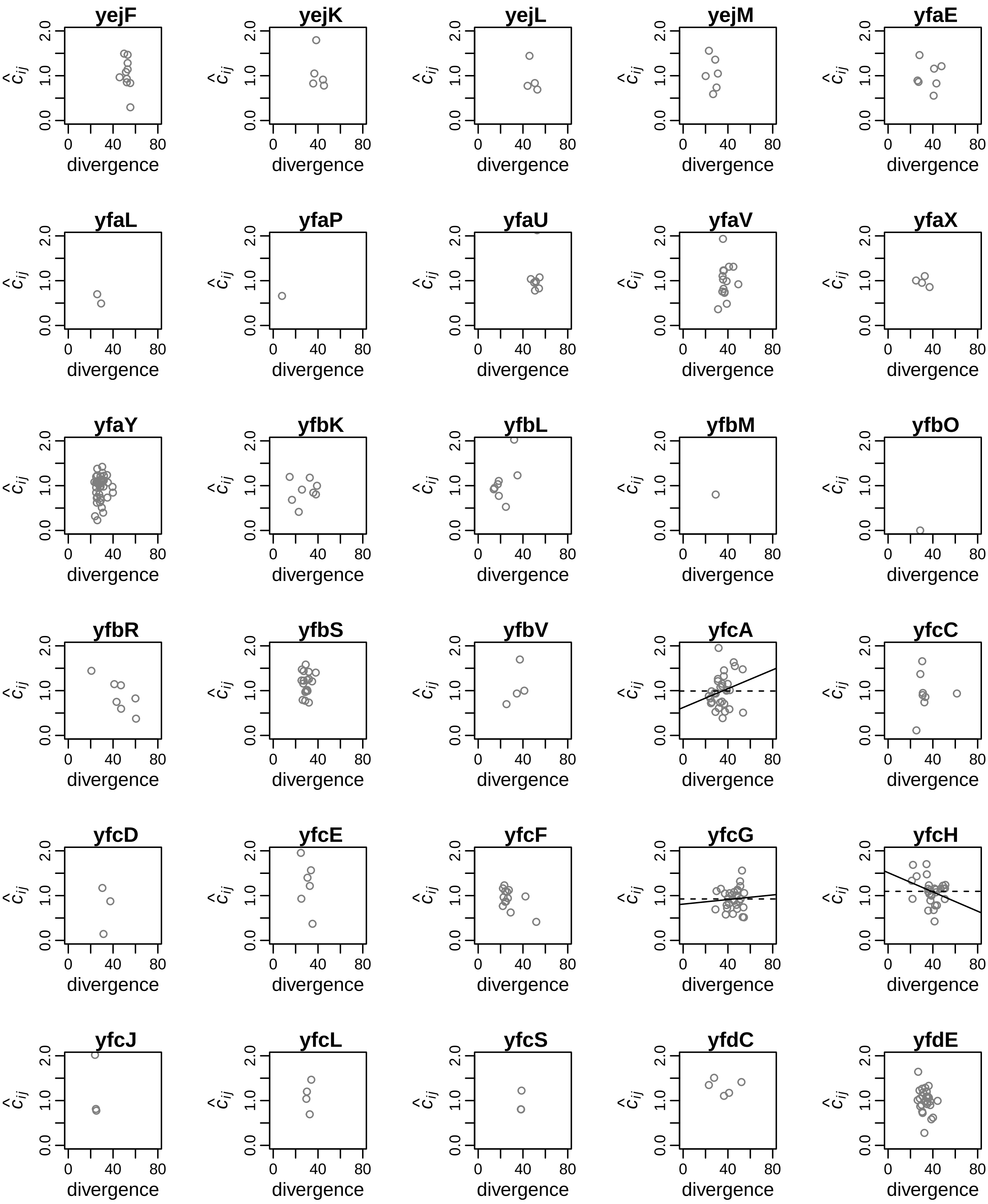
yejF
yejK
yejL
yejM
yfaE
yfaL
yfaP
yfaU
yfaV
yfaX
yfaY
yfbK
yfbL
yfbM
yfbO
yfbR
yfbS
yfbV
yfcA
yfcC
yfcD
yfcE
yfcF
yfcG
yfcH
yfcJ
yfcL
yfcS
yfdC
yfdE
$\hat{c}_{ij}$
0.0
1.0
2.0
0
40
80
divergence

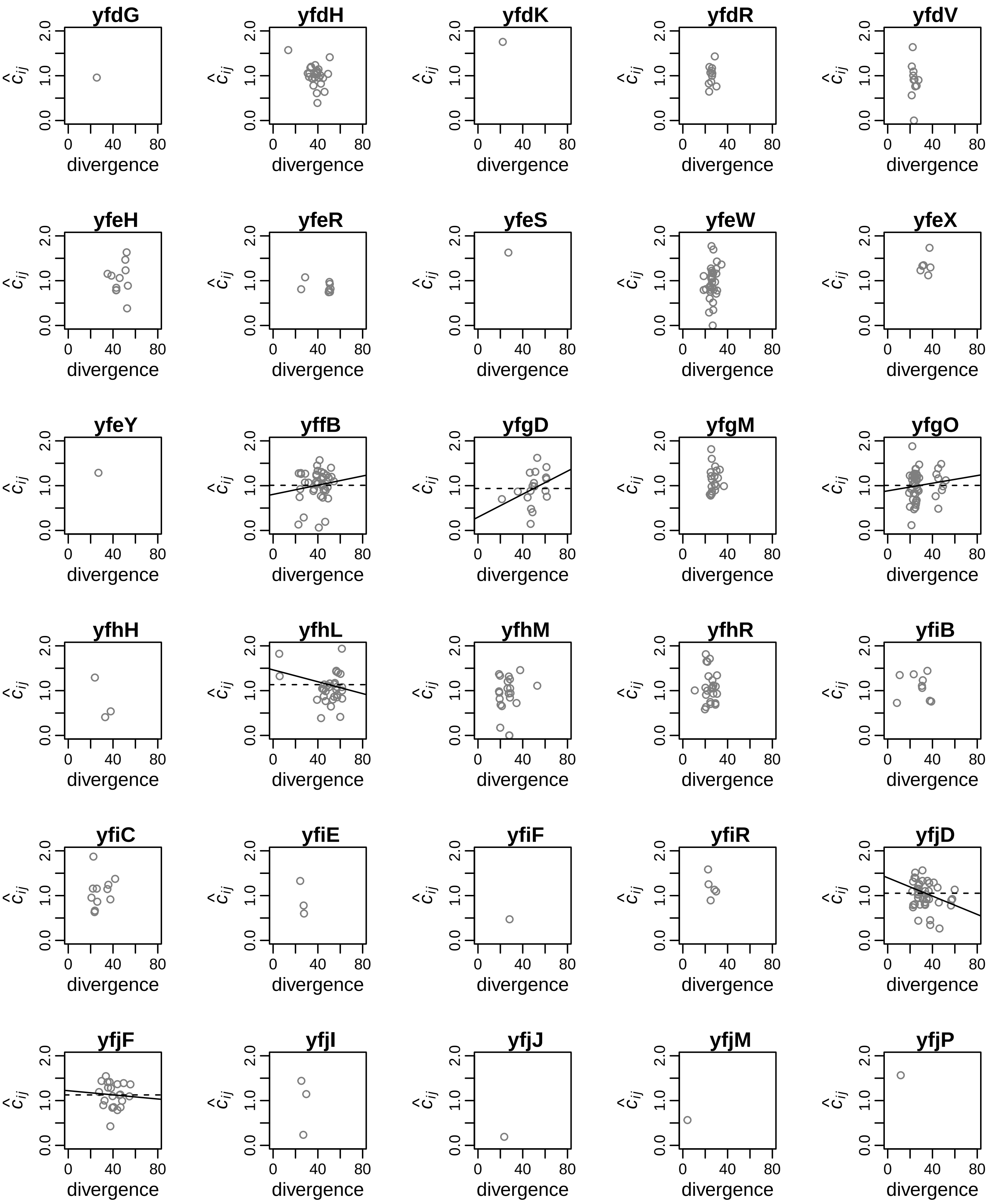
yfdG
yfdH
yfdK
yfdR
yfdV
yfeH
yfeR
yfeS
yfeW
yfeX
yfeY
yffB
yfgD
yfgM
yfgO
yfhH
yfhL
yfhM
yfhR
yfiB
yfiC
yfiE
yfiF
yfiR
yfjD
yfjF
yfjI
yfjJ
yfjM
yfjP
$\hat{c}_{ij}$
0.0
1.0
2.0
0
40
80
divergence

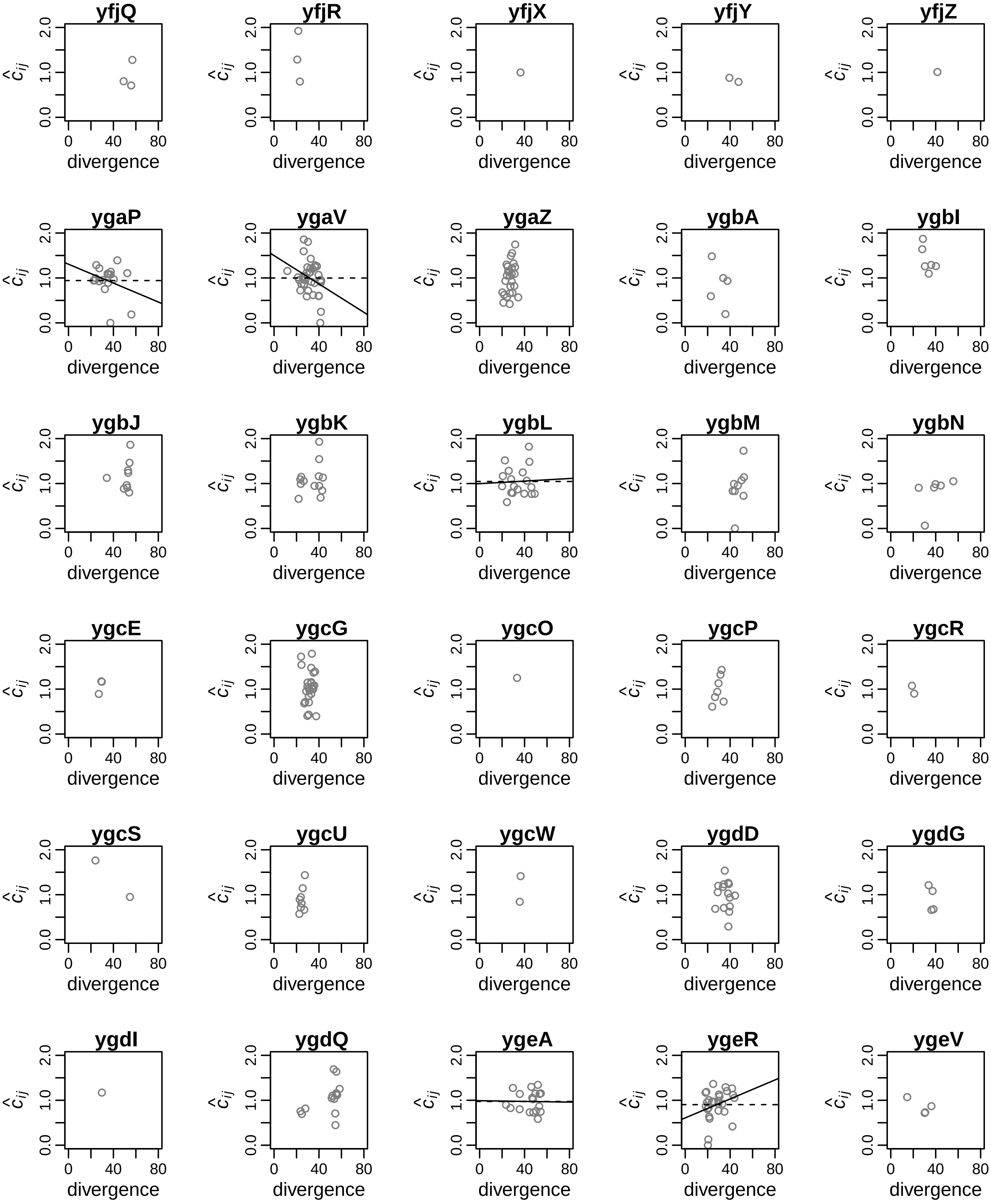
yfjQ
yfjR
yfjX
yfjY
yfjZ
ygaP
ygaV
ygaZ
ygbA
ygbI
ygbJ
ygbK
ygbL
ygbM
ygbN
ygcE
ygcG
ygcO
ygcP
ygcR
ygcS
ygcU
ygcW
ygdD
ygdG
ygdI
ygdQ
ygeA
ygeR
ygeV
$\hat{c}_{ij}$
0.0
1.0
2.0
0
40
80
divergence

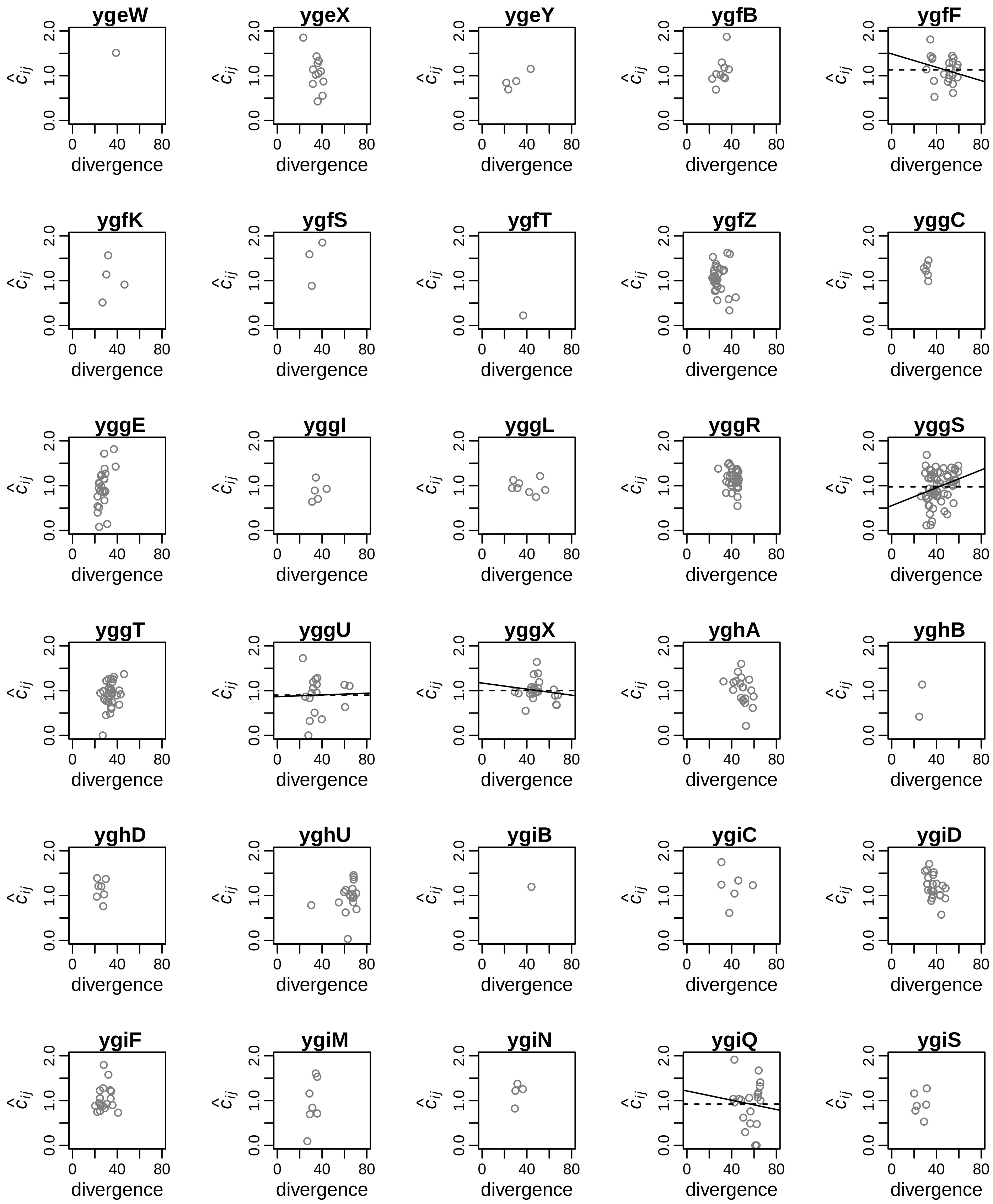
ygeW
ygeX
ygeY
ygfB
ygfF
ygfK
ygfS
ygfT
ygfZ
yggC
yggE
yggI
yggL
yggR
yggS
yggT
yggU
yggX
yghA
yghB
yghD
yghU
ygiB
ygiC
ygiD
ygiF
ygiM
ygiN
ygiQ
ygiS
$\hat{c}_{ij}$
0.0
1.0
2.0
0
40
80
divergence

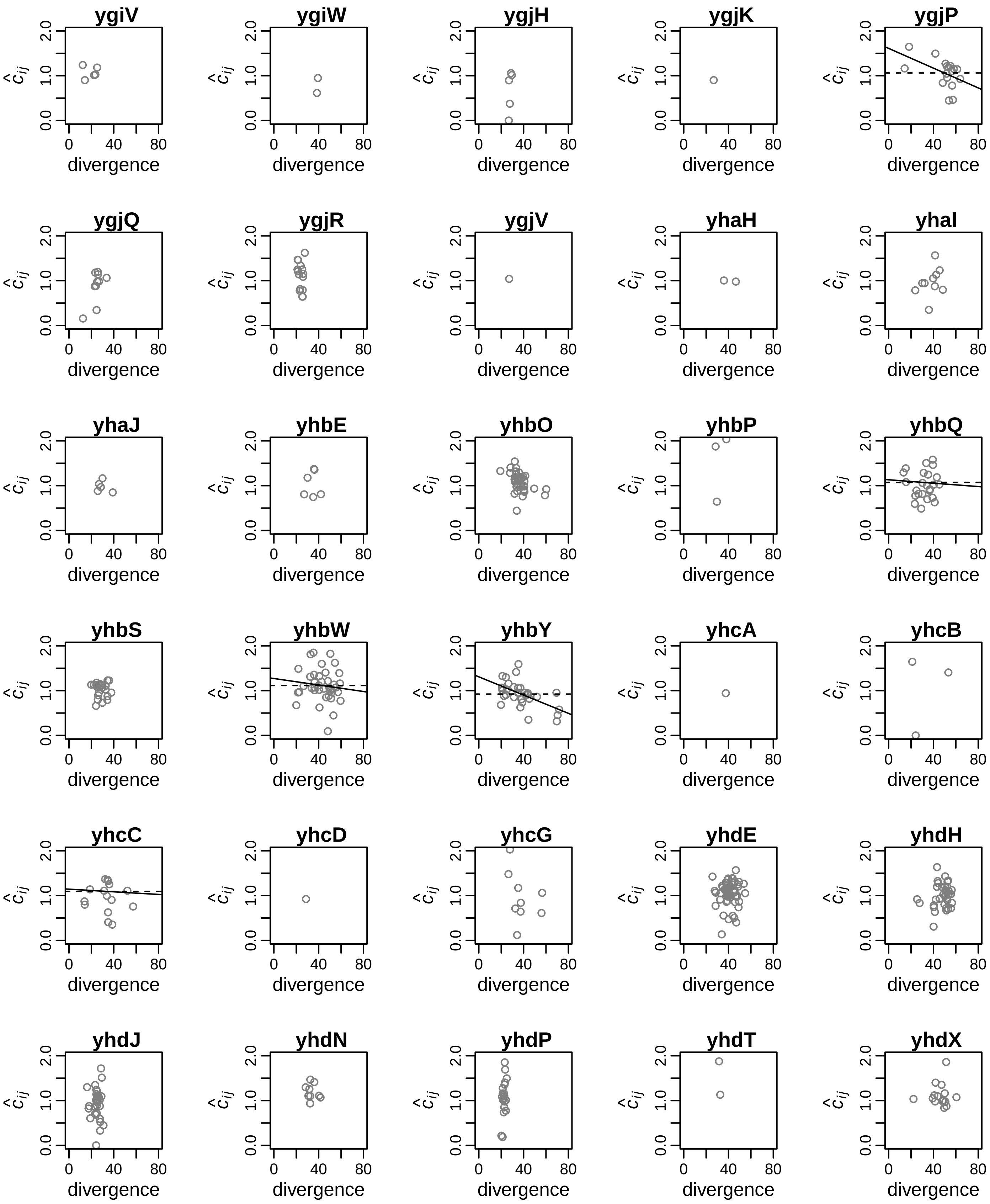
ygiV
ygiW
ygjH
ygjK
ygjP
ygjQ
ygjR
ygjV
yhaH
yhaI
yhaJ
yhbE
yhbO
yhbP
yhbQ
yhbS
yhbW
yhbY
yhcA
yhcB
yhcC
yhcD
yhcG
yhdE
yhdH
yhdJ
yhdN
yhdP
yhdT
yhdX
$\hat{c}_{ij}$
0.0
1.0
2.0
0
40
80
divergence

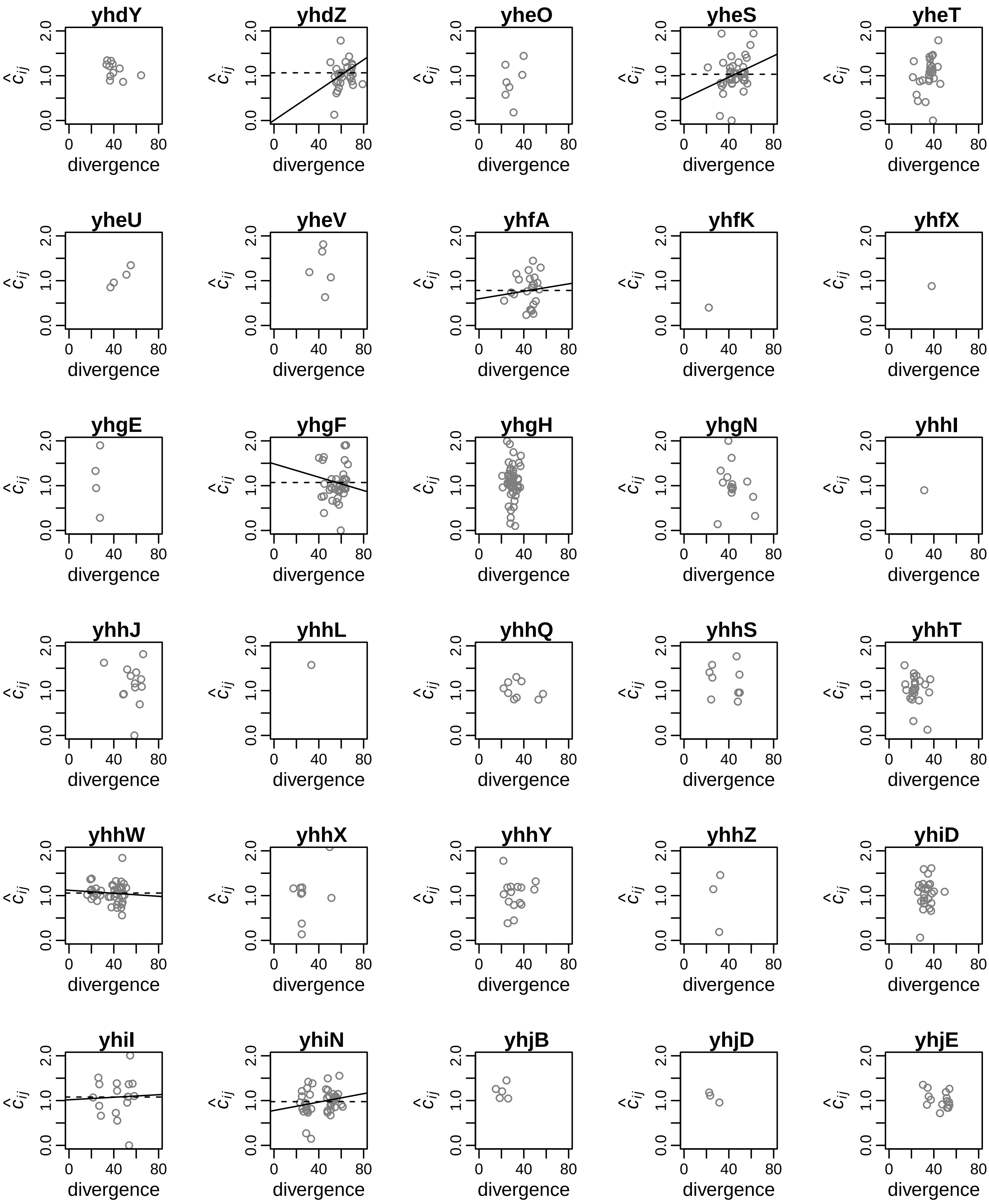
yhdY
yhdZ
yheO
yheS
yheT
yheU
yheV
yhfA
yhfK
yhfX
yhgE
yhgF
yhgH
yhgN
yhhI
yhhJ
yhhL
yhhQ
yhhS
yhhT
yhhW
yhhX
yhhY
yhhZ
yhiD
yhiI
yhiN
yhjB
yhjD
yhjE
$\hat{c}_{ij}$
0.0
1.0
2.0
0
40
80
divergence

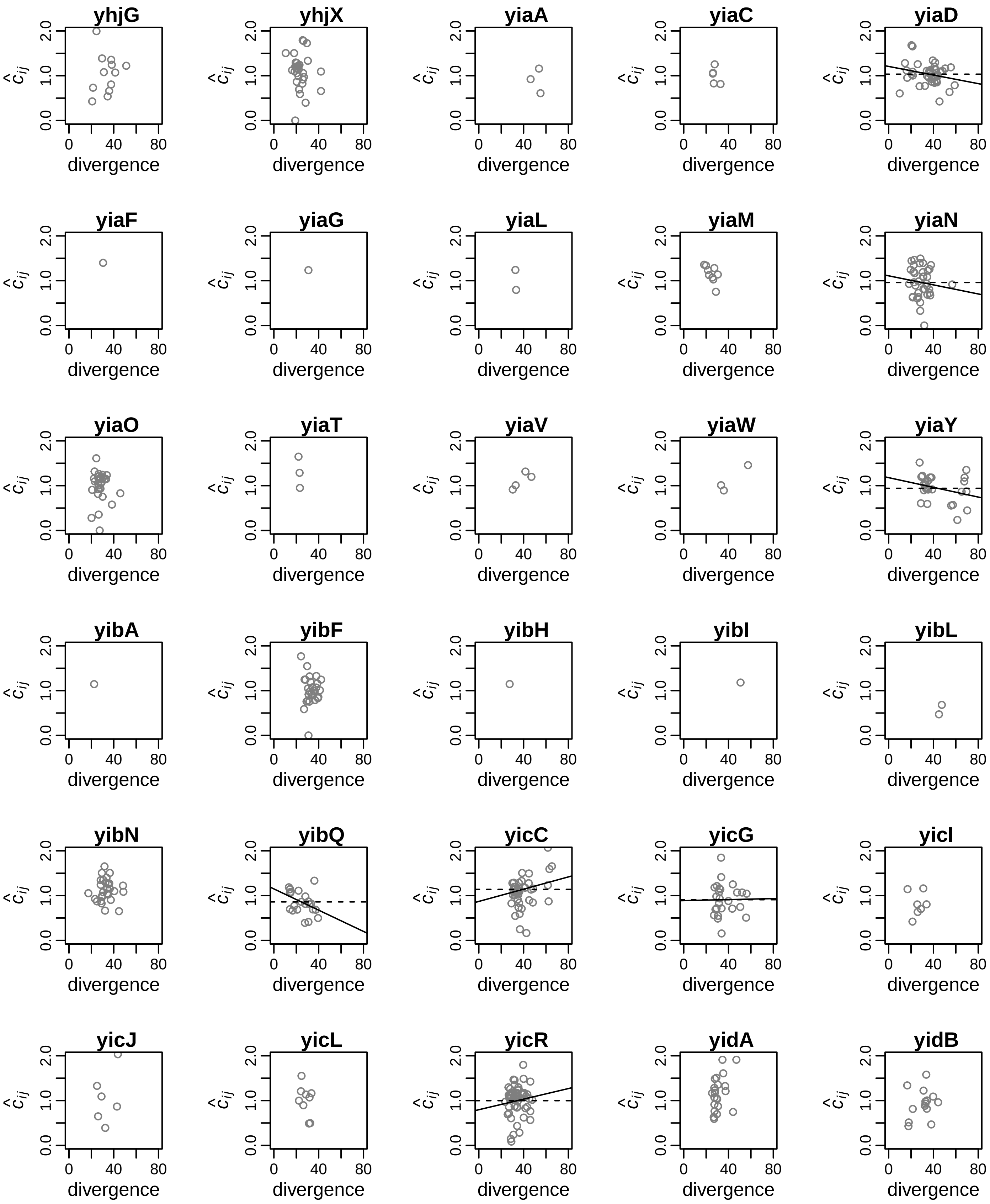
yhjG
yhjX
yiaA
yiaC
yiaD
yiaF
yiaG
yiaL
yiaM
yiaN
yiaO
yiaT
yiaV
yiaW
yiaY
yibA
yibF
yibH
yibI
yibL
yibN
yibQ
yicC
yicG
yicI
yicJ
yicL
yicR
yidA
yidB
$\hat{c}_{ij}$
divergence
0.0
1.0
2.0
0
40
80

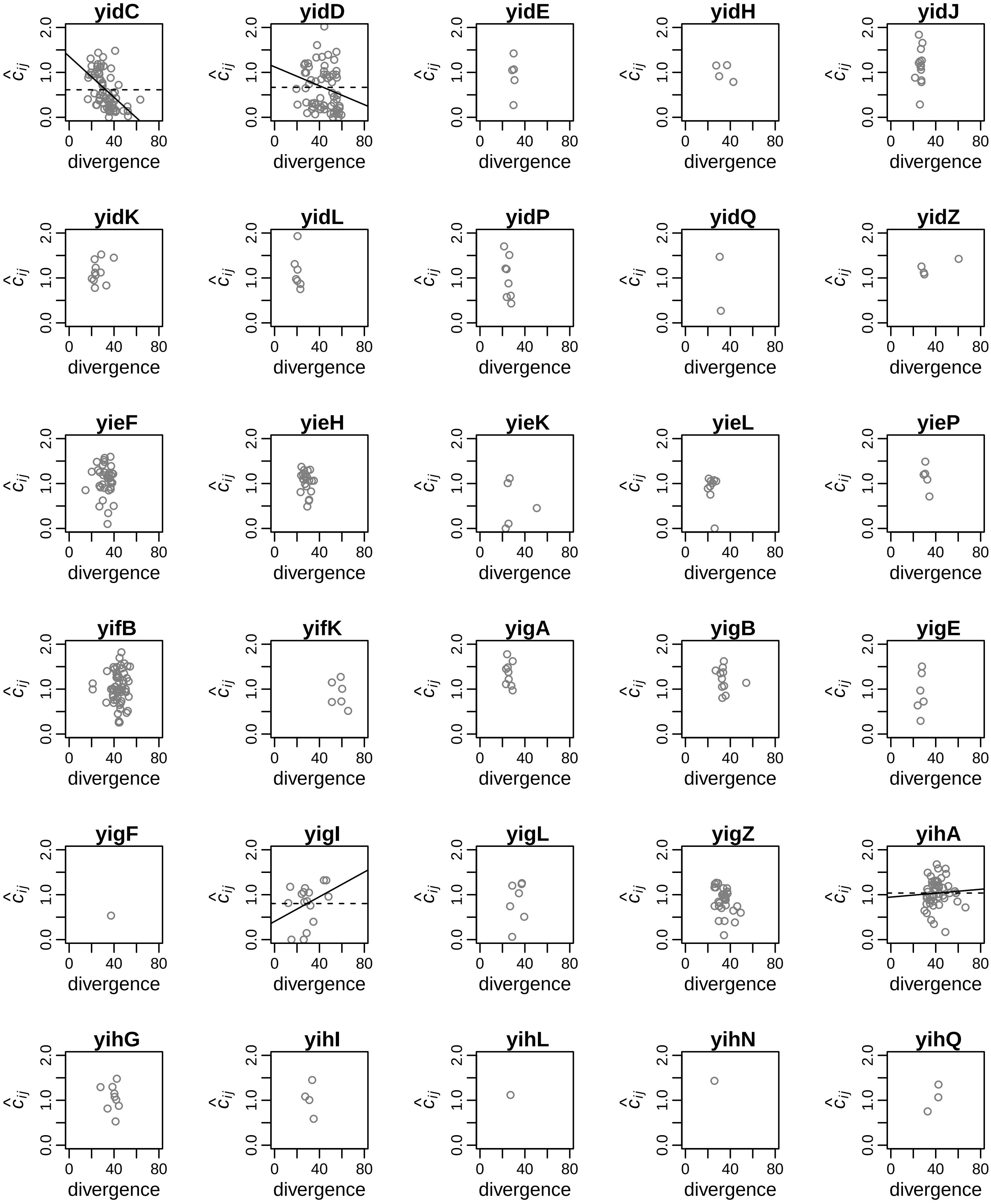
yidC
yidD
yidE
yidH
yidJ
yidK
yidL
yidP
yidQ
yidZ
yieF
yieH
yieK
yieL
yieP
yifB
yifK
yigA
yigB
yigE
yigF
yigI
yigL
yigZ
yihA
yihG
yihI
yihL
yihN
yihQ
$\hat{c}_{ij}$
0.0
1.0
2.0
0
40
80
divergence

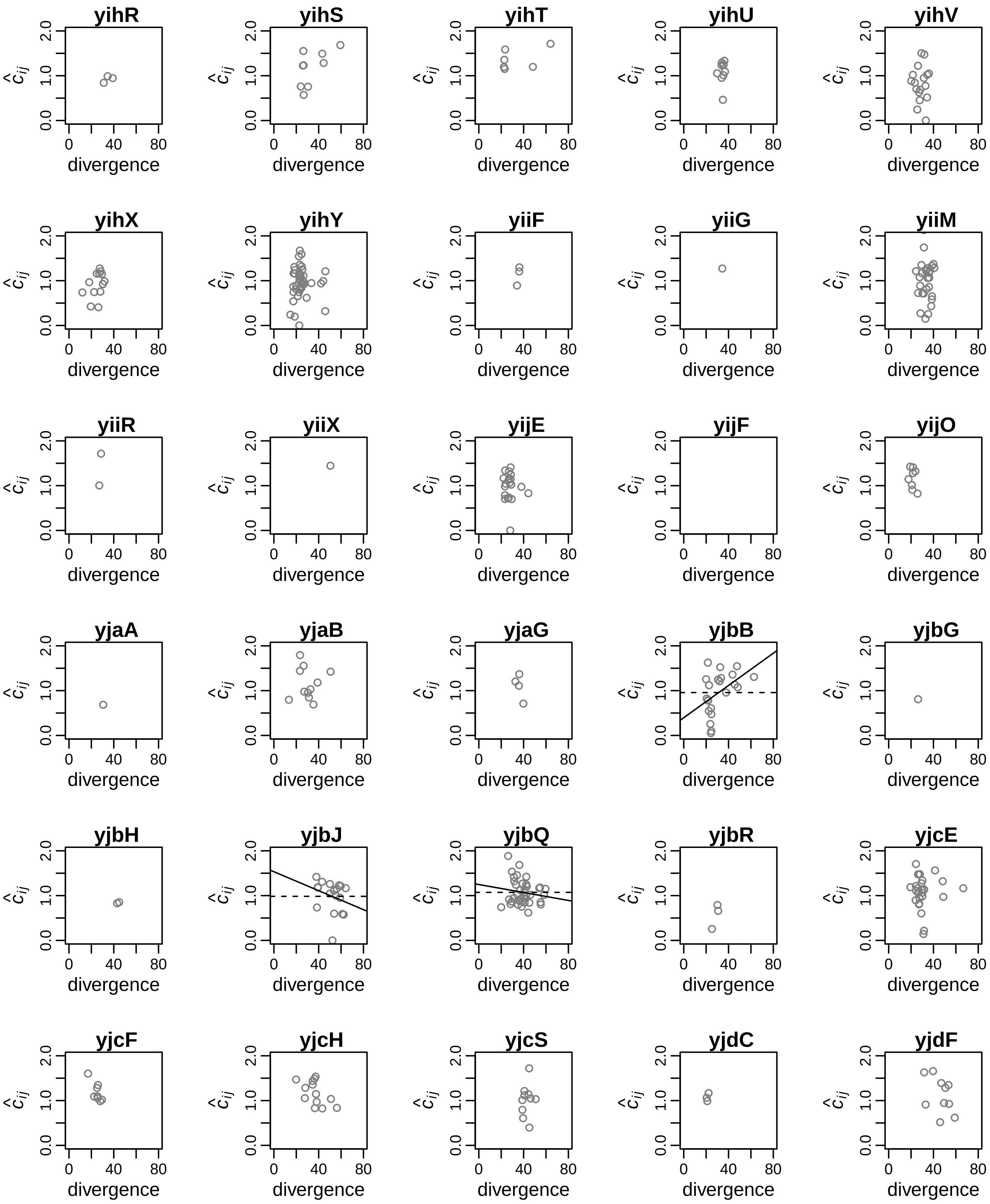
yihR
yihS
yihT
yihU
yihV
yihX
yihY
yiiF
yiiG
yiiM
yiiR
yiiX
yijE
yijF
yijO
yjaA
yjaB
yjaG
yjbB
yjbG
yjbH
yjbJ
yjbQ
yjbR
yjcE
yjcF
yjcH
yjcS
yjdC
yjdF
$\hat{c}_{ij}$
0.0 1.0 2.0
0 40 80
divergence

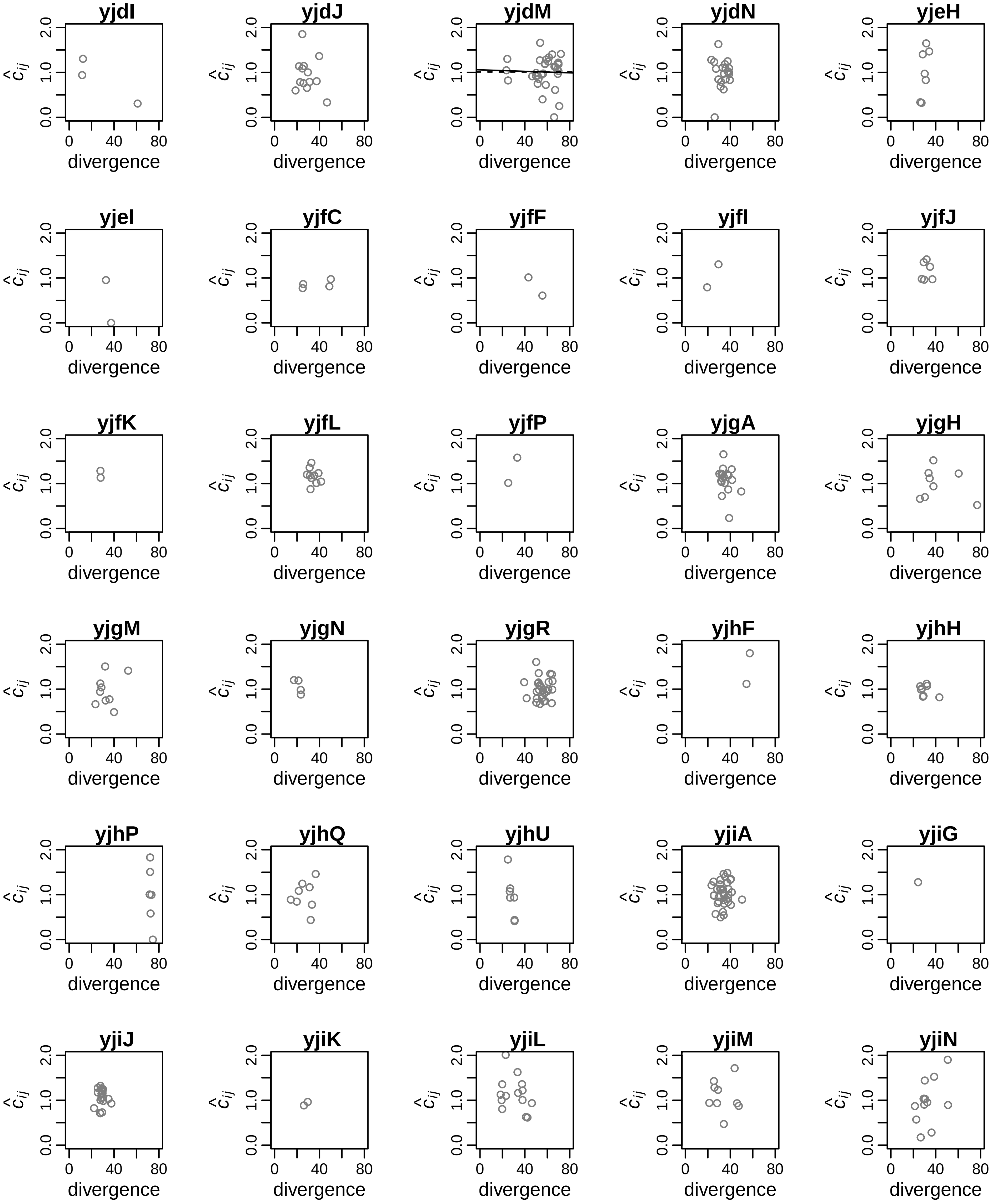
yjdI
yjdJ
yjdM
yjdN
yjeH
yjeI
yjfC
yjfF
yjfI
yjfJ
yjfK
yjfL
yjfP
yjgA
yjgH
yjgM
yjgN
yjgR
yjhF
yjhH
yjhP
yjhQ
yjhU
yjiA
yjiG
yjiJ
yjiK
yjiL
yjiM
yjiN
$\hat{c}_{ij}$
divergence
0.0
1.0
2.0
0
40
80

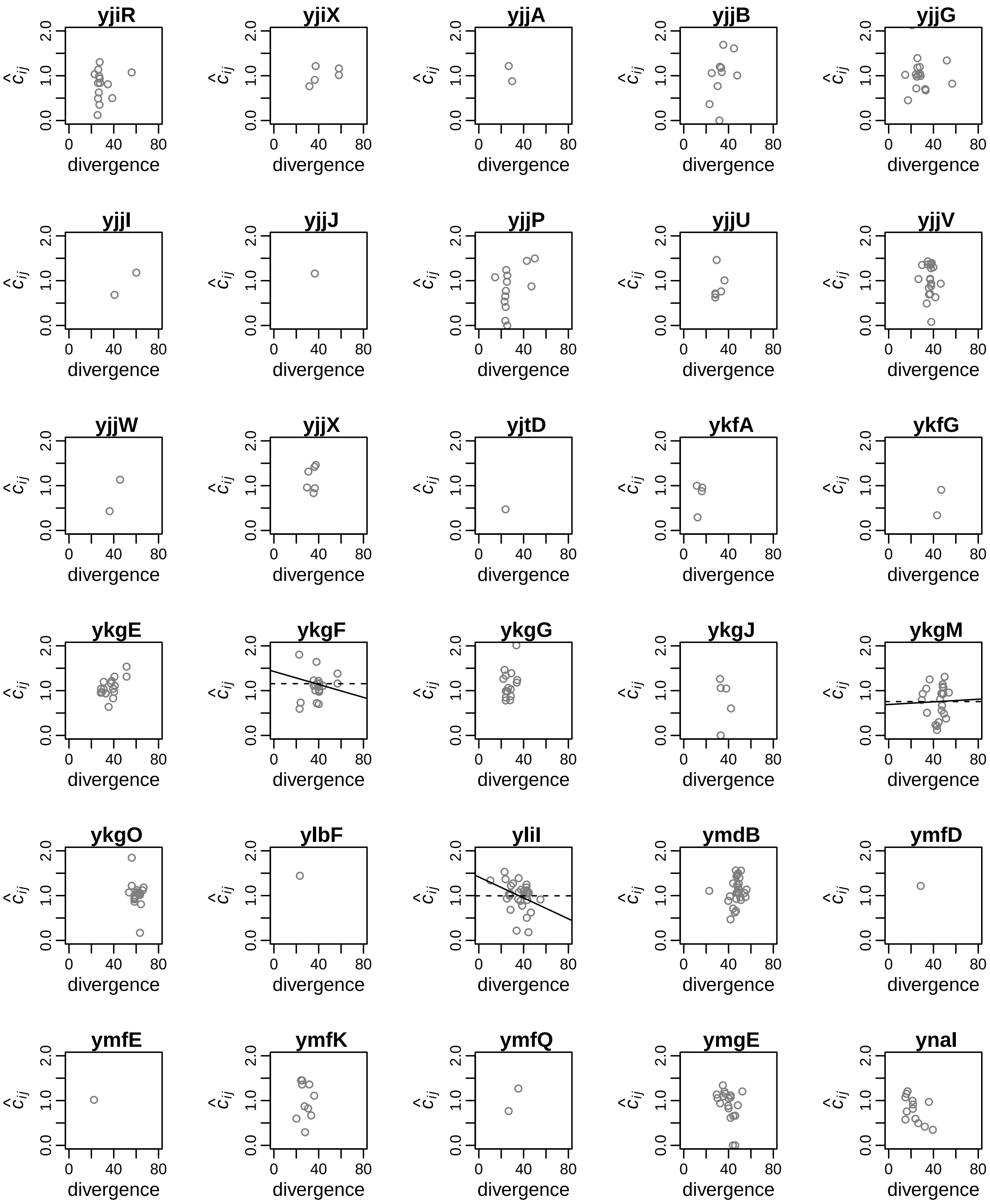
yjiR
yjiX
yjjA
yjjB
yjjG
yjjI
yjjJ
yjjP
yjjU
yjjV
yjjW
yjjX
yjtD
ykfA
ykfG
ykgE
ykgF
ykgG
ykgJ
ykgM
ykgO
ylbF
yliI
ymdB
ymfD
ymfE
ymfK
ymfQ
ymgE
ynaI
$\hat{c}_{ij}$
0.0 1.0 2.0
0 40 80
divergence

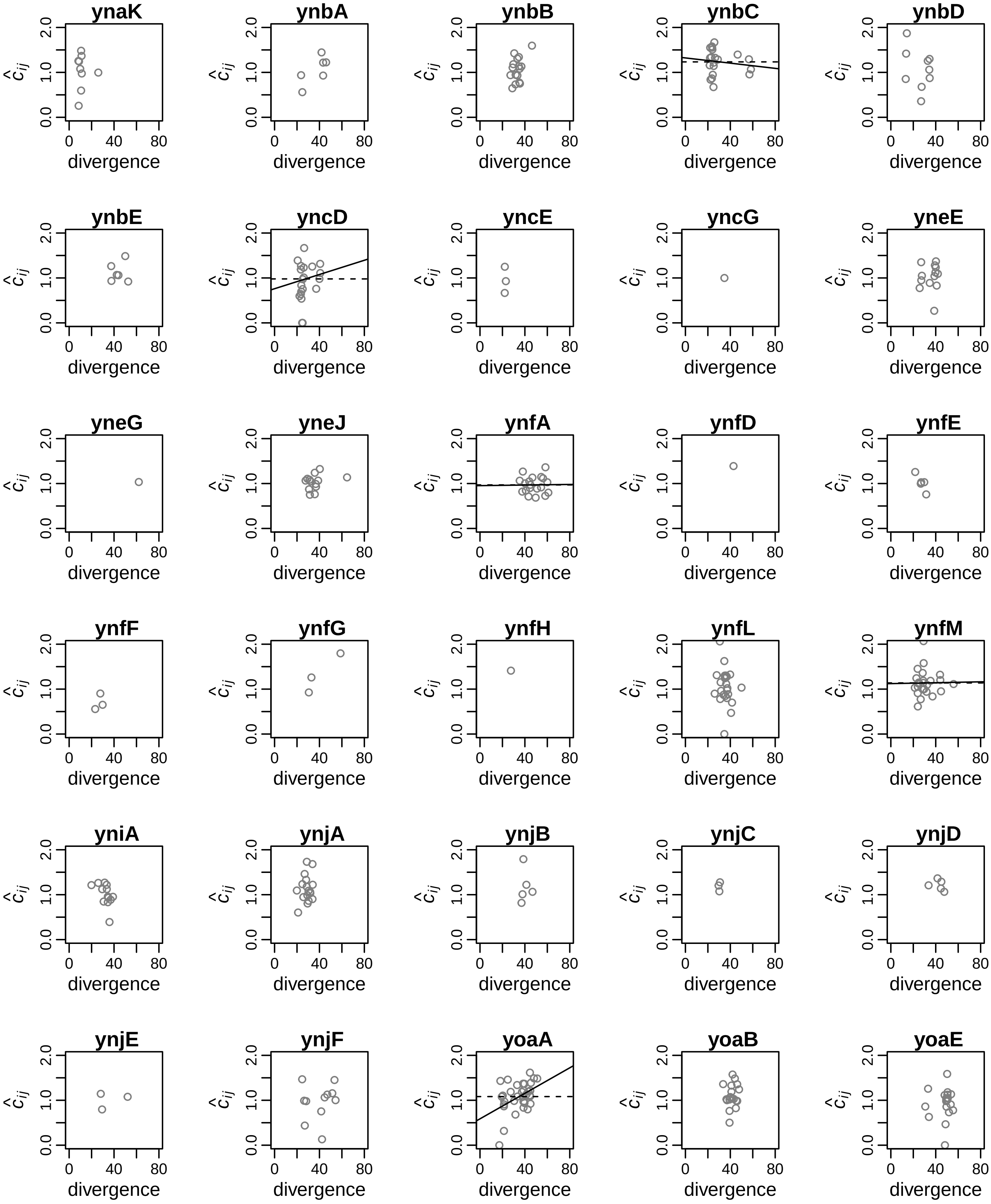
ynaK
ynbA
ynbB
ynbC
ynbD
ynbE
yncD
yncE
yncG
yneE
yneG
yneJ
ynfA
ynfD
ynfE
ynfF
ynfG
ynfH
ynfL
ynfM
yniA
ynjA
ynjB
ynjC
ynjD
ynjE
ynjF
yoaA
yoaB
yoaE
$\hat{c}_{ij}$
0.0
1.0
2.0
0
40
80
divergence

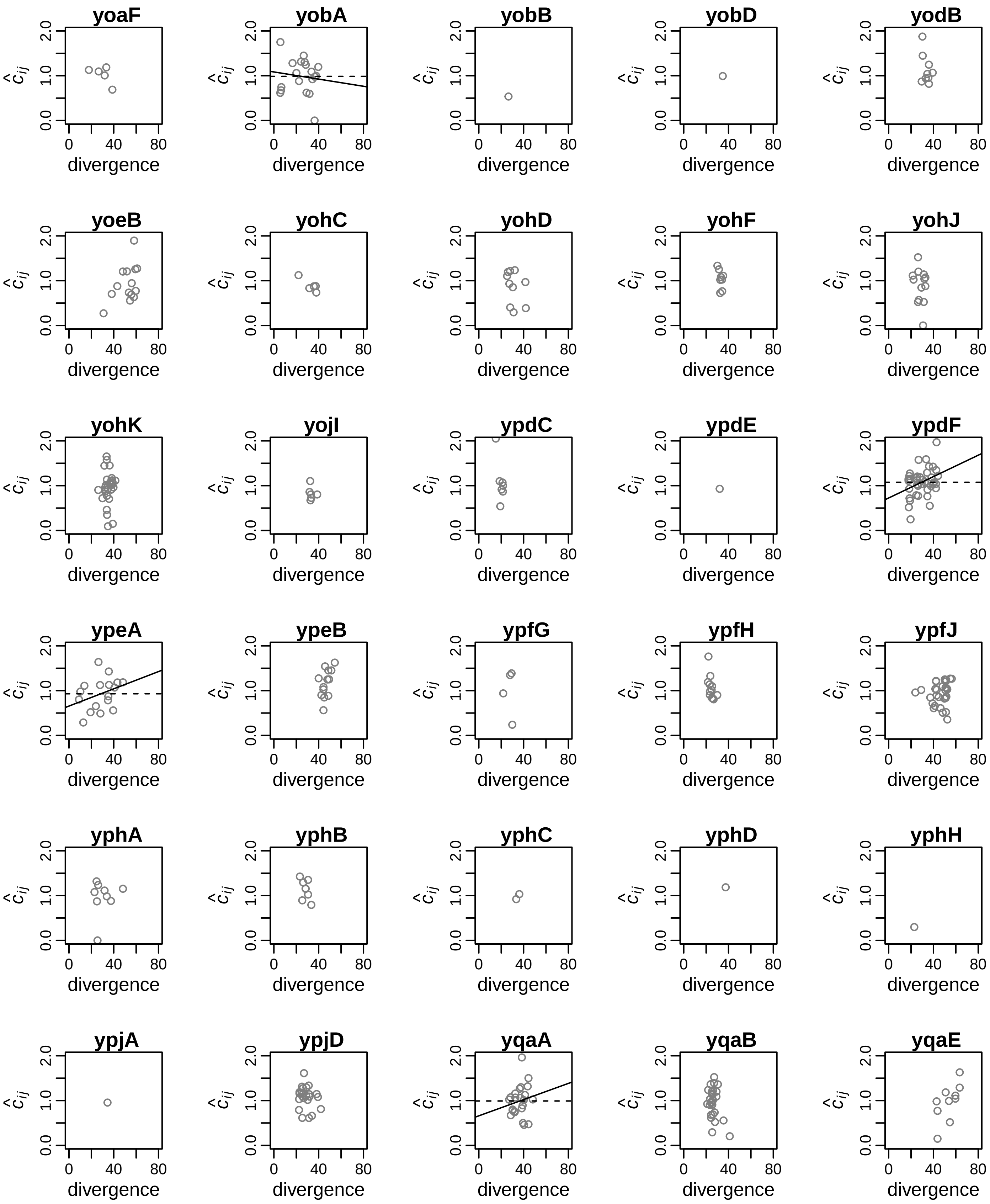
yoaF
yobA
yobB
yobD
yodB
yoeB
yohC
yohD
yohF
yohJ
yohK
yojI
ypdC
ypdE
ypdF
ypeA
ypeB
ypfG
ypfH
ypfJ
yphA
yphB
yphC
yphD
yphH
ypjA
ypjD
yqaA
yqaB
yqaE
$\hat{c}_{ij}$
0.0
1.0
2.0
0
40
80
divergence

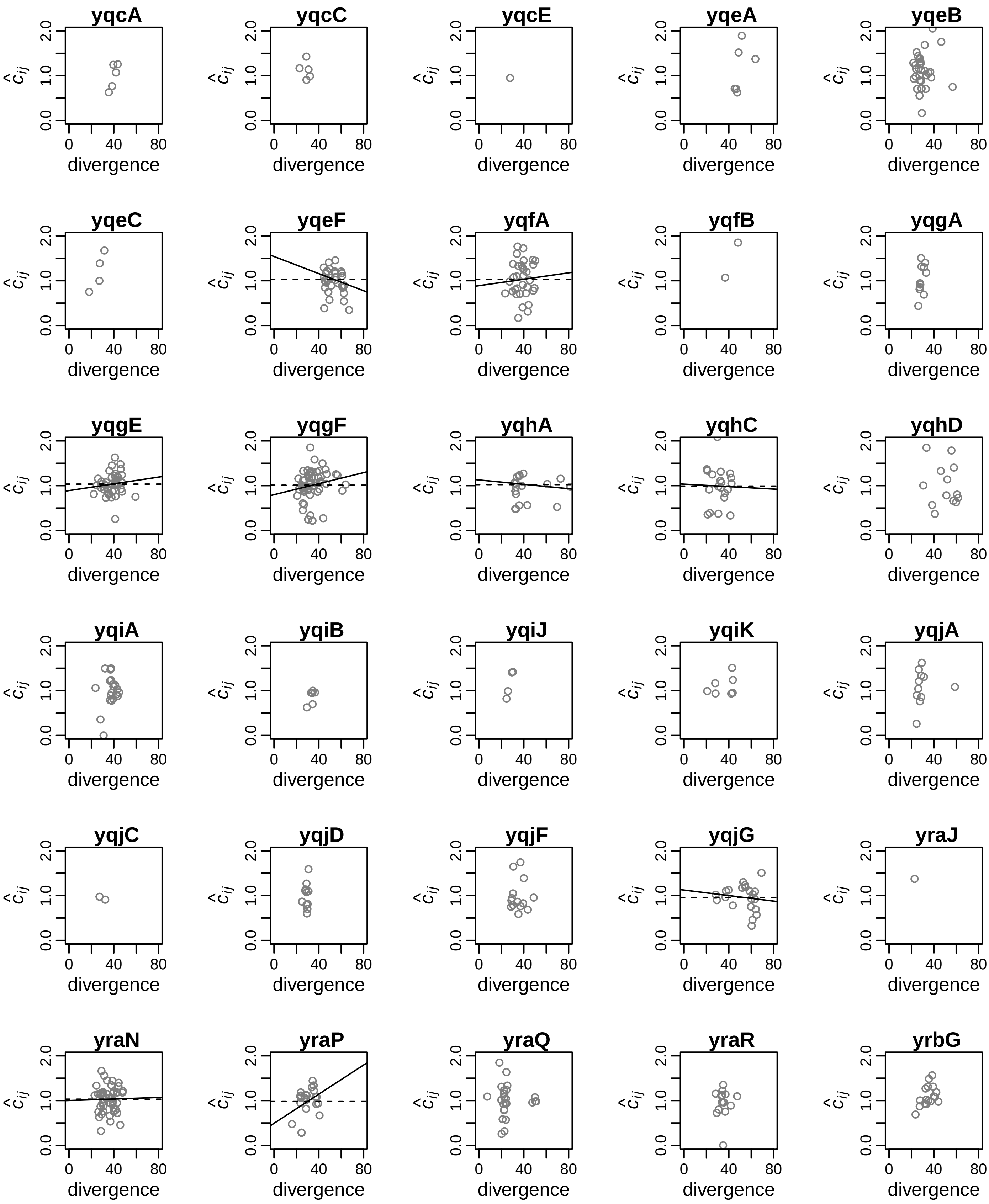
yqcA
yqcC
yqcE
yqeA
yqeB
yqeC
yqeF
yqfA
yqfB
yqgA
yqgE
yqgF
yqhA
yqhC
yqhD
yqiA
yqiB
yqiJ
yqiK
yqjA
yqjC
yqjD
yqjF
yqjG
yraJ
yraN
yraP
yraQ
yraR
yrbG
$\hat{c}_{ij}$
0.0
1.0
2.0
0
40
80
divergence

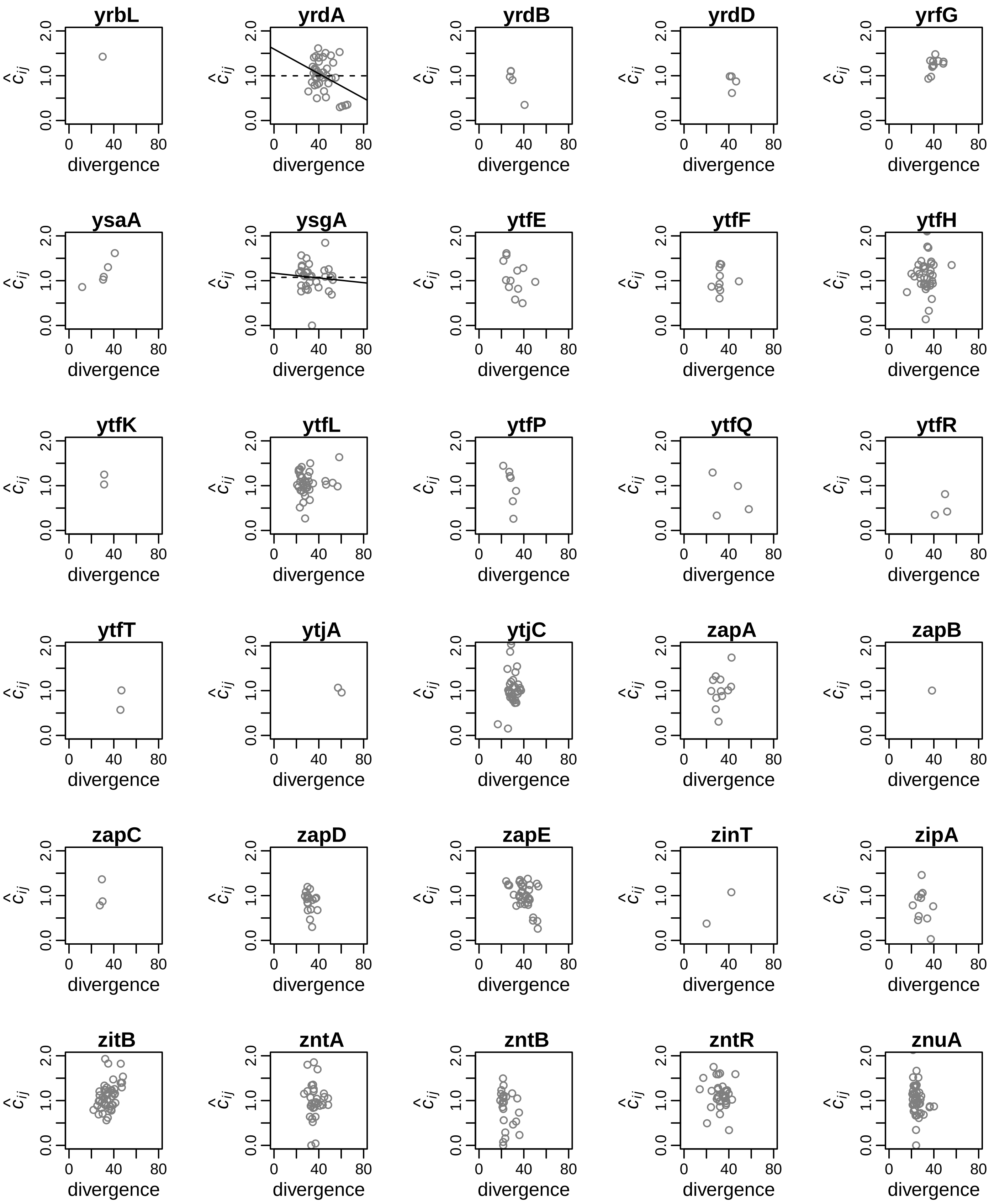
yrbL
yrdA
yrdB
yrdD
yrfG
ysaA
ysgA
ytfE
ytfF
ytfH
ytfK
ytfL
ytfP
ytfQ
ytfR
ytfT
ytjA
ytjC
zapA
zapB
zapC
zapD
zapE
zinT
zipA
zitB
zntA
zntB
zntR
znuA
$\hat{c}_{ij}$
0.0
1.0
2.0
0
40
80
divergence

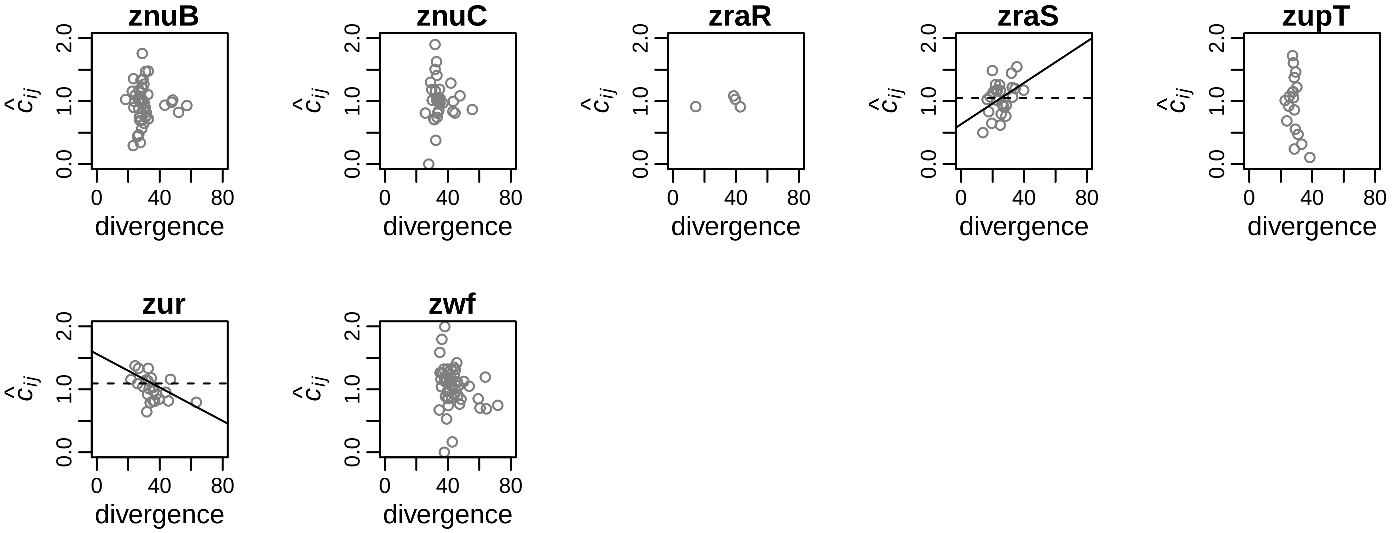

znuB
znuC
zraR
zraS
zupT
zur
zwf
$\hat{c}_{ij}$
divergence
0.0
1.0
2.0
0
40
80

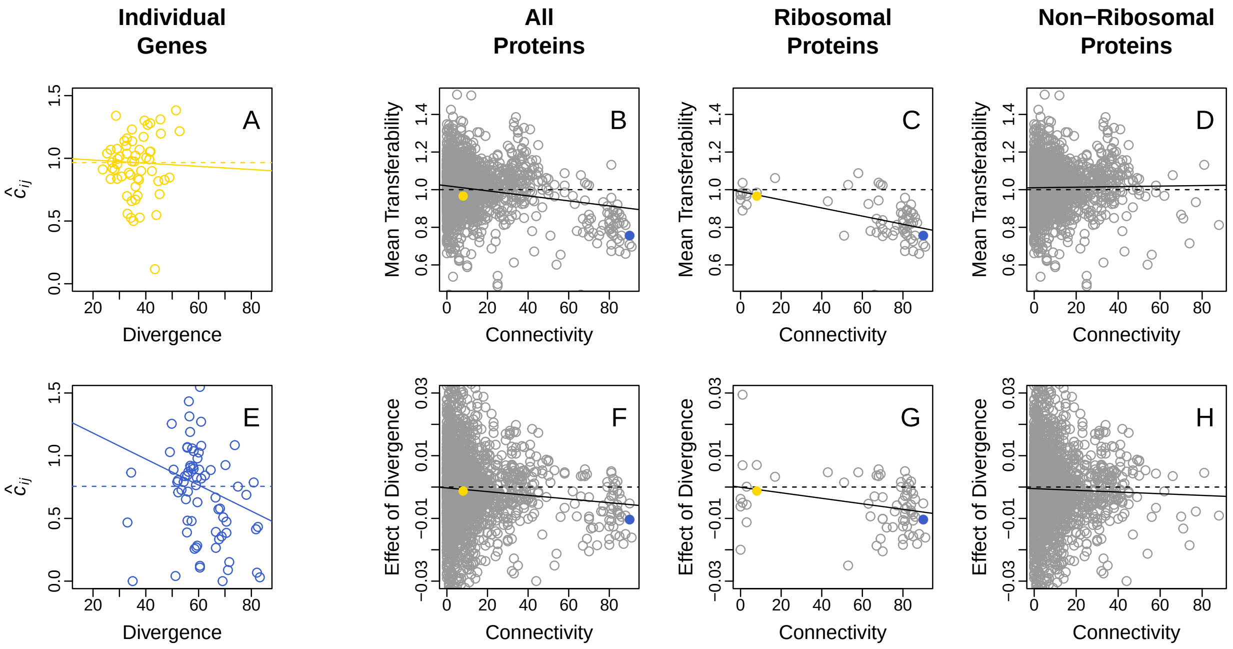


**Figure S5. Connectivity and divergence effects on transferability in a less-stringent analysis of 2420 sets of orthologous genes.** All panels are as described in fig. 3, except that more genes were included in the analysis. In the fig. 3 analysis, each set of orthologous genes was required to include at least 16 orthologs and to exhibit a standard deviation greater than 12% amino acid difference. By contrast, in this analysis each set of orthologous genes was required to include a minimum of only 5 orthologs and a standard deviation greater than 5% amino acid difference. 2420 orthologous gene sets met these less-stringent criteria. As in fig. 3, in this larger gene set we found a significant negative relationship between mean transferability and connectivity among the complete collection of 2420 protein-coding genes (panel B; estimate = -0.0013, $F_{1,2418} = 69.04$, $p = 2.2 \times 10^{-16}$) and among the 68 ribosomal genes (panel C; estimate = -0.0022, $F_{1,66} = 27.39$, $p = 1.8 \times 10^{-6}$), but not among the 2352 non-ribosomal genes (panel D; estimate = 0.0001, $F_{1,1045} = 0.41$, $p = 0.5231$). We also found a negative divergence $\times$ connectivity interaction, that was significant among the complete collection of 2420 genes (panel F; estimate = $-5.9 \times 10^{-5}$,

$F_{1,2418} = 6.134$, $p = 0.0133$) and the 68 ribosomal genes (panel G; estimate $= -8.8 \times 10^{-5}$, $F_{1,66} = 6.047$, $p = 0.01656$), but not the 2352 non-ribosomal genes (panel H; estimate $= -2.7 \times 10^{-5}$, $F_{1,2350} = 0.614$, $p = 0.4336$).
